## Supplementary table 7 for "DAP12 deficiency alters microglia-oligodendrocyte communication and enhances resilience against tau toxicity"

| Figure7E_GSEA_correlation_between_mouse _oligodendrocytes_and_human_oligodendrocyte_population | | | | | | | | | | | |
|---|---|---|---|---|---|---|---|---|---|---|---|
| Cluster_num | NAME | GS<br> follow link to MSigDB | SIZE | ES | NES | NOM p-val | FDR q-val | FWER p-val | RANK AT MAX | LEADING EDGE | 0 equals infinity value stands for high significance |
| mOL1 | HOL3 | HOL3 | 70 | -0.13202061 | -0.91086036 | 0.59398496 | 1 | 0.291 | 657 | tags=93%, list=73%, signal=319% | |
| mOL1 | HOL4 | HOL4 | 153 | -0.09869152 | -0.8215476 | 0.7592593 | 0.7113475 | 0.351 | 276 | tags=40%, list=31%, signal=48% | |
| mOL1 | HOL2 | HOL2 | 133 | 0.47467968 | 2.6209013 | 0 | 0 | 0 | 211 | tags=49%, list=23%, signal=54% | |
| mOL1 | HOL1 | HOL1 | 311 | 0.17749754 | 1.0811751 | 0.35569105 | 0.5471752 | 0.855 | 237 | tags=29%, list=26%, signal=26% | |
| mOL1 | HOL5 | HOL5 | 14 | 0.27046114 | 0.88939786 | 0.60678923 | 0.64911574 | 0.989 | 10 | tags=7%, list=1%, signal=7% | |
| mOL4 | HOL1 | HOL1 | 1 | 1 | 1.1913265 | 0 | 0.50343406 | 0.249 | 0 | tags=100%, list=0%, signal=75% | |
| mOL4 | HOL5 | HOL5 | 1 | -1 | -1.2064409 | 0 | 0.47326204 | 0.251 | 2 | tags=100%, list=50%, signal=150% | |
| mOL2 | HOL2 | HOL2 | 49 | -0.7189196 | -1.9252855 | 0 | 0 | 0 | 54 | tags=47%, list=19%, signal=48% | |
| mOL2 | HOL3 | HOL3 | 20 | -0.4904893 | -1.081384 | 0.3503521 | 0.6803107 | 0.633 | 56 | tags=30%, list=20%, signal=35% | |
| mOL2 | HOL4 | HOL4 | 26 | -0.38885194 | -0.92468286 | 0.58434784 | 0.8124278 | 0.873 | 47 | tags=15%, list=17%, signal=17% | |
| mOL2 | HOL5 | HOL5 | 1 | -0.5017794 | -0.6614262 | 0.9958159 | 0.95744735 | 0.982 | 142 | tags=100%, list=50%, signal=201% | |
| mOL2 | HOL1 | HOL1 | 117 | 0.3286479 | 1.0364422 | 0.3840399 | 0.4087957 | 0.642 | 42 | tags=22%, list=15%, signal=15% | |
| mOL5 | HOL1 | HOL1 | 307 | -0.23475203 | --- | --- | 1 | 0 | 151 | tags=18%, list=10%, signal=16% | |
| mOL5 | HOL3 | HOL3 | 124 | 0.5348707 | 1.7200477 | 0.041067763 | 0.009600001 | 0.012 | 97 | tags=12%, list=6%, signal=12% | |
| mOL5 | HOL5 | HOL5 | 28 | 0.3765294 | 1.0238177 | 0.4533947 | 0.8941662 | 0.937 | 201 | tags=18%, list=13%, signal=20% | |
| mOL5 | HOL4 | HOL4 | 326 | 0.2658045 | 0.8767577 | 0.779 | 0.9619296 | 0.998 | 513 | tags=32%, list=32%, signal=38% | |
| mOL5 | HOL2 | HOL2 | 182 | 0.2323642 | 0.7459075 | 0.8973843 | 0.8895969 | 1 | 395 | tags=26%, list=25%, signal=31% | |
| mOL3 | HOL1 | HOL1 | 271 | -0.16439 | -1.4537516 | 0.033264033 | 0.16185829 | 0.129 | 421 | tags=62%, list=57%, signal=91% | |
| mOL3 | HOL5 | HOL5 | 12 | -0.21741582 | -0.7755205 | 0.7166362 | 1 | 0.936 | 73 | tags=25%, list=10%, signal=27% | |
| mOL3 | HOL2 | HOL2 | 111 | -0.0929599 | -0.6934509 | 0.9129594 | 0.8953632 | 0.96 | 364 | tags=55%, list=49%, signal=92% | |
| mOL3 | HOL3 | HOL3 | 67 | 0.404484 | 2.6498325 | 0 | 0 | 0 | 173 | tags=45%, list=23%, signal=53% | |
| mOL3 | HOL4 | HOL4 | 112 | 0.18656817 | 1.4028509 | 0.07966457 | 0.077818364 | 0.174 | 153 | tags=34%, list=21%, signal=36% | |

| Figure7H_spatial_transcriptome_analysis_iOli_markergenes | | | | | | | | | |
|---|---|---|---|---|---|---|---|---|---|
| | group | category | mean | sd | p | p.adj | p.signif | method | comparison |
| 41 | annotation_sixgroups_geneset3_OLIG_nonoise_nolevel4 | AT8 | 0.3 | 0.12201346 | 1.18E-21 | 1.20E-21 | **** | Wilcoxon | vs non_AT8_level1 |
| 42 | annotation_sixgroups_geneset3_OLIG_nonoise_nolevel4 | non_AT8_level1 | 0.2 | 0.127667509 | 1.18E-21 | 1.20E-21 | **** | Wilcoxon | vs non_AT8_level1 |
| 43 | annotation_sixgroups_geneset3_OLIG_nonoise_nolevel4 | AT8 | 0.3 | 0.12201346 | 0.001996553 | 0.002 | ** | Wilcoxon | vs non_AT8_level2 |
| 44 | annotation_sixgroups_geneset3_OLIG_nonoise_nolevel4 | non_AT8_level2 | 0.2 | 0.141827529 | 0.001996553 | 0.002 | ** | Wilcoxon | vs non_AT8_level2 |
| 45 | annotation_sixgroups_geneset3_OLIG_nonoise_nolevel4 | AT8 | 0.3 | 0.12201346 | 1.08E-14 | 1.10E-14 | **** | Wilcoxon | vs non_AT8_level3 |
| 46 | annotation_sixgroups_geneset3_OLIG_nonoise_nolevel4 | non_AT8_level3 | 0.2 | 0.142913154 | 1.08E-14 | 1.10E-14 | **** | Wilcoxon | vs non_AT8_level3 |
| 47 | annotation_sixgroups_geneset3_OLIG_nonoise_nolevel4 | AT8 | 0.3 | 0.12201346 | 8.65E-168 | 8.60E-168 | **** | Wilcoxon | AT8 vs control |
| 48 | annotation_sixgroups_geneset3_OLIG_nonoise_nolevel4 | control | 0.2 | 0.13388826 | 8.65E-168 | 8.60E-168 | **** | Wilcoxon | AT8 vs control |
| 49 | annotation_sixgroups_geneset3_OLIG_nonoise_nolevel4 | non_AT8_level1 | 0.2 | 0.127667509 | 2.00E-07 | 2.00E-07 | **** | Wilcoxon | level1 vs non_AT8_level2 |
| 50 | annotation_sixgroups_geneset3_OLIG_nonoise_nolevel4 | non_AT8_level2 | 0.2 | 0.141827529 | 2.00E-07 | 2.00E-07 | **** | Wilcoxon | level1 vs non_AT8_level2 |
| 51 | annotation_sixgroups_geneset3_OLIG_nonoise_nolevel4 | non_AT8_level1 | 0.2 | 0.127667509 | 0.327602909 | 0.33 | ns | Wilcoxon | level1 vs non_AT8_level3 |
| 52 | annotation_sixgroups_geneset3_OLIG_nonoise_nolevel4 | non_AT8_level3 | 0.2 | 0.142913154 | 0.327602909 | 0.33 | ns | Wilcoxon | level1 vs non_AT8_level3 |
| 53 | annotation_sixgroups_geneset3_OLIG_nonoise_nolevel4 | non_AT8_level1 | 0.2 | 0.127667509 | 3.28E-93 | 3.30E-93 | **** | Wilcoxon | AT8_level1 vs control |
| 54 | annotation_sixgroups_geneset3_OLIG_nonoise_nolevel4 | control | 0.2 | 0.13388826 | 3.28E-93 | 3.30E-93 | **** | Wilcoxon | AT8_level1 vs control |
| 55 | annotation_sixgroups_geneset3_OLIG_nonoise_nolevel4 | non_AT8_level2 | 0.2 | 0.141827529 | 3.04E-06 | 3.00E-06 | **** | Wilcoxon | level2 vs non_AT8_level3 |
| 56 | annotation_sixgroups_geneset3_OLIG_nonoise_nolevel4 | non_AT8_level3 | 0.2 | 0.142913154 | 3.04E-06 | 3.00E-06 | **** | Wilcoxon | level2 vs non_AT8_level3 |
| 57 | annotation_sixgroups_geneset3_OLIG_nonoise_nolevel4 | non_AT8_level2 | 0.2 | 0.141827529 | 1.06E-96 | 1.10E-96 | **** | Wilcoxon | AT8_level2 vs control |
| 58 | annotation_sixgroups_geneset3_OLIG_nonoise_nolevel4 | control | 0.2 | 0.13388826 | 1.06E-96 | 1.10E-96 | **** | Wilcoxon | AT8_level2 vs control |
| 59 | annotation_sixgroups_geneset3_OLIG_nonoise_nolevel4 | non_AT8_level3 | 0.2 | 0.142913154 | 3.77E-30 | 3.80E-30 | **** | Wilcoxon | AT8_level3 vs control |
| 60 | annotation_sixgroups_geneset3_OLIG_nonoise_nolevel4 | control | 0.2 | 0.13388826 | 3.77E-30 | 3.80E-30 | **** | Wilcoxon | AT8_level3 vs control |

| Figure7IJ_Oigo2_Mef2c_staining_human_brain-tissue | |
|---|---|
| cell numebrs/mm2 | |
| NON-AD | AD |
| 13 | 6 |
| 8 | 8 |
| 10 | 7 |
| 2 | 13 |
| 7 | 13 |
| 8 | 6 |
| 4 | 2 |
| 7 | 4 |
| 4 | 4 |
| 4 | 5 |
| 9 | 15 |
| 3 | 15 |
| 6 | 8 |
| 9 | 10 |
| 9 | 11 |
| 14 | 8 |
| 3 | 8 |
| 7 | 7 |
| 3 | 6 |
| 7 | 9 |
| 5 | 8 |
| 10 | 16 |
| 6 | 20 |
| 11 | 9 |
| 6 | 13 |

| Figure7D_human oligodendrocyte cluster3 DEGs versus others | | | | | | |
|---|---|---|---|---|---|---|
| | p_val | avg_log2FC | pct.1 | pct.2 | p_val_adj | 22895729E-300 to infinity value |
| NRXN1 | 0 | 2.854889625 | 0.7 | 0.212 | 5.57E-300 | |
| ATP2B2 | 0 | 1.473356801 | 0.2 | 0.026 | 5.57E-300 | |
| GPM6A | 0 | 2.827126739 | 0.6 | 0.141 | 5.57E-300 | |
| ADCY2 | 0 | 1.9000949 | 0.3 | 0.06 | 5.57E-300 | |
| TENM2 | 0 | 1.962174525 | 0.4 | 0.068 | 5.57E-300 | |
| NRCAM | 0 | 2.012063285 | 0.4 | 0.066 | 5.57E-300 | |
| SLC1A2 | 0 | 2.663619282 | 0.4 | 0.11 | 5.57E-300 | |
| CELF2 | 0 | 1.859229663 | 0.5 | 0.098 | 5.57E-300 | |
| NRG3 | 0 | 3.312136762 | 0.7 | 0.155 | 5.57E-300 | |
| SYT1 | 0 | 3.024120605 | 0.6 | 0.177 | 5.57E-300 | |
| FAM155A | 0 | 2.584896087 | 0.6 | 0.121 | 5.57E-300 | |
| DLGAP1 | 0 | 2.293016122 | 0.5 | 0.122 | 5.57E-300 | |
| PLCB1 | 0 | 2.173056284 | 0.5 | 0.11 | 5.57E-300 | |
| ADGRV1 | 1.93E-293 | 2.378482414 | 0.4 | 0.072 | 5.57E-289 | |
| GRIN2B | 3.28E-285 | 1.409840174 | 0.2 | 0.027 | 9.49E-281 | |
| SNTG1 | 4.38E-281 | 1.790399159 | 0.4 | 0.074 | 1.27E-276 | |
| FGF14 | 1.29E-276 | 2.088234281 | 0.5 | 0.136 | 3.74E-272 | |
| RYR3 | 1.31E-268 | 1.948479597 | 0.3 | 0.045 | 3.78E-264 | |
| KCNQ5 | 2.35E-267 | 2.102223214 | 0.4 | 0.076 | 6.78E-263 | |
| OPCML | 1.89E-259 | 2.288171484 | 0.4 | 0.121 | 5.45E-255 | |
| MYT1L | 5.03E-259 | 1.418492578 | 0.2 | 0.037 | 1.45E-254 | |
| PTPRZ1 | 3.55E-258 | 1.615921211 | 0.3 | 0.043 | 1.03E-253 | |
| CNTN1 | 6.28E-258 | 1.724588275 | 0.4 | 0.113 | 1.82E-253 | |
| KCNMA1 | 8.50E-257 | 1.093608113 | 0.4 | 0.08 | 2.46E-252 | |
| AGBL4 | 4.52E-254 | 1.737066291 | 0.3 | 0.072 | 1.31E-249 | |
| ADCY8 | 2.28E-251 | 0.724873717 | 0.1 | 0.008 | 6.58E-247 | |
| RALYL | 1.76E-250 | 2.03650524 | 0.3 | 0.072 | 5.09E-246 | |
| KCND2 | 3.84E-249 | 1.488738472 | 0.3 | 0.042 | 1.11E-244 | |
| FMN2 | 6.46E-243 | 1.942006727 | 0.4 | 0.089 | 1.87E-238 | |
| MGAT4C | 1.36E-242 | 1.389010924 | 0.3 | 0.041 | 3.92E-238 | |
| GABRB2 | 2.75E-241 | 1.655146209 | 0.3 | 0.043 | 7.95E-237 | |
| CHL1 | 5.32E-241 | 1.511957705 | 0.2 | 0.037 | 1.54E-236 | |
| FRMPD4 | 6.77E-230 | 1.257374401 | 0.2 | 0.034 | 1.96E-225 | |
| NEBL | 6.81E-230 | 1.63484712 | 0.3 | 0.063 | 1.97E-225 | |
| RNF219-AS1 | 1.11E-229 | 1.222203121 | 0.2 | 0.022 | 3.21E-225 | |
| EPHA6 | 1.51E-228 | 1.345530158 | 0.2 | 0.039 | 4.38E-224 | |
| GLIS3 | 8.36E-228 | 1.576394933 | 0.2 | 0.039 | 2.42E-223 | |
| RBFOX1 | 1.64E-223 | 2.150284064 | 0.6 | 0.258 | 4.73E-219 | |
| GRIP1 | 1.73E-222 | 1.437729717 | 0.3 | 0.051 | 4.99E-218 | |
| STXBP5L | 2.36E-220 | 1.612665719 | 0.3 | 0.069 | 6.83E-216 | |
| KCNJ3 | 9.41E-219 | 1.10867111 | 0.2 | 0.018 | 2.72E-214 | |
| HS6ST3 | 1.45E-217 | 1.608089274 | 0.2 | 0.044 | 4.20E-213 | |
| CADPS | 3.60E-217 | 1.611928959 | 0.3 | 0.055 | 1.04E-212 | |
| LINC00299 | 3.23E-214 | 0.68038324 | 0.1 | 0.006 | 9.34E-210 | |
| DCLK1 | 3.97E-213 | 1.447990878 | 0.3 | 0.051 | 1.15E-208 | |
| CCSER1 | 2.79E-211 | 1.714048433 | 0.3 | 0.089 | 8.08E-207 | |
| SLC4A10 | 3.53E-210 | 1.436790088 | 0.2 | 0.043 | 1.02E-205 | |
| GRIA1 | 6.06E-210 | 1.05299528 | 0.2 | 0.023 | 1.75E-205 | |
| SOX5 | 3.41E-209 | 1.789583471 | 0.4 | 0.127 | 9.87E-205 | |
| GRM5 | 5.25E-208 | 1.619246542 | 0.3 | 0.058 | 1.52E-203 | |

| | | | | | |
|---|---|---|---|---|---|
| RFX4 | 2.49E-207 | 1.24272696 | 0.2 | 0.019 | 7.20E-203 |
| EFNA5 | 6.68E-205 | 1.212189546 | 0.2 | 0.035 | 1.93E-200 |
| MEF2C | 4.41E-204 | 1.705015754 | 0.3 | 0.081 | 1.28E-199 |
| SYN2 | 7.76E-203 | 1.303324735 | 0.2 | 0.031 | 2.24E-198 |
| FGF12 | 4.94E-201 | 1.663337298 | 0.3 | 0.089 | 1.43E-196 |
| ASIC2 | 8.88E-200 | 1.35175227 | 0.2 | 0.039 | 2.57E-195 |
| GABRB1 | 7.95E-199 | 1.665241763 | 0.3 | 0.078 | 2.30E-194 |
| NELL2 | 4.46E-198 | 1.485990438 | 0.2 | 0.042 | 1.29E-193 |
| ROBO2 | 1.08E-197 | 1.984483372 | 0.4 | 0.111 | 3.11E-193 |
| GABBR2 | 2.27E-197 | 1.224959361 | 0.2 | 0.027 | 6.56E-193 |
| SCN2A | 3.90E-195 | 1.597811057 | 0.2 | 0.039 | 1.13E-190 |
| MAPK4 | 1.60E-192 | 1.018683821 | 0.1 | 0.012 | 4.64E-188 |
| CSGALNACT1 | 3.28E-192 | 1.13143695 | 0.2 | 0.028 | 9.49E-188 |
| MED12L | 1.80E-191 | 1.309880037 | 0.2 | 0.039 | 5.20E-187 |
| ATRNL1 | 7.22E-191 | 1.84442603 | 0.3 | 0.089 | 2.09E-186 |
| NKAIN3 | 3.52E-186 | 1.322674375 | 0.2 | 0.029 | 1.02E-181 |
| TRPM3 | 5.12E-186 | 1.482917155 | 0.2 | 0.051 | 1.48E-181 |
| RIMS2 | 6.46E-184 | 1.648249608 | 0.3 | 0.083 | 1.87E-179 |
| KCNIP4 | 7.78E-184 | 2.276755189 | 0.6 | 0.326 | 2.25E-179 |
| DLGAP2 | 1.72E-182 | 1.436063258 | 0.3 | 0.054 | 4.97E-178 |
| RGS20 | 3.56E-182 | 0.888770722 | 0.1 | 0.011 | 1.03E-177 |
| SPARCL1 | 2.86E-180 | 2.123391846 | 0.4 | 0.158 | 8.27E-176 |
| THRB | 7.52E-180 | 1.251926115 | 0.3 | 0.059 | 2.17E-175 |
| LMO3 | 3.55E-175 | 1.053609147 | 0.2 | 0.023 | 1.03E-170 |
| CNKSR2 | 6.08E-175 | 1.154913662 | 0.2 | 0.036 | 1.76E-170 |
| CAMK2B | 7.96E-174 | 1.176447496 | 0.2 | 0.03 | 2.30E-169 |
| DPP10 | 3.26E-173 | 2.579233145 | 0.6 | 0.296 | 9.42E-169 |
| SLC4A4 | 1.52E-172 | 0.595621561 | 0.2 | 0.033 | 4.39E-168 |
| CNTNAP5 | 3.22E-172 | 1.385654889 | 0.2 | 0.05 | 9.30E-168 |
| CARMIL1 | 1.57E-171 | 1.137660814 | 0.2 | 0.031 | 4.54E-167 |
| CTNNA2 | 2.95E-170 | 1.404567279 | 0.6 | 0.289 | 8.52E-166 |
| BRINP1 | 5.10E-170 | 0.832695928 | 0.1 | 0.016 | 1.47E-165 |
| FAT3 | 7.13E-169 | 1.065076394 | 0.2 | 0.028 | 2.06E-164 |
| CACNB2 | 7.27E-168 | 1.504853806 | 0.3 | 0.086 | 2.10E-163 |
| PCLO | 4.30E-167 | 1.500504205 | 0.3 | 0.081 | 1.24E-162 |
| PIP4K2A | 1.03E-166 | -0.797500053 | 0.7 | 0.967 | 2.99E-162 |
| CACNA2D3 | 2.78E-165 | 1.156788395 | 0.3 | 0.063 | 8.03E-161 |
| PITPNC1 | 4.83E-165 | 1.83544528 | 0.3 | 0.105 | 1.39E-160 |
| CDH10 | 1.37E-163 | 1.032451034 | 0.2 | 0.023 | 3.95E-159 |
| ABLIM1 | 5.81E-163 | 1.154639752 | 0.2 | 0.047 | 1.68E-158 |
| CACNA1B | 1.17E-162 | 1.239262117 | 0.2 | 0.034 | 3.38E-158 |
| SNAP25 | 3.03E-162 | 2.389158861 | 0.4 | 0.146 | 8.77E-158 |
| CABLES1 | 3.67E-161 | 0.994588249 | 0.1 | 0.018 | 1.06E-156 |
| GABRB3 | 2.22E-159 | 1.208768356 | 0.2 | 0.03 | 6.41E-155 |
| FLRT2 | 3.68E-159 | 0.933552172 | 0.2 | 0.035 | 1.06E-154 |
| TOX | 3.27E-158 | 1.070059691 | 0.2 | 0.028 | 9.46E-154 |
| SHROOM3 | 1.16E-157 | 0.827047326 | 0.1 | 0.014 | 3.36E-153 |
| LGI1 | 4.11E-157 | 0.881134823 | 0.1 | 0.016 | 1.19E-152 |
| SLC8A1 | 2.62E-156 | 1.754574845 | 0.3 | 0.073 | 7.58E-152 |
| DSCAM | 2.86E-155 | 1.497133895 | 0.3 | 0.072 | 8.26E-151 |
| HPSE2 | 5.89E-154 | 1.156119948 | 0.2 | 0.028 | 1.70E-149 |
| PCDH7 | 1.64E-152 | 1.495629939 | 0.3 | 0.097 | 4.75E-148 |
| DNAH7 | 4.21E-151 | 0.95758625 | 0.1 | 0.018 | 1.22E-146 |
| CSMD1 | 4.81E-151 | 2.246126136 | 0.7 | 0.427 | 1.39E-146 |
| GPC5 | 5.73E-151 | 1.937563057 | 0.4 | 0.163 | 1.66E-146 |
| KCNC2 | 4.28E-149 | 1.180693323 | 0.2 | 0.03 | 1.24E-144 |
| PTCHD1-AS | 2.07E-148 | 0.888507623 | 0.2 | 0.025 | 5.97E-144 |
| ST6GALNAC5 | 5.09E-147 | 0.917265108 | 0.2 | 0.026 | 1.47E-142 |
| LRFN5 | 1.70E-145 | 1.194082881 | 0.2 | 0.046 | 4.92E-141 |
| PRKG1 | 2.51E-145 | 1.11423072 | 0.3 | 0.082 | 7.25E-141 |
| SCN1A | 2.92E-144 | 1.206688526 | 0.2 | 0.029 | 8.44E-140 |
| ELMO1 | 4.09E-142 | -0.742502317 | 0.7 | 0.958 | 1.18E-137 |
| GRIK2 | 1.36E-141 | 1.591036221 | 0.3 | 0.11 | 3.93E-137 |
| SLC35F1 | 2.76E-141 | 1.409481828 | 0.3 | 0.069 | 7.97E-137 |
| UNC5D | 4.25E-141 | 1.269424776 | 0.2 | 0.049 | 1.23E-136 |
| NTRK3 | 6.23E-140 | 1.034064999 | 0.3 | 0.071 | 1.80E-135 |
| BMPR1B | 6.42E-140 | 1.121466045 | 0.2 | 0.025 | 1.86E-135 |
| VSNL1 | 3.30E-139 | 1.917671698 | 0.2 | 0.046 | 9.55E-135 |
| MTUS2 | 4.83E-139 | 1.002071639 | 0.2 | 0.04 | 1.40E-134 |
| CAMK1D | 9.29E-139 | 1.213006388 | 0.2 | 0.058 | 2.68E-134 |
| PLP1 | 1.82E-138 | -0.906317798 | 0.7 | 0.926 | 5.25E-134 |
| KIAA1549L | 7.76E-138 | 0.986365683 | 0.1 | 0.02 | 2.24E-133 |
| TF | 1.05E-136 | -0.865648254 | 0.6 | 0.904 | 3.03E-132 |
| GRIN1 | 1.34E-136 | 1.545450995 | 0.2 | 0.063 | 3.88E-132 |
| C1orf21 | 2.67E-135 | 0.910171971 | 0.2 | 0.032 | 7.71E-131 |
| RNF220 | 5.33E-134 | -0.796452399 | 0.6 | 0.964 | 1.54E-129 |
| CDH18 | 5.47E-134 | 1.467576122 | 0.2 | 0.045 | 1.58E-129 |
| QKI | 5.59E-134 | -0.644809796 | 0.8 | 0.987 | 1.62E-129 |
| BRINP3 | 3.58E-133 | 0.804750212 | 0.1 | 0.025 | 1.03E-128 |
| GABRA1 | 7.70E-133 | 1.22716531 | 0.1 | 0.02 | 2.23E-128 |
| MBP | 4.60E-132 | -0.698805301 | 0.7 | 0.963 | 1.33E-127 |
| RBFOX3 | 1.34E-131 | 0.910491877 | 0.1 | 0.016 | 3.89E-127 |
| BASP1 | 3.14E-131 | 1.566186047 | 0.2 | 0.041 | 9.08E-127 |
| MEF2C-AS1 | 4.20E-130 | 0.920044516 | 0.2 | 0.034 | 1.21E-125 |
| MAP7 | 1.91E-129 | -0.711251308 | 0.6 | 0.943 | 5.52E-125 |
| ST18 | 9.36E-129 | -0.754162134 | 0.6 | 0.952 | 2.71E-124 |
| IL1RAPL1 | 1.03E-128 | -0.683722177 | 0.8 | 0.967 | 2.97E-124 |
| HCN1 | 1.07E-128 | 1.038329054 | 0.2 | 0.035 | 3.09E-124 |
| DGKG | 1.34E-128 | 1.360576407 | 0.2 | 0.05 | 3.88E-124 |
| OLFM3 | 7.65E-127 | 0.928988568 | 0.1 | 0.021 | 2.21E-122 |
| CBLN2 | 1.09E-125 | 0.886750836 | 0.1 | 0.016 | 3.16E-121 |
| DOCK5 | 2.95E-125 | -0.735467821 | 0.6 | 0.941 | 8.52E-121 |
| SH3GL2 | 3.84E-125 | 1.044009119 | 0.1 | 0.023 | 1.11E-120 |
| ZNF385D | 8.44E-125 | 1.293073856 | 0.2 | 0.043 | 2.44E-120 |
| DOCK10 | 3.46E-124 | -0.715240807 | 0.6 | 0.948 | 1.00E-119 |
| SRRM4 | 1.10E-123 | 0.726368411 | 0.1 | 0.015 | 3.18E-119 |
| KCNH8 | 6.73E-123 | -0.762379272 | 0.6 | 0.909 | 1.94E-118 |
| ENC1 | 1.86E-122 | 1.579626089 | 0.2 | 0.032 | 5.38E-118 |
| THSD7A | 2.58E-121 | 0.882288409 | 0.1 | 0.026 | 7.47E-117 |
| ADGRL2 | 5.93E-121 | 1.024485147 | 0.2 | 0.043 | 1.71E-116 |
| MAP3K5 | 1.01E-120 | 1.04138157 | 0.2 | 0.045 | 2.93E-116 |
| SLC44A1 | 1.54E-120 | -0.683777658 | 0.7 | 0.962 | 4.44E-116 |
| FRMD5 | 3.12E-120 | -0.706687038 | 0.7 | 0.954 | 9.01E-116 |
| CTNNA3 | 3.48E-120 | -0.698752855 | 0.7 | 0.957 | 1.01E-115 |
| MT-CO3 | 3.66E-120 | 1.266911351 | 0.9 | 0.871 | 1.06E-115 |
| CNTN5 | 1.32E-119 | 1.536969255 | 0.3 | 0.09 | 3.80E-115 |
| HECW1 | 1.83E-119 | 1.231699086 | 0.2 | 0.029 | 5.29E-115 |
| CDH9 | 2.70E-119 | 0.660648134 | 0.1 | 0.014 | 7.80E-115 |
| UGT8 | 3.57E-119 | -0.751662929 | 0.5 | 0.838 | 1.03E-114 |
| TMEM165 | 4.10E-118 | -0.677066542 | 0.6 | 0.948 | 1.19E-113 |
| PDE4B | 2.90E-116 | -0.629130583 | 0.7 | 0.966 | 8.39E-112 |
| PHLPP1 | 7.81E-116 | -0.677636398 | 0.7 | 0.958 | 2.26E-111 |
| PLCL1 | 9.12E-116 | -0.751057375 | 0.6 | 0.945 | 2.63E-111 |
| SYBU | 4.11E-115 | 0.847959547 | 0.1 | 0.019 | 1.19E-110 |
| FRMD4B | 4.43E-115 | -0.699098878 | 0.6 | 0.935 | 1.28E-110 |
| NRGN | 5.02E-115 | 1.94927562 | 0.2 | 0.061 | 1.45E-110 |
| SYNPR | 1.73E-114 | 0.787635987 | 0.1 | 0.022 | 4.99E-110 |
| C10orf90 | 1.77E-114 | -0.716595758 | 0.6 | 0.923 | 5.13E-110 |
| FAM107B | 2.89E-114 | -0.733429533 | 0.5 | 0.862 | 8.35E-110 |
| MOG | 3.31E-114 | -0.708283027 | 0.5 | 0.846 | 9.56E-110 |
| PRUNE2 | 4.08E-114 | -0.721851177 | 0.6 | 0.937 | 1.18E-109 |
| FAM189A2 | 6.35E-114 | 1.05329856 | 0.1 | 0.016 | 1.84E-109 |
| ELAVL4 | 8.41E-114 | 0.8245993 | 0.1 | 0.015 | 2.43E-109 |
| UNC5C | 9.65E-114 | -0.794502811 | 0.7 | 0.938 | 2.79E-109 |
| EDIL3 | 2.36E-113 | -0.660525865 | 0.7 | 0.955 | 6.82E-109 |
| TMEM144 | 2.80E-113 | -0.698124386 | 0.6 | 0.929 | 8.11E-109 |
| CNDP1 | 1.16E-112 | -0.906474658 | 0.5 | 0.796 | 3.35E-108 |
| SLAIN1 | 1.27E-112 | -0.75079867 | 0.6 | 0.914 | 3.68E-108 |
| DNM3 | 1.52E-111 | -0.615379808 | 0.7 | 0.961 | 4.39E-107 |
| PLEKHH1 | 2.11E-111 | -0.667552099 | 0.6 | 0.898 | 6.10E-107 |
| MAP2 | 2.30E-111 | 1.36055068 | 0.3 | 0.103 | 6.66E-107 |
| SHISA9 | 5.98E-111 | 0.809014649 | 0.1 | 0.018 | 1.73E-106 |
| BCAS1 | 7.85E-111 | -0.716606272 | 0.5 | 0.865 | 2.27E-106 |
| ERBIN | 3.68E-110 | -0.674299509 | 0.7 | 0.959 | 1.06E-105 |
| CELF4 | 3.83E-110 | 0.882456468 | 0.1 | 0.028 | 1.11E-105 |
| AC092691.1 | 5.19E-110 | 1.545545577 | 0.5 | 0.218 | 1.50E-105 |
| PLD1 | 1.01E-109 | -0.766451431 | 0.5 | 0.811 | 2.91E-105 |
| PDE8A | 1.49E-109 | -0.70733783 | 0.6 | 0.931 | 4.31E-105 |

| | | | | | |
|---|---|---|---|---|---|
| ABR | 1.71E-109 | 1.072150093 | 0.2 | 0.045 | 4.93E-105 |
| LRRTM4 | 3.24E-109 | 1.398786653 | 0.5 | 0.287 | 9.37E-105 |
| NCKAP5 | 5.92E-109 | -0.727682114 | 0.7 | 0.936 | 1.71E-104 |
| GALNTL6 | 9.65E-109 | 1.079788594 | 0.2 | 0.04 | 2.79E-104 |
| SHTN1 | 4.38E-108 | -0.674337916 | 0.6 | 0.936 | 1.27E-103 |
| ARHGAP26 | 5.68E-108 | 1.110080524 | 0.3 | 0.093 | 1.64E-103 |
| PDZRN3 | 6.63E-108 | 0.696106899 | 0.2 | 0.043 | 1.92E-103 |
| MAN2A1 | 1.76E-106 | -0.711476374 | 0.6 | 0.938 | 5.10E-102 |
| CABP1 | 2.40E-106 | 0.943672599 | 0.1 | 0.02 | 6.93E-102 |
| SIK3 | 3.53E-106 | -0.637013076 | 0.7 | 0.979 | 1.02E-101 |
| RORA | 4.07E-106 | 1.169399538 | 0.6 | 0.36 | 1.18E-101 |
| STMN2 | 5.24E-106 | 1.142851059 | 0.1 | 0.02 | 1.51E-101 |
| CAMK2A | 1.70E-105 | 1.53024911 | 0.2 | 0.049 | 4.91E-101 |
| PICALM | 3.78E-105 | -0.678685456 | 0.6 | 0.905 | 1.09E-100 |
| NETO1 | 1.44E-104 | 0.749583156 | 0.1 | 0.017 | 4.17E-100 |
| 1-Mar | 3.56E-104 | -0.661023088 | 0.7 | 0.935 | 1.03E-99 |
| RGS7 | 4.34E-104 | 1.256555711 | 0.3 | 0.134 | 1.25E-99 |
| DLG1 | 4.79E-104 | -0.685120448 | 0.6 | 0.945 | 1.38E-99 |
| SPOCK3 | 9.65E-104 | -0.690870522 | 0.6 | 0.93 | 2.79E-99 |
| PDE1C | 1.07E-103 | -0.688180339 | 0.6 | 0.932 | 3.09E-99 |
| SLC24A2 | 1.24E-103 | -0.620824725 | 0.7 | 0.953 | 3.57E-99 |
| SCD | 1.39E-103 | -0.879930843 | 0.4 | 0.726 | 4.01E-99 |
| PHYHIP | 2.11E-103 | 1.152818425 | 0.1 | 0.025 | 6.10E-99 |
| ENPP2 | 4.87E-103 | -0.641424346 | 0.6 | 0.903 | 1.41E-98 |
| PXK | 6.60E-103 | -0.643958058 | 0.6 | 0.911 | 1.91E-98 |
| ZNF536 | 1.34E-102 | -0.663065589 | 0.6 | 0.923 | 3.87E-98 |
| XKR4 | 1.99E-102 | 0.947767973 | 0.2 | 0.033 | 5.75E-98 |
| PTPRK | 3.48E-102 | -0.664586319 | 0.7 | 0.949 | 1.01E-97 |
| PDE3B | 5.72E-102 | 0.81413245 | 0.1 | 0.03 | 1.65E-97 |
| DKK3 | 6.01E-102 | 1.164381387 | 0.2 | 0.043 | 1.74E-97 |
| FRYL | 6.13E-102 | -0.646237994 | 0.6 | 0.953 | 1.77E-97 |
| GABRA2 | 1.59E-101 | 0.75958274 | 0.1 | 0.023 | 4.61E-97 |
| LINGO2 | 3.95E-101 | 1.420542035 | 0.2 | 0.051 | 1.14E-96 |
| CLMN | 8.66E-101 | -0.648317599 | 0.6 | 0.925 | 2.50E-96 |
| CAMK4 | 1.07E-100 | 0.826089204 | 0.1 | 0.024 | 3.10E-96 |
| ZNF804A | 2.14E-100 | 0.661295494 | 0.1 | 0.02 | 6.19E-96 |
| DPYD | 7.48E-100 | -0.673576954 | 0.6 | 0.943 | 2.16E-95 |
| PCDH9 | 1.37E-99 | -0.511183657 | 0.9 | 0.976 | 3.95E-95 |
| FA2H | 2.18E-99 | -0.642719125 | 0.5 | 0.804 | 6.31E-95 |
| MMP16 | 3.10E-99 | 0.938454404 | 0.1 | 0.032 | 8.96E-95 |
| SGK1 | 8.08E-99 | -0.652880059 | 0.6 | 0.911 | 2.33E-94 |
| SLC14A1 | 1.13E-98 | 0.758242256 | 0.1 | 0.018 | 3.26E-94 |
| KIAA1217 | 5.71E-98 | 1.161915275 | 0.2 | 0.043 | 1.65E-93 |
| CLDND1 | 6.89E-98 | -0.660333925 | 0.5 | 0.837 | 1.99E-93 |
| MYRF | 1.37E-97 | -0.628419163 | 0.4 | 0.782 | 3.96E-93 |
| PTPRT | 3.44E-97 | 1.090714352 | 0.2 | 0.041 | 9.95E-93 |
| DTNA | 5.08E-97 | 1.954386957 | 0.6 | 0.343 | 1.47E-92 |
| MAG | 7.02E-97 | -0.713414296 | 0.4 | 0.729 | 2.03E-92 |
| GABRG2 | 7.75E-97 | 0.86194034 | 0.1 | 0.017 | 2.24E-92 |
| CREB5 | 1.05E-96 | -0.656016704 | 0.6 | 0.902 | 3.02E-92 |
| DOCK1 | 1.82E-96 | -0.621836959 | 0.6 | 0.892 | 5.26E-92 |
| TCF12 | 1.96E-96 | -0.618937902 | 0.7 | 0.961 | 5.66E-92 |
| SNCB | 2.80E-96 | 1.052161997 | 0.1 | 0.019 | 8.09E-92 |
| GALNT17 | 9.69E-96 | 1.080224492 | 0.2 | 0.044 | 2.80E-91 |
| QDPR | 1.46E-95 | -0.723585575 | 0.5 | 0.845 | 4.22E-91 |
| GFAP | 1.91E-95 | 1.824586734 | 0.2 | 0.077 | 5.51E-91 |
| SHROOM4 | 2.06E-95 | -0.667310029 | 0.5 | 0.81 | 5.96E-91 |
| CLASP2 | 2.30E-95 | -0.56023405 | 0.7 | 0.955 | 6.64E-91 |
| AGAP1 | 4.43E-95 | -0.580564367 | 0.7 | 0.969 | 1.28E-90 |
| DNAJC6 | 9.20E-95 | -0.646448931 | 0.6 | 0.909 | 2.66E-90 |
| NEFL | 9.62E-95 | 1.401307024 | 0.1 | 0.021 | 2.78E-90 |
| SLC17A7 | 2.03E-94 | 0.991441007 | 0.1 | 0.017 | 5.85E-90 |
| ANKRD30BL | 2.47E-94 | 0.746250964 | 0.1 | 0.022 | 7.15E-90 |
| ZNF804B | 3.12E-94 | 0.708225859 | 0.1 | 0.017 | 9.02E-90 |
| TMTC2 | 3.39E-94 | -0.655698713 | 0.7 | 0.946 | 9.81E-90 |
| SLCO3A1 | 5.07E-94 | -0.621123177 | 0.6 | 0.901 | 1.47E-89 |
| SYNJ2 | 5.70E-94 | -0.660795287 | 0.5 | 0.817 | 1.65E-89 |
| PCSK2 | 6.51E-94 | 0.913027071 | 0.1 | 0.023 | 1.88E-89 |
| WNK2 | 7.86E-94 | 0.527092203 | 0.1 | 0.016 | 2.27E-89 |
| IQCA1 | 9.84E-94 | 0.719003002 | 0.1 | 0.018 | 2.84E-89 |
| TTLL7 | 2.42E-93 | -0.602514551 | 0.7 | 0.951 | 6.98E-89 |
| ABCA2 | 2.94E-93 | -0.589733668 | 0.6 | 0.894 | 8.50E-89 |
| CDK18 | 5.51E-93 | -0.637171544 | 0.5 | 0.814 | 1.59E-88 |
| ABCA8 | 7.94E-93 | -0.664048825 | 0.5 | 0.785 | 2.30E-88 |
| ZNF385B | 1.36E-92 | 0.980678285 | 0.2 | 0.037 | 3.94E-88 |
| ADIPOR2 | 2.70E-92 | -0.650243483 | 0.5 | 0.813 | 7.80E-88 |
| PRDM16 | 6.28E-92 | 0.744534589 | 0.1 | 0.017 | 1.81E-87 |
| CNP | 6.63E-91 | -0.654443286 | 0.5 | 0.839 | 1.92E-86 |
| DNM1 | 9.86E-91 | 1.570614934 | 0.3 | 0.088 | 2.85E-86 |
| DGKB | 1.05E-90 | 1.083034051 | 0.2 | 0.054 | 3.03E-86 |
| NOL4 | 5.16E-90 | 0.814678066 | 0.1 | 0.03 | 1.49E-85 |
| NMNAT2 | 5.41E-90 | 0.829824909 | 0.1 | 0.024 | 1.56E-85 |
| DIP2B | 7.95E-90 | -0.586683789 | 0.6 | 0.941 | 2.30E-85 |
| OSBPL6 | 8.83E-90 | 0.777340164 | 0.2 | 0.038 | 2.55E-85 |
| CERCAM | 1.41E-89 | -0.595797662 | 0.5 | 0.867 | 4.09E-85 |
| GAB1 | 1.89E-89 | -0.606562138 | 0.6 | 0.918 | 5.47E-85 |
| NXPH1 | 2.20E-89 | 0.79879022 | 0.1 | 0.019 | 6.36E-85 |
| FARP1 | 4.90E-89 | 1.00513274 | 0.2 | 0.047 | 1.42E-84 |
| GPM6B | 7.50E-89 | -0.613403457 | 0.7 | 0.93 | 2.17E-84 |
| TENM4 | 9.66E-89 | 1.070863865 | 0.2 | 0.048 | 2.79E-84 |
| HHIP | 3.18E-88 | -0.670179848 | 0.5 | 0.808 | 9.19E-84 |
| MOBP | 3.28E-88 | -0.677415275 | 0.6 | 0.867 | 9.48E-84 |
| KCTD16 | 4.25E-88 | 1.119572546 | 0.2 | 0.071 | 1.23E-83 |
| SPOCK1 | 4.42E-88 | -0.622478606 | 0.7 | 0.929 | 1.28E-83 |
| NR2F1-AS1 | 1.78E-87 | 0.795157789 | 0.2 | 0.043 | 5.14E-83 |
| CPE | 2.28E-87 | 1.381080123 | 0.3 | 0.104 | 6.59E-83 |
| AC073050.1 | 3.10E-87 | 1.019766264 | 0.2 | 0.079 | 8.96E-83 |
| PLLP | 4.31E-87 | -0.641236658 | 0.4 | 0.756 | 1.25E-82 |
| PTPRG | 5.57E-87 | 0.448453992 | 0.2 | 0.073 | 1.61E-82 |
| SYN1 | 1.14E-86 | 0.956756446 | 0.1 | 0.021 | 3.31E-82 |
| GSN | 1.30E-86 | -0.655588287 | 0.5 | 0.794 | 3.76E-82 |
| UNC80 | 2.00E-86 | 0.819783947 | 0.1 | 0.027 | 5.77E-82 |
| SH3GL3 | 2.20E-86 | -0.631062374 | 0.6 | 0.88 | 6.35E-82 |
| AC026316.5 | 2.37E-86 | -0.70308709 | 0.5 | 0.849 | 6.85E-82 |
| LPAR1 | 3.28E-86 | -0.612197229 | 0.5 | 0.839 | 9.47E-82 |
| CNTN4 | 4.12E-86 | 1.074669824 | 0.2 | 0.056 | 1.19E-81 |
| GRIK1 | 6.78E-86 | 1.058320107 | 0.2 | 0.045 | 1.96E-81 |
| JAM3 | 4.73E-85 | -0.581944745 | 0.5 | 0.797 | 1.37E-80 |
| COL4A5 | 1.28E-84 | -0.587910178 | 0.5 | 0.831 | 3.69E-80 |
| PAK3 | 2.48E-84 | 1.039431607 | 0.2 | 0.041 | 7.17E-80 |
| CHST11 | 3.68E-84 | 0.953004214 | 0.1 | 0.037 | 1.06E-79 |
| ZEB2 | 4.28E-84 | -0.565640341 | 0.7 | 0.949 | 1.24E-79 |
| DOCK9 | 6.85E-84 | -0.562404247 | 0.6 | 0.873 | 1.98E-79 |
| APBB2 | 7.48E-84 | -0.573930359 | 0.6 | 0.918 | 2.16E-79 |
| CNTN2 | 7.97E-84 | -0.573561619 | 0.5 | 0.842 | 2.30E-79 |
| LAMP2 | 8.29E-84 | -0.565336519 | 0.5 | 0.849 | 2.40E-79 |
| CDK19 | 8.92E-84 | -0.622948497 | 0.6 | 0.879 | 2.58E-79 |
| USP54 | 1.05E-83 | -0.565020063 | 0.6 | 0.877 | 3.04E-79 |
| KALRN | 1.11E-83 | 1.338942678 | 0.4 | 0.168 | 3.20E-79 |
| PHLDB1 | 2.40E-83 | -0.571935921 | 0.5 | 0.803 | 6.93E-79 |
| SH3TC2 | 2.73E-83 | -0.643266778 | 0.4 | 0.695 | 7.88E-79 |
| HIP1 | 2.22E-82 | -0.616020383 | 0.5 | 0.856 | 6.40E-78 |
| LINC00639 | 3.21E-82 | -0.662638244 | 0.4 | 0.744 | 9.28E-78 |
| CADPS2 | 3.34E-82 | 1.114777961 | 0.2 | 0.053 | 9.64E-78 |
| HIPK2 | 6.09E-82 | -0.565396898 | 0.6 | 0.879 | 1.76E-77 |
| LPGAT1 | 8.75E-82 | -0.615022237 | 0.5 | 0.827 | 2.53E-77 |
| DDAH1 | 2.13E-81 | 0.655836743 | 0.1 | 0.023 | 6.15E-77 |
| DIP2C | 4.50E-81 | -0.531311575 | 0.6 | 0.935 | 1.30E-76 |
| NFASC | 7.57E-81 | -0.536867762 | 0.6 | 0.892 | 2.19E-76 |
| IQGAP1 | 7.76E-81 | -0.573550687 | 0.4 | 0.755 | 2.24E-76 |
| KCNMB4 | 1.03E-80 | -0.766455524 | 0.4 | 0.715 | 2.98E-76 |
| SLCO1A2 | 1.85E-80 | -0.567861812 | 0.5 | 0.81 | 5.35E-76 |
| PEX5L | 1.94E-80 | -0.611456789 | 0.7 | 0.934 | 5.60E-76 |
| MAP4K4 | 2.21E-80 | -0.546290216 | 0.7 | 0.937 | 6.40E-76 |
| EPHA4 | 3.09E-80 | 0.839043141 | 0.1 | 0.024 | 8.94E-76 |
| OLFM1 | 4.41E-80 | 1.152499143 | 0.1 | 0.026 | 1.27E-75 |
| GRAMD1B | 5.01E-80 | 0.698344608 | 0.1 | 0.02 | 1.45E-75 |
| SH3D19 | 6.72E-80 | -0.60287827 | 0.5 | 0.824 | 1.94E-75 |

| | | | | | |
|---|---|---|---|---|---|
| TTYH2 | 1.15E-79 | -0.547585831 | 0.4 | 0.742 | 3.33E-75 |
| AGTPBP1 | 1.21E-79 | -0.621398143 | 0.5 | 0.78 | 3.48E-75 |
| HAPLN2 | 1.29E-79 | -0.651400456 | 0.4 | 0.678 | 3.71E-75 |
| PSEN1 | 1.34E-79 | -0.535770157 | 0.5 | 0.786 | 3.87E-75 |
| PCSK6 | 1.35E-79 | -0.67017262 | 0.5 | 0.798 | 3.91E-75 |
| POLR2F | 1.45E-79 | -0.638155466 | 0.5 | 0.819 | 4.19E-75 |
| KIF13B | 1.49E-79 | -0.588829972 | 0.5 | 0.756 | 4.31E-75 |
| ARHGAP21 | 7.32E-79 | -0.548211171 | 0.7 | 0.938 | 2.11E-74 |
| MYO1D | 1.46E-78 | -0.586789603 | 0.5 | 0.825 | 4.23E-74 |
| ANK3 | 1.55E-78 | -0.514070925 | 0.7 | 0.954 | 4.49E-74 |
| ZEB1 | 3.08E-78 | 0.637499541 | 0.2 | 0.057 | 8.89E-74 |
| ANO4 | 3.38E-78 | -0.638015824 | 0.6 | 0.863 | 9.76E-74 |
| DAAM2 | 4.01E-78 | -0.533875288 | 0.5 | 0.847 | 1.16E-73 |
| POU6F2 | 5.74E-78 | 0.650629617 | 0.1 | 0.024 | 1.66E-73 |
| TNR | 3.78E-77 | 0.805162674 | 0.1 | 0.026 | 1.09E-72 |
| CARNS1 | 4.52E-77 | -0.573101571 | 0.4 | 0.71 | 1.31E-72 |
| KLHL32 | 5.03E-77 | -0.609429462 | 0.6 | 0.875 | 1.45E-72 |
| C9orf3 | 1.16E-76 | -0.542170848 | 0.6 | 0.884 | 3.34E-72 |
| LSAMP | 1.43E-76 | 0.914454264 | 0.9 | 0.841 | 4.13E-72 |
| HDAC2-AS2 | 2.25E-76 | -0.685543949 | 0.5 | 0.78 | 6.51E-72 |
| RYBP | 2.67E-76 | -0.565499374 | 0.5 | 0.787 | 7.72E-72 |
| RGS6 | 3.29E-76 | 0.761861112 | 0.1 | 0.038 | 9.50E-72 |
| DLC1 | 3.54E-76 | -0.783569574 | 0.5 | 0.781 | 1.02E-71 |
| CDHR3 | 7.90E-76 | 0.997622701 | 0.1 | 0.027 | 2.28E-71 |
| MIR181A1HG | 1.37E-75 | -0.583966451 | 0.5 | 0.798 | 3.96E-71 |
| 7-Sep | 3.29E-75 | -0.530017174 | 0.7 | 0.942 | 9.50E-71 |
| NPC1 | 1.28E-74 | -0.556837839 | 0.4 | 0.685 | 3.71E-70 |
| HIF3A | 2.19E-74 | 0.919021277 | 0.1 | 0.037 | 6.33E-70 |
| NKX6-2 | 6.99E-74 | -0.619936218 | 0.4 | 0.647 | 2.02E-69 |
| PALM2 | 8.50E-74 | -0.659870916 | 0.5 | 0.812 | 2.46E-69 |
| DSCAML1 | 9.51E-74 | -0.579540181 | 0.5 | 0.833 | 2.75E-69 |
| FMNL2 | 9.60E-74 | -0.511895436 | 0.7 | 0.958 | 2.77E-69 |
| TULP4 | 1.09E-73 | -0.524941577 | 0.5 | 0.848 | 3.14E-69 |
| ZNF708 | 1.49E-73 | -0.538695775 | 0.4 | 0.675 | 4.30E-69 |
| SLC12A2 | 1.86E-73 | -0.537418645 | 0.4 | 0.752 | 5.36E-69 |
| AQP4 | 2.34E-73 | 0.949779821 | 0.1 | 0.037 | 6.75E-69 |
| LINC00499 | 3.03E-73 | 0.705911777 | 0.1 | 0.02 | 8.77E-69 |
| PNMA2 | 7.26E-73 | 1.01626522 | 0.1 | 0.028 | 2.10E-68 |
| ATG4C | 8.00E-73 | -0.532638201 | 0.5 | 0.758 | 2.31E-68 |
| PKP4 | 8.29E-73 | -0.584162198 | 0.5 | 0.825 | 2.40E-68 |
| TJP1 | 9.45E-73 | -0.5182243 | 0.6 | 0.94 | 2.73E-68 |
| KCNQ3 | 1.14E-72 | 1.160409274 | 0.2 | 0.09 | 3.31E-68 |
| MAP4K5 | 1.69E-72 | -0.53078624 | 0.6 | 0.874 | 4.88E-68 |
| GAP43 | 3.78E-72 | 0.87450607 | 0.1 | 0.021 | 1.09E-67 |
| MCTP1 | 8.11E-72 | 0.933045832 | 0.1 | 0.035 | 2.34E-67 |
| SRRM3 | 1.08E-71 | 0.639039062 | 0.1 | 0.02 | 3.12E-67 |
| SORBS2 | 1.11E-71 | 0.6969588 | 0.1 | 0.038 | 3.21E-67 |
| AMPH | 1.35E-71 | 0.751660739 | 0.1 | 0.023 | 3.89E-67 |
| PTBP2 | 1.57E-71 | -0.491723243 | 0.6 | 0.881 | 4.55E-67 |
| CDH19 | 1.57E-71 | -0.634182098 | 0.4 | 0.696 | 4.55E-67 |
| FAM19A1 | 2.06E-71 | 0.734845912 | 0.1 | 0.032 | 5.94E-67 |
| SELENOP | 6.24E-71 | -0.630205912 | 0.4 | 0.644 | 1.80E-66 |
| FOXN2 | 1.77E-70 | -0.516167643 | 0.4 | 0.687 | 5.11E-66 |
| MTUS1 | 4.26E-70 | -0.569429628 | 0.5 | 0.804 | 1.23E-65 |
| DLEU2 | 4.43E-70 | -0.509220692 | 0.5 | 0.831 | 1.28E-65 |
| KCNB2 | 5.83E-70 | 0.62858244 | 0.1 | 0.022 | 1.68E-65 |
| SORBS1 | 5.99E-70 | 1.441267091 | 0.4 | 0.237 | 1.73E-65 |
| FGFR2 | 7.14E-70 | -0.522053692 | 0.5 | 0.813 | 2.06E-65 |
| TRIM2 | 8.89E-70 | -0.494634636 | 0.7 | 0.929 | 2.57E-65 |
| COLGALT2 | 1.47E-69 | -0.598119036 | 0.5 | 0.738 | 4.25E-65 |
| CHD5 | 2.65E-69 | 0.665641238 | 0.1 | 0.022 | 7.66E-65 |
| CNTNAP4 | 3.97E-69 | -0.701239678 | 0.5 | 0.745 | 1.15E-64 |
| CAP2 | 5.60E-69 | 0.700832712 | 0.1 | 0.021 | 1.62E-64 |
| CD22 | 6.47E-69 | -0.535565785 | 0.4 | 0.695 | 1.87E-64 |
| CLIC4 | 9.40E-69 | -0.538759916 | 0.4 | 0.717 | 2.72E-64 |
| KDM6A | 1.41E-68 | -0.511032667 | 0.5 | 0.759 | 4.08E-64 |
| ANLN | 1.60E-68 | -0.586943256 | 0.4 | 0.727 | 4.63E-64 |
| PPP2R2B | 2.12E-68 | -0.569124646 | 0.8 | 0.95 | 6.11E-64 |
| SUN2 | 2.76E-68 | -0.469666485 | 0.4 | 0.727 | 7.98E-64 |
| AKAP6 | 4.25E-68 | -0.487120227 | 0.7 | 0.924 | 1.23E-63 |
| SLC22A23 | 1.49E-67 | -0.53764034 | 0.5 | 0.764 | 4.30E-63 |
| VRK2 | 1.53E-67 | -0.615794182 | 0.4 | 0.698 | 4.42E-63 |
| ATP8A1 | 2.53E-67 | -0.508244963 | 0.6 | 0.905 | 7.31E-63 |
| SYN3 | 3.31E-67 | 0.984688156 | 0.2 | 0.044 | 9.55E-63 |
| LINC01608 | 4.17E-67 | -0.604465931 | 0.5 | 0.785 | 1.21E-62 |
| AATK | 5.36E-67 | -0.462692547 | 0.5 | 0.787 | 1.55E-62 |
| GABRG3 | 5.46E-67 | 0.778858314 | 0.1 | 0.037 | 1.58E-62 |
| DCLK2 | 5.88E-67 | 1.197785643 | 0.2 | 0.08 | 1.70E-62 |
| SCN3A | 1.01E-66 | 0.76260119 | 0.1 | 0.023 | 2.93E-62 |
| ALCAM | 1.09E-66 | -0.527501773 | 0.6 | 0.889 | 3.14E-62 |
| FAM19A2 | 1.64E-66 | 1.104775035 | 0.2 | 0.091 | 4.74E-62 |
| FAM13C | 1.65E-66 | -0.548943703 | 0.4 | 0.737 | 4.77E-62 |
| SEMA6A | 1.71E-66 | -0.516001205 | 0.5 | 0.764 | 4.95E-62 |
| ATP1A2 | 1.72E-66 | 0.724258598 | 0.2 | 0.048 | 4.96E-62 |
| SGK3 | 1.93E-66 | -0.465611579 | 0.5 | 0.76 | 5.58E-62 |
| CCK | 2.76E-66 | 1.10048897 | 0.1 | 0.028 | 7.96E-62 |
| TMEM63A | 2.87E-66 | -0.533679867 | 0.4 | 0.647 | 8.28E-62 |
| ATP10B | 3.22E-66 | -0.527754003 | 0.5 | 0.733 | 9.30E-62 |
| IGSF11 | 3.50E-66 | -0.498892686 | 0.5 | 0.747 | 1.01E-61 |
| APLP1 | 6.54E-66 | -0.507597936 | 0.6 | 0.822 | 1.89E-61 |
| PTPRN2 | 6.94E-66 | 1.146552122 | 0.2 | 0.063 | 2.01E-61 |
| DLG2 | 1.02E-65 | -0.340576058 | 0.8 | 0.984 | 2.96E-61 |
| PRKAR1B | 1.13E-65 | 1.078170087 | 0.1 | 0.038 | 3.26E-61 |
| CHN1 | 1.56E-65 | 1.857892282 | 0.3 | 0.179 | 4.50E-61 |
| FUT8 | 1.76E-65 | -0.509925859 | 0.6 | 0.891 | 5.09E-61 |
| TBC1D12 | 1.81E-65 | -0.604081633 | 0.3 | 0.563 | 5.22E-61 |
| SGMS1 | 2.19E-65 | -0.460103303 | 0.5 | 0.813 | 6.32E-61 |
| MEGF10 | 2.52E-65 | -0.508406836 | 0.5 | 0.745 | 7.29E-61 |
| TPD52L1 | 3.10E-65 | 0.863748659 | 0.2 | 0.05 | 8.97E-61 |
| ELF2 | 4.92E-65 | -0.459191366 | 0.5 | 0.786 | 1.42E-60 |
| TMEM132B | 6.26E-65 | 0.915918369 | 0.2 | 0.078 | 1.81E-60 |
| RNF130 | 1.06E-64 | -0.480415882 | 0.6 | 0.882 | 3.07E-60 |
| CD9 | 1.38E-64 | -0.479477464 | 0.3 | 0.581 | 3.99E-60 |
| TSHZ2 | 1.46E-64 | 0.804220754 | 0.1 | 0.025 | 4.22E-60 |
| GLDN | 1.46E-64 | -0.523187473 | 0.4 | 0.691 | 4.23E-60 |
| DICER1 | 1.48E-64 | -0.436004208 | 0.5 | 0.801 | 4.27E-60 |
| ARHGAP23 | 1.81E-64 | -0.45494463 | 0.4 | 0.668 | 5.24E-60 |
| TMEM132D | 2.57E-64 | 0.929395824 | 0.2 | 0.046 | 7.43E-60 |
| FOLH1 | 2.89E-64 | -0.580895408 | 0.4 | 0.696 | 8.34E-60 |
| GJA1 | 4.52E-64 | 0.84696168 | 0.1 | 0.032 | 1.31E-59 |
| DAPK1 | 7.76E-64 | 0.655130167 | 0.1 | 0.033 | 2.24E-59 |
| MAL | 1.41E-63 | -0.550737433 | 0.4 | 0.642 | 4.06E-59 |
| LANCL1 | 1.45E-63 | -0.494295534 | 0.5 | 0.788 | 4.20E-59 |
| MAGI2 | 1.63E-63 | -0.415287029 | 0.8 | 0.983 | 4.70E-59 |
| RBPJ | 2.13E-63 | -0.463377663 | 0.5 | 0.798 | 6.16E-59 |
| TMTC4 | 2.96E-63 | -0.541748174 | 0.3 | 0.613 | 8.55E-59 |
| ARAP2 | 3.27E-63 | -0.494708196 | 0.6 | 0.933 | 9.44E-59 |
| RFTN2 | 4.38E-63 | -0.507764766 | 0.5 | 0.807 | 1.27E-58 |
| NCAM2 | 5.30E-63 | -0.520072067 | 0.8 | 0.953 | 1.53E-58 |
| KSR2 | 6.84E-63 | 0.722908355 | 0.1 | 0.024 | 1.98E-58 |
| PPP2R2A | 6.90E-63 | -0.464260635 | 0.5 | 0.755 | 1.99E-58 |
| COBL | 1.21E-62 | -0.472581925 | 0.6 | 0.869 | 3.48E-58 |
| KIF13A | 1.22E-62 | -0.452781309 | 0.5 | 0.819 | 3.54E-58 |
| RNF13 | 2.23E-62 | -0.467930698 | 0.5 | 0.751 | 6.45E-58 |
| ZBTB20 | 2.37E-62 | -0.442905752 | 0.7 | 0.981 | 6.84E-58 |
| SLC25A13 | 2.71E-62 | -0.566300481 | 0.4 | 0.664 | 7.83E-58 |
| GPR37 | 2.85E-62 | -0.5111212 | 0.4 | 0.687 | 8.24E-58 |
| MACROD2 | 3.03E-62 | 1.109641526 | 0.4 | 0.183 | 8.76E-58 |
| STK39 | 3.89E-62 | -0.47509907 | 0.4 | 0.696 | 1.13E-57 |
| FAM107A | 8.95E-62 | 0.810051898 | 0.1 | 0.025 | 2.59E-57 |
| MVB12B | 9.12E-62 | -0.475131234 | 0.5 | 0.779 | 2.64E-57 |
| LIMCH1 | 1.47E-61 | -0.452501671 | 0.7 | 0.921 | 4.24E-57 |
| PTPRD | 1.51E-61 | -0.285421105 | 0.8 | 0.96 | 4.36E-57 |
| PTPRJ | 1.83E-61 | -0.500661614 | 0.5 | 0.754 | 5.29E-57 |
| HS3ST5 | 2.57E-61 | -0.546492154 | 0.4 | 0.725 | 7.42E-57 |
| MOB3B | 2.73E-61 | -0.476299386 | 0.4 | 0.723 | 7.88E-57 |
| TMEFF2 | 3.03E-61 | -0.535432981 | 0.6 | 0.858 | 8.75E-57 |
| RDX | 3.15E-61 | -0.477156852 | 0.4 | 0.721 | 9.11E-57 |

| | | | | | |
|---|---|---|---|---|---|
| ADCY1 | 6.67E-61 | 0.83155143 | 0.1 | 0.023 | 1.93E-56 |
| BIN1 | 9.44E-61 | -0.420075326 | 0.5 | 0.814 | 2.73E-56 |
| NDRG1 | 9.92E-61 | -0.453774596 | 0.4 | 0.729 | 2.87E-56 |
| SGCZ | 1.37E-60 | 0.882318799 | 0.2 | 0.065 | 3.95E-56 |
| KAT2B | 1.41E-60 | -0.44120281 | 0.4 | 0.722 | 4.08E-56 |
| PTK2 | 2.17E-60 | -0.453264804 | 0.7 | 0.947 | 6.27E-56 |
| RAPGEF5 | 2.30E-60 | -0.45263507 | 0.6 | 0.875 | 6.65E-56 |
| GRM3 | 2.55E-60 | -0.500082743 | 0.6 | 0.871 | 7.36E-56 |
| CCNY | 2.73E-60 | -0.473476615 | 0.5 | 0.74 | 7.89E-56 |
| GNAI1 | 2.88E-60 | -0.462110413 | 0.5 | 0.74 | 8.33E-56 |
| AGPAT4 | 4.23E-60 | -0.531123991 | 0.4 | 0.711 | 1.22E-55 |
| RAB6B | 4.28E-60 | 0.877501267 | 0.1 | 0.029 | 1.24E-55 |
| NUTM2A-AS1 | 4.93E-60 | -0.482935588 | 0.5 | 0.78 | 1.42E-55 |
| PIK3C2B | 5.31E-60 | -0.522207335 | 0.3 | 0.597 | 1.53E-55 |
| OSBPL1A | 5.86E-60 | -0.48621837 | 0.6 | 0.837 | 1.69E-55 |
| 4-Sep | 5.93E-60 | -0.410447036 | 0.4 | 0.642 | 1.71E-55 |
| SEMA4D | 6.28E-60 | -0.462087568 | 0.5 | 0.751 | 1.82E-55 |
| MYO1E | 6.76E-60 | -0.495646546 | 0.4 | 0.697 | 1.95E-55 |
| CACHD1 | 9.73E-60 | 0.719402145 | 0.1 | 0.027 | 2.81E-55 |
| NAV3 | 1.38E-59 | 1.354532277 | 0.6 | 0.426 | 3.99E-55 |
| AC016597.1 | 2.22E-59 | -0.482719693 | 0.3 | 0.587 | 6.41E-55 |
| NHSL1 | 2.72E-59 | 0.624547993 | 0.1 | 0.03 | 7.87E-55 |
| PREX1 | 4.15E-59 | -0.386067903 | 0.4 | 0.727 | 1.20E-54 |
| PLPPR1 | 5.28E-59 | -0.653221869 | 0.3 | 0.564 | 1.53E-54 |
| TNRC6C | 6.79E-59 | -0.373370506 | 0.5 | 0.795 | 1.96E-54 |
| SERINC5 | 8.02E-59 | -0.447087543 | 0.4 | 0.69 | 2.32E-54 |
| CHRM3 | 1.24E-58 | 1.019013905 | 0.2 | 0.084 | 3.59E-54 |
| PDE7B | 1.61E-58 | 0.211177226 | 0.2 | 0.052 | 4.64E-54 |
| MARCKSL1 | 2.49E-58 | -0.502562426 | 0.4 | 0.621 | 7.19E-54 |
| MT-ATP6 | 4.25E-58 | 0.891060297 | 0.8 | 0.819 | 1.23E-53 |
| RANBP3L | 4.26E-58 | 0.589377944 | 0.1 | 0.026 | 1.23E-53 |
| WIPF1 | 4.28E-58 | -0.54365075 | 0.3 | 0.561 | 1.24E-53 |
| CDC14B | 4.41E-58 | -0.4497309 | 0.5 | 0.768 | 1.27E-53 |
| PLA2G16 | 8.29E-58 | -0.40549918 | 0.5 | 0.734 | 2.40E-53 |
| CCP110 | 8.90E-58 | -0.446843087 | 0.5 | 0.751 | 2.57E-53 |
| MYO6 | 1.02E-57 | -0.381901549 | 0.6 | 0.853 | 2.94E-53 |
| ANKIB1 | 1.05E-57 | -0.417824221 | 0.6 | 0.847 | 3.03E-53 |
| PAM | 1.13E-57 | 0.732625179 | 0.2 | 0.055 | 3.26E-53 |
| OXR1 | 1.21E-57 | 1.266019499 | 0.4 | 0.224 | 3.51E-53 |
| ENOSF1 | 1.87E-57 | -0.48873762 | 0.4 | 0.626 | 5.41E-53 |
| UBE4B | 2.91E-57 | -0.423384375 | 0.5 | 0.73 | 8.42E-53 |
| SORT1 | 3.46E-57 | -0.428388734 | 0.6 | 0.875 | 1.00E-52 |
| ITCH | 3.76E-57 | -0.394263295 | 0.5 | 0.774 | 1.09E-52 |
| TUBA1A | 4.46E-57 | -0.330205877 | 0.4 | 0.681 | 1.29E-52 |
| OSBPL9 | 7.31E-57 | -0.428032223 | 0.4 | 0.711 | 2.11E-52 |
| TSPAN7 | 8.54E-57 | 1.342550711 | 0.2 | 0.062 | 2.47E-52 |
| NDRG4 | 8.86E-57 | 1.133723629 | 0.1 | 0.044 | 2.56E-52 |
| NKAIN2 | 9.52E-57 | -0.551032443 | 0.7 | 0.896 | 2.75E-52 |
| PPP6R3 | 1.05E-56 | -0.372189796 | 0.4 | 0.699 | 3.04E-52 |
| ZNF638 | 2.05E-56 | -0.425931494 | 0.7 | 0.938 | 5.92E-52 |
| SLC22A15 | 2.34E-56 | -0.459946474 | 0.4 | 0.676 | 6.75E-52 |
| CRYAB | 2.46E-56 | -0.564899488 | 0.4 | 0.689 | 7.11E-52 |
| PPM1B | 2.49E-56 | -0.400985731 | 0.5 | 0.768 | 7.19E-52 |
| ERMN | 2.57E-56 | -0.481434674 | 0.3 | 0.585 | 7.42E-52 |
| TFEB | 3.31E-56 | -0.459073381 | 0.3 | 0.554 | 9.55E-52 |
| CTTNBP2 | 3.90E-56 | -0.404032225 | 0.4 | 0.721 | 1.13E-51 |
| LRRC1 | 5.12E-56 | -0.507384019 | 0.4 | 0.637 | 1.48E-51 |
| ACER3 | 7.79E-56 | -0.463397237 | 0.4 | 0.707 | 2.25E-51 |
| LIPA | 1.23E-55 | -0.392401495 | 0.4 | 0.707 | 3.55E-51 |
| NCAM1 | 1.46E-55 | -0.402548783 | 0.7 | 0.94 | 4.23E-51 |
| PIEZO2 | 1.99E-55 | -0.618457804 | 0.4 | 0.674 | 5.75E-51 |
| KIAA1324L | 2.70E-55 | -0.481281026 | 0.4 | 0.633 | 7.79E-51 |
| AMER2 | 2.89E-55 | -0.442630997 | 0.5 | 0.82 | 8.34E-51 |
| CACNA1A | 3.29E-55 | 1.005352382 | 0.2 | 0.074 | 9.50E-51 |
| LRP2 | 3.38E-55 | -0.6377594 | 0.4 | 0.603 | 9.76E-51 |
| ST6GALNAC3 | 3.41E-55 | -0.574368387 | 0.5 | 0.73 | 9.84E-51 |
| RNF150 | 3.93E-55 | 0.687117575 | 0.1 | 0.033 | 1.14E-50 |
| ZNF652 | 3.95E-55 | -0.464777822 | 0.4 | 0.704 | 1.14E-50 |
| ARID4B | 4.69E-55 | -0.367856924 | 0.5 | 0.778 | 1.36E-50 |
| ZDHHC20 | 1.21E-54 | -0.586937945 | 0.4 | 0.67 | 3.50E-50 |
| CHD7 | 1.31E-54 | -0.442764176 | 0.4 | 0.675 | 3.78E-50 |
| RAB40B | 1.47E-54 | -0.426248448 | 0.4 | 0.619 | 4.24E-50 |
| ZBTB16 | 1.76E-54 | -0.474729186 | 0.5 | 0.775 | 5.10E-50 |
| FAM124A | 1.85E-54 | -0.469596787 | 0.4 | 0.651 | 5.35E-50 |
| OPALIN | 2.36E-54 | -0.548835771 | 0.4 | 0.618 | 6.81E-50 |
| TP53TG5 | 2.62E-54 | -0.483935237 | 0.3 | 0.603 | 7.56E-50 |
| FAF1 | 3.72E-54 | -0.390642446 | 0.6 | 0.831 | 1.08E-49 |
| MT-CO1 | 4.13E-54 | 1.046439251 | 0.8 | 0.831 | 1.19E-49 |
| GNG7 | 4.24E-54 | -0.50428554 | 0.4 | 0.67 | 1.23E-49 |
| CACNA1C | 4.58E-54 | 0.847528896 | 0.3 | 0.109 | 1.32E-49 |
| SAMD4B | 4.58E-54 | -0.416517698 | 0.4 | 0.637 | 1.32E-49 |
| FANCL | 4.59E-54 | -0.408953762 | 0.3 | 0.556 | 1.33E-49 |
| PRR5L | 4.66E-54 | -0.485063792 | 0.4 | 0.609 | 1.35E-49 |
| SECISBP2L | 4.68E-54 | -0.365114682 | 0.5 | 0.783 | 1.35E-49 |
| STARD13 | 4.70E-54 | -0.46080072 | 0.5 | 0.76 | 1.36E-49 |
| ATXN1 | 5.07E-54 | -0.443791682 | 0.6 | 0.874 | 1.47E-49 |
| EPN2 | 6.45E-54 | -0.389455057 | 0.5 | 0.803 | 1.86E-49 |
| ATP1B2 | 6.88E-54 | 0.847253343 | 0.1 | 0.034 | 1.99E-49 |
| MBNL2 | 6.94E-54 | -0.408670318 | 0.6 | 0.899 | 2.01E-49 |
| GPRC5B | 7.80E-54 | -0.389673313 | 0.5 | 0.812 | 2.26E-49 |
| WSB1 | 8.73E-54 | -0.371658166 | 0.6 | 0.868 | 2.52E-49 |
| PMP22 | 9.06E-54 | -0.383993329 | 0.3 | 0.586 | 2.62E-49 |
| RNF144A | 9.78E-54 | -0.366733752 | 0.5 | 0.749 | 2.83E-49 |
| YPEL2 | 9.97E-54 | -0.479341853 | 0.4 | 0.622 | 2.88E-49 |
| ZKSCAN1 | 1.33E-53 | -0.38201313 | 0.4 | 0.711 | 3.84E-49 |
| TRAPPC10 | 1.54E-53 | -0.41387111 | 0.3 | 0.587 | 4.46E-49 |
| KIF6 | 1.63E-53 | -0.537385846 | 0.4 | 0.679 | 4.72E-49 |
| AUTS2 | 2.11E-53 | -0.439836674 | 0.7 | 0.942 | 6.11E-49 |
| TRPC4AP | 2.17E-53 | -0.321203742 | 0.4 | 0.707 | 6.28E-49 |
| KCNH7 | 2.30E-53 | 0.799294902 | 0.1 | 0.045 | 6.66E-49 |
| GAB2 | 2.53E-53 | -0.444038183 | 0.5 | 0.788 | 7.32E-49 |
| PPP1R16B | 2.58E-53 | -0.42530352 | 0.4 | 0.707 | 7.45E-49 |
| BTBD3 | 2.92E-53 | -0.399512556 | 0.4 | 0.64 | 8.44E-49 |
| FAM102A | 3.77E-53 | -0.420317695 | 0.3 | 0.563 | 1.09E-48 |
| PSME4 | 3.81E-53 | -0.362088403 | 0.4 | 0.692 | 1.10E-48 |
| AQP4-AS1 | 4.64E-53 | 0.789318354 | 0.1 | 0.039 | 1.34E-48 |
| USP47 | 4.66E-53 | -0.336382237 | 0.5 | 0.767 | 1.35E-48 |
| GNA14 | 7.73E-53 | 1.235909666 | 0.2 | 0.051 | 2.23E-48 |
| EVA1C | 8.61E-53 | -0.527827754 | 0.4 | 0.687 | 2.49E-48 |
| WDR20 | 9.96E-53 | -0.445326628 | 0.4 | 0.625 | 2.88E-48 |
| USP32 | 1.08E-52 | -0.408551389 | 0.4 | 0.704 | 3.11E-48 |
| ZCCHC24 | 1.52E-52 | -0.375759147 | 0.4 | 0.667 | 4.39E-48 |
| PPP1R14A | 1.69E-52 | -0.466061398 | 0.2 | 0.465 | 4.87E-48 |
| PACS2 | 1.72E-52 | -0.403248552 | 0.4 | 0.654 | 4.96E-48 |
| RGCC | 1.95E-52 | -0.492423593 | 0.4 | 0.598 | 5.62E-48 |
| CDK6 | 2.18E-52 | -0.486337507 | 0.3 | 0.593 | 6.31E-48 |
| AL359091.1 | 4.10E-52 | -0.477118962 | 0.4 | 0.652 | 1.18E-47 |
| NRG1 | 4.87E-52 | 1.143264571 | 0.1 | 0.047 | 1.41E-47 |
| RHOU | 7.40E-52 | -0.449410752 | 0.4 | 0.603 | 2.14E-47 |
| KIRREL3 | 8.77E-52 | -0.483586325 | 0.6 | 0.824 | 2.53E-47 |
| DUBR | 9.72E-52 | -0.422118684 | 0.4 | 0.669 | 2.81E-47 |
| ZNF644 | 1.19E-51 | -0.352190157 | 0.4 | 0.714 | 3.44E-47 |
| LDB3 | 1.33E-51 | -0.505711448 | 0.4 | 0.596 | 3.84E-47 |
| SIRT2 | 1.36E-51 | -0.410269991 | 0.3 | 0.522 | 3.92E-47 |
| APOD | 1.84E-51 | -0.568303435 | 0.3 | 0.543 | 5.31E-47 |
| SOX6 | 2.44E-51 | 0.844814667 | 0.2 | 0.087 | 7.05E-47 |
| SSH2 | 2.56E-51 | -0.41761431 | 0.6 | 0.861 | 7.39E-47 |
| SEMA3B | 2.80E-51 | -0.391408639 | 0.4 | 0.667 | 8.09E-47 |
| EXOC6B | 4.01E-51 | -0.396538884 | 0.7 | 0.938 | 1.16E-46 |
| AC110285.1 | 5.64E-51 | -0.513038204 | 0.3 | 0.545 | 1.63E-46 |
| CPOX | 6.99E-51 | -0.456066499 | 0.3 | 0.553 | 2.02E-46 |
| CBR1 | 7.91E-51 | -0.391415919 | 0.2 | 0.467 | 2.29E-46 |
| CA2 | 8.03E-51 | -0.538614366 | 0.3 | 0.5 | 2.32E-46 |
| LINC00320 | 8.19E-51 | -0.459210991 | 0.4 | 0.702 | 2.37E-46 |
| NSMCE2 | 8.63E-51 | -0.436452424 | 0.5 | 0.74 | 2.49E-46 |
| RNASE1 | 9.81E-51 | -0.463728929 | 0.3 | 0.505 | 2.84E-46 |
| AC096564.1 | 1.07E-50 | -0.510052465 | 0.3 | 0.512 | 3.09E-46 |
| LINC01170 | 1.25E-50 | -0.649495684 | 0.3 | 0.591 | 3.60E-46 |

| | | | | | |
|---|---|---|---|---|---|
| JPX | 2.18E-50 | -0.370288852 | 0.4 | 0.688 | 6.29E-46 |
| SAMD12 | 2.27E-50 | -0.480038519 | 0.5 | 0.789 | 6.56E-46 |
| GRID1 | 2.96E-50 | -0.39887729 | 0.6 | 0.85 | 8.55E-46 |
| SCD5 | 3.77E-50 | -0.388193662 | 0.6 | 0.887 | 1.09E-45 |
| PARD3 | 4.05E-50 | 0.777488117 | 0.2 | 0.099 | 1.17E-45 |
| CCDC88A | 5.65E-50 | -0.389830351 | 0.7 | 0.939 | 1.63E-45 |
| LAMA2 | 6.41E-50 | -1.019059013 | 0.3 | 0.56 | 1.85E-45 |
| CNTNAP3B | 7.93E-50 | 0.714681692 | 0.1 | 0.032 | 2.29E-45 |
| LRRFIP2 | 1.07E-49 | -0.33810282 | 0.4 | 0.704 | 3.09E-45 |
| ZFYVE16 | 1.10E-49 | -0.385987249 | 0.5 | 0.792 | 3.18E-45 |
| DUSP16 | 1.13E-49 | -0.522360412 | 0.3 | 0.559 | 3.25E-45 |
| RIMS1 | 1.22E-49 | 1.082412509 | 0.2 | 0.106 | 3.53E-45 |
| ASPA | 1.85E-49 | -0.431616023 | 0.4 | 0.65 | 5.36E-45 |
| C12orf76 | 2.38E-49 | -0.398784929 | 0.4 | 0.632 | 6.89E-45 |
| NLGN1 | 3.25E-49 | -0.471812504 | 0.7 | 0.906 | 9.40E-45 |
| OTUD7A | 4.20E-49 | -0.427503865 | 0.6 | 0.893 | 1.21E-44 |
| MPP5 | 4.28E-49 | -0.415336316 | 0.3 | 0.518 | 1.24E-44 |
| CHKA | 4.85E-49 | -0.37516802 | 0.4 | 0.689 | 1.40E-44 |
| TSPYL1 | 4.91E-49 | 1.181816196 | 0.2 | 0.07 | 1.42E-44 |
| EPS15 | 4.92E-49 | -0.328615386 | 0.5 | 0.762 | 1.42E-44 |
| KANSL1 | 6.19E-49 | -0.343837381 | 0.5 | 0.775 | 1.79E-44 |
| UVRAG | 6.35E-49 | -0.368684356 | 0.4 | 0.709 | 1.84E-44 |
| ZNF565 | 1.11E-48 | -0.403156974 | 0.3 | 0.493 | 3.21E-44 |
| ATP11A | 1.48E-48 | -0.348984703 | 0.4 | 0.642 | 4.26E-44 |
| CYTH1 | 1.51E-48 | -0.362594592 | 0.4 | 0.623 | 4.37E-44 |
| CTNNA1 | 1.94E-48 | -0.349749756 | 0.4 | 0.641 | 5.61E-44 |
| PCDH9-AS2 | 2.14E-48 | -0.448074638 | 0.2 | 0.441 | 6.19E-44 |
| FEZ1 | 2.80E-48 | -0.310966041 | 0.4 | 0.6 | 8.10E-44 |
| DENND5A | 2.84E-48 | -0.316754322 | 0.5 | 0.721 | 8.22E-44 |
| SCARB2 | 2.96E-48 | -0.322485866 | 0.4 | 0.678 | 8.56E-44 |
| ANKRD44 | 3.16E-48 | -0.38423339 | 0.4 | 0.685 | 9.14E-44 |
| FNBP1 | 3.16E-48 | -0.384466516 | 0.6 | 0.876 | 9.15E-44 |
| NAIP | 3.93E-48 | -0.355804193 | 0.4 | 0.679 | 1.14E-43 |
| PPP1R12B | 4.59E-48 | -0.375566975 | 0.5 | 0.781 | 1.33E-43 |
| FNIP2 | 5.23E-48 | -0.414991147 | 0.4 | 0.662 | 1.51E-43 |
| ZDHHC9 | 6.22E-48 | -0.449066146 | 0.3 | 0.509 | 1.80E-43 |
| POGZ | 6.79E-48 | -0.2922065 | 0.5 | 0.768 | 1.96E-43 |
| HECW2 | 7.47E-48 | -0.378369312 | 0.5 | 0.796 | 2.16E-43 |
| RALGAPA1 | 1.08E-47 | -0.325125266 | 0.5 | 0.777 | 3.12E-43 |
| PLEKHG3 | 1.10E-47 | -0.431614734 | 0.3 | 0.544 | 3.17E-43 |
| ZSWIM6 | 1.77E-47 | -0.448075297 | 0.6 | 0.886 | 5.12E-43 |
| RICTOR | 1.81E-47 | -0.333465025 | 0.5 | 0.753 | 5.22E-43 |
| STRN | 2.83E-47 | -0.40533037 | 0.4 | 0.591 | 8.17E-43 |
| SLC31A2 | 3.30E-47 | -0.438322971 | 0.2 | 0.468 | 9.55E-43 |
| ITGB8 | 3.34E-47 | -0.323597016 | 0.4 | 0.651 | 9.65E-43 |
| CPB2-AS1 | 4.74E-47 | -0.480969254 | 0.3 | 0.58 | 1.37E-42 |
| SLF2 | 5.13E-47 | -0.329775347 | 0.4 | 0.618 | 1.48E-42 |
| CUEDC1 | 6.61E-47 | -0.419136755 | 0.3 | 0.571 | 1.91E-42 |
| HEPN1 | 7.11E-47 | -0.288702521 | 0.2 | 0.369 | 2.06E-42 |
| SORCS1 | 7.85E-47 | 0.97493749 | 0.1 | 0.042 | 2.27E-42 |
| CERS6 | 1.00E-46 | 0.869153188 | 0.2 | 0.066 | 2.89E-42 |
| PKNOX2 | 1.27E-46 | 0.633591581 | 0.1 | 0.034 | 3.68E-42 |
| P2RX7 | 1.28E-46 | -0.346156908 | 0.4 | 0.697 | 3.69E-42 |
| GLTP | 1.42E-46 | -0.433048067 | 0.3 | 0.494 | 4.11E-42 |
| ELL2 | 1.80E-46 | -0.283362238 | 0.5 | 0.745 | 5.21E-42 |
| PPP2R3A | 3.00E-46 | -0.436392364 | 0.4 | 0.593 | 8.66E-42 |
| ELF1 | 4.03E-46 | -0.509859779 | 0.2 | 0.467 | 1.17E-41 |
| SPAG9 | 4.37E-46 | -0.267162343 | 0.5 | 0.771 | 1.26E-41 |
| ARHGEF7 | 4.64E-46 | -0.317022015 | 0.5 | 0.743 | 1.34E-41 |
| RERE | 5.39E-46 | -0.321125738 | 0.6 | 0.895 | 1.56E-41 |
| NRDC | 5.40E-46 | -0.286325616 | 0.4 | 0.589 | 1.56E-41 |
| TMEM108 | 5.60E-46 | 0.725577409 | 0.2 | 0.081 | 1.62E-41 |
| ADAM10 | 5.69E-46 | -0.38547281 | 0.4 | 0.683 | 1.64E-41 |
| ZNF146 | 5.70E-46 | -0.383665857 | 0.3 | 0.556 | 1.65E-41 |
| SOX10 | 6.13E-46 | -0.45121604 | 0.3 | 0.489 | 1.77E-41 |
| RETREG1 | 6.73E-46 | -0.350092964 | 0.5 | 0.723 | 1.95E-41 |
| MOSPD2 | 7.70E-46 | -0.43900214 | 0.2 | 0.442 | 2.22E-41 |
| DYM | 8.59E-46 | -0.311846495 | 0.4 | 0.698 | 2.48E-41 |
| MON2 | 8.61E-46 | -0.281994527 | 0.5 | 0.76 | 2.49E-41 |
| NT5DC1 | 1.16E-45 | -0.449609452 | 0.4 | 0.649 | 3.35E-41 |
| WNK1 | 1.66E-45 | -0.311632023 | 0.5 | 0.781 | 4.81E-41 |
| PPP1R21 | 1.68E-45 | -0.342971152 | 0.4 | 0.697 | 4.84E-41 |
| FAR1 | 1.79E-45 | -0.327902425 | 0.4 | 0.657 | 5.16E-41 |
| ZNF608 | 1.89E-45 | -0.418592028 | 0.3 | 0.577 | 5.47E-41 |
| GRM7 | 2.49E-45 | 0.969536299 | 0.3 | 0.129 | 7.18E-41 |
| TACC1 | 2.87E-45 | -0.52211569 | 0.5 | 0.757 | 8.28E-41 |
| TSC22D4 | 3.38E-45 | -0.37862593 | 0.3 | 0.556 | 9.78E-41 |
| CNOT2 | 3.64E-45 | -0.326100588 | 0.4 | 0.663 | 1.05E-40 |
| S100B | 3.71E-45 | -0.421011163 | 0.2 | 0.403 | 1.07E-40 |
| ZNF280D | 7.63E-45 | -0.320389269 | 0.4 | 0.637 | 2.20E-40 |
| NCOA7 | 8.38E-45 | -0.337101376 | 0.5 | 0.783 | 2.42E-40 |
| CCNH | 9.12E-45 | -0.364904834 | 0.6 | 0.842 | 2.63E-40 |
| NPAS2 | 9.77E-45 | 0.633752544 | 0.1 | 0.032 | 2.82E-40 |
| STXBP3 | 1.07E-44 | -0.346629282 | 0.3 | 0.562 | 3.10E-40 |
| NIPBL | 1.48E-44 | -0.298938977 | 0.5 | 0.762 | 4.28E-40 |
| AC118549.1 | 2.76E-44 | -0.352403249 | 0.4 | 0.671 | 7.97E-40 |
| AC012593.1 | 3.96E-44 | -0.681566725 | 0.3 | 0.535 | 1.14E-39 |
| CD46 | 4.45E-44 | -0.361437818 | 0.3 | 0.506 | 1.29E-39 |
| SPATA6 | 4.58E-44 | -0.349981458 | 0.4 | 0.657 | 1.32E-39 |
| PCDH15 | 7.66E-44 | 0.903533113 | 0.1 | 0.044 | 2.21E-39 |
| ZHX2 | 8.25E-44 | -0.385946551 | 0.5 | 0.728 | 2.38E-39 |
| MT-CO2 | 8.84E-44 | 0.647552615 | 0.8 | 0.898 | 2.55E-39 |
| ANAPC5 | 1.33E-43 | -0.306312776 | 0.4 | 0.629 | 3.85E-39 |
| KLHL4 | 1.49E-43 | -0.529830882 | 0.4 | 0.573 | 4.29E-39 |
| RASGRP3 | 1.65E-43 | -0.510881473 | 0.3 | 0.482 | 4.75E-39 |
| BCAS3 | 1.98E-43 | -0.407508389 | 0.5 | 0.788 | 5.73E-39 |
| C1QTNF3-AMACR | 2.74E-43 | -0.39080795 | 0.5 | 0.697 | 7.92E-39 |
| AFDN | 3.04E-43 | -0.292140596 | 0.4 | 0.641 | 8.79E-39 |
| NEO1 | 4.50E-43 | -0.297758707 | 0.6 | 0.822 | 1.30E-38 |
| CSRP1 | 4.53E-43 | -0.350304872 | 0.4 | 0.6 | 1.31E-38 |
| SRCIN1 | 4.54E-43 | -0.360619553 | 0.5 | 0.721 | 1.31E-38 |
| TSPAN15 | 6.51E-43 | -0.407812765 | 0.2 | 0.439 | 1.88E-38 |
| LINC00511 | 7.16E-43 | 0.805663867 | 0.1 | 0.052 | 2.07E-38 |
| ZNF704 | 7.36E-43 | -0.392662467 | 0.5 | 0.743 | 2.13E-38 |
| PKD2 | 7.46E-43 | -0.265700403 | 0.4 | 0.68 | 2.15E-38 |
| ARFGEF3 | 7.54E-43 | -0.327182814 | 0.5 | 0.814 | 2.18E-38 |
| TTLL11 | 7.92E-43 | -0.364805567 | 0.3 | 0.564 | 2.29E-38 |
| PDCD6IP | 1.01E-42 | -0.301792985 | 0.3 | 0.535 | 2.91E-38 |
| PROX1 | 1.12E-42 | -0.432606222 | 0.2 | 0.421 | 3.22E-38 |
| PWWP2A | 1.20E-42 | -0.325534008 | 0.3 | 0.566 | 3.46E-38 |
| TMEM98 | 1.21E-42 | -0.461978482 | 0.3 | 0.511 | 3.50E-38 |
| FAM13B | 1.25E-42 | -0.329866402 | 0.4 | 0.694 | 3.60E-38 |
| TMC7 | 1.32E-42 | -0.432158192 | 0.3 | 0.486 | 3.80E-38 |
| LINC01630 | 1.38E-42 | -0.467821895 | 0.4 | 0.601 | 3.98E-38 |
| GPC6 | 1.88E-42 | 0.266860684 | 0.1 | 0.04 | 5.43E-38 |
| SNX1 | 1.93E-42 | -0.330298681 | 0.4 | 0.578 | 5.59E-38 |
| FSTL5 | 2.00E-42 | 0.672044376 | 0.1 | 0.041 | 5.78E-38 |
| TMEM235 | 2.06E-42 | -0.408076928 | 0.3 | 0.507 | 5.96E-38 |
| TARSL2 | 2.21E-42 | -0.340282756 | 0.4 | 0.666 | 6.40E-38 |
| TECPR2 | 2.23E-42 | -0.35362641 | 0.4 | 0.667 | 6.44E-38 |
| KMT2E | 2.36E-42 | -0.234059402 | 0.5 | 0.752 | 6.83E-38 |
| AC006059.1 | 2.37E-42 | -0.490526908 | 0.4 | 0.591 | 6.85E-38 |
| VPS35L | 2.60E-42 | -0.338205217 | 0.4 | 0.649 | 7.51E-38 |
| LINC00609 | 2.74E-42 | -0.51674306 | 0.4 | 0.616 | 7.93E-38 |
| HBS1L | 2.86E-42 | -0.349999247 | 0.4 | 0.616 | 8.26E-38 |
| TBC1D5 | 2.95E-42 | -0.347725837 | 0.6 | 0.844 | 8.52E-38 |
| CECR2 | 3.31E-42 | -0.385099725 | 0.3 | 0.561 | 9.57E-38 |
| CALD1 | 3.67E-42 | -0.437063685 | 0.4 | 0.596 | 1.06E-37 |
| MCAM | 3.90E-42 | -0.441148851 | 0.2 | 0.385 | 1.13E-37 |
| NALCN | 4.54E-42 | -0.334432437 | 0.6 | 0.819 | 1.31E-37 |
| BBX | 4.87E-42 | -0.321287372 | 0.5 | 0.745 | 1.41E-37 |
| KANK1 | 6.38E-42 | -0.428800869 | 0.5 | 0.72 | 1.84E-37 |
| RAP1A | 7.09E-42 | -0.271616242 | 0.4 | 0.619 | 2.05E-37 |
| FBXL7 | 7.40E-42 | -0.421524053 | 0.6 | 0.852 | 2.14E-37 |
| ADCY5 | 9.19E-42 | -0.354492974 | 0.3 | 0.492 | 2.66E-37 |
| ATM | 9.52E-42 | -0.286377105 | 0.4 | 0.637 | 2.75E-37 |
| DYNC1I2 | 1.02E-41 | -0.285789789 | 0.3 | 0.542 | 2.96E-37 |
| MTMR7 | 1.02E-41 | -0.416077062 | 0.3 | 0.514 | 2.96E-37 |

| | | | | | |
|---|---|---|---|---|---|
| CPM | 1.11E-41 | -0.487917682 | 0.3 | 0.533 | 3.20E-37 |
| CPQ | 1.53E-41 | -0.365216242 | 0.4 | 0.666 | 4.42E-37 |
| ENOX1 | 1.62E-41 | -0.372497023 | 0.6 | 0.861 | 4.68E-37 |
| TMED10 | 1.75E-41 | -0.243338414 | 0.3 | 0.475 | 5.06E-37 |
| GNAO1 | 1.79E-41 | -0.385849405 | 0.6 | 0.858 | 5.16E-37 |
| TANC1 | 1.86E-41 | -0.395849036 | 0.3 | 0.514 | 5.37E-37 |
| MT-ND4 | 2.10E-41 | 0.696832304 | 0.8 | 0.831 | 6.05E-37 |
| GATM | 2.34E-41 | -0.310694066 | 0.3 | 0.512 | 6.78E-37 |
| AP3B1 | 2.57E-41 | -0.329538277 | 0.4 | 0.637 | 7.42E-37 |
| NAPEPLD | 2.74E-41 | -0.269530381 | 0.3 | 0.563 | 7.91E-37 |
| BBS9 | 2.93E-41 | -0.401633162 | 0.4 | 0.62 | 8.46E-37 |
| ST3GAL5 | 3.45E-41 | -0.330505106 | 0.3 | 0.496 | 9.97E-37 |
| CPEB2 | 3.87E-41 | -0.32318278 | 0.4 | 0.676 | 1.12E-36 |
| PPM1H | 4.21E-41 | -0.345854695 | 0.5 | 0.757 | 1.22E-36 |
| ERBB3 | 4.27E-41 | -0.417677888 | 0.3 | 0.46 | 1.23E-36 |
| BACH2 | 4.36E-41 | -0.376925585 | 0.4 | 0.664 | 1.26E-36 |
| KHDRBS2 | 4.65E-41 | 1.124494337 | 0.3 | 0.154 | 1.34E-36 |
| SNX29 | 4.81E-41 | -0.293014319 | 0.4 | 0.623 | 1.39E-36 |
| KIZ | 5.29E-41 | -0.337008488 | 0.4 | 0.58 | 1.53E-36 |
| FAXDC2 | 6.02E-41 | -0.413317457 | 0.4 | 0.587 | 1.74E-36 |
| GRB10 | 6.43E-41 | -0.447692424 | 0.3 | 0.48 | 1.86E-36 |
| LINC00486 | 7.39E-41 | -0.198190494 | 0.4 | 0.611 | 2.14E-36 |
| OLIG1 | 1.47E-40 | -0.36569023 | 0.3 | 0.484 | 4.26E-36 |
| LINC02256 | 1.53E-40 | -0.318136677 | 0.2 | 0.438 | 4.41E-36 |
| SESTD1 | 1.64E-40 | -0.275463325 | 0.4 | 0.646 | 4.74E-36 |
| NXPE3 | 1.74E-40 | -0.328311906 | 0.3 | 0.542 | 5.04E-36 |
| CPNE2 | 2.08E-40 | -0.354425283 | 0.3 | 0.515 | 6.02E-36 |
| CACNA2D1 | 2.68E-40 | -0.345835017 | 0.6 | 0.817 | 7.74E-36 |
| LRCH3 | 2.70E-40 | -0.281188488 | 0.4 | 0.657 | 7.80E-36 |
| MIRLET7BHG | 3.06E-40 | -0.283949521 | 0.3 | 0.547 | 8.84E-36 |
| CDK17 | 3.91E-40 | -0.283650037 | 0.4 | 0.652 | 1.13E-35 |
| OTUD7B | 4.62E-40 | -0.2539345 | 0.4 | 0.66 | 1.34E-35 |
| TRIP12 | 5.83E-40 | -0.231176524 | 0.4 | 0.618 | 1.68E-35 |
| AGO3 | 5.87E-40 | -0.244921533 | 0.4 | 0.607 | 1.70E-35 |
| LMCD1-AS1 | 6.06E-40 | -0.533690023 | 0.4 | 0.632 | 1.75E-35 |
| AC090502.1 | 6.80E-40 | -0.567044666 | 0.3 | 0.468 | 1.97E-35 |
| SDCCAG8 | 7.10E-40 | -0.300729956 | 0.5 | 0.723 | 2.05E-35 |
| GPBP1 | 7.80E-40 | -0.244124837 | 0.4 | 0.662 | 2.25E-35 |
| BBS2 | 7.95E-40 | -0.25600474 | 0.4 | 0.614 | 2.30E-35 |
| RAP1GDS1 | 9.67E-40 | -0.25343175 | 0.5 | 0.695 | 2.79E-35 |
| RECK | 9.82E-40 | -0.377878733 | 0.2 | 0.446 | 2.84E-35 |
| ST8SIA1 | 9.93E-40 | 0.533786308 | 0.1 | 0.051 | 2.87E-35 |
| SF1 | 9.97E-40 | -0.226331961 | 0.4 | 0.591 | 2.88E-35 |
| PEAK1 | 1.13E-39 | -0.294408781 | 0.4 | 0.673 | 3.26E-35 |
| PHC3 | 1.22E-39 | -0.244585206 | 0.4 | 0.603 | 3.52E-35 |
| FBN1 | 1.42E-39 | -0.390533584 | 0.3 | 0.521 | 4.09E-35 |
| ATG13 | 1.52E-39 | -0.307563803 | 0.3 | 0.505 | 4.40E-35 |
| TAOK1 | 1.71E-39 | -0.178033277 | 0.5 | 0.696 | 4.95E-35 |
| ANKRD28 | 1.72E-39 | -0.288934373 | 0.5 | 0.756 | 4.96E-35 |
| MARK3 | 1.75E-39 | -0.197486928 | 0.4 | 0.677 | 5.05E-35 |
| MXI1 | 1.79E-39 | -0.325452706 | 0.5 | 0.695 | 5.17E-35 |
| FKBP5 | 2.08E-39 | -0.403103087 | 0.5 | 0.682 | 6.02E-35 |
| HTRA1 | 2.23E-39 | -0.290036519 | 0.4 | 0.659 | 6.43E-35 |
| ACYP2 | 2.60E-39 | -0.27756056 | 0.5 | 0.732 | 7.51E-35 |
| CBFB | 2.77E-39 | -0.359601053 | 0.3 | 0.529 | 7.99E-35 |
| UBR3 | 3.18E-39 | -0.227256033 | 0.5 | 0.756 | 9.20E-35 |
| PRMT2 | 3.35E-39 | -0.266407496 | 0.3 | 0.503 | 9.67E-35 |
| PALM2-AKAP2 | 3.38E-39 | -0.460758782 | 0.3 | 0.543 | 9.78E-35 |
| HEPACAM | 3.52E-39 | -0.299213292 | 0.4 | 0.627 | 1.02E-34 |
| KDM4C | 3.89E-39 | -0.212152928 | 0.5 | 0.76 | 1.13E-34 |
| GIGYF2 | 4.05E-39 | -0.243489428 | 0.4 | 0.614 | 1.17E-34 |
| YWHAQ | 4.13E-39 | -0.25828562 | 0.4 | 0.583 | 1.19E-34 |
| SNX9 | 4.99E-39 | -0.232493418 | 0.4 | 0.604 | 1.44E-34 |
| HIP1R | 5.10E-39 | -0.261300655 | 0.4 | 0.585 | 1.47E-34 |
| ENDOD1 | 5.27E-39 | -0.307485217 | 0.2 | 0.436 | 1.52E-34 |
| HSPA2 | 5.57E-39 | -0.336827447 | 0.3 | 0.465 | 1.61E-34 |
| MTMR10 | 5.63E-39 | -0.364395708 | 0.3 | 0.489 | 1.63E-34 |
| LSS | 5.69E-39 | -0.285225638 | 0.3 | 0.551 | 1.64E-34 |
| AC011287.1 | 6.31E-39 | 0.748936169 | 0.1 | 0.032 | 1.82E-34 |
| ZNF207 | 6.39E-39 | -0.204973408 | 0.4 | 0.592 | 1.85E-34 |
| RASSF2 | 8.27E-39 | -0.328297129 | 0.3 | 0.46 | 2.39E-34 |
| WDFY2 | 9.92E-39 | -0.284625596 | 0.4 | 0.689 | 2.87E-34 |
| EFHD1 | 1.29E-38 | -0.330016479 | 0.4 | 0.643 | 3.74E-34 |
| ABHD17B | 1.46E-38 | -0.290487334 | 0.3 | 0.565 | 4.21E-34 |
| RALGPS1 | 1.61E-38 | -0.334107725 | 0.3 | 0.526 | 4.66E-34 |
| NLK | 1.92E-38 | -0.28630068 | 0.5 | 0.76 | 5.54E-34 |
| SLC5A11 | 2.78E-38 | -0.564380399 | 0.3 | 0.552 | 8.04E-34 |
| B3GAT1 | 3.27E-38 | -0.262098803 | 0.3 | 0.536 | 9.45E-34 |
| REEP3 | 3.43E-38 | -0.323428665 | 0.3 | 0.536 | 9.91E-34 |
| TLE4 | 4.28E-38 | -0.315491232 | 0.5 | 0.682 | 1.24E-33 |
| TMEM87A | 5.43E-38 | -0.325969902 | 0.3 | 0.562 | 1.57E-33 |
| CDK5RAP2 | 7.33E-38 | -0.274068788 | 0.3 | 0.572 | 2.12E-33 |
| HIBCH | 7.38E-38 | -0.256099168 | 0.4 | 0.681 | 2.13E-33 |
| MAPK10 | 7.74E-38 | 1.041950616 | 0.6 | 0.517 | 2.24E-33 |
| AC009063.2 | 8.06E-38 | -0.43305734 | 0.3 | 0.53 | 2.33E-33 |
| ZBED5 | 8.54E-38 | -0.298496076 | 0.3 | 0.543 | 2.47E-33 |
| SPTLC2 | 8.62E-38 | -0.278265459 | 0.4 | 0.581 | 2.49E-33 |
| PCA3 | 9.10E-38 | -0.420665793 | 0.1 | 0.305 | 2.63E-33 |
| AIF1L | 1.04E-37 | -0.420485914 | 0.1 | 0.288 | 2.99E-33 |
| AFF3 | 1.04E-37 | 0.676339647 | 0.2 | 0.075 | 3.02E-33 |
| WLS | 1.08E-37 | 0.834021047 | 0.1 | 0.046 | 3.12E-33 |
| FAM126A | 1.17E-37 | -0.438288766 | 0.3 | 0.45 | 3.39E-33 |
| OLMALINC | 1.27E-37 | -0.425681047 | 0.3 | 0.471 | 3.66E-33 |
| LCOR | 1.45E-37 | -0.246421783 | 0.4 | 0.663 | 4.18E-33 |
| SLTM | 1.57E-37 | -0.229549798 | 0.3 | 0.541 | 4.55E-33 |
| ZNRF3 | 1.60E-37 | 1.091045839 | 0.2 | 0.104 | 4.61E-33 |
| ELOVL5 | 1.61E-37 | -0.2151546 | 0.3 | 0.501 | 4.66E-33 |
| LRP4 | 1.63E-37 | -0.219497037 | 0.4 | 0.6 | 4.71E-33 |
| CRTC3 | 2.20E-37 | -0.259614705 | 0.4 | 0.605 | 6.36E-33 |
| ARHGAP1 | 2.26E-37 | -0.31784265 | 0.3 | 0.523 | 6.54E-33 |
| TENT2 | 2.42E-37 | -0.253448407 | 0.4 | 0.619 | 7.00E-33 |
| PTPN13 | 2.53E-37 | -0.380629209 | 0.4 | 0.557 | 7.31E-33 |
| L3MBTL4 | 2.59E-37 | -0.335819441 | 0.5 | 0.704 | 7.50E-33 |
| IKZF2 | 2.63E-37 | -0.333508156 | 0.3 | 0.557 | 7.60E-33 |
| VMP1 | 2.65E-37 | -0.263487735 | 0.3 | 0.549 | 7.66E-33 |
| EEF1A2 | 2.83E-37 | 0.990303909 | 0.1 | 0.039 | 8.19E-33 |
| CORO1C | 3.31E-37 | -0.405257716 | 0.2 | 0.435 | 9.57E-33 |
| UBE2E2 | 3.38E-37 | -0.353714243 | 0.6 | 0.833 | 9.77E-33 |
| SREBF2 | 3.62E-37 | -0.182139896 | 0.4 | 0.642 | 1.05E-32 |
| POLK | 3.62E-37 | -0.256890777 | 0.3 | 0.531 | 1.05E-32 |
| ENOX2 | 3.66E-37 | -0.425291738 | 0.3 | 0.527 | 1.06E-32 |
| TPPP | 4.87E-37 | -0.185216071 | 0.4 | 0.603 | 1.41E-32 |
| C1GALT1 | 5.35E-37 | -0.36539703 | 0.2 | 0.445 | 1.55E-32 |
| CYP27A1 | 5.79E-37 | -0.332965762 | 0.2 | 0.44 | 1.67E-32 |
| RTN4 | 5.80E-37 | -0.270370002 | 0.7 | 0.93 | 1.68E-32 |
| FCHSD2 | 6.73E-37 | -0.435644606 | 0.5 | 0.731 | 1.95E-32 |
| NDE1 | 7.93E-37 | -0.371538025 | 0.3 | 0.506 | 2.29E-32 |
| KCND3 | 9.36E-37 | 0.773711236 | 0.2 | 0.066 | 2.70E-32 |
| KLF12 | 9.49E-37 | 0.611462318 | 0.1 | 0.049 | 2.74E-32 |
| PLEKHB1 | 1.10E-36 | -0.311537872 | 0.3 | 0.504 | 3.18E-32 |
| ARGLU1 | 1.17E-36 | -0.140315378 | 0.5 | 0.781 | 3.38E-32 |
| SYNDIG1 | 1.28E-36 | -0.331810654 | 0.4 | 0.586 | 3.69E-32 |
| PTN | 1.36E-36 | -0.342170095 | 0.3 | 0.501 | 3.92E-32 |
| ITGA2 | 1.62E-36 | -0.446809853 | 0.3 | 0.535 | 4.69E-32 |
| RAB30 | 1.79E-36 | -0.302264217 | 0.4 | 0.595 | 5.16E-32 |
| VWA1 | 1.83E-36 | -0.312074825 | 0.2 | 0.395 | 5.30E-32 |
| KIDINS220 | 2.12E-36 | -0.187510757 | 0.5 | 0.74 | 6.14E-32 |
| LAMP1 | 2.45E-36 | -0.254337153 | 0.3 | 0.473 | 7.08E-32 |
| MGAT5 | 2.49E-36 | -0.39892069 | 0.4 | 0.587 | 7.21E-32 |
| KAT6A | 2.50E-36 | -0.223309129 | 0.4 | 0.605 | 7.23E-32 |
| VPS13B | 2.64E-36 | -0.186588996 | 0.4 | 0.66 | 7.62E-32 |
| PTMA | 2.72E-36 | -0.20050441 | 0.5 | 0.716 | 7.86E-32 |
| COL9A2 | 3.03E-36 | -0.360809368 | 0.2 | 0.367 | 8.77E-32 |
| RIF1 | 3.10E-36 | -0.190083266 | 0.4 | 0.58 | 8.95E-32 |
| BACE1 | 3.17E-36 | -0.231226871 | 0.2 | 0.437 | 9.17E-32 |
| CALY | 4.01E-36 | 0.86645556 | 0.1 | 0.035 | 1.16E-31 |
| ZDHHC11B | 4.72E-36 | -0.318373037 | 0.3 | 0.487 | 1.36E-31 |
| OSBP2 | 4.84E-36 | -0.371562672 | 0.4 | 0.612 | 1.40E-31 |

| | | | | | |
|---|---|---|---|---|---|
| RTKN | 4.86E-36 | -0.380973675 | 0.2 | 0.419 | 1.40E-31 |
| GATAD2B | 5.11E-36 | -0.232040284 | 0.4 | 0.591 | 1.48E-31 |
| GTDC1 | 5.28E-36 | -0.316828369 | 0.4 | 0.674 | 1.53E-31 |
| CDK13 | 5.86E-36 | -0.176743863 | 0.4 | 0.668 | 1.69E-31 |
| SLC11A2 | 7.18E-36 | -0.217858849 | 0.3 | 0.474 | 2.08E-31 |
| SNX30 | 8.56E-36 | -0.322115501 | 0.3 | 0.545 | 2.48E-31 |
| TRA2A | 9.11E-36 | -0.248715732 | 0.5 | 0.703 | 2.63E-31 |
| PHACTR3 | 1.01E-35 | -0.310550132 | 0.6 | 0.804 | 2.91E-31 |
| SENP6 | 1.05E-35 | -0.185441799 | 0.4 | 0.683 | 3.02E-31 |
| TMCC3 | 1.14E-35 | -0.365925509 | 0.2 | 0.411 | 3.28E-31 |
| KIF2A | 1.29E-35 | -0.231193102 | 0.4 | 0.582 | 3.74E-31 |
| NCOA6 | 1.34E-35 | -0.269837236 | 0.4 | 0.565 | 3.88E-31 |
| LDB2 | 1.49E-35 | 0.928370058 | 0.2 | 0.106 | 4.30E-31 |
| AC007566.1 | 1.52E-35 | -0.311673032 | 0.3 | 0.461 | 4.40E-31 |
| LRRC63 | 1.60E-35 | -0.48521537 | 0.2 | 0.419 | 4.63E-31 |
| ZNF98 | 2.02E-35 | 0.680573386 | 0.1 | 0.045 | 5.84E-31 |
| MKRN3 | 2.76E-35 | -0.306240249 | 0.3 | 0.524 | 7.98E-31 |
| AMD1 | 3.12E-35 | -0.318912148 | 0.4 | 0.565 | 9.02E-31 |
| CD55 | 3.69E-35 | -0.338022117 | 0.3 | 0.468 | 1.07E-30 |
| APPBP2 | 3.71E-35 | -0.231875683 | 0.3 | 0.567 | 1.07E-30 |
| SVEP1 | 3.90E-35 | -0.56436145 | 0.2 | 0.421 | 1.13E-30 |
| ARHGAP22 | 4.48E-35 | -0.404742437 | 0.2 | 0.408 | 1.29E-30 |
| PRIMA1 | 4.98E-35 | -0.416970782 | 0.2 | 0.338 | 1.44E-30 |
| CLU | 5.79E-35 | 1.624909251 | 0.5 | 0.361 | 1.67E-30 |
| PBRM1 | 5.98E-35 | -0.251471055 | 0.3 | 0.53 | 1.73E-30 |
| SPP1 | 6.55E-35 | -0.173215598 | 0.5 | 0.728 | 1.89E-30 |
| TLK2 | 8.10E-35 | -0.285951657 | 0.3 | 0.481 | 2.34E-30 |
| NINJ2 | 8.11E-35 | -0.431391246 | 0.3 | 0.467 | 2.34E-30 |
| RPS6KA5 | 8.83E-35 | -0.260465041 | 0.4 | 0.567 | 2.55E-30 |
| STXBP1 | 1.28E-34 | 1.338008631 | 0.3 | 0.155 | 3.69E-30 |
| RPTOR | 1.48E-34 | -0.221917553 | 0.3 | 0.499 | 4.27E-30 |
| NUDT4 | 1.61E-34 | 0.697725584 | 0.1 | 0.037 | 4.65E-30 |
| CLIP4 | 1.99E-34 | -0.229423807 | 0.4 | 0.647 | 5.75E-30 |
| MYO5A | 2.12E-34 | -0.177473656 | 0.5 | 0.729 | 6.14E-30 |
| HP1BP3 | 2.23E-34 | -0.168748828 | 0.4 | 0.685 | 6.43E-30 |
| MPDZ | 2.45E-34 | -0.237139071 | 0.4 | 0.654 | 7.08E-30 |
| AGO2 | 2.96E-34 | -0.229924584 | 0.3 | 0.548 | 8.55E-30 |
| PAXBP1 | 3.11E-34 | -0.252827049 | 0.4 | 0.565 | 8.98E-30 |
| URI1 | 4.53E-34 | -0.249614484 | 0.3 | 0.512 | 1.31E-29 |
| ARID2 | 5.20E-34 | -0.277644222 | 0.4 | 0.583 | 1.50E-29 |
| SHPRH | 5.48E-34 | -0.250262109 | 0.4 | 0.568 | 1.58E-29 |
| EMC2 | 5.62E-34 | -0.247726268 | 0.2 | 0.426 | 1.63E-29 |
| IBTK | 5.92E-34 | -0.242981099 | 0.3 | 0.472 | 1.71E-29 |
| ZNF532 | 6.03E-34 | -0.195428961 | 0.4 | 0.624 | 1.74E-29 |
| DNM2 | 6.50E-34 | -0.286857528 | 0.3 | 0.496 | 1.88E-29 |
| ANKRD10 | 6.87E-34 | -0.137895062 | 0.4 | 0.566 | 1.99E-29 |
| KRIT1 | 7.08E-34 | -0.257880025 | 0.3 | 0.457 | 2.05E-29 |
| PHIP | 7.20E-34 | -0.247880267 | 0.5 | 0.77 | 2.08E-29 |
| FCHO1 | 7.60E-34 | -0.326330211 | 0.2 | 0.392 | 2.20E-29 |
| WHRN | 7.77E-34 | -0.332616547 | 0.3 | 0.451 | 2.25E-29 |
| CDC42BPA | 8.64E-34 | -0.300020369 | 0.7 | 0.93 | 2.50E-29 |
| APP | 1.01E-33 | -0.179150906 | 0.7 | 0.913 | 2.91E-29 |
| SPPL3 | 1.08E-33 | -0.217128091 | 0.4 | 0.658 | 3.11E-29 |
| HDAC4 | 1.08E-33 | -0.240912212 | 0.4 | 0.612 | 3.12E-29 |
| KMT2C | 1.12E-33 | -0.224590122 | 0.6 | 0.843 | 3.24E-29 |
| DMXL1 | 1.36E-33 | -0.206171837 | 0.4 | 0.586 | 3.94E-29 |
| SEC14L5 | 1.48E-33 | -0.377820201 | 0.3 | 0.459 | 4.27E-29 |
| GARNL3 | 1.49E-33 | -0.221173272 | 0.4 | 0.599 | 4.30E-29 |
| MYO9B | 1.58E-33 | -0.2459292 | 0.4 | 0.583 | 4.55E-29 |
| DMTF1 | 1.60E-33 | -0.163426103 | 0.4 | 0.577 | 4.63E-29 |
| FTO | 1.69E-33 | -0.242833791 | 0.5 | 0.745 | 4.90E-29 |
| NASP | 1.76E-33 | -0.252741817 | 0.3 | 0.541 | 5.10E-29 |
| AP002370.2 | 1.87E-33 | -0.354854154 | 0.1 | 0.296 | 5.40E-29 |
| SLC38A2 | 1.99E-33 | -0.231123442 | 0.3 | 0.524 | 5.74E-29 |
| KIF1B | 2.11E-33 | -0.268418734 | 0.6 | 0.894 | 6.10E-29 |
| ZMYM2 | 2.30E-33 | -0.216093602 | 0.5 | 0.717 | 6.64E-29 |
| RAB30-AS1 | 2.40E-33 | -0.229027783 | 0.3 | 0.497 | 6.93E-29 |
| SIK2 | 2.47E-33 | -0.305503901 | 0.4 | 0.609 | 7.15E-29 |
| AC106791.1 | 2.50E-33 | -0.386078763 | 0.3 | 0.468 | 7.24E-29 |
| TSPAN5 | 2.64E-33 | -0.311756492 | 0.5 | 0.764 | 7.62E-29 |
| ARHGEF2 | 2.76E-33 | -0.235390738 | 0.3 | 0.506 | 7.97E-29 |
| IWS1 | 2.76E-33 | -0.204416794 | 0.3 | 0.516 | 7.97E-29 |
| COPA | 2.93E-33 | -0.187942477 | 0.4 | 0.591 | 8.48E-29 |
| SLC35F5 | 3.00E-33 | -0.211547525 | 0.2 | 0.417 | 8.66E-29 |
| MTMR2 | 3.05E-33 | -0.322504892 | 0.3 | 0.507 | 8.82E-29 |
| JMJD1C | 3.34E-33 | -0.212225527 | 0.6 | 0.83 | 9.65E-29 |
| FGF1 | 3.42E-33 | -0.311115329 | 0.3 | 0.508 | 9.88E-29 |
| SYNE1 | 3.48E-33 | 1.277951107 | 0.5 | 0.395 | 1.00E-28 |
| ZNF397 | 3.93E-33 | -0.295791757 | 0.3 | 0.477 | 1.14E-28 |
| SLC2A13 | 4.03E-33 | 0.891623978 | 0.2 | 0.097 | 1.17E-28 |
| WASF2 | 4.11E-33 | -0.268914917 | 0.3 | 0.449 | 1.19E-28 |
| SPG11 | 4.15E-33 | -0.220603036 | 0.3 | 0.537 | 1.20E-28 |
| LARP6 | 5.04E-33 | -0.281780396 | 0.2 | 0.423 | 1.46E-28 |
| TMEM123 | 5.47E-33 | -0.366808388 | 0.2 | 0.429 | 1.58E-28 |
| MBOAT2 | 5.57E-33 | -0.174701143 | 0.4 | 0.643 | 1.61E-28 |
| FTH1 | 5.70E-33 | -0.322916719 | 0.6 | 0.81 | 1.65E-28 |
| ABHD6 | 5.81E-33 | -0.301541253 | 0.2 | 0.43 | 1.68E-28 |
| DYSF | 6.10E-33 | -0.356590741 | 0.4 | 0.557 | 1.76E-28 |
| UBE2H | 6.71E-33 | -0.198748887 | 0.3 | 0.493 | 1.94E-28 |
| NEK7 | 8.59E-33 | -0.289237163 | 0.3 | 0.532 | 2.48E-28 |
| TOM1L2 | 1.09E-32 | -0.238731994 | 0.5 | 0.693 | 3.14E-28 |
| SLC9A9 | 1.13E-32 | -0.26808058 | 0.5 | 0.723 | 3.26E-28 |
| USP40 | 1.45E-32 | -0.260025262 | 0.3 | 0.489 | 4.20E-28 |
| FBXO7 | 1.57E-32 | -0.254096627 | 0.3 | 0.45 | 4.53E-28 |
| TTC37 | 1.79E-32 | -0.160848704 | 0.3 | 0.526 | 5.18E-28 |
| SOX8 | 2.38E-32 | -0.310216642 | 0.2 | 0.417 | 6.86E-28 |
| LRRC8B | 2.55E-32 | -0.199775501 | 0.3 | 0.505 | 7.36E-28 |
| KCNJ2 | 3.02E-32 | -0.383932956 | 0.2 | 0.417 | 8.72E-28 |
| PHYKPL | 3.05E-32 | -0.215357896 | 0.2 | 0.345 | 8.82E-28 |
| STARD9 | 3.26E-32 | -0.286768083 | 0.3 | 0.486 | 9.42E-28 |
| FTCDNL1 | 3.73E-32 | -0.375673973 | 0.2 | 0.348 | 1.08E-27 |
| ANKRD17 | 4.73E-32 | -0.222565617 | 0.5 | 0.704 | 1.37E-27 |
| N4BP2L2 | 4.94E-32 | -0.195710059 | 0.6 | 0.875 | 1.43E-27 |
| INPP4B | 5.66E-32 | 0.676936794 | 0.2 | 0.12 | 1.63E-27 |
| MAT2A | 5.72E-32 | -0.190440393 | 0.4 | 0.586 | 1.65E-27 |
| FNBP4 | 6.32E-32 | -0.194493105 | 0.3 | 0.551 | 1.83E-27 |
| ATF7IP | 6.68E-32 | -0.224783036 | 0.3 | 0.505 | 1.93E-27 |
| ADD3 | 6.99E-32 | -0.214141815 | 0.5 | 0.747 | 2.02E-27 |
| NFE2L2 | 7.79E-32 | -0.197358909 | 0.4 | 0.651 | 2.25E-27 |
| EXOC1 | 8.07E-32 | -0.239619163 | 0.3 | 0.458 | 2.33E-27 |
| LINC01792 | 8.53E-32 | -0.398564256 | 0.2 | 0.403 | 2.47E-27 |
| GATAD1 | 8.61E-32 | -0.254794431 | 0.3 | 0.448 | 2.49E-27 |
| ANKH | 9.51E-32 | -0.232348819 | 0.4 | 0.572 | 2.75E-27 |
| NFIX | 9.97E-32 | -0.149689005 | 0.5 | 0.683 | 2.88E-27 |
| BCL2L1 | 1.04E-31 | -0.248901923 | 0.3 | 0.486 | 3.02E-27 |
| RBM5 | 1.37E-31 | -0.226884483 | 0.3 | 0.515 | 3.96E-27 |
| STAG1 | 1.45E-31 | -0.272833805 | 0.4 | 0.667 | 4.19E-27 |
| RHOBTB1 | 1.52E-31 | -0.358245619 | 0.3 | 0.437 | 4.40E-27 |
| DNAJB2 | 1.53E-31 | -0.182713067 | 0.3 | 0.518 | 4.41E-27 |
| ZNF451 | 1.53E-31 | -0.202643304 | 0.4 | 0.627 | 4.42E-27 |
| NEK3 | 1.79E-31 | -0.312451473 | 0.2 | 0.408 | 5.18E-27 |
| ACSS2 | 1.96E-31 | -0.260236082 | 0.2 | 0.402 | 5.68E-27 |
| FARP2 | 2.30E-31 | -0.271822127 | 0.3 | 0.459 | 6.64E-27 |
| CDC42SE2 | 2.41E-31 | -0.230192886 | 0.3 | 0.542 | 6.97E-27 |
| SLC38A9 | 2.49E-31 | -0.317642244 | 0.3 | 0.47 | 7.18E-27 |
| MITF | 2.69E-31 | -0.381587157 | 0.3 | 0.48 | 7.78E-27 |
| CEP170 | 2.82E-31 | -0.180052144 | 0.4 | 0.658 | 8.16E-27 |
| PHYHIPL | 2.92E-31 | -0.278793005 | 0.5 | 0.672 | 8.44E-27 |
| WDR70 | 3.41E-31 | -0.160699478 | 0.4 | 0.629 | 9.84E-27 |
| AC004158.1 | 3.41E-31 | -0.39543127 | 0.3 | 0.431 | 9.87E-27 |
| PHACTR4 | 3.56E-31 | -0.215281707 | 0.4 | 0.602 | 1.03E-26 |
| FHIT | 4.09E-31 | -0.379749961 | 0.6 | 0.79 | 1.18E-26 |
| ABL1 | 4.13E-31 | -0.274384436 | 0.3 | 0.479 | 1.19E-26 |
| PRR18 | 4.44E-31 | -0.277115587 | 0.2 | 0.403 | 1.28E-26 |
| DIP2A | 4.53E-31 | -0.249643276 | 0.3 | 0.511 | 1.31E-26 |
| FNDC3A | 4.56E-31 | -0.202622433 | 0.3 | 0.542 | 1.32E-26 |
| SH3PXD2A | 5.04E-31 | -0.212081063 | 0.2 | 0.419 | 1.46E-26 |
| USP48 | 6.04E-31 | -0.208836426 | 0.3 | 0.504 | 1.75E-26 |

| | | | | | |
|---|---|---|---|---|---|
| AKAP12 | 6.43E-31 | 0.54772291 | 0.1 | 0.036 | 1.86E-26 |
| GALNT13 | 7.53E-31 | -0.37552182 | 0.4 | 0.575 | 2.18E-26 |
| ITSN2 | 8.09E-31 | -0.181637724 | 0.4 | 0.585 | 2.34E-26 |
| UNC5B | 8.33E-31 | -0.299575243 | 0.2 | 0.389 | 2.41E-26 |
| HBP1 | 8.41E-31 | -0.215377964 | 0.2 | 0.418 | 2.43E-26 |
| MTSS1 | 8.70E-31 | -0.295042992 | 0.3 | 0.522 | 2.51E-26 |
| RFC1 | 9.95E-31 | -0.235857317 | 0.2 | 0.42 | 2.88E-26 |
| KLF13 | 1.06E-30 | -0.182919086 | 0.3 | 0.442 | 3.08E-26 |
| ZNF609 | 1.09E-30 | -0.199138861 | 0.4 | 0.648 | 3.15E-26 |
| THOC2 | 1.16E-30 | -0.207754937 | 0.3 | 0.496 | 3.35E-26 |
| MAN2A2 | 1.36E-30 | -0.249143854 | 0.2 | 0.374 | 3.94E-26 |
| SMURF1 | 1.40E-30 | -0.322671851 | 0.3 | 0.516 | 4.04E-26 |
| DIXDC1 | 1.47E-30 | -0.211209682 | 0.4 | 0.566 | 4.24E-26 |
| DEPTOR | 1.62E-30 | -0.302254954 | 0.3 | 0.492 | 4.67E-26 |
| ZFAND3 | 1.86E-30 | -0.173392749 | 0.5 | 0.727 | 5.38E-26 |
| EEPD1 | 1.97E-30 | -0.371972921 | 0.2 | 0.404 | 5.70E-26 |
| UBR2 | 2.21E-30 | -0.145491577 | 0.3 | 0.532 | 6.40E-26 |
| SLC44A5 | 2.30E-30 | 0.698208119 | 0.1 | 0.046 | 6.66E-26 |
| NIPAL3 | 2.38E-30 | -0.24127982 | 0.3 | 0.467 | 6.89E-26 |
| HPN-AS1 | 2.43E-30 | -0.309251629 | 0.3 | 0.456 | 7.02E-26 |
| ACOT11 | 2.58E-30 | 0.634234869 | 0.1 | 0.045 | 7.45E-26 |
| ACTN4 | 2.65E-30 | -0.189281281 | 0.3 | 0.543 | 7.66E-26 |
| MICAL3 | 2.74E-30 | -0.205211037 | 0.5 | 0.733 | 7.93E-26 |
| ADSS | 3.12E-30 | -0.282714163 | 0.3 | 0.425 | 9.02E-26 |
| XPO6 | 3.13E-30 | -0.241267248 | 0.2 | 0.415 | 9.04E-26 |
| INPP1 | 3.39E-30 | -0.214383807 | 0.2 | 0.391 | 9.79E-26 |
| PKN2 | 3.80E-30 | -0.187763663 | 0.4 | 0.555 | 1.10E-25 |
| PPP4R2 | 4.16E-30 | -0.187265137 | 0.3 | 0.529 | 1.20E-25 |
| CERS2 | 4.53E-30 | -0.292787496 | 0.1 | 0.285 | 1.31E-25 |
| NAV1 | 4.55E-30 | -0.203314915 | 0.5 | 0.743 | 1.31E-25 |
| FHOD3 | 4.66E-30 | 0.462291898 | 0.1 | 0.036 | 1.35E-25 |
| GPSM2 | 5.02E-30 | -0.258602923 | 0.3 | 0.493 | 1.45E-25 |
| FXR1 | 5.20E-30 | -0.209704252 | 0.3 | 0.529 | 1.50E-25 |
| SLC48A1 | 5.32E-30 | -0.200842275 | 0.3 | 0.467 | 1.54E-25 |
| KIAA1755 | 5.98E-30 | -0.247396149 | 0.4 | 0.582 | 1.73E-25 |
| GOLGA7 | 6.36E-30 | -0.284424684 | 0.2 | 0.409 | 1.84E-25 |
| GCLC | 6.53E-30 | -0.336528689 | 0.2 | 0.404 | 1.89E-25 |
| INF2 | 7.01E-30 | -0.195960298 | 0.3 | 0.453 | 2.03E-25 |
| PHC2 | 7.02E-30 | -0.243984665 | 0.3 | 0.472 | 2.03E-25 |
| BEST1 | 7.48E-30 | -0.303302763 | 0.2 | 0.338 | 2.16E-25 |
| BTBD7 | 7.49E-30 | -0.232280149 | 0.3 | 0.502 | 2.16E-25 |
| STK3 | 8.20E-30 | -0.1737111 | 0.4 | 0.629 | 2.37E-25 |
| MFF | 9.24E-30 | -0.155100053 | 0.3 | 0.44 | 2.67E-25 |
| ELOVL7 | 9.95E-30 | -0.42958966 | 0.2 | 0.318 | 2.88E-25 |
| CPSF6 | 9.97E-30 | -0.1750454 | 0.3 | 0.54 | 2.88E-25 |
| UBA6 | 9.98E-30 | -0.157462686 | 0.2 | 0.427 | 2.88E-25 |
| AASS | 1.09E-29 | -0.185434999 | 0.2 | 0.34 | 3.15E-25 |
| ANKRD12 | 1.21E-29 | -0.156357191 | 0.6 | 0.799 | 3.50E-25 |
| WDFY3 | 1.30E-29 | -0.208444907 | 0.6 | 0.802 | 3.76E-25 |
| MDM4 | 1.38E-29 | -0.170525452 | 0.3 | 0.56 | 3.98E-25 |
| SPART | 1.45E-29 | -0.141778201 | 0.3 | 0.54 | 4.19E-25 |
| ACSL1 | 1.46E-29 | -0.194469573 | 0.2 | 0.374 | 4.22E-25 |
| TBC1D2 | 1.49E-29 | -0.330095583 | 0.2 | 0.341 | 4.30E-25 |
| ECPAS | 1.69E-29 | -0.174270421 | 0.4 | 0.59 | 4.87E-25 |
| PDE4D | 1.71E-29 | 0.846537077 | 0.6 | 0.454 | 4.94E-25 |
| ABCG1 | 1.77E-29 | -0.340310798 | 0.2 | 0.399 | 5.11E-25 |
| ARL17B | 1.93E-29 | -0.236378192 | 0.2 | 0.385 | 5.59E-25 |
| UBXN4 | 2.21E-29 | -0.143849762 | 0.3 | 0.475 | 6.38E-25 |
| PIGU | 2.31E-29 | -0.251296444 | 0.2 | 0.401 | 6.69E-25 |
| CREB3L2 | 2.32E-29 | -0.329518457 | 0.2 | 0.376 | 6.72E-25 |
| EVI5L | 2.52E-29 | -0.211339423 | 0.2 | 0.409 | 7.28E-25 |
| ROCK1 | 2.57E-29 | -0.191848256 | 0.3 | 0.508 | 7.44E-25 |
| HSD17B4 | 2.90E-29 | -0.195134664 | 0.3 | 0.544 | 8.38E-25 |
| HS3ST4 | 3.12E-29 | 0.688004676 | 0.1 | 0.039 | 9.02E-25 |
| SRPK2 | 3.44E-29 | -0.216491793 | 0.6 | 0.842 | 9.94E-25 |
| MEIS1 | 3.45E-29 | -0.306511645 | 0.3 | 0.438 | 9.98E-25 |
| ATE1 | 3.92E-29 | -0.204828767 | 0.4 | 0.623 | 1.13E-24 |
| FUS | 4.08E-29 | -0.178366817 | 0.3 | 0.522 | 1.18E-24 |
| WWP1 | 4.47E-29 | -0.206422069 | 0.3 | 0.495 | 1.29E-24 |
| ATP11B | 4.56E-29 | -0.221341039 | 0.3 | 0.461 | 1.32E-24 |
| SCAF8 | 4.72E-29 | -0.174746611 | 0.3 | 0.501 | 1.36E-24 |
| ZC3H6 | 4.87E-29 | -0.286688785 | 0.3 | 0.489 | 1.41E-24 |
| AK5 | 5.11E-29 | -0.288359418 | 0.6 | 0.806 | 1.48E-24 |
| PDE7A | 5.81E-29 | -0.227456283 | 0.3 | 0.531 | 1.68E-24 |
| RCAN2 | 5.94E-29 | -0.247941863 | 0.4 | 0.594 | 1.72E-24 |
| LGI3 | 6.25E-29 | -0.178869819 | 0.2 | 0.401 | 1.81E-24 |
| DIS3L2 | 6.35E-29 | -0.23122822 | 0.4 | 0.562 | 1.83E-24 |
| JAKMIP3 | 7.01E-29 | -0.298067455 | 0.2 | 0.385 | 2.03E-24 |
| FDFT1 | 7.63E-29 | -0.196517903 | 0.3 | 0.482 | 2.20E-24 |
| CDC42BPB | 7.96E-29 | -0.145466141 | 0.3 | 0.524 | 2.30E-24 |
| CHD1L | 8.12E-29 | -0.265098272 | 0.2 | 0.373 | 2.35E-24 |
| CTBP2 | 8.37E-29 | -0.268796919 | 0.2 | 0.418 | 2.42E-24 |
| LITAF | 9.66E-29 | -0.339445512 | 0.2 | 0.31 | 2.79E-24 |
| ARIH2 | 1.26E-28 | -0.172417873 | 0.3 | 0.525 | 3.64E-24 |
| INSR | 1.42E-28 | -0.21497545 | 0.4 | 0.63 | 4.10E-24 |
| ACSL4 | 1.50E-28 | -0.162792171 | 0.2 | 0.417 | 4.34E-24 |
| SEMA3C | 1.59E-28 | -0.438801993 | 0.2 | 0.342 | 4.61E-24 |
| PCCA | 1.66E-28 | -0.205707428 | 0.4 | 0.608 | 4.80E-24 |
| TMPRSS5 | 1.74E-28 | -0.397228455 | 0.2 | 0.306 | 5.03E-24 |
| ETNK1 | 1.80E-28 | -0.150172766 | 0.3 | 0.496 | 5.21E-24 |
| PAN3 | 1.86E-28 | -0.183876963 | 0.4 | 0.594 | 5.36E-24 |
| VEZF1 | 1.86E-28 | -0.275941507 | 0.2 | 0.407 | 5.38E-24 |
| SLC20A2 | 2.28E-28 | -0.159186831 | 0.4 | 0.561 | 6.59E-24 |
| LGR5 | 2.53E-28 | -0.42051088 | 0.2 | 0.366 | 7.32E-24 |
| PLXDC2 | 2.67E-28 | -0.296298251 | 0.6 | 0.832 | 7.71E-24 |
| CADM2 | 2.67E-28 | -0.335258504 | 0.9 | 0.961 | 7.73E-24 |
| C9orf85 | 3.27E-28 | -0.284476038 | 0.3 | 0.443 | 9.44E-24 |
| XRRA1 | 3.33E-28 | -0.251693566 | 0.2 | 0.413 | 9.62E-24 |
| CDH20 | 3.53E-28 | -0.315255461 | 0.6 | 0.873 | 1.02E-23 |
| COMMD10 | 3.56E-28 | -0.294832836 | 0.3 | 0.492 | 1.03E-23 |
| MT-ND1 | 3.83E-28 | 0.976733424 | 0.6 | 0.625 | 1.11E-23 |
| FAM222A | 3.98E-28 | -0.338984306 | 0.2 | 0.373 | 1.15E-23 |
| LRRC37A3 | 4.05E-28 | -0.164933752 | 0.3 | 0.451 | 1.17E-23 |
| DTNB | 4.10E-28 | -0.222991666 | 0.4 | 0.617 | 1.18E-23 |
| TPD52 | 4.29E-28 | -0.362843509 | 0.3 | 0.417 | 1.24E-23 |
| POC1B | 4.56E-28 | -0.31699622 | 0.2 | 0.372 | 1.32E-23 |
| ELMO1-AS1 | 4.76E-28 | -0.325499986 | 0.1 | 0.203 | 1.38E-23 |
| C21orf91 | 4.81E-28 | -0.31074998 | 0.2 | 0.332 | 1.39E-23 |
| AL139294.1 | 5.80E-28 | -0.300978354 | 0.1 | 0.229 | 1.68E-23 |
| KIAA1958 | 6.01E-28 | -0.219201294 | 0.4 | 0.57 | 1.74E-23 |
| TJAP1 | 6.55E-28 | -0.258340266 | 0.3 | 0.433 | 1.89E-23 |
| SH3BP4 | 6.86E-28 | -0.316300384 | 0.2 | 0.375 | 1.98E-23 |
| MSRA | 6.97E-28 | 0.999905303 | 0.3 | 0.146 | 2.01E-23 |
| PPP2R5A | 7.52E-28 | -0.187683442 | 0.2 | 0.413 | 2.17E-23 |
| LRP6 | 7.55E-28 | -0.203167269 | 0.3 | 0.52 | 2.18E-23 |
| SS18 | 8.19E-28 | -0.243587718 | 0.2 | 0.369 | 2.37E-23 |
| CLASRP | 8.42E-28 | -0.250202967 | 0.2 | 0.383 | 2.43E-23 |
| CDK14 | 8.55E-28 | 0.770342008 | 0.2 | 0.124 | 2.47E-23 |
| TMEM178B | 8.75E-28 | 0.767582137 | 0.2 | 0.136 | 2.53E-23 |
| ILKAP | 9.80E-28 | -0.246371593 | 0.3 | 0.442 | 2.83E-23 |
| FADS1 | 1.00E-27 | -0.260768996 | 0.2 | 0.341 | 2.90E-23 |
| CRADD | 1.04E-27 | -0.364489846 | 0.4 | 0.562 | 2.99E-23 |
| EXOC4 | 1.04E-27 | -0.216409521 | 0.5 | 0.716 | 3.00E-23 |
| TTL | 1.06E-27 | -0.235745769 | 0.3 | 0.451 | 3.06E-23 |
| RORB | 1.26E-27 | 1.019527363 | 0.2 | 0.129 | 3.63E-23 |
| CD82 | 1.26E-27 | -0.288912945 | 0.1 | 0.293 | 3.64E-23 |
| PHTF2 | 1.31E-27 | -0.217172083 | 0.3 | 0.42 | 3.79E-23 |
| AL078590.3 | 1.41E-27 | -0.37603904 | 0.2 | 0.355 | 4.07E-23 |
| FIGN | 1.46E-27 | -0.343015776 | 0.4 | 0.563 | 4.22E-23 |
| NCS1 | 1.69E-27 | 0.584645975 | 0.1 | 0.038 | 4.89E-23 |
| SGCE | 1.77E-27 | -0.244967738 | 0.3 | 0.423 | 5.12E-23 |
| ALG13 | 1.86E-27 | -0.278381344 | 0.2 | 0.341 | 5.39E-23 |
| MBNL1 | 1.92E-27 | -0.233584932 | 0.5 | 0.744 | 5.54E-23 |
| AC090517.4 | 2.13E-27 | -0.323788165 | 0.2 | 0.414 | 6.14E-23 |
| THADA | 2.50E-27 | -0.240314259 | 0.3 | 0.444 | 7.23E-23 |
| AC106869.1 | 2.57E-27 | -0.359784791 | 0.2 | 0.401 | 7.41E-23 |
| UBL3 | 2.67E-27 | -0.163334026 | 0.4 | 0.544 | 7.70E-23 |
| CDKAL1 | 2.68E-27 | -0.201843331 | 0.4 | 0.602 | 7.75E-23 |

| | | | | | |
|---|---|---|---|---|---|
| ROBO1 | 3.06E-27 | -0.249959942 | 0.6 | 0.798 | 8.83E-23 |
| CPPED1 | 3.29E-27 | -0.318156543 | 0.2 | 0.341 | 9.51E-23 |
| ZBTB44 | 3.60E-27 | -0.206501918 | 0.3 | 0.499 | 1.04E-22 |
| IVNS1ABP | 3.74E-27 | -0.236660805 | 0.2 | 0.34 | 1.08E-22 |
| AC084032.1 | 3.90E-27 | -0.381478949 | 0.1 | 0.273 | 1.13E-22 |
| VSTM2B | 4.04E-27 | -0.232503009 | 0.2 | 0.357 | 1.17E-22 |
| PLA2G4C | 4.07E-27 | -0.335630943 | 0.2 | 0.413 | 1.18E-22 |
| SYT9 | 4.17E-27 | -0.253508435 | 0.3 | 0.479 | 1.21E-22 |
| KLK6 | 4.23E-27 | -0.294068227 | 0.2 | 0.314 | 1.22E-22 |
| TRAPPC9 | 4.41E-27 | -0.199986292 | 0.4 | 0.553 | 1.27E-22 |
| BAZ2B | 4.56E-27 | -0.145633718 | 0.6 | 0.867 | 1.32E-22 |
| ANKRD36 | 4.70E-27 | -0.142535004 | 0.4 | 0.637 | 1.36E-22 |
| GREB1L | 4.71E-27 | -0.363939652 | 0.3 | 0.423 | 1.36E-22 |
| UACA | 5.01E-27 | -0.396970623 | 0.2 | 0.348 | 1.45E-22 |
| KLF3 | 5.82E-27 | -0.272895433 | 0.2 | 0.316 | 1.68E-22 |
| PDK4 | 6.35E-27 | -0.359568623 | 0.2 | 0.348 | 1.84E-22 |
| EMC10 | 6.44E-27 | -0.214791919 | 0.2 | 0.343 | 1.86E-22 |
| TMC6 | 6.96E-27 | -0.237202726 | 0.2 | 0.361 | 2.01E-22 |
| RALY | 7.07E-27 | -0.267390024 | 0.2 | 0.344 | 2.04E-22 |
| SASH1 | 7.24E-27 | -0.167032391 | 0.5 | 0.719 | 2.09E-22 |
| FBXL5 | 8.48E-27 | -0.140936984 | 0.3 | 0.511 | 2.45E-22 |
| NCOR2 | 8.55E-27 | -0.177472897 | 0.3 | 0.486 | 2.47E-22 |
| PALLD | 1.01E-26 | 0.63710522 | 0.1 | 0.039 | 2.93E-22 |
| RNF213 | 1.02E-26 | -0.166325861 | 0.2 | 0.353 | 2.94E-22 |
| LSM14A | 1.04E-26 | -0.165716036 | 0.3 | 0.506 | 3.00E-22 |
| MT-CYB | 1.08E-26 | 0.685799714 | 0.8 | 0.789 | 3.13E-22 |
| ADAMTS14 | 1.17E-26 | -0.328870537 | 0.1 | 0.275 | 3.38E-22 |
| ENAH | 1.25E-26 | -0.18293277 | 0.4 | 0.552 | 3.61E-22 |
| HIBADH | 1.27E-26 | -0.344479647 | 0.2 | 0.366 | 3.66E-22 |
| ARID1B | 1.30E-26 | -0.181838664 | 0.6 | 0.837 | 3.75E-22 |
| LINC00877 | 1.30E-26 | -0.354779809 | 0.2 | 0.367 | 3.76E-22 |
| STAT3 | 1.34E-26 | -0.207918637 | 0.3 | 0.426 | 3.87E-22 |
| MBTD1 | 1.39E-26 | -0.189919026 | 0.2 | 0.415 | 4.00E-22 |
| SRP54 | 1.82E-26 | -0.230429824 | 0.2 | 0.333 | 5.27E-22 |
| SH3TC2-DT | 1.99E-26 | -0.42310771 | 0.2 | 0.312 | 5.75E-22 |
| RBL2 | 2.03E-26 | -0.191982307 | 0.3 | 0.421 | 5.86E-22 |
| KLHL3 | 2.13E-26 | -0.284447017 | 0.3 | 0.44 | 6.16E-22 |
| KNOP1 | 2.14E-26 | -0.281632654 | 0.3 | 0.449 | 6.19E-22 |
| NAALADL2 | 2.49E-26 | -0.328516748 | 0.5 | 0.641 | 7.21E-22 |
| GABPB1-AS1 | 2.50E-26 | -0.243490633 | 0.3 | 0.475 | 7.22E-22 |
| TIAM1 | 2.84E-26 | -0.258359796 | 0.4 | 0.562 | 8.22E-22 |
| WASHC4 | 2.98E-26 | -0.153974804 | 0.2 | 0.395 | 8.61E-22 |
| IPO13 | 3.16E-26 | -0.254543378 | 0.2 | 0.343 | 9.15E-22 |
| CLIP2 | 3.47E-26 | -0.159452194 | 0.4 | 0.616 | 1.00E-21 |
| CDKN1C | 3.67E-26 | -0.263574385 | 0.2 | 0.326 | 1.06E-21 |
| XRN2 | 3.70E-26 | -0.186648312 | 0.2 | 0.349 | 1.07E-21 |
| AC105383.1 | 3.71E-26 | -0.322393305 | 0.2 | 0.348 | 1.07E-21 |
| RC3H1 | 4.25E-26 | -0.231010178 | 0.3 | 0.442 | 1.23E-21 |
| USP31 | 4.44E-26 | -0.220761178 | 0.3 | 0.457 | 1.28E-21 |
| SMAD2 | 4.87E-26 | -0.220812097 | 0.3 | 0.412 | 1.41E-21 |
| NRF1 | 5.02E-26 | -0.163705454 | 0.3 | 0.465 | 1.45E-21 |
| YTHDC1 | 5.12E-26 | -0.166509704 | 0.3 | 0.472 | 1.48E-21 |
| BORCS5 | 5.19E-26 | -0.243209718 | 0.2 | 0.401 | 1.50E-21 |
| FAM172A | 5.42E-26 | -0.187654859 | 0.5 | 0.718 | 1.57E-21 |
| GCA | 5.44E-26 | -0.196645584 | 0.2 | 0.333 | 1.57E-21 |
| DAPK2 | 5.65E-26 | -0.235859186 | 0.2 | 0.41 | 1.63E-21 |
| NCDN | 5.72E-26 | 1.031065152 | 0.2 | 0.088 | 1.65E-21 |
| SOS2 | 5.73E-26 | -0.189355415 | 0.3 | 0.435 | 1.66E-21 |
| NR2C2 | 5.77E-26 | -0.251679631 | 0.2 | 0.409 | 1.67E-21 |
| STAG2 | 5.97E-26 | -0.186616166 | 0.3 | 0.506 | 1.73E-21 |
| PRKACB | 6.49E-26 | -0.186452221 | 0.6 | 0.782 | 1.88E-21 |
| NCBP3 | 6.62E-26 | -0.237691673 | 0.2 | 0.387 | 1.91E-21 |
| ADAM19 | 6.96E-26 | -0.385938983 | 0.1 | 0.272 | 2.01E-21 |
| MYO18A | 7.92E-26 | -0.225470334 | 0.2 | 0.404 | 2.29E-21 |
| MAP3K2 | 7.92E-26 | -0.167529867 | 0.3 | 0.515 | 2.29E-21 |
| LINC01184 | 8.25E-26 | -0.277357331 | 0.3 | 0.47 | 2.38E-21 |
| VPS54 | 8.60E-26 | -0.213185279 | 0.3 | 0.436 | 2.48E-21 |
| MLH3 | 9.07E-26 | -0.178748407 | 0.3 | 0.447 | 2.62E-21 |
| TAPT1 | 1.00E-25 | -0.178246417 | 0.2 | 0.372 | 2.90E-21 |
| CNKSR3 | 1.02E-25 | -0.360802697 | 0.2 | 0.404 | 2.94E-21 |
| UGDH-AS1 | 1.07E-25 | -0.260300028 | 0.2 | 0.316 | 3.11E-21 |
| MTREX | 1.09E-25 | -0.183788339 | 0.3 | 0.425 | 3.14E-21 |
| NOVA1 | 1.17E-25 | -0.160387627 | 0.5 | 0.713 | 3.38E-21 |
| TBC1D22A | 1.33E-25 | -0.151983079 | 0.3 | 0.513 | 3.85E-21 |
| LINC00844 | 1.37E-25 | -0.311831977 | 0.2 | 0.371 | 3.97E-21 |
| AMBRA1 | 1.55E-25 | -0.141472442 | 0.4 | 0.584 | 4.48E-21 |
| TBCA | 1.64E-25 | -0.206750977 | 0.3 | 0.455 | 4.73E-21 |
| PPA2 | 1.69E-25 | -0.160678072 | 0.3 | 0.525 | 4.89E-21 |
| CUX1 | 2.10E-25 | -0.233253782 | 0.4 | 0.555 | 6.07E-21 |
| ENTPD3-AS1 | 2.24E-25 | -0.270443154 | 0.2 | 0.318 | 6.47E-21 |
| CPD | 2.32E-25 | -0.19078005 | 0.2 | 0.384 | 6.72E-21 |
| ZBTB37 | 2.34E-25 | -0.243530802 | 0.2 | 0.315 | 6.76E-21 |
| WDSUB1 | 2.37E-25 | -0.200902606 | 0.1 | 0.259 | 6.84E-21 |
| TAOK3 | 2.38E-25 | -0.157811449 | 0.5 | 0.669 | 6.87E-21 |
| BOK | 2.39E-25 | -0.285640766 | 0.2 | 0.364 | 6.92E-21 |
| ELP2 | 2.65E-25 | -0.171646246 | 0.3 | 0.435 | 7.67E-21 |
| TMEM184B | 3.01E-25 | -0.21122868 | 0.3 | 0.466 | 8.71E-21 |
| THUMPD3-AS1 | 3.47E-25 | -0.200584253 | 0.2 | 0.36 | 1.00E-20 |
| ERMP1 | 3.88E-25 | -0.256380493 | 0.2 | 0.328 | 1.12E-20 |
| INSIG1 | 3.88E-25 | -0.257708037 | 0.2 | 0.307 | 1.12E-20 |
| ZZEF1 | 3.97E-25 | -0.195136275 | 0.2 | 0.396 | 1.15E-20 |
| EPHB1 | 4.04E-25 | 0.484060148 | 0.1 | 0.054 | 1.17E-20 |
| ARMC8 | 4.06E-25 | -0.150466714 | 0.3 | 0.467 | 1.17E-20 |
| FSCN1 | 4.08E-25 | -0.184942691 | 0.2 | 0.379 | 1.18E-20 |
| PTP4A2 | 4.34E-25 | -0.280266282 | 0.2 | 0.296 | 1.25E-20 |
| WWOX | 4.55E-25 | -0.315854851 | 0.7 | 0.86 | 1.31E-20 |
| SREBF1 | 4.61E-25 | -0.155269598 | 0.2 | 0.413 | 1.33E-20 |
| PAPOLA | 6.07E-25 | -0.144905481 | 0.3 | 0.483 | 1.75E-20 |
| EML2 | 6.22E-25 | -0.208942932 | 0.3 | 0.418 | 1.80E-20 |
| CTDSPL2 | 7.03E-25 | -0.169515384 | 0.2 | 0.391 | 2.03E-20 |
| LYPLAL1 | 7.12E-25 | -0.154736336 | 0.3 | 0.421 | 2.06E-20 |
| AGFG1 | 7.13E-25 | -0.140319812 | 0.3 | 0.43 | 2.06E-20 |
| SH3RF1 | 7.70E-25 | -0.288205323 | 0.3 | 0.484 | 2.23E-20 |
| PNPT1 | 7.78E-25 | -0.197514295 | 0.3 | 0.436 | 2.25E-20 |
| SNX6 | 7.92E-25 | -0.204542747 | 0.2 | 0.364 | 2.29E-20 |
| TOGARAM1 | 7.98E-25 | -0.181437818 | 0.4 | 0.54 | 2.31E-20 |
| RTTN | 8.13E-25 | -0.235932057 | 0.3 | 0.457 | 2.35E-20 |
| EIF4G3 | 8.41E-25 | -0.173087708 | 0.6 | 0.827 | 2.43E-20 |
| AMOTL2 | 8.43E-25 | -0.271593537 | 0.2 | 0.322 | 2.44E-20 |
| FAM49B | 8.45E-25 | -0.153904639 | 0.4 | 0.589 | 2.44E-20 |
| MIER1 | 8.54E-25 | -0.207029708 | 0.2 | 0.328 | 2.47E-20 |
| CEP95 | 8.73E-25 | -0.203003628 | 0.2 | 0.332 | 2.52E-20 |
| NUP214 | 9.69E-25 | -0.1920689 | 0.3 | 0.422 | 2.80E-20 |
| TAF1 | 1.01E-24 | -0.189867058 | 0.2 | 0.406 | 2.91E-20 |
| TPP2 | 1.12E-24 | -0.161057468 | 0.3 | 0.404 | 3.24E-20 |
| MEGF9 | 1.15E-24 | -0.174307714 | 0.4 | 0.6 | 3.32E-20 |
| CACNA1D | 1.18E-24 | 0.619006276 | 0.1 | 0.062 | 3.42E-20 |
| ANKRD13C | 1.19E-24 | -0.175983516 | 0.3 | 0.41 | 3.45E-20 |
| BRMS1L | 1.38E-24 | -0.279331076 | 0.2 | 0.375 | 4.00E-20 |
| AL354809.1 | 1.59E-24 | -0.474068601 | 0.2 | 0.397 | 4.59E-20 |
| CREBBP | 1.67E-24 | -0.158930987 | 0.3 | 0.528 | 4.82E-20 |
| PPFIBP2 | 1.67E-24 | -0.330496791 | 0.2 | 0.374 | 4.84E-20 |
| PDE5A | 1.78E-24 | -0.223623657 | 0.1 | 0.2 | 5.15E-20 |
| SRSF4 | 2.12E-24 | -0.215443066 | 0.2 | 0.351 | 6.11E-20 |
| LRP1B | 2.14E-24 | 0.687442548 | 0.8 | 0.773 | 6.18E-20 |
| FAM177A1 | 2.26E-24 | -0.182526335 | 0.2 | 0.387 | 6.54E-20 |
| CRYBG3 | 2.34E-24 | -0.293130346 | 0.2 | 0.333 | 6.76E-20 |
| PSRC1 | 2.39E-24 | -0.211337786 | 0.1 | 0.287 | 6.91E-20 |
| SHC4 | 2.50E-24 | -0.266169478 | 0.2 | 0.352 | 7.24E-20 |
| UTRN | 2.51E-24 | 0.340197701 | 0.1 | 0.05 | 7.27E-20 |
| LCORL | 2.52E-24 | -0.217560457 | 0.3 | 0.487 | 7.29E-20 |
| FRMD4A | 2.57E-24 | 1.235144313 | 0.5 | 0.416 | 7.43E-20 |
| CEP97 | 2.64E-24 | -0.20617247 | 0.3 | 0.435 | 7.63E-20 |
| DDX17 | 2.79E-24 | -0.174188907 | 0.7 | 0.922 | 8.08E-20 |
| ZCCHC7 | 3.17E-24 | -0.220953202 | 0.4 | 0.577 | 9.18E-20 |
| TCP11L2 | 3.78E-24 | -0.307674291 | 0.2 | 0.379 | 1.09E-19 |
| CYBRD1 | 3.94E-24 | -0.261943418 | 0.1 | 0.231 | 1.14E-19 |
| PRTFDC1 | 4.09E-24 | -0.250265578 | 0.3 | 0.431 | 1.18E-19 |

| | | | | | |
|---|---|---|---|---|---|
| NBEAL1 | 4.13E-24 | -0.180980783 | 0.3 | 0.424 | 1.19E-19 |
| ZNF462 | 4.20E-24 | -0.15559045 | 0.3 | 0.436 | 1.21E-19 |
| RRBP1 | 4.60E-24 | -0.192011809 | 0.2 | 0.336 | 1.33E-19 |
| DNAH17 | 4.90E-24 | -0.317933331 | 0.1 | 0.224 | 1.42E-19 |
| DPF3 | 4.92E-24 | -0.154656151 | 0.3 | 0.429 | 1.42E-19 |
| TPTEP2-CSNK1E | 5.08E-24 | -0.209293278 | 0.2 | 0.383 | 1.47E-19 |
| TXNIP | 5.69E-24 | -0.31116294 | 0.3 | 0.47 | 1.65E-19 |
| DHX15 | 5.80E-24 | -0.144172916 | 0.2 | 0.39 | 1.68E-19 |
| DSEL | 5.83E-24 | -0.280089397 | 0.2 | 0.317 | 1.69E-19 |
| USP14 | 6.14E-24 | -0.139163488 | 0.2 | 0.386 | 1.77E-19 |
| DIRAS2 | 6.64E-24 | 0.76821934 | 0.1 | 0.041 | 1.92E-19 |
| ARHGAP24 | 6.68E-24 | 0.687468029 | 0.2 | 0.121 | 1.93E-19 |
| TBL1XR1 | 6.72E-24 | -0.142296283 | 0.5 | 0.674 | 1.94E-19 |
| ARHGAP17 | 6.80E-24 | -0.260692698 | 0.3 | 0.405 | 1.97E-19 |
| CAMK2G | 7.39E-24 | 0.839165663 | 0.2 | 0.123 | 2.14E-19 |
| AC078881.1 | 8.67E-24 | -0.410897902 | 0.2 | 0.309 | 2.51E-19 |
| LIPE | 9.43E-24 | -0.268530583 | 0.2 | 0.307 | 2.72E-19 |
| SH3PXD2B | 9.59E-24 | -0.168601056 | 0.3 | 0.431 | 2.77E-19 |
| KIAA1328 | 1.07E-23 | -0.211234033 | 0.3 | 0.529 | 3.08E-19 |
| INVS | 1.09E-23 | -0.178709314 | 0.4 | 0.557 | 3.15E-19 |
| PPP4R1 | 1.11E-23 | -0.181791196 | 0.2 | 0.331 | 3.22E-19 |
| PTPDC1 | 1.13E-23 | -0.235242415 | 0.2 | 0.398 | 3.27E-19 |
| DPYSL5 | 1.37E-23 | -0.380816014 | 0.3 | 0.443 | 3.95E-19 |
| C1orf198 | 1.40E-23 | -0.181441743 | 0.2 | 0.403 | 4.04E-19 |
| RNH1 | 1.46E-23 | -0.146090602 | 0.2 | 0.333 | 4.22E-19 |
| TMEM59L | 1.49E-23 | 1.083638081 | 0.2 | 0.101 | 4.30E-19 |
| SFT2D1 | 1.54E-23 | -0.24139536 | 0.1 | 0.274 | 4.46E-19 |
| PLPP2 | 1.55E-23 | -0.209159403 | 0.2 | 0.307 | 4.49E-19 |
| C3orf58 | 1.75E-23 | -0.202475262 | 0.3 | 0.41 | 5.05E-19 |
| TALDO1 | 1.94E-23 | -0.172073237 | 0.2 | 0.304 | 5.61E-19 |
| TRPM6 | 2.12E-23 | -0.326556251 | 0.1 | 0.269 | 6.12E-19 |
| SLC6A6 | 2.20E-23 | -0.285239217 | 0.1 | 0.204 | 6.34E-19 |
| MTRR | 2.21E-23 | -0.283991392 | 0.1 | 0.233 | 6.38E-19 |
| KIAA1147 | 2.46E-23 | -0.227487483 | 0.2 | 0.383 | 7.10E-19 |
| ARHGEF37 | 2.55E-23 | -0.216141882 | 0.2 | 0.341 | 7.37E-19 |
| GALNT15 | 2.67E-23 | -0.305166206 | 0.1 | 0.267 | 7.73E-19 |
| SPSB1 | 2.69E-23 | -0.283158417 | 0.1 | 0.255 | 7.77E-19 |
| NENF | 3.30E-23 | -0.203735195 | 0.1 | 0.268 | 9.53E-19 |
| HCN2 | 3.36E-23 | -0.241486677 | 0.1 | 0.276 | 9.71E-19 |
| THY1 | 3.36E-23 | 1.189054447 | 0.1 | 0.071 | 9.72E-19 |
| ADD1 | 3.99E-23 | -0.170479375 | 0.5 | 0.741 | 1.15E-18 |
| TECR | 4.05E-23 | -0.202604037 | 0.2 | 0.352 | 1.17E-18 |
| PDE1A | 4.34E-23 | -0.307610029 | 0.4 | 0.629 | 1.25E-18 |
| CFL2 | 4.37E-23 | -0.21196168 | 0.2 | 0.328 | 1.26E-18 |
| PLPP1 | 4.87E-23 | -0.180211434 | 0.3 | 0.496 | 1.41E-18 |
| LINC02241 | 4.92E-23 | -0.172706655 | 0.3 | 0.413 | 1.42E-18 |
| ITPR2 | 5.32E-23 | 1.063644187 | 0.4 | 0.253 | 1.54E-18 |
| MFHAS1 | 5.41E-23 | 0.640358094 | 0.1 | 0.05 | 1.56E-18 |
| STXBP6 | 5.77E-23 | -0.203746286 | 0.4 | 0.545 | 1.67E-18 |
| SLC39A11 | 5.85E-23 | -0.174710197 | 0.5 | 0.738 | 1.69E-18 |
| ATAD2B | 5.93E-23 | -0.209107084 | 0.3 | 0.438 | 1.71E-18 |
| LINC01135 | 6.29E-23 | -0.337501298 | 0.2 | 0.356 | 1.82E-18 |
| TRPC1 | 6.50E-23 | -0.137698672 | 0.3 | 0.512 | 1.88E-18 |
| PCBP4 | 6.85E-23 | -0.149173287 | 0.2 | 0.345 | 1.98E-18 |
| CST3 | 6.91E-23 | 0.864672922 | 0.2 | 0.094 | 2.00E-18 |
| CADM4 | 7.24E-23 | -0.173896907 | 0.2 | 0.336 | 2.09E-18 |
| ANKUB1 | 8.13E-23 | -0.207102451 | 0.1 | 0.239 | 2.35E-18 |
| ELOVL1 | 8.58E-23 | -0.231403234 | 0.2 | 0.291 | 2.48E-18 |
| TESK2 | 9.28E-23 | -0.258063186 | 0.3 | 0.469 | 2.68E-18 |
| DNHD1 | 9.29E-23 | -0.187796925 | 0.2 | 0.33 | 2.69E-18 |
| PRKCQ | 1.02E-22 | -0.280578378 | 0.2 | 0.342 | 2.94E-18 |
| VTI1A | 1.05E-22 | -0.17442677 | 0.4 | 0.562 | 3.05E-18 |
| AC027644.4 | 1.14E-22 | -0.206084666 | 0.3 | 0.425 | 3.30E-18 |
| WDR44 | 1.21E-22 | -0.203154074 | 0.2 | 0.381 | 3.49E-18 |
| ANXA4 | 1.36E-22 | -0.283207576 | 0.3 | 0.419 | 3.94E-18 |
| TMEM135 | 1.42E-22 | -0.164734367 | 0.3 | 0.469 | 4.09E-18 |
| ZNF189 | 1.79E-22 | -0.177951342 | 0.2 | 0.325 | 5.17E-18 |
| CES4A | 1.85E-22 | -0.244345709 | 0.2 | 0.315 | 5.36E-18 |
| STOX2 | 1.86E-22 | -0.179332326 | 0.4 | 0.53 | 5.38E-18 |
| SPINT2 | 2.01E-22 | -0.22361644 | 0.1 | 0.166 | 5.81E-18 |
| 8-Mar | 2.17E-22 | -0.249641244 | 0.2 | 0.312 | 6.29E-18 |
| EYA3 | 2.41E-22 | -0.205428066 | 0.2 | 0.332 | 6.96E-18 |
| MID1IP1 | 3.27E-22 | -0.261496293 | 0.2 | 0.302 | 9.46E-18 |
| SLC6A8 | 3.53E-22 | -0.16788613 | 0.2 | 0.365 | 1.02E-17 |
| CYLD | 3.81E-22 | -0.148055988 | 0.3 | 0.443 | 1.10E-17 |
| PDE6B | 3.84E-22 | -0.336155688 | 0.1 | 0.233 | 1.11E-17 |
| EML6 | 3.90E-22 | 0.61526817 | 0.1 | 0.069 | 1.13E-17 |
| WWP2 | 4.12E-22 | -0.21635469 | 0.3 | 0.436 | 1.19E-17 |
| B2M | 4.89E-22 | -0.202495594 | 0.2 | 0.372 | 1.41E-17 |
| HDAC11 | 5.17E-22 | -0.194444069 | 0.2 | 0.305 | 1.49E-17 |
| TNS1 | 5.28E-22 | -0.265556313 | 0.1 | 0.245 | 1.53E-17 |
| COMMD1 | 5.45E-22 | -0.267907071 | 0.3 | 0.408 | 1.57E-17 |
| AP1S2 | 5.73E-22 | -0.248023689 | 0.2 | 0.288 | 1.66E-17 |
| CYP2J2 | 5.97E-22 | -0.241838119 | 0.2 | 0.302 | 1.72E-17 |
| AL139383.1 | 5.97E-22 | -0.318675054 | 0.2 | 0.347 | 1.73E-17 |
| ARHGEF4 | 6.07E-22 | 0.898872826 | 0.2 | 0.101 | 1.75E-17 |
| TMEM151A | 6.12E-22 | -0.150800627 | 0.2 | 0.295 | 1.77E-17 |
| SEC31A | 6.57E-22 | -0.167704079 | 0.2 | 0.379 | 1.90E-17 |
| TFCP2 | 6.66E-22 | -0.182923681 | 0.3 | 0.447 | 1.92E-17 |
| SETBP1 | 7.00E-22 | 0.593317004 | 0.2 | 0.082 | 2.02E-17 |
| SLC13A3 | 7.30E-22 | -0.218146812 | 0.3 | 0.43 | 2.11E-17 |
| PHLDA3 | 7.36E-22 | -0.190669626 | 0.1 | 0.269 | 2.13E-17 |
| PPFIBP1 | 8.35E-22 | -0.199455729 | 0.3 | 0.485 | 2.41E-17 |
| RHBDL2 | 8.49E-22 | -0.20493024 | 0.1 | 0.262 | 2.45E-17 |
| SLC25A27 | 9.19E-22 | -0.161056973 | 0.3 | 0.418 | 2.66E-17 |
| MAGI3 | 9.56E-22 | 0.769082738 | 0.2 | 0.092 | 2.76E-17 |
| LINC01505 | 1.00E-21 | -0.559093746 | 0.2 | 0.295 | 2.90E-17 |
| FAM214A | 1.02E-21 | -0.140933904 | 0.3 | 0.475 | 2.96E-17 |
| CYP7B1 | 1.08E-21 | -0.32917256 | 0.2 | 0.372 | 3.13E-17 |
| S100PBP | 1.13E-21 | -0.15919943 | 0.2 | 0.362 | 3.26E-17 |
| FOXO1 | 1.15E-21 | -0.232181271 | 0.3 | 0.475 | 3.33E-17 |
| MED27 | 1.21E-21 | -0.144890394 | 0.3 | 0.462 | 3.49E-17 |
| SLC4A8 | 1.30E-21 | -0.195842064 | 0.3 | 0.436 | 3.77E-17 |
| ICK | 1.65E-21 | -0.242427793 | 0.2 | 0.298 | 4.78E-17 |
| LRRC8C-DT | 1.67E-21 | -0.278617218 | 0.2 | 0.326 | 4.84E-17 |
| NSRP1 | 1.74E-21 | -0.143255397 | 0.2 | 0.382 | 5.02E-17 |
| FBXL4 | 1.83E-21 | -0.156753144 | 0.2 | 0.377 | 5.29E-17 |
| NUBPL | 1.97E-21 | -0.184545198 | 0.3 | 0.433 | 5.70E-17 |
| RAP1B | 2.10E-21 | -0.157115521 | 0.2 | 0.33 | 6.06E-17 |
| TSEN15 | 2.13E-21 | -0.19761847 | 0.2 | 0.361 | 6.16E-17 |
| SNAP91 | 2.19E-21 | 0.90799198 | 0.2 | 0.141 | 6.32E-17 |
| CALM3 | 2.20E-21 | 1.328633675 | 0.2 | 0.118 | 6.37E-17 |
| LIFR | 2.24E-21 | -0.143935691 | 0.5 | 0.682 | 6.48E-17 |
| ZDHHC14 | 2.36E-21 | -0.17604298 | 0.3 | 0.467 | 6.81E-17 |
| PIGC | 2.47E-21 | -0.193048642 | 0.1 | 0.25 | 7.14E-17 |
| DHX29 | 2.53E-21 | -0.146173537 | 0.2 | 0.308 | 7.32E-17 |
| AC063944.1 | 2.57E-21 | -0.303776151 | 0.2 | 0.316 | 7.43E-17 |
| RSRC1 | 2.66E-21 | -0.139064236 | 0.3 | 0.456 | 7.70E-17 |
| GOLPH3 | 2.98E-21 | -0.139913985 | 0.2 | 0.339 | 8.61E-17 |
| SH3BP5 | 3.17E-21 | -0.141749482 | 0.1 | 0.27 | 9.15E-17 |
| ICE2 | 3.34E-21 | -0.176326261 | 0.2 | 0.337 | 9.66E-17 |
| PLD5 | 3.37E-21 | -0.334211276 | 0.3 | 0.483 | 9.74E-17 |
| ASTN1 | 4.00E-21 | 0.755015618 | 0.2 | 0.11 | 1.16E-16 |
| EFS | 4.00E-21 | -0.177826185 | 0.1 | 0.245 | 1.16E-16 |
| ZNF718 | 4.27E-21 | -0.156861029 | 0.2 | 0.332 | 1.23E-16 |
| TTC7A | 4.28E-21 | -0.148526686 | 0.3 | 0.429 | 1.24E-16 |
| PCSK7 | 4.28E-21 | -0.189096255 | 0.2 | 0.351 | 1.24E-16 |
| FAM114A1 | 4.32E-21 | -0.211158693 | 0.1 | 0.277 | 1.25E-16 |
| TMEM64 | 4.71E-21 | -0.28983064 | 0.2 | 0.347 | 1.36E-16 |
| ADAMTS4 | 4.72E-21 | -0.293297412 | 0.1 | 0.241 | 1.37E-16 |
| BCAR1 | 5.02E-21 | -0.162943287 | 0.1 | 0.276 | 1.45E-16 |
| FBXO28 | 5.86E-21 | -0.279766073 | 0.2 | 0.276 | 1.69E-16 |
| PLS3 | 6.53E-21 | -0.245918786 | 0.2 | 0.338 | 1.89E-16 |
| REPS2 | 6.71E-21 | -0.169378924 | 0.3 | 0.478 | 1.94E-16 |
| NEK9 | 7.41E-21 | -0.151113288 | 0.2 | 0.333 | 2.14E-16 |
| FAM133B | 8.28E-21 | -0.202642807 | 0.2 | 0.328 | 2.39E-16 |
| SERPINB6 | 8.42E-21 | -0.174414965 | 0.2 | 0.315 | 2.43E-16 |
| GALNT6 | 8.46E-21 | -0.253015141 | 0.1 | 0.272 | 2.45E-16 |

| | | | | | |
|---|---|---|---|---|---|
| MYO15B | 8.48E-21 | -0.221417786 | 0.1 | 0.274 | 2.45E-16 |
| TTLL5 | 8.74E-21 | -0.161211778 | 0.2 | 0.391 | 2.52E-16 |
| CMC2 | 9.61E-21 | -0.178053231 | 0.2 | 0.289 | 2.78E-16 |
| AF165147.1 | 1.08E-20 | -0.308249632 | 0.2 | 0.347 | 3.12E-16 |
| TAPT1-AS1 | 1.08E-20 | -0.141220855 | 0.4 | 0.534 | 3.13E-16 |
| XPO4 | 1.13E-20 | -0.144022608 | 0.2 | 0.354 | 3.26E-16 |
| CMTM5 | 1.17E-20 | -0.244631411 | 0.1 | 0.252 | 3.38E-16 |
| LRRFIP1 | 1.26E-20 | 0.489992875 | 0.1 | 0.049 | 3.65E-16 |
| KCNAB1 | 1.31E-20 | -0.461973609 | 0.3 | 0.398 | 3.78E-16 |
| ARPC5 | 1.31E-20 | -0.205572264 | 0.1 | 0.253 | 3.79E-16 |
| CFLAR | 1.38E-20 | -0.17365296 | 0.2 | 0.36 | 3.98E-16 |
| NHLRC3 | 1.40E-20 | -0.138436376 | 0.2 | 0.355 | 4.06E-16 |
| ZNF365 | 1.43E-20 | -0.24577015 | 0.3 | 0.515 | 4.13E-16 |
| ARHGAP44 | 1.51E-20 | 0.557427123 | 0.1 | 0.056 | 4.37E-16 |
| RILPL1 | 1.53E-20 | -0.138675475 | 0.2 | 0.384 | 4.42E-16 |
| TRMT13 | 1.56E-20 | -0.171261557 | 0.2 | 0.346 | 4.52E-16 |
| SMARCC1 | 1.62E-20 | -0.166349998 | 0.3 | 0.475 | 4.68E-16 |
| CEP112 | 1.63E-20 | 0.419633998 | 0.1 | 0.049 | 4.70E-16 |
| TMCC1 | 1.63E-20 | -0.168970795 | 0.4 | 0.569 | 4.72E-16 |
| SOGA1 | 1.69E-20 | -0.145820519 | 0.3 | 0.411 | 4.89E-16 |
| CDH11 | 1.73E-20 | -0.247654413 | 0.2 | 0.373 | 5.01E-16 |
| CERS4 | 1.80E-20 | -0.147645449 | 0.3 | 0.43 | 5.21E-16 |
| TATDN3 | 1.83E-20 | -0.20699971 | 0.2 | 0.301 | 5.28E-16 |
| ENO4 | 1.87E-20 | -0.208286904 | 0.1 | 0.226 | 5.41E-16 |
| STEAP1B | 2.19E-20 | -0.28488397 | 0.1 | 0.257 | 6.33E-16 |
| YWHAH | 2.21E-20 | 1.1629214 | 0.1 | 0.078 | 6.37E-16 |
| TMEM206 | 2.36E-20 | -0.235868256 | 0.1 | 0.255 | 6.83E-16 |
| UFL1 | 2.40E-20 | -0.181632253 | 0.2 | 0.363 | 6.92E-16 |
| AL138881.1 | 2.47E-20 | -0.301379261 | 0.1 | 0.244 | 7.14E-16 |
| PDGFA | 2.55E-20 | -0.178934829 | 0.1 | 0.224 | 7.38E-16 |
| FAM69A | 2.57E-20 | -0.145124634 | 0.4 | 0.567 | 7.43E-16 |
| CCDC122 | 2.58E-20 | -0.238839128 | 0.2 | 0.395 | 7.46E-16 |
| SEMA6A-AS1 | 2.64E-20 | -0.262994676 | 0.1 | 0.165 | 7.64E-16 |
| AC139887.2 | 2.89E-20 | -0.169724071 | 0.3 | 0.419 | 8.34E-16 |
| DAB1 | 2.89E-20 | 0.858544139 | 0.3 | 0.237 | 8.37E-16 |
| CSGALNACT2 | 3.01E-20 | -0.154387236 | 0.1 | 0.262 | 8.71E-16 |
| PADI2 | 3.02E-20 | -0.15756556 | 0.3 | 0.428 | 8.73E-16 |
| PPM1E | 3.06E-20 | 0.598979509 | 0.1 | 0.055 | 8.84E-16 |
| EMSY | 3.28E-20 | -0.216158601 | 0.3 | 0.392 | 9.49E-16 |
| AFMID | 3.32E-20 | -0.195593601 | 0.2 | 0.39 | 9.60E-16 |
| TTC27 | 3.41E-20 | -0.217616486 | 0.2 | 0.279 | 9.85E-16 |
| FBXO32 | 4.14E-20 | -0.237106147 | 0.2 | 0.311 | 1.20E-15 |
| CDC42EP1 | 4.15E-20 | -0.238423161 | 0.2 | 0.289 | 1.20E-15 |
| XIST | 4.64E-20 | -0.351844381 | 0.3 | 0.416 | 1.34E-15 |
| RFFL | 4.79E-20 | -0.225323484 | 0.1 | 0.249 | 1.39E-15 |
| GIPC1 | 4.90E-20 | -0.176075144 | 0.2 | 0.316 | 1.42E-15 |
| AC004690.2 | 5.06E-20 | -0.273029263 | 0.1 | 0.269 | 1.46E-15 |
| TRHDE | 5.22E-20 | 0.676557278 | 0.2 | 0.095 | 1.51E-15 |
| TPRN | 5.23E-20 | -0.180705741 | 0.2 | 0.297 | 1.51E-15 |
| LINC00862 | 5.36E-20 | -0.309076746 | 0.2 | 0.373 | 1.55E-15 |
| SLC35D2 | 5.85E-20 | -0.284991695 | 0.1 | 0.25 | 1.69E-15 |
| PDCD4 | 7.57E-20 | -0.183106069 | 0.2 | 0.278 | 2.19E-15 |
| GAL3ST1 | 7.59E-20 | -0.26671396 | 0.1 | 0.232 | 2.19E-15 |
| ZNF83 | 8.71E-20 | -0.175055714 | 0.2 | 0.377 | 2.52E-15 |
| NTRK2 | 8.75E-20 | 0.886643921 | 0.5 | 0.48 | 2.53E-15 |
| CXXC5 | 9.19E-20 | -0.147128394 | 0.2 | 0.342 | 2.66E-15 |
| G3BP1 | 9.27E-20 | -0.165044136 | 0.2 | 0.327 | 2.68E-15 |
| ALDH3A2 | 9.85E-20 | -0.160407024 | 0.2 | 0.306 | 2.85E-15 |
| PCCB | 1.01E-19 | -0.169453791 | 0.2 | 0.363 | 2.91E-15 |
| APPL2 | 1.02E-19 | -0.148432669 | 0.3 | 0.394 | 2.95E-15 |
| LINC00854 | 1.11E-19 | -0.295795774 | 0.2 | 0.333 | 3.20E-15 |
| MTERF1 | 1.12E-19 | -0.205313681 | 0.2 | 0.367 | 3.22E-15 |
| BAIAP2L2 | 1.23E-19 | -0.164697805 | 0.1 | 0.236 | 3.55E-15 |
| AC245297.3 | 1.27E-19 | -0.147219495 | 0.5 | 0.653 | 3.68E-15 |
| DMD | 1.32E-19 | -0.217528601 | 0.6 | 0.756 | 3.82E-15 |
| PBX3 | 1.46E-19 | -0.298022869 | 0.2 | 0.366 | 4.23E-15 |
| PTPN12 | 1.47E-19 | -0.154483331 | 0.2 | 0.355 | 4.24E-15 |
| RFX3-AS1 | 1.62E-19 | -0.16674512 | 0.3 | 0.43 | 4.68E-15 |
| CFTR | 1.86E-19 | -0.227206829 | 0.1 | 0.21 | 5.39E-15 |
| IFIT3 | 1.89E-19 | -0.162910058 | 0.1 | 0.2 | 5.45E-15 |
| UBA3 | 1.99E-19 | -0.142931968 | 0.2 | 0.334 | 5.75E-15 |
| BDNF-AS | 2.00E-19 | -0.215795893 | 0.2 | 0.321 | 5.77E-15 |
| LRMDA | 2.01E-19 | 0.710270642 | 0.2 | 0.099 | 5.80E-15 |
| TOB2 | 2.11E-19 | -0.139933019 | 0.3 | 0.41 | 6.10E-15 |
| IFT88 | 2.15E-19 | -0.210810923 | 0.3 | 0.397 | 6.23E-15 |
| FAM69C | 2.17E-19 | -0.255053446 | 0.1 | 0.261 | 6.27E-15 |
| CEND1 | 2.42E-19 | 0.796412423 | 0.1 | 0.048 | 7.00E-15 |
| TPM4 | 2.81E-19 | -0.189496164 | 0.2 | 0.296 | 8.13E-15 |
| RAB21 | 2.99E-19 | -0.185732458 | 0.2 | 0.333 | 8.64E-15 |
| SMOC1 | 3.03E-19 | -0.338557326 | 0.1 | 0.231 | 8.77E-15 |
| LRRC37A2 | 3.12E-19 | -0.144955221 | 0.1 | 0.242 | 9.02E-15 |
| GDI2 | 3.17E-19 | -0.176184584 | 0.2 | 0.285 | 9.15E-15 |
| LINC00882 | 3.38E-19 | -0.217079759 | 0.4 | 0.554 | 9.77E-15 |
| DENND2A | 3.50E-19 | -0.23071413 | 0.3 | 0.406 | 1.01E-14 |
| ZIC1 | 4.32E-19 | -0.199419147 | 0.3 | 0.39 | 1.25E-14 |
| DAAM1 | 4.58E-19 | 0.809725506 | 0.2 | 0.145 | 1.32E-14 |
| RASGRF2 | 4.72E-19 | -0.243293408 | 0.3 | 0.408 | 1.36E-14 |
| MDGA2 | 4.74E-19 | 0.632777259 | 0.3 | 0.233 | 1.37E-14 |
| NFKB1 | 5.26E-19 | -0.174144952 | 0.3 | 0.416 | 1.52E-14 |
| PRKAR2A | 5.37E-19 | -0.149663578 | 0.3 | 0.437 | 1.55E-14 |
| LINC01411 | 5.76E-19 | -0.217666696 | 0.2 | 0.331 | 1.67E-14 |
| RAB8B | 5.77E-19 | -0.199161097 | 0.2 | 0.268 | 1.67E-14 |
| KPNA3 | 5.89E-19 | -0.144249399 | 0.2 | 0.336 | 1.70E-14 |
| SPNS2 | 6.09E-19 | -0.172814366 | 0.2 | 0.377 | 1.76E-14 |
| WDPCP | 6.24E-19 | -0.183319849 | 0.5 | 0.668 | 1.80E-14 |
| CHADL | 6.25E-19 | -0.242481374 | 0.2 | 0.292 | 1.81E-14 |
| SYNGR2 | 6.54E-19 | -0.218734382 | 0.1 | 0.186 | 1.89E-14 |
| ADGRB3 | 6.90E-19 | 0.499064321 | 0.8 | 0.848 | 1.99E-14 |
| CCDC66 | 7.12E-19 | -0.173381285 | 0.2 | 0.384 | 2.06E-14 |
| PARN | 7.26E-19 | -0.161889819 | 0.2 | 0.371 | 2.10E-14 |
| CBLB | 7.55E-19 | -0.246817251 | 0.3 | 0.384 | 2.18E-14 |
| DCP1A | 7.64E-19 | -0.160815609 | 0.2 | 0.268 | 2.21E-14 |
| SOX2 | 7.92E-19 | -0.142317206 | 0.2 | 0.376 | 2.29E-14 |
| FN3K | 8.26E-19 | -0.186156794 | 0.1 | 0.218 | 2.39E-14 |
| KRCC1 | 8.47E-19 | -0.146321645 | 0.2 | 0.287 | 2.45E-14 |
| MED14 | 8.60E-19 | -0.1989795 | 0.2 | 0.33 | 2.49E-14 |
| SF3B2 | 8.83E-19 | -0.146725709 | 0.2 | 0.29 | 2.55E-14 |
| ODR4 | 8.85E-19 | -0.168515367 | 0.2 | 0.305 | 2.56E-14 |
| NADSYN1 | 8.91E-19 | -0.166259937 | 0.1 | 0.27 | 2.58E-14 |
| AP003086.2 | 8.98E-19 | -0.171695175 | 0.1 | 0.261 | 2.59E-14 |
| NFKBIZ | 9.89E-19 | -0.195638203 | 0.2 | 0.282 | 2.86E-14 |
| C1orf122 | 1.08E-18 | -0.175907795 | 0.1 | 0.263 | 3.11E-14 |
| EFCAB14 | 1.16E-18 | -0.179784795 | 0.1 | 0.253 | 3.36E-14 |
| SNX2 | 1.42E-18 | -0.146232028 | 0.1 | 0.236 | 4.09E-14 |
| SAR1B | 1.63E-18 | -0.178573668 | 0.2 | 0.286 | 4.72E-14 |
| EGFR | 1.69E-18 | 0.336520636 | 0.1 | 0.055 | 4.89E-14 |
| CPEB1 | 1.87E-18 | -0.238796111 | 0.2 | 0.321 | 5.41E-14 |
| ST3GAL4 | 1.90E-18 | -0.222874641 | 0.2 | 0.294 | 5.50E-14 |
| ABTB2 | 1.91E-18 | -0.344721495 | 0.2 | 0.285 | 5.51E-14 |
| LRP4-AS1 | 1.93E-18 | -0.210214454 | 0 | 0.134 | 5.57E-14 |
| LDLRAD3 | 1.93E-18 | -0.335815051 | 0.3 | 0.409 | 5.59E-14 |
| LURAP1L-AS1 | 2.12E-18 | -0.362860359 | 0.3 | 0.408 | 6.13E-14 |
| GJB1 | 2.28E-18 | -0.212657918 | 0.1 | 0.189 | 6.60E-14 |
| ANKRD23 | 2.59E-18 | -0.222433501 | 0.2 | 0.281 | 7.49E-14 |
| IRF2 | 2.95E-18 | -0.17635745 | 0.3 | 0.384 | 8.53E-14 |
| FAM221A | 2.96E-18 | -0.158650455 | 0.1 | 0.224 | 8.55E-14 |
| CSMD2 | 2.96E-18 | -0.19815483 | 0.3 | 0.418 | 8.56E-14 |
| CYB5R4 | 3.21E-18 | -0.219574598 | 0.2 | 0.353 | 9.28E-14 |
| FAM184A | 3.32E-18 | 0.697870622 | 0.1 | 0.08 | 9.58E-14 |
| AL162511.1 | 4.04E-18 | -0.214488839 | 0.1 | 0.193 | 1.17E-13 |
| ASCC1 | 4.14E-18 | -0.170666972 | 0.2 | 0.303 | 1.20E-13 |
| NEGR1 | 4.25E-18 | 0.757316025 | 0.5 | 0.462 | 1.23E-13 |
| POLA1 | 4.41E-18 | -0.154489838 | 0.2 | 0.374 | 1.27E-13 |
| R3HCC1L | 4.64E-18 | -0.199496318 | 0.1 | 0.238 | 1.34E-13 |
| HID1 | 5.03E-18 | -0.138229976 | 0.2 | 0.366 | 1.45E-13 |
| PLOD3 | 5.72E-18 | -0.193262494 | 0.1 | 0.218 | 1.65E-13 |
| LEMD3 | 5.74E-18 | -0.169083115 | 0.1 | 0.231 | 1.66E-13 |
| OLIG2 | 5.97E-18 | -0.160302989 | 0.2 | 0.269 | 1.73E-13 |

| | | | | | |
|---|---|---|---|---|---|
| AC096677.1 | 7.11E-18 | -0.234970109 | 0.1 | 0.246 | 2.05E-13 |
| PPP2R3C | 7.47E-18 | -0.14563741 | 0.2 | 0.315 | 2.16E-13 |
| SP4 | 7.84E-18 | -0.194144105 | 0.2 | 0.284 | 2.27E-13 |
| EFNA1 | 7.90E-18 | -0.188537054 | 0.1 | 0.21 | 2.28E-13 |
| CAPS2 | 8.04E-18 | -0.194125562 | 0.2 | 0.363 | 2.32E-13 |
| AL392023.2 | 8.50E-18 | -0.329134817 | 0.1 | 0.256 | 2.46E-13 |
| SCFD2 | 8.56E-18 | -0.144212817 | 0.3 | 0.426 | 2.47E-13 |
| TGFA | 9.12E-18 | -0.237693507 | 0.1 | 0.206 | 2.64E-13 |
| BBIP1 | 9.22E-18 | -0.163339352 | 0.2 | 0.293 | 2.67E-13 |
| ECHDC1 | 1.03E-17 | -0.150501798 | 0.1 | 0.244 | 2.98E-13 |
| C21orf62-AS1 | 1.04E-17 | -0.199033869 | 0.2 | 0.338 | 3.01E-13 |
| HEG1 | 1.08E-17 | -0.227812695 | 0.1 | 0.172 | 3.13E-13 |
| PIGN | 1.09E-17 | -0.148839883 | 0.2 | 0.375 | 3.14E-13 |
| ENTPD1-AS1 | 1.14E-17 | -0.183075296 | 0.3 | 0.453 | 3.30E-13 |
| SPTBN4 | 1.17E-17 | 0.711095923 | 0.2 | 0.1 | 3.39E-13 |
| SPATA22 | 1.41E-17 | -0.233558709 | 0.1 | 0.197 | 4.08E-13 |
| SLC6A1-AS1 | 1.42E-17 | -0.164990418 | 0.1 | 0.172 | 4.12E-13 |
| PIGF | 1.46E-17 | -0.18913473 | 0.1 | 0.251 | 4.23E-13 |
| LANCL1-AS1 | 1.47E-17 | -0.168521539 | 0.1 | 0.142 | 4.25E-13 |
| 10-Sep | 1.49E-17 | -0.201766522 | 0.1 | 0.252 | 4.32E-13 |
| SGIP1 | 1.53E-17 | -0.142272285 | 0.5 | 0.653 | 4.42E-13 |
| MKL2 | 1.62E-17 | 0.781277921 | 0.2 | 0.117 | 4.69E-13 |
| AL358216.1 | 1.67E-17 | -0.227790878 | 0.1 | 0.212 | 4.84E-13 |
| KCTD8 | 1.81E-17 | -0.288261946 | 0.3 | 0.448 | 5.23E-13 |
| CAMKMT | 1.89E-17 | -0.281556068 | 0.3 | 0.47 | 5.46E-13 |
| NEAT1 | 1.98E-17 | -0.193069121 | 0.7 | 0.925 | 5.72E-13 |
| VLDLR-AS1 | 2.03E-17 | -0.318863463 | 0.2 | 0.265 | 5.85E-13 |
| AP000688.2 | 2.04E-17 | -0.229795365 | 0.1 | 0.17 | 5.90E-13 |
| RAD51B | 2.07E-17 | -0.298986209 | 0.4 | 0.533 | 5.97E-13 |
| LMAN1 | 2.09E-17 | -0.155897698 | 0.1 | 0.23 | 6.04E-13 |
| RHOG | 2.53E-17 | -0.203656899 | 0.1 | 0.232 | 7.32E-13 |
| SPTAN1 | 2.75E-17 | 0.942577843 | 0.2 | 0.154 | 7.94E-13 |
| MTFR1 | 3.13E-17 | -0.23133331 | 0.2 | 0.299 | 9.05E-13 |
| RASAL1 | 3.33E-17 | -0.260365885 | 0.2 | 0.294 | 9.62E-13 |
| SUCLG2 | 3.70E-17 | -0.218053546 | 0.2 | 0.264 | 1.07E-12 |
| KIF19 | 4.16E-17 | -0.197796727 | 0.2 | 0.272 | 1.20E-12 |
| TATDN1 | 4.49E-17 | -0.200834784 | 0.2 | 0.261 | 1.30E-12 |
| LINC01470 | 4.93E-17 | -0.228782527 | 0.1 | 0.168 | 1.42E-12 |
| DOCK3 | 4.95E-17 | -0.160661745 | 0.6 | 0.819 | 1.43E-12 |
| AC092958.1 | 5.31E-17 | -0.25691365 | 0.1 | 0.255 | 1.54E-12 |
| SORL1 | 5.41E-17 | 0.897462149 | 0.2 | 0.139 | 1.56E-12 |
| ATP8A2 | 5.69E-17 | 0.756125474 | 0.2 | 0.158 | 1.64E-12 |
| AL591895.1 | 6.01E-17 | -0.197643496 | 0.2 | 0.318 | 1.74E-12 |
| THBS2 | 6.10E-17 | -0.255231451 | 0.1 | 0.236 | 1.76E-12 |
| HTATIP2 | 6.63E-17 | -0.151222594 | 0.1 | 0.207 | 1.92E-12 |
| FAM19A4 | 6.70E-17 | -0.324926664 | 0.1 | 0.213 | 1.94E-12 |
| TSGA10 | 6.92E-17 | -0.172268906 | 0.2 | 0.373 | 2.00E-12 |
| TANGO6 | 6.95E-17 | -0.236100869 | 0.2 | 0.32 | 2.01E-12 |
| GTF2H2C | 7.51E-17 | -0.155761799 | 0.2 | 0.273 | 2.17E-12 |
| CDH1 | 7.51E-17 | -0.253339694 | 0.1 | 0.239 | 2.17E-12 |
| MTHFD2L | 7.68E-17 | -0.155405073 | 0.2 | 0.294 | 2.22E-12 |
| TSC22D3 | 8.15E-17 | -0.151601465 | 0.2 | 0.345 | 2.35E-12 |
| SLC45A3 | 8.21E-17 | -0.195527434 | 0.1 | 0.213 | 2.37E-12 |
| ZNF407 | 8.49E-17 | -0.150272968 | 0.3 | 0.393 | 2.45E-12 |
| TNS2 | 9.04E-17 | -0.146568557 | 0.1 | 0.217 | 2.61E-12 |
| AC023590.1 | 9.19E-17 | -0.178126476 | 0.1 | 0.154 | 2.65E-12 |
| GXYLT2 | 9.38E-17 | -0.258743879 | 0.1 | 0.252 | 2.71E-12 |
| PLIN3 | 9.47E-17 | -0.184376033 | 0.1 | 0.179 | 2.74E-12 |
| FAM196B | 1.04E-16 | -0.285276779 | 0.2 | 0.313 | 3.00E-12 |
| SCN8A | 1.26E-16 | 0.721100584 | 0.1 | 0.085 | 3.65E-12 |
| PAG1 | 1.32E-16 | -0.186651975 | 0.2 | 0.316 | 3.82E-12 |
| CDYL | 1.34E-16 | -0.164887129 | 0.2 | 0.341 | 3.88E-12 |
| ERGIC2 | 1.50E-16 | -0.173648537 | 0.2 | 0.276 | 4.34E-12 |
| SFRP1 | 1.52E-16 | -0.24948233 | 0.1 | 0.223 | 4.39E-12 |
| SPARC | 1.53E-16 | -0.179272828 | 0.1 | 0.171 | 4.43E-12 |
| RBAK-RBAKDN | 1.69E-16 | -0.158564238 | 0.1 | 0.257 | 4.88E-12 |
| ARNT | 1.80E-16 | -0.146738641 | 0.2 | 0.261 | 5.20E-12 |
| DPYD-AS1 | 2.05E-16 | -0.209407781 | 0.1 | 0.152 | 5.92E-12 |
| ZNF493 | 2.24E-16 | -0.217373193 | 0.1 | 0.182 | 6.47E-12 |
| CXorf36 | 2.28E-16 | -0.2828955 | 0.1 | 0.182 | 6.60E-12 |
| GBE1 | 2.75E-16 | -0.196658371 | 0.1 | 0.241 | 7.95E-12 |
| EFCAB2 | 2.84E-16 | -0.198836314 | 0.2 | 0.363 | 8.21E-12 |
| MLLT3 | 3.45E-16 | -0.150891536 | 0.3 | 0.477 | 9.97E-12 |
| SQLE | 3.47E-16 | -0.189401151 | 0.1 | 0.232 | 1.00E-11 |
| UCHL1 | 3.90E-16 | 1.266786882 | 0.2 | 0.137 | 1.13E-11 |
| CLSTN2 | 3.96E-16 | 0.638155944 | 0.1 | 0.077 | 1.14E-11 |
| NKD1 | 4.18E-16 | -0.259393864 | 0.1 | 0.155 | 1.21E-11 |
| TMEM125 | 4.31E-16 | -0.166012373 | 0.1 | 0.192 | 1.25E-11 |
| NBPF1 | 4.63E-16 | -0.175855164 | 0.2 | 0.254 | 1.34E-11 |
| AL512625.3 | 4.91E-16 | -0.189145123 | 0.1 | 0.192 | 1.42E-11 |
| COX19 | 5.22E-16 | -0.165231022 | 0.1 | 0.179 | 1.51E-11 |
| CARHSP1 | 5.36E-16 | -0.168501228 | 0.1 | 0.223 | 1.55E-11 |
| HDAC5 | 5.53E-16 | -0.138945295 | 0.1 | 0.24 | 1.60E-11 |
| PPP3CA | 5.66E-16 | 0.872430165 | 0.5 | 0.485 | 1.64E-11 |
| DNAJC1 | 6.02E-16 | -0.181657775 | 0.3 | 0.429 | 1.74E-11 |
| UTY | 6.20E-16 | -0.165462886 | 0.3 | 0.441 | 1.79E-11 |
| SMC6 | 6.55E-16 | -0.17807986 | 0.2 | 0.32 | 1.89E-11 |
| LINC00476 | 6.62E-16 | -0.225012553 | 0.2 | 0.285 | 1.91E-11 |
| ADAM23 | 6.99E-16 | 0.848904132 | 0.2 | 0.138 | 2.02E-11 |
| SH3RF3 | 7.98E-16 | 0.52306072 | 0.1 | 0.053 | 2.31E-11 |
| ADAM12 | 8.62E-16 | -0.227972487 | 0.2 | 0.31 | 2.49E-11 |
| MAD1L1 | 8.85E-16 | -0.21425783 | 0.2 | 0.264 | 2.56E-11 |
| AC087564.1 | 9.39E-16 | -0.180917241 | 0.3 | 0.429 | 2.71E-11 |
| NUP107 | 9.94E-16 | -0.182502953 | 0.1 | 0.238 | 2.87E-11 |
| LINC01099 | 1.01E-15 | -0.553254552 | 0.1 | 0.148 | 2.93E-11 |
| CDKL1 | 1.02E-15 | -0.201398068 | 0.2 | 0.347 | 2.94E-11 |
| TAF3 | 1.06E-15 | -0.193488263 | 0.2 | 0.295 | 3.06E-11 |
| DUSP10 | 1.14E-15 | -0.209585913 | 0.2 | 0.294 | 3.29E-11 |
| CHST3 | 1.15E-15 | -0.156793835 | 0.2 | 0.311 | 3.33E-11 |
| YIPF6 | 1.29E-15 | -0.142726196 | 0.1 | 0.158 | 3.72E-11 |
| SUPT20H | 1.30E-15 | -0.152165295 | 0.2 | 0.269 | 3.75E-11 |
| AC006030.1 | 1.39E-15 | -0.205363528 | 0.1 | 0.244 | 4.02E-11 |
| BARD1 | 1.82E-15 | -0.166536605 | 0.1 | 0.184 | 5.27E-11 |
| SCGB2B2 | 1.95E-15 | -0.192983519 | 0.2 | 0.325 | 5.63E-11 |
| RMDN1 | 2.03E-15 | -0.160572011 | 0.2 | 0.27 | 5.86E-11 |
| KIFC2 | 2.23E-15 | 0.690421841 | 0.1 | 0.052 | 6.43E-11 |
| CETN3 | 2.37E-15 | -0.144201873 | 0.1 | 0.132 | 6.85E-11 |
| RSPRY1 | 2.38E-15 | -0.138783029 | 0.2 | 0.255 | 6.88E-11 |
| CDH12 | 2.53E-15 | 0.911527778 | 0.2 | 0.15 | 7.31E-11 |
| GJC2 | 2.56E-15 | -0.13928424 | 0.1 | 0.166 | 7.40E-11 |
| AATF | 2.65E-15 | -0.150765352 | 0.1 | 0.232 | 7.66E-11 |
| PRPF39 | 2.66E-15 | -0.166808179 | 0.2 | 0.27 | 7.70E-11 |
| WIPI1 | 2.78E-15 | -0.217369961 | 0.1 | 0.193 | 8.04E-11 |
| SLC1A3 | 2.88E-15 | 1.348491332 | 0.6 | 0.625 | 8.32E-11 |
| DISC1 | 3.00E-15 | -0.295857934 | 0.2 | 0.279 | 8.67E-11 |
| GLCE | 3.04E-15 | -0.159352386 | 0.2 | 0.292 | 8.80E-11 |
| DOCK4 | 3.12E-15 | -0.171884212 | 0.7 | 0.917 | 9.02E-11 |
| LRIG3 | 3.16E-15 | -0.173417926 | 0.1 | 0.157 | 9.14E-11 |
| ANKMY1 | 3.18E-15 | -0.175586463 | 0.1 | 0.228 | 9.20E-11 |
| TRIM62 | 3.23E-15 | -0.207807938 | 0.1 | 0.227 | 9.35E-11 |
| FOXP1 | 3.44E-15 | -0.211334534 | 0.5 | 0.718 | 9.95E-11 |
| COL9A3 | 3.50E-15 | -0.157909715 | 0.1 | 0.24 | 1.01E-10 |
| ZNF431 | 3.61E-15 | -0.154434095 | 0.2 | 0.264 | 1.04E-10 |
| AAMDC | 3.92E-15 | -0.167106393 | 0.1 | 0.224 | 1.13E-10 |
| C4orf48 | 3.99E-15 | -0.164145667 | 0.1 | 0.194 | 1.15E-10 |
| CYP20A1 | 4.52E-15 | -0.158218141 | 0.2 | 0.272 | 1.31E-10 |
| NAP1L3 | 4.96E-15 | 0.795672221 | 0.1 | 0.069 | 1.43E-10 |
| AC053527.1 | 5.66E-15 | -0.14669909 | 0.2 | 0.36 | 1.64E-10 |
| GANC | 5.73E-15 | -0.162940616 | 0.2 | 0.348 | 1.66E-10 |
| KCNK10 | 5.83E-15 | -0.261003033 | 0.1 | 0.211 | 1.68E-10 |
| ATXN3 | 6.06E-15 | -0.140337568 | 0.2 | 0.274 | 1.75E-10 |
| PAQR4 | 6.31E-15 | -0.151637255 | 0.1 | 0.246 | 1.82E-10 |
| AC066613.2 | 6.66E-15 | -0.195626489 | 0.1 | 0.206 | 1.93E-10 |
| GTF2F2 | 7.28E-15 | -0.204330599 | 0.2 | 0.294 | 2.10E-10 |
| PHACTR1 | 8.15E-15 | 0.822044636 | 0.4 | 0.365 | 2.36E-10 |
| ZFP14 | 8.49E-15 | -0.151980763 | 0.2 | 0.307 | 2.45E-10 |
| CLCA4 | 8.52E-15 | -0.225026121 | 0.1 | 0.189 | 2.46E-10 |
| TRIM59 | 8.67E-15 | -0.242138973 | 0.1 | 0.177 | 2.51E-10 |

| | | | | | |
|---|---|---|---|---|---|
| GPR158 | 8.71E-15 | 0.711396855 | 0.1 | 0.061 | 2.52E-10 |
| MRPS27 | 8.79E-15 | -0.159610323 | 0.1 | 0.217 | 2.54E-10 |
| TCFL5 | 9.61E-15 | -0.141232862 | 0.1 | 0.216 | 2.78E-10 |
| ATP9A | 9.89E-15 | 0.160372837 | 0.4 | 0.519 | 2.86E-10 |
| PDZRN4 | 1.14E-14 | 0.594834084 | 0.2 | 0.1 | 3.28E-10 |
| C15orf41 | 1.20E-14 | -0.185116705 | 0.2 | 0.331 | 3.46E-10 |
| ATL3 | 1.27E-14 | -0.140400824 | 0.1 | 0.195 | 3.68E-10 |
| RNF114 | 1.33E-14 | -0.173714338 | 0.1 | 0.202 | 3.83E-10 |
| SV2B | 1.39E-14 | 0.901615975 | 0.2 | 0.11 | 4.02E-10 |
| AGMO | 1.50E-14 | -0.331804348 | 0.1 | 0.177 | 4.33E-10 |
| CNTNAP2 | 1.56E-14 | 0.46049459 | 0.8 | 0.719 | 4.52E-10 |
| SLC30A7 | 1.58E-14 | -0.171929814 | 0.1 | 0.247 | 4.56E-10 |
| AC018647.1 | 1.66E-14 | -0.173566052 | 0.1 | 0.2 | 4.80E-10 |
| GPIHBP1 | 1.74E-14 | -0.211625354 | 0.1 | 0.161 | 5.02E-10 |
| METTL16 | 1.78E-14 | -0.161002666 | 0.2 | 0.316 | 5.14E-10 |
| NDUFAF2 | 1.89E-14 | -0.188412746 | 0.2 | 0.283 | 5.47E-10 |
| RPS6KB1 | 1.96E-14 | -0.208214053 | 0.1 | 0.203 | 5.67E-10 |
| SLC16A1 | 1.97E-14 | -0.189259284 | 0.1 | 0.157 | 5.71E-10 |
| HNRNPA3 | 1.99E-14 | 0.180625423 | 0.3 | 0.49 | 5.76E-10 |
| AMPD3 | 2.02E-14 | -0.170149178 | 0.2 | 0.305 | 5.85E-10 |
| AC007364.1 | 2.25E-14 | -0.307569235 | 0.2 | 0.264 | 6.49E-10 |
| EFCAB11 | 2.32E-14 | -0.292722638 | 0.2 | 0.251 | 6.72E-10 |
| HOMER1 | 2.37E-14 | 0.516626913 | 0.1 | 0.058 | 6.84E-10 |
| FZD3 | 2.47E-14 | 0.624442102 | 0.1 | 0.074 | 7.15E-10 |
| ABHD5 | 2.63E-14 | -0.149518485 | 0.1 | 0.159 | 7.59E-10 |
| IPO9-AS1 | 2.72E-14 | -0.165853121 | 0.1 | 0.212 | 7.86E-10 |
| AC233296.1 | 2.84E-14 | -0.346804337 | 0.2 | 0.288 | 8.20E-10 |
| NCALD | 2.99E-14 | 0.812910823 | 0.2 | 0.108 | 8.63E-10 |
| AC073342.1 | 3.69E-14 | -0.161358433 | 0.1 | 0.139 | 1.07E-09 |
| DCAF16 | 3.98E-14 | -0.151884085 | 0.1 | 0.206 | 1.15E-09 |
| LBR | 4.07E-14 | -0.167156818 | 0.1 | 0.226 | 1.18E-09 |
| AC004918.1 | 4.11E-14 | -0.167474228 | 0.1 | 0.206 | 1.19E-09 |
| MRPL48 | 4.17E-14 | -0.153075703 | 0.1 | 0.24 | 1.21E-09 |
| LDLRAP1 | 4.36E-14 | -0.162299758 | 0.2 | 0.249 | 1.26E-09 |
| SLC25A18 | 4.68E-14 | 0.530537248 | 0.1 | 0.068 | 1.35E-09 |
| AC008537.1 | 5.28E-14 | -0.157496208 | 0.1 | 0.244 | 1.53E-09 |
| NOS1 | 5.30E-14 | -0.218886029 | 0.1 | 0.158 | 1.53E-09 |
| CEP85 | 5.34E-14 | -0.174473044 | 0.1 | 0.201 | 1.54E-09 |
| GREM1 | 6.22E-14 | -0.198975921 | 0.2 | 0.296 | 1.80E-09 |
| CEP128 | 6.51E-14 | -0.195154043 | 0.2 | 0.299 | 1.88E-09 |
| PRIMPOL | 6.92E-14 | -0.156558703 | 0.1 | 0.181 | 2.00E-09 |
| KEL | 7.11E-14 | -0.160987457 | 0.1 | 0.179 | 2.06E-09 |
| GPR137C | 7.90E-14 | -0.165654938 | 0.3 | 0.433 | 2.28E-09 |
| RAPGEF4 | 7.94E-14 | 0.50698787 | 0.2 | 0.107 | 2.29E-09 |
| METTL25 | 7.95E-14 | -0.143876131 | 0.2 | 0.307 | 2.30E-09 |
| PCSK1N | 9.40E-14 | 0.998502674 | 0.1 | 0.088 | 2.72E-09 |
| SUMF1 | 9.46E-14 | -0.161391741 | 0.4 | 0.551 | 2.73E-09 |
| TMEM258 | 1.07E-13 | -0.149668561 | 0.1 | 0.179 | 3.08E-09 |
| VPS13A | 1.09E-13 | 0.661992297 | 0.2 | 0.091 | 3.16E-09 |
| ZNF274 | 1.10E-13 | -0.161763203 | 0.1 | 0.192 | 3.18E-09 |
| MATN2 | 1.14E-13 | -0.156301114 | 0.3 | 0.365 | 3.30E-09 |
| EIF1 | 1.29E-13 | 0.267111153 | 0.3 | 0.48 | 3.73E-09 |
| CFDP1 | 1.34E-13 | -0.142451063 | 0.2 | 0.3 | 3.86E-09 |
| SFT2D2 | 1.35E-13 | -0.18453742 | 0.1 | 0.202 | 3.90E-09 |
| LAYN | 1.47E-13 | -0.162159736 | 0.1 | 0.182 | 4.25E-09 |
| MYLIP | 1.51E-13 | -0.144262131 | 0.2 | 0.263 | 4.37E-09 |
| SGK2 | 1.59E-13 | -0.14358516 | 0.1 | 0.158 | 4.59E-09 |
| ZNF253 | 1.62E-13 | -0.161308223 | 0.1 | 0.197 | 4.68E-09 |
| HPN | 1.66E-13 | -0.156171474 | 0.1 | 0.179 | 4.80E-09 |
| AP001347.1 | 1.66E-13 | 0.481375813 | 0.1 | 0.067 | 4.81E-09 |
| LINC01010 | 1.75E-13 | -0.232223824 | 0.1 | 0.124 | 5.06E-09 |
| ST3GAL6 | 1.78E-13 | -0.179748289 | 0.2 | 0.334 | 5.15E-09 |
| SEM1 | 1.99E-13 | -0.150626527 | 0.2 | 0.358 | 5.76E-09 |
| SYT11 | 2.01E-13 | 0.219141886 | 0.5 | 0.62 | 5.81E-09 |
| IP6K3 | 2.11E-13 | -0.168637341 | 0.1 | 0.182 | 6.10E-09 |
| RSU1 | 2.32E-13 | -0.154552664 | 0.1 | 0.241 | 6.70E-09 |
| GALNT18 | 2.49E-13 | 0.661262474 | 0.1 | 0.086 | 7.18E-09 |
| CCZ1B | 2.51E-13 | -0.139639085 | 0.1 | 0.162 | 7.25E-09 |
| MYH10 | 2.55E-13 | 0.710107581 | 0.1 | 0.075 | 7.37E-09 |
| DYNC1H1 | 2.56E-13 | 0.246698138 | 0.4 | 0.522 | 7.40E-09 |
| MT-ND3 | 2.84E-13 | 0.4397786 | 0.7 | 0.785 | 8.21E-09 |
| SAV1 | 2.91E-13 | -0.1808699 | 0.1 | 0.158 | 8.41E-09 |
| GRIA3 | 3.10E-13 | 0.709300758 | 0.2 | 0.145 | 8.96E-09 |
| TAF2 | 3.12E-13 | -0.138396788 | 0.2 | 0.262 | 9.01E-09 |
| APOE | 3.34E-13 | 0.799483032 | 0.1 | 0.086 | 9.65E-09 |
| ITM2B | 3.38E-13 | 0.274821598 | 0.4 | 0.546 | 9.77E-09 |
| MANBA | 3.46E-13 | -0.172983953 | 0.2 | 0.24 | 1.00E-08 |
| HNRNPK | 3.48E-13 | 0.176302864 | 0.3 | 0.417 | 1.01E-08 |
| SPATA5 | 3.84E-13 | -0.170546098 | 0.2 | 0.329 | 1.11E-08 |
| PHF8 | 3.84E-13 | -0.149158688 | 0.1 | 0.211 | 1.11E-08 |
| AL034405.1 | 3.86E-13 | -0.306007698 | 0.1 | 0.234 | 1.12E-08 |
| ITPR1 | 4.31E-13 | 0.721174099 | 0.2 | 0.117 | 1.25E-08 |
| ARRDC2 | 4.89E-13 | -0.137752919 | 0.1 | 0.216 | 1.41E-08 |
| PIGK | 5.15E-13 | -0.183423459 | 0.2 | 0.274 | 1.49E-08 |
| SYP | 5.24E-13 | 0.953398616 | 0.2 | 0.122 | 1.52E-08 |
| RAB6A | 5.27E-13 | 0.159264307 | 0.3 | 0.483 | 1.52E-08 |
| AP002495.2 | 5.60E-13 | -0.198149865 | 0.1 | 0.225 | 1.62E-08 |
| PHGDH | 5.60E-13 | -0.159791211 | 0.1 | 0.226 | 1.62E-08 |
| AC079594.2 | 6.19E-13 | -0.258936244 | 0.1 | 0.228 | 1.79E-08 |
| PGAP1 | 6.64E-13 | -0.144811915 | 0.2 | 0.276 | 1.92E-08 |
| MYRIP | 6.86E-13 | 0.671798415 | 0.2 | 0.145 | 1.98E-08 |
| MIR181A2HG | 6.96E-13 | -0.208853973 | 0.1 | 0.21 | 2.01E-08 |
| TFB1M | 7.16E-13 | -0.144715105 | 0.1 | 0.234 | 2.07E-08 |
| AC069228.1 | 7.43E-13 | -0.204416297 | 0.1 | 0.238 | 2.15E-08 |
| MRPS22 | 8.04E-13 | -0.149430942 | 0.1 | 0.15 | 2.32E-08 |
| FRRS1L | 8.33E-13 | 0.749320152 | 0.1 | 0.069 | 2.41E-08 |
| SMYD3 | 8.43E-13 | 0.52756061 | 0.4 | 0.301 | 2.44E-08 |
| FAT4 | 8.44E-13 | -0.177123465 | 0.2 | 0.273 | 2.44E-08 |
| C2orf92 | 8.75E-13 | -0.166127572 | 0.1 | 0.14 | 2.53E-08 |
| RBM19 | 8.78E-13 | -0.138878988 | 0.1 | 0.171 | 2.54E-08 |
| COPG2 | 8.86E-13 | -0.143189294 | 0.1 | 0.241 | 2.56E-08 |
| HSD11B1 | 8.94E-13 | -0.237945061 | 0.1 | 0.194 | 2.58E-08 |
| LLGL2 | 9.18E-13 | -0.195967426 | 0.1 | 0.196 | 2.65E-08 |
| CPEB1-AS1 | 9.77E-13 | -0.213192798 | 0.1 | 0.191 | 2.82E-08 |
| SON | 1.45E-12 | 0.144982067 | 0.5 | 0.645 | 4.19E-08 |
| FMN1 | 1.55E-12 | 0.703216982 | 0.1 | 0.091 | 4.47E-08 |
| APOLD1 | 1.57E-12 | -0.286163709 | 0.2 | 0.24 | 4.53E-08 |
| NPTX2 | 1.61E-12 | -0.15589247 | 0.1 | 0.141 | 4.65E-08 |
| SERTAD2 | 1.68E-12 | -0.159032537 | 0.1 | 0.232 | 4.84E-08 |
| RPL13 | 2.37E-12 | 0.181526404 | 0.2 | 0.355 | 6.86E-08 |
| PACRG-AS3 | 2.62E-12 | -0.240747025 | 0.1 | 0.2 | 7.56E-08 |
| FAM227B | 2.68E-12 | -0.140046661 | 0.1 | 0.225 | 7.74E-08 |
| XKR6 | 2.74E-12 | 0.576121473 | 0.2 | 0.133 | 7.90E-08 |
| MCC | 2.81E-12 | 0.321882912 | 0.1 | 0.058 | 8.11E-08 |
| SLC30A6 | 2.91E-12 | -0.148825387 | 0.1 | 0.177 | 8.41E-08 |
| NRXN3 | 3.24E-12 | -0.243360763 | 0.8 | 0.888 | 9.38E-08 |
| BAZ1A | 3.45E-12 | -0.14956432 | 0.1 | 0.23 | 9.97E-08 |
| RNF138 | 3.56E-12 | -0.151712412 | 0.1 | 0.187 | 1.03E-07 |
| HSPA1A | 3.90E-12 | 0.175831202 | 0.2 | 0.284 | 1.13E-07 |
| SYT16 | 4.01E-12 | 0.527241169 | 0.1 | 0.081 | 1.16E-07 |
| POLR3G | 4.22E-12 | -0.139992892 | 0.1 | 0.158 | 1.22E-07 |
| ENPP6 | 4.75E-12 | -0.194393023 | 0.1 | 0.139 | 1.37E-07 |
| ALG14 | 5.17E-12 | -0.143018372 | 0.1 | 0.198 | 1.50E-07 |
| ZNF727 | 5.51E-12 | -0.158989373 | 0.1 | 0.118 | 1.59E-07 |
| AC027808.1 | 7.29E-12 | -0.150040128 | 0.1 | 0.162 | 2.11E-07 |
| MSH6 | 7.32E-12 | -0.146584056 | 0.1 | 0.202 | 2.12E-07 |
| PDSS2 | 7.68E-12 | -0.165859062 | 0.2 | 0.314 | 2.22E-07 |
| OSBPL5 | 8.44E-12 | -0.176088113 | 0.1 | 0.192 | 2.44E-07 |
| ATP1B1 | 8.70E-12 | 0.339480004 | 0.5 | 0.743 | 2.51E-07 |
| LINC00601 | 1.01E-11 | -0.154639515 | 0.1 | 0.173 | 2.91E-07 |
| CLSTN1 | 1.07E-11 | 1.232136598 | 0.4 | 0.331 | 3.10E-07 |
| SEMA5A | 1.21E-11 | -0.194257892 | 0.2 | 0.248 | 3.50E-07 |
| MAST4 | 1.21E-11 | 0.739345596 | 0.3 | 0.237 | 3.51E-07 |
| C8orf59 | 1.22E-11 | -0.163777544 | 0.1 | 0.127 | 3.52E-07 |
| DPY19L2 | 1.31E-11 | 0.505079758 | 0.1 | 0.063 | 3.78E-07 |
| TAF5L | 1.37E-11 | -0.152518289 | 0.1 | 0.134 | 3.96E-07 |
| AL160269.1 | 1.48E-11 | -0.1488747 | 0.1 | 0.144 | 4.29E-07 |
| AC110285.2 | 1.53E-11 | -0.151468731 | 0.2 | 0.239 | 4.43E-07 |

| | | | | | |
|---|---|---|---|---|---|
| ANKRD18A | 1.57E-11 | -0.218336525 | 0.1 | 0.188 | 4.53E-07 |
| CHSY3 | 1.58E-11 | 0.391701948 | 0.2 | 0.12 | 4.56E-07 |
| MFSD14B | 1.64E-11 | -0.171640966 | 0.1 | 0.174 | 4.73E-07 |
| SHANK2 | 1.67E-11 | 0.507186352 | 0.1 | 0.081 | 4.83E-07 |
| ADAMTSL3 | 1.69E-11 | -0.197224943 | 0.2 | 0.313 | 4.87E-07 |
| PRCD | 2.14E-11 | -0.139661922 | 0 | 0.106 | 6.20E-07 |
| TAX1BP1 | 2.21E-11 | 0.150086061 | 0.3 | 0.438 | 6.38E-07 |
| AL078590.2 | 3.12E-11 | -0.179374344 | 0.1 | 0.168 | 9.03E-07 |
| LINC01877 | 3.17E-11 | -0.196987234 | 0.1 | 0.141 | 9.16E-07 |
| CDKN2A | 3.22E-11 | -0.184084673 | 0.1 | 0.13 | 9.31E-07 |
| PRRC2B | 3.38E-11 | 0.227711836 | 0.3 | 0.449 | 9.77E-07 |
| IMMP1L | 3.41E-11 | -0.142793013 | 0.2 | 0.238 | 9.85E-07 |
| CCNE2 | 3.48E-11 | -0.155457384 | 0.1 | 0.116 | 1.01E-06 |
| FANCB | 3.94E-11 | -0.188743891 | 0.1 | 0.146 | 1.14E-06 |
| AF241726.2 | 4.74E-11 | -0.207716064 | 0.1 | 0.189 | 1.37E-06 |
| SEPSECS | 5.19E-11 | -0.144660504 | 0 | 0.106 | 1.50E-06 |
| ARSB | 5.20E-11 | -0.166581046 | 0.2 | 0.244 | 1.50E-06 |
| KSR1 | 5.35E-11 | -0.172644415 | 0.1 | 0.23 | 1.55E-06 |
| CIT | 6.84E-11 | 0.70225084 | 0.1 | 0.08 | 1.98E-06 |
| CCDC146 | 7.41E-11 | -0.170215212 | 0.2 | 0.282 | 2.14E-06 |
| OGFRL1 | 8.02E-11 | 0.623435674 | 0.1 | 0.09 | 2.32E-06 |
| FAM49A | 8.03E-11 | 0.617270088 | 0.2 | 0.104 | 2.32E-06 |
| SFXN5 | 8.32E-11 | 0.765076798 | 0.2 | 0.162 | 2.40E-06 |
| REST | 9.13E-11 | -0.155480396 | 0.1 | 0.15 | 2.64E-06 |
| IPO11 | 1.01E-10 | -0.1529848 | 0.1 | 0.169 | 2.92E-06 |
| PSG8-AS1 | 1.04E-10 | -0.150761609 | 0.1 | 0.108 | 3.00E-06 |
| PKM | 1.06E-10 | 1.085834043 | 0.2 | 0.166 | 3.06E-06 |
| APLF | 1.12E-10 | -0.171161399 | 0.1 | 0.172 | 3.24E-06 |
| OPHN1 | 1.20E-10 | 0.770917973 | 0.3 | 0.196 | 3.46E-06 |
| CHRAC1 | 1.21E-10 | -0.153359914 | 0.1 | 0.109 | 3.50E-06 |
| HECTD4 | 1.27E-10 | 0.141792031 | 0.3 | 0.475 | 3.68E-06 |
| EEF1A1 | 1.33E-10 | 0.220229179 | 0.3 | 0.385 | 3.84E-06 |
| PPFIA1 | 1.34E-10 | 0.202466166 | 0.2 | 0.357 | 3.87E-06 |
| CANX | 1.34E-10 | 0.182486916 | 0.3 | 0.392 | 3.88E-06 |
| ATP2B1 | 1.35E-10 | 1.071448739 | 0.4 | 0.31 | 3.91E-06 |
| VLDLR | 1.44E-10 | -0.147267506 | 0.1 | 0.163 | 4.16E-06 |
| CLCN3 | 1.88E-10 | 0.157699345 | 0.3 | 0.42 | 5.44E-06 |
| SAMD4A | 2.25E-10 | 0.717951353 | 0.2 | 0.157 | 6.50E-06 |
| TMEM30A | 2.50E-10 | 0.197404927 | 0.2 | 0.273 | 7.23E-06 |
| TCERG1 | 2.59E-10 | 0.147802718 | 0.3 | 0.367 | 7.47E-06 |
| NDFIP1 | 2.64E-10 | 0.256501742 | 0.4 | 0.484 | 7.63E-06 |
| TMEM245 | 3.13E-10 | 0.175883802 | 0.3 | 0.379 | 9.04E-06 |
| AC113414.1 | 3.23E-10 | -0.23130965 | 0.1 | 0.186 | 9.34E-06 |
| RNF125 | 3.30E-10 | -0.145513164 | 0.1 | 0.188 | 9.52E-06 |
| HERC1 | 3.43E-10 | 0.176636643 | 0.5 | 0.676 | 9.91E-06 |
| ARL8B | 3.71E-10 | 0.179751299 | 0.2 | 0.337 | 1.07E-05 |
| EPS8 | 4.50E-10 | -0.239473319 | 0.2 | 0.23 | 1.30E-05 |
| TCF25 | 4.61E-10 | 0.151768443 | 0.3 | 0.453 | 1.33E-05 |
| PRKAR1A | 4.63E-10 | 0.212320538 | 0.2 | 0.334 | 1.34E-05 |
| PI4KA | 4.95E-10 | 0.181770938 | 0.4 | 0.491 | 1.43E-05 |
| CAVIN4 | 4.99E-10 | -0.140487132 | 0.1 | 0.118 | 1.44E-05 |
| SMARCAL1 | 5.84E-10 | -0.156157177 | 0.1 | 0.203 | 1.69E-05 |
| TMEM38B | 5.89E-10 | -0.153928346 | 0.1 | 0.138 | 1.70E-05 |
| TSPYL2 | 6.06E-10 | 0.732467201 | 0.2 | 0.107 | 1.75E-05 |
| AHCYL2 | 6.13E-10 | 0.65715275 | 0.3 | 0.27 | 1.77E-05 |
| SLC9B1 | 6.26E-10 | -0.162351787 | 0.1 | 0.154 | 1.81E-05 |
| MAGEC3 | 6.89E-10 | -0.162713711 | 0.1 | 0.163 | 1.99E-05 |
| MGMT | 6.96E-10 | -0.154684367 | 0.1 | 0.175 | 2.01E-05 |
| MAP2K6 | 7.62E-10 | -0.142105299 | 0.1 | 0.182 | 2.20E-05 |
| LINC02177 | 7.91E-10 | -0.143196589 | 0.1 | 0.179 | 2.28E-05 |
| LINC01299 | 8.65E-10 | -0.193645757 | 0.1 | 0.137 | 2.50E-05 |
| GNAS | 1.34E-09 | 1.223868506 | 0.4 | 0.384 | 3.88E-05 |
| GUK1 | 1.37E-09 | 0.180734823 | 0.1 | 0.216 | 3.96E-05 |
| PPP2CB | 1.48E-09 | 0.147133802 | 0.2 | 0.287 | 4.29E-05 |
| PRRC2C | 1.56E-09 | 0.21685224 | 0.5 | 0.678 | 4.50E-05 |
| LMO4 | 1.89E-09 | 1.056217725 | 0.2 | 0.157 | 5.45E-05 |
| ADAMTS1 | 1.96E-09 | -0.144934949 | 0.1 | 0.129 | 5.65E-05 |
| LINC02226 | 2.01E-09 | -0.166861489 | 0 | 0.102 | 5.80E-05 |
| SLC35F3 | 2.25E-09 | 0.447390317 | 0.1 | 0.084 | 6.50E-05 |
| CSMD3 | 2.28E-09 | 0.775638059 | 0.4 | 0.334 | 6.60E-05 |
| SERINC3 | 2.46E-09 | 0.20008193 | 0.2 | 0.33 | 7.10E-05 |
| SLC6A1 | 2.49E-09 | 0.246414757 | 0.4 | 0.517 | 7.19E-05 |
| NAT8L | 2.66E-09 | 0.73537326 | 0.1 | 0.099 | 7.68E-05 |
| RPS15 | 3.49E-09 | 0.202227542 | 0.2 | 0.29 | 0.000100908 |
| PPP1R12A | 3.58E-09 | 0.141474357 | 0.4 | 0.485 | 0.000103325 |
| GRK3 | 3.59E-09 | 0.525405862 | 0.2 | 0.143 | 0.000103803 |
| SV2A | 3.80E-09 | 0.807520402 | 0.1 | 0.09 | 0.000109916 |
| AP000282.1 | 4.00E-09 | -0.228237033 | 0.1 | 0.18 | 0.000115516 |
| EIF5 | 4.24E-09 | 0.18508797 | 0.2 | 0.297 | 0.000122461 |
| OGA | 4.32E-09 | 0.162131043 | 0.4 | 0.498 | 0.000124882 |
| SESN3 | 5.14E-09 | 0.598427874 | 0.1 | 0.08 | 0.000148455 |
| EDA | 5.29E-09 | -0.270783817 | 0.2 | 0.223 | 0.00015284 |
| NIPSNAP2 | 5.45E-09 | 0.138112976 | 0.2 | 0.296 | 0.000157504 |
| C11orf58 | 5.86E-09 | 0.145709641 | 0.2 | 0.315 | 0.000169298 |
| C11orf65 | 6.15E-09 | -0.17942115 | 0.1 | 0.158 | 0.000177623 |
| NR1D2 | 6.44E-09 | 0.200072064 | 0.2 | 0.332 | 0.00018606 |
| LINC01322 | 6.61E-09 | 0.479644096 | 0.1 | 0.07 | 0.000191076 |
| FYN | 6.64E-09 | 0.172578603 | 0.5 | 0.675 | 0.000191996 |
| TMEM126B | 6.91E-09 | -0.141229321 | 0.1 | 0.113 | 0.000199788 |
| AC104461.1 | 7.03E-09 | -0.138347403 | 0.1 | 0.11 | 0.000203291 |
| CPB1 | 7.06E-09 | -0.163592032 | 0.1 | 0.164 | 0.000204176 |
| XRCC6 | 7.73E-09 | 0.187931457 | 0.1 | 0.203 | 0.000223489 |
| NKAIN1 | 9.35E-09 | -0.177591914 | 0.1 | 0.174 | 0.000270182 |
| STPG2 | 1.06E-08 | -0.189676274 | 0.2 | 0.263 | 0.000307522 |
| LRP1 | 1.17E-08 | 0.581216969 | 0.1 | 0.096 | 0.000338302 |
| FADS2 | 1.28E-08 | 0.144430742 | 0.1 | 0.155 | 0.000370032 |
| FAM241A | 1.32E-08 | -0.168420422 | 0.1 | 0.131 | 0.000382378 |
| AL445250.1 | 1.35E-08 | -0.184732401 | 0.2 | 0.245 | 0.000391444 |
| TMOD2 | 1.35E-08 | 0.170634678 | 0.3 | 0.409 | 0.000391502 |
| GALNT2 | 1.37E-08 | 0.222129214 | 0.2 | 0.3 | 0.0003969 |
| ARID4A | 1.37E-08 | 0.183796811 | 0.3 | 0.346 | 0.000396905 |
| ADAMTS18 | 1.38E-08 | -0.303450733 | 0.1 | 0.184 | 0.000400037 |
| LINC01006 | 1.46E-08 | -0.162034597 | 0.1 | 0.176 | 0.00042154 |
| SCAMP1 | 1.56E-08 | 0.156761111 | 0.2 | 0.23 | 0.000449747 |
| SDK1 | 1.64E-08 | 0.306217964 | 0.1 | 0.073 | 0.000473735 |
| SREK1 | 1.86E-08 | 0.1591645 | 0.3 | 0.357 | 0.000538945 |
| KHDRBS1 | 1.92E-08 | 0.159268274 | 0.3 | 0.351 | 0.000554331 |
| AAK1 | 2.17E-08 | 0.223207093 | 0.4 | 0.487 | 0.000626147 |
| JADE3 | 2.27E-08 | -0.181080828 | 0.1 | 0.133 | 0.000656791 |
| ATP6V0A1 | 2.67E-08 | 0.317628918 | 0.3 | 0.421 | 0.000772843 |
| TRIO | 2.88E-08 | 0.698285787 | 0.4 | 0.366 | 0.00083332 |
| ACTG1 | 3.25E-08 | 0.462595008 | 0.3 | 0.388 | 0.000939528 |
| FAM13A | 3.39E-08 | 0.541393344 | 0.1 | 0.094 | 0.000980067 |
| DNAJB14 | 3.39E-08 | 0.137984267 | 0.3 | 0.353 | 0.00098016 |
| ATP6V0B | 3.71E-08 | 0.163254282 | 0.2 | 0.245 | 0.00107168 |
| XRCC4 | 3.78E-08 | -0.13848385 | 0.1 | 0.212 | 0.001093775 |
| DZIP3 | 3.80E-08 | 0.190072014 | 0.2 | 0.337 | 0.001098302 |
| PFKL | 3.90E-08 | 0.15952368 | 0.1 | 0.214 | 0.0011272 |
| MIR4300HG | 4.07E-08 | 0.493999826 | 0.1 | 0.062 | 0.001175654 |
| AC022075.1 | 4.13E-08 | -0.137873722 | 0.1 | 0.102 | 0.001194113 |
| TMEM106B | 4.24E-08 | 0.18538758 | 0.2 | 0.318 | 0.001225659 |
| CAMK2N1 | 5.40E-08 | 0.356013989 | 0.4 | 0.567 | 0.001560621 |
| DCC | 5.74E-08 | 0.299037101 | 0.3 | 0.222 | 0.001658474 |
| CSPP1 | 5.74E-08 | 0.195840863 | 0.3 | 0.353 | 0.001660227 |
| EPB41L3 | 5.77E-08 | 0.231414806 | 0.2 | 0.34 | 0.001666744 |
| ARHGAP35 | 5.85E-08 | 0.140394258 | 0.3 | 0.356 | 0.001689999 |
| VGLL4 | 6.06E-08 | 0.190645778 | 0.2 | 0.34 | 0.001752731 |
| CAB39 | 6.51E-08 | 0.168287214 | 0.2 | 0.303 | 0.001881444 |
| LINC00632 | 6.71E-08 | 0.195764667 | 0.3 | 0.423 | 0.001938287 |
| RAB18 | 6.96E-08 | 0.141113862 | 0.2 | 0.241 | 0.002012716 |
| PDPK1 | 7.89E-08 | 0.187481965 | 0.2 | 0.31 | 0.0022791 |
| GUCY1A2 | 7.99E-08 | 0.519267277 | 0.2 | 0.108 | 0.002310512 |
| ZC4H2 | 8.54E-08 | -0.147106221 | 0.1 | 0.206 | 0.002467281 |
| FAM135B | 8.78E-08 | 0.488621276 | 0.1 | 0.081 | 0.002536462 |
| IREB2 | 9.13E-08 | 0.150340048 | 0.2 | 0.286 | 0.00263853 |
| NDST3 | 9.27E-08 | 0.537298627 | 0.1 | 0.071 | 0.002680287 |
| MARF1 | 9.29E-08 | 0.143574142 | 0.2 | 0.271 | 0.002683723 |
| TLL2 | 9.43E-08 | -0.157489269 | 0.1 | 0.143 | 0.002724865 |

| | | | | | |
|---|---|---|---|---|---|
| TMEM232 | 9.84E-08 | 0.595253246 | 0.2 | 0.172 | 0.002844876 |
| GMDS | 1.07E-07 | -0.18839573 | 0.2 | 0.226 | 0.003089132 |
| PCSK5 | 1.14E-07 | 0.300088753 | 0.1 | 0.074 | 0.003289795 |
| IMMP2L | 1.16E-07 | -0.138157431 | 0.4 | 0.526 | 0.003341939 |
| IDS | 1.18E-07 | 1.293116307 | 0.3 | 0.294 | 0.003404364 |
| RIMKLB | 1.18E-07 | 0.291806225 | 0.1 | 0.064 | 0.003423906 |
| PCNT | 1.27E-07 | 0.160430449 | 0.1 | 0.212 | 0.003660607 |
| EYA4 | 1.42E-07 | -0.161199473 | 0.1 | 0.156 | 0.004111242 |
| DNAJB6 | 1.51E-07 | 0.204754245 | 0.3 | 0.362 | 0.004362267 |
| SELENOI | 1.66E-07 | 0.143197808 | 0.2 | 0.221 | 0.004809908 |
| PNPLA8 | 1.70E-07 | 0.152092302 | 0.2 | 0.255 | 0.00492361 |
| NIN | 1.80E-07 | 0.189136622 | 0.3 | 0.369 | 0.00520142 |
| LONRF2 | 1.92E-07 | 0.703910193 | 0.2 | 0.141 | 0.005545632 |
| ACP7 | 2.01E-07 | -0.14461894 | 0.1 | 0.1 | 0.005812607 |
| TERF2IP | 2.06E-07 | 0.203967792 | 0.4 | 0.457 | 0.005947146 |
| LURAP1L | 2.07E-07 | -0.192830259 | 0.1 | 0.168 | 0.005989004 |
| ATRX | 2.33E-07 | 0.198181646 | 0.5 | 0.707 | 0.006740378 |
| USP22 | 2.50E-07 | 0.317327619 | 0.3 | 0.394 | 0.007234966 |
| ARL6IP5 | 2.54E-07 | 0.255191941 | 0.3 | 0.345 | 0.0073472 |
| YWHAE | 2.61E-07 | 0.284026689 | 0.3 | 0.434 | 0.007546715 |
| DST | 2.82E-07 | 0.250455388 | 0.8 | 0.978 | 0.008139287 |
| FRK | 2.89E-07 | -0.151461547 | 0.1 | 0.156 | 0.008354304 |
| PWAR6 | 2.90E-07 | 0.918889557 | 0.3 | 0.209 | 0.00838374 |
| SUN1 | 3.34E-07 | 0.175297041 | 0.2 | 0.307 | 0.009655968 |
| PRIM2 | 3.48E-07 | -0.137830533 | 0.1 | 0.196 | 0.010057021 |
| TIAM2 | 3.62E-07 | 0.139128925 | 0.3 | 0.362 | 0.010469988 |
| AHCYL1 | 3.68E-07 | 0.889119225 | 0.3 | 0.284 | 0.010648337 |
| MORF4L1 | 3.98E-07 | 0.273306296 | 0.3 | 0.398 | 0.011513342 |
| CALR | 4.14E-07 | 0.142813404 | 0.1 | 0.198 | 0.011962567 |
| EID1 | 4.26E-07 | 0.321390337 | 0.2 | 0.281 | 0.012297765 |
| ITGAV | 4.31E-07 | 0.185042271 | 0.2 | 0.307 | 0.012450595 |
| DOCK7 | 4.40E-07 | 0.647618875 | 0.2 | 0.155 | 0.012707336 |
| RPL6 | 5.06E-07 | 0.158028145 | 0.1 | 0.17 | 0.014611218 |
| AC109927.1 | 5.23E-07 | -0.138225052 | 0.1 | 0.168 | 0.015129634 |
| LYST | 5.56E-07 | 0.143744716 | 0.3 | 0.365 | 0.016077178 |
| CSRNP3 | 5.57E-07 | 0.698461826 | 0.3 | 0.204 | 0.016104194 |
| 11-Sep | 5.69E-07 | 0.469119413 | 0.1 | 0.098 | 0.016447974 |
| APOO | 5.87E-07 | 0.194301508 | 0.2 | 0.249 | 0.016963823 |
| CNTN3 | 6.06E-07 | 0.470782827 | 0.1 | 0.082 | 0.017499982 |
| PI4KB | 6.21E-07 | 0.151679749 | 0.2 | 0.225 | 0.017935572 |
| RYR2 | 6.37E-07 | 0.777046526 | 0.4 | 0.332 | 0.018421942 |
| DGCR2 | 6.55E-07 | 0.174685107 | 0.2 | 0.252 | 0.01893578 |
| MAP1B | 6.86E-07 | 1.167200079 | 0.5 | 0.551 | 0.019821506 |
| ETV5 | 7.68E-07 | -0.149670847 | 0.1 | 0.128 | 0.022208967 |
| ALK | 8.24E-07 | 0.50382494 | 0.1 | 0.07 | 0.023823761 |
| PLEKHA6 | 1.17E-06 | 0.481538322 | 0.1 | 0.094 | 0.033782913 |
| RASAL2 | 1.27E-06 | 0.479359013 | 0.3 | 0.214 | 0.036733369 |
| CCNI | 1.32E-06 | 0.286217078 | 0.2 | 0.331 | 0.038101979 |
| ZFAND5 | 1.41E-06 | 0.289199486 | 0.2 | 0.32 | 0.040859773 |
| BMPR1A | 1.44E-06 | 0.202013566 | 0.2 | 0.32 | 0.041563186 |
| ABCA3 | 1.47E-06 | 0.148360232 | 0.2 | 0.229 | 0.042611188 |
| IPO5 | 1.58E-06 | 0.16537213 | 0.2 | 0.22 | 0.04561806 |
| CLTC | 1.63E-06 | 0.356791292 | 0.3 | 0.452 | 0.047056963 |
| HTT | 1.65E-06 | 0.180154965 | 0.3 | 0.339 | 0.047784807 |
| ASAP1 | 1.83E-06 | 0.18022545 | 0.4 | 0.456 | 0.052924267 |
| NCKAP1 | 1.95E-06 | 0.211350072 | 0.3 | 0.427 | 0.056490119 |
| DDX3X | 2.02E-06 | 0.248655771 | 0.2 | 0.295 | 0.058456664 |
| SKIL | 2.04E-06 | 0.15451942 | 0.1 | 0.14 | 0.059047629 |
| DCAF5 | 2.35E-06 | 0.145089927 | 0.2 | 0.3 | 0.067911896 |
| SDHC | 2.36E-06 | 0.141778693 | 0.1 | 0.191 | 0.068141339 |
| SHOC2 | 2.58E-06 | 0.198976116 | 0.2 | 0.266 | 0.07463186 |
| TTC7B | 2.94E-06 | 0.190801756 | 0.3 | 0.346 | 0.08485942 |
| CLIP1 | 3.24E-06 | 0.735536483 | 0.2 | 0.2 | 0.093511846 |
| CDH4 | 3.27E-06 | 0.467746013 | 0.1 | 0.086 | 0.094470065 |
| AEBP2 | 3.35E-06 | 0.260454319 | 0.3 | 0.377 | 0.096768926 |
| ITM2C | 3.36E-06 | 0.738141518 | 0.1 | 0.106 | 0.097176918 |
| DDX42 | 3.41E-06 | 0.172438083 | 0.1 | 0.193 | 0.098675326 |
| EEF2K | 3.49E-06 | 0.279995654 | 0.1 | 0.075 | 0.100820713 |
| MTCH1 | 3.50E-06 | 0.267723976 | 0.3 | 0.387 | 0.101125603 |
| RHEB | 3.62E-06 | 0.154007289 | 0.1 | 0.194 | 0.104714227 |
| HSPA9 | 3.62E-06 | 0.165256642 | 0.2 | 0.271 | 0.104752011 |
| RAPGEF3 | 3.69E-06 | 0.185175358 | 0.2 | 0.318 | 0.106764432 |
| HSP90AA1 | 3.74E-06 | 0.310186801 | 0.6 | 0.816 | 0.108167819 |
| SACS | 3.90E-06 | 0.233973099 | 0.2 | 0.285 | 0.112578185 |
| ENO2 | 4.16E-06 | 1.115770098 | 0.2 | 0.184 | 0.120288165 |
| ABCA5 | 4.40E-06 | 0.227182485 | 0.2 | 0.308 | 0.127034785 |
| CNBP | 4.54E-06 | 0.165352511 | 0.2 | 0.231 | 0.131271652 |
| MRAS | 4.95E-06 | 0.409164447 | 0.1 | 0.079 | 0.142971512 |
| ATPAF1 | 5.08E-06 | 0.143480761 | 0.1 | 0.201 | 0.146747819 |
| BRINP2 | 5.19E-06 | 0.325644533 | 0.1 | 0.069 | 0.150025471 |
| OSBPL8 | 5.22E-06 | 0.176980745 | 0.3 | 0.363 | 0.150835921 |
| BEX1 | 5.33E-06 | 0.814780498 | 0.1 | 0.083 | 0.153994332 |
| NDFIP2 | 5.48E-06 | 0.156891672 | 0.3 | 0.336 | 0.158275954 |
| PCMT1 | 5.74E-06 | 0.247689562 | 0.2 | 0.264 | 0.165913858 |
| ZNF248 | 5.82E-06 | 0.140431611 | 0.3 | 0.372 | 0.16827473 |
| AC034195.1 | 5.93E-06 | -0.205624663 | 0.1 | 0.176 | 0.171250418 |
| EXOSC6 | 6.20E-06 | 0.163041439 | 0.1 | 0.122 | 0.179198268 |
| NAP1L1 | 6.27E-06 | 0.323814888 | 0.3 | 0.431 | 0.181335466 |
| PPIG | 6.30E-06 | 0.223926131 | 0.3 | 0.359 | 0.182136909 |
| CTNND2 | 6.85E-06 | 0.347816908 | 0.8 | 0.856 | 0.198013589 |
| B3GAT3 | 7.62E-06 | 0.184340111 | 0.1 | 0.152 | 0.22034604 |
| DPYSL2 | 7.92E-06 | 0.239698592 | 0.6 | 0.786 | 0.228766601 |
| ABCA6 | 8.18E-06 | -0.146675726 | 0.1 | 0.179 | 0.23649894 |
| MACF1 | 8.88E-06 | 0.797185708 | 0.7 | 0.892 | 0.256576179 |
| KAT2A | 9.09E-06 | 0.175999987 | 0.2 | 0.217 | 0.262842272 |
| GCC2 | 9.50E-06 | 0.197315987 | 0.2 | 0.311 | 0.274666578 |
| DYNLL2 | 9.68E-06 | 0.203065235 | 0.1 | 0.17 | 0.279742671 |
| MYO9A | 1.03E-05 | 0.20051808 | 0.4 | 0.516 | 0.297020978 |
| PRICKLE2 | 1.03E-05 | 0.486380985 | 0.2 | 0.113 | 0.297511815 |
| TMEM33 | 1.10E-05 | 0.153176021 | 0.1 | 0.193 | 0.31786511 |
| RUBCN | 1.15E-05 | 0.149349739 | 0.1 | 0.195 | 0.331188301 |
| ATP6V0E2 | 1.17E-05 | 0.233502689 | 0.1 | 0.167 | 0.336998201 |
| AKAP13 | 1.21E-05 | 0.27624877 | 0.1 | 0.072 | 0.350351687 |
| NAA20 | 1.41E-05 | 0.138525937 | 0.1 | 0.114 | 0.406347203 |
| CELF1 | 1.41E-05 | 0.258648089 | 0.3 | 0.393 | 0.407912995 |
| TPD52L2 | 1.46E-05 | 0.21798728 | 0.1 | 0.168 | 0.421029506 |
| GABBR1 | 1.46E-05 | 0.685765089 | 0.2 | 0.13 | 0.421386293 |
| TUBA1B | 1.52E-05 | 1.234717327 | 0.3 | 0.222 | 0.437974223 |
| IFNGR2 | 1.56E-05 | 0.17554736 | 0.1 | 0.133 | 0.450096015 |
| BDP1 | 1.58E-05 | 0.262687381 | 0.4 | 0.452 | 0.456518166 |
| CTDSPL | 1.62E-05 | 0.424904063 | 0.1 | 0.105 | 0.469633985 |
| NUMB | 1.63E-05 | 0.179167709 | 0.3 | 0.348 | 0.471753459 |
| CLPTM1 | 1.67E-05 | 0.284023591 | 0.1 | 0.183 | 0.481832351 |
| RAB28 | 1.69E-05 | 0.137852868 | 0.2 | 0.311 | 0.488714317 |
| PTK2B | 1.86E-05 | 0.633463737 | 0.1 | 0.104 | 0.536385532 |
| ATP13A4 | 1.86E-05 | 0.440604985 | 0.2 | 0.118 | 0.538427091 |
| APC | 1.92E-05 | 0.28074897 | 0.4 | 0.454 | 0.555473394 |
| AKAP8L | 1.92E-05 | 0.182730882 | 0.2 | 0.289 | 0.556032318 |
| REEP5 | 2.08E-05 | 0.391568419 | 0.2 | 0.271 | 0.600281108 |
| ZNF25 | 2.08E-05 | 0.166967367 | 0.1 | 0.187 | 0.600419028 |
| SCAMP5 | 2.16E-05 | 0.612396114 | 0.2 | 0.146 | 0.623350733 |
| EIF1AX | 2.22E-05 | 0.293533834 | 0.2 | 0.269 | 0.64040415 |
| HIPK1 | 2.29E-05 | 0.142115847 | 0.1 | 0.162 | 0.661940697 |
| RTN1 | 2.30E-05 | 0.987742852 | 0.5 | 0.454 | 0.66471417 |
| RPS6KA2 | 2.36E-05 | 0.723772511 | 0.5 | 0.457 | 0.682086617 |
| MYEF2 | 2.49E-05 | 0.249989194 | 0.1 | 0.195 | 0.719513905 |
| HSPA12A | 2.56E-05 | 0.236018596 | 0.3 | 0.362 | 0.739135483 |
| YTHDF2 | 2.57E-05 | 0.154767729 | 0.1 | 0.145 | 0.742770316 |
| FAM66D | 2.60E-05 | 0.140546965 | 0.2 | 0.305 | 0.751859969 |
| BICD1 | 2.61E-05 | 0.210743341 | 0.4 | 0.528 | 0.754958418 |
| SRP9 | 2.65E-05 | 0.181121378 | 0.1 | 0.19 | 0.76710441 |
| ZMAT1 | 2.67E-05 | 0.344269293 | 0.3 | 0.437 | 0.77043082 |
| CYFIP2 | 2.67E-05 | 0.765412805 | 0.2 | 0.212 | 0.772595221 |
| ACACB | 2.68E-05 | 0.160965998 | 0.2 | 0.29 | 0.775101772 |
| BTBD8 | 2.88E-05 | 0.249881682 | 0.3 | 0.419 | 0.833494024 |
| CELSR2 | 2.99E-05 | 0.276452491 | 0.2 | 0.242 | 0.865115882 |
| UNC79 | 3.04E-05 | 0.164338836 | 0.3 | 0.384 | 0.877656921 |
| PGAP2 | 3.06E-05 | 0.220187464 | 0.1 | 0.15 | 0.883367251 |

| | | | | | |
|---|---|---|---|---|---|
| CCT4 | 3.10E-05 | 0.146693669 | 0.1 | 0.2 | 0.894810928 |
| CKB | 3.20E-05 | 0.95543465 | 0.5 | 0.487 | 0.925870908 |
| NUPL2 | 3.22E-05 | 0.138292352 | 0.1 | 0.156 | 0.929554893 |
| JARID2 | 3.22E-05 | 0.191941416 | 0.3 | 0.436 | 0.931918298 |
| TP53BP1 | 3.26E-05 | 0.168195178 | 0.2 | 0.27 | 0.940966999 |
| NLGN4X | 3.29E-05 | 0.589444913 | 0.2 | 0.138 | 0.949844548 |
| UBE2N | 3.30E-05 | 0.156385951 | 0.1 | 0.179 | 0.954223241 |
| MGST3 | 3.35E-05 | 0.158883019 | 0.2 | 0.211 | 0.968672715 |
| CCDC92 | 3.37E-05 | 0.224302376 | 0.3 | 0.328 | 0.972808101 |
| ASPH | 3.37E-05 | 0.7686992 | 0.3 | 0.283 | 0.974318278 |
| MTRNR2L8 | 3.40E-05 | 0.383083994 | 0.1 | 0.076 | 0.9816518 |
| ADAM22 | 3.72E-05 | 0.214554753 | 0.3 | 0.376 | 1 |
| TCF7L2 | 3.79E-05 | 0.585414216 | 0.2 | 0.205 | 1 |
| AC124312.1 | 3.86E-05 | 1.028030946 | 0.5 | 0.486 | 1 |
| DSTN | 3.94E-05 | 0.238134476 | 0.1 | 0.176 | 1 |
| CFL1 | 4.21E-05 | 0.481646971 | 0.2 | 0.306 | 1 |
| ITPKB | 4.34E-05 | 0.213713482 | 0.4 | 0.476 | 1 |
| RPL32 | 4.60E-05 | 0.179446092 | 0.1 | 0.191 | 1 |
| ABL2 | 4.66E-05 | 0.192222698 | 0.2 | 0.281 | 1 |
| ZBTB4 | 4.70E-05 | 0.211461951 | 0.2 | 0.217 | 1 |
| ZNF91 | 4.71E-05 | 0.307125086 | 0.4 | 0.458 | 1 |
| RBBP6 | 4.71E-05 | 0.218640186 | 0.3 | 0.357 | 1 |
| KIF1BP | 4.83E-05 | 0.141834592 | 0.1 | 0.12 | 1 |
| SPIRE1 | 4.89E-05 | 0.213771172 | 0.3 | 0.428 | 1 |
| PAK5 | 5.14E-05 | 0.388007212 | 0.1 | 0.089 | 1 |
| NUDT3 | 5.22E-05 | 0.513619541 | 0.2 | 0.12 | 1 |
| ERC2 | 5.24E-05 | 0.531682142 | 0.3 | 0.228 | 1 |
| PABPN1 | 5.26E-05 | 0.177228954 | 0.2 | 0.299 | 1 |
| ACTR2 | 5.26E-05 | 0.272785574 | 0.2 | 0.266 | 1 |
| GTF2A1 | 5.30E-05 | 0.17782147 | 0.1 | 0.189 | 1 |
| UBB | 5.32E-05 | 0.274210397 | 0.2 | 0.312 | 1 |
| THRA | 5.33E-05 | 0.194970023 | 0.2 | 0.297 | 1 |
| WASL | 5.44E-05 | 0.325597404 | 0.2 | 0.225 | 1 |
| RPS11 | 5.48E-05 | 0.141270433 | 0.1 | 0.145 | 1 |
| AC098850.3 | 5.48E-05 | 0.313923162 | 0.3 | 0.437 | 1 |
| AKAP11 | 5.74E-05 | 0.392954165 | 0.3 | 0.338 | 1 |
| NACA | 5.91E-05 | 0.146574348 | 0.1 | 0.163 | 1 |
| YPEL3 | 6.02E-05 | 0.395559153 | 0.3 | 0.324 | 1 |
| GRIA2 | 6.20E-05 | 0.651259772 | 0.6 | 0.598 | 1 |
| HYDIN | 6.60E-05 | 0.160666184 | 0.3 | 0.349 | 1 |
| TOMM70 | 6.72E-05 | 0.170227065 | 0.1 | 0.142 | 1 |
| ZNF428 | 6.87E-05 | 0.150355419 | 0.1 | 0.112 | 1 |
| TIMM23B | 7.01E-05 | 0.180048458 | 0.2 | 0.286 | 1 |
| PGRMC1 | 7.30E-05 | 0.316080905 | 0.2 | 0.239 | 1 |
| NFAT5 | 7.92E-05 | 0.254571173 | 0.4 | 0.556 | 1 |
| CABIN1 | 8.09E-05 | 0.230288371 | 0.2 | 0.213 | 1 |
| BEX4 | 8.24E-05 | 0.374994586 | 0.2 | 0.228 | 1 |
| SDF4 | 8.32E-05 | 0.164777961 | 0.1 | 0.142 | 1 |
| PDLIM5 | 8.40E-05 | 0.147463054 | 0.2 | 0.248 | 1 |
| PFKFB3 | 8.43E-05 | 0.240245196 | 0.2 | 0.283 | 1 |
| GRIN2A | 8.62E-05 | 0.732783491 | 0.4 | 0.358 | 1 |
| LARGE1 | 8.83E-05 | 0.141581807 | 0.3 | 0.42 | 1 |
| VPS29 | 9.59E-05 | 0.153398995 | 0.1 | 0.173 | 1 |
| ABI2 | 0.000113525 | 0.270142407 | 0.3 | 0.414 | 1 |
| CHRM5 | 0.00011855 | -0.200483123 | 0.2 | 0.208 | 1 |
| VPS45 | 0.00011899 | 0.159393354 | 0.2 | 0.281 | 1 |
| DNAJC5 | 0.000119952 | 0.148049538 | 0.2 | 0.205 | 1 |
| ATP2A2 | 0.000120595 | 0.432352771 | 0.3 | 0.399 | 1 |
| NMT1 | 0.000124117 | 0.173084904 | 0.2 | 0.205 | 1 |
| TMEM132C | 0.000124535 | -0.154126415 | 0.2 | 0.221 | 1 |
| GABARAPL2 | 0.000128095 | 0.224879747 | 0.2 | 0.215 | 1 |
| MIR29B2CHG | 0.000134294 | 0.19197316 | 0.2 | 0.256 | 1 |
| MORF4L2 | 0.000143488 | 0.23345746 | 0.2 | 0.31 | 1 |
| PCBP1 | 0.000153688 | 0.203570854 | 0.2 | 0.233 | 1 |
| AC105402.3 | 0.000155375 | 0.548662797 | 0.2 | 0.217 | 1 |
| MEF2D | 0.000159506 | 0.194959547 | 0.2 | 0.208 | 1 |
| RIC8B | 0.000165459 | 0.180728553 | 0.2 | 0.248 | 1 |
| DAP3 | 0.000170744 | 0.143032313 | 0.2 | 0.25 | 1 |
| ZBTB7A | 0.000180461 | 0.231537734 | 0.2 | 0.288 | 1 |
| WBP2 | 0.000182835 | 0.290030629 | 0.3 | 0.335 | 1 |
| CAPNS1 | 0.000182843 | 0.179248248 | 0.1 | 0.155 | 1 |
| PLEKHA5 | 0.000183396 | 0.571639649 | 0.3 | 0.244 | 1 |
| GRIA4 | 0.000188891 | 0.682464523 | 0.4 | 0.338 | 1 |
| BTBD10 | 0.000223483 | 0.179255262 | 0.1 | 0.182 | 1 |
| MED29 | 0.000237704 | 0.139872419 | 0.1 | 0.11 | 1 |
| ARHGEF11 | 0.000241759 | 0.197140934 | 0.2 | 0.236 | 1 |
| PSMD12 | 0.00024311 | 0.183963558 | 0.1 | 0.146 | 1 |
| NPTXR | 0.000252421 | 0.721903756 | 0.2 | 0.129 | 1 |
| SMAP2 | 0.000252669 | 0.200889593 | 0.1 | 0.167 | 1 |
| GAK | 0.000258307 | 0.221726014 | 0.2 | 0.238 | 1 |
| ZC2HC1A | 0.000267268 | 0.1579816 | 0.1 | 0.178 | 1 |
| TBC1D9B | 0.000267456 | 0.183245143 | 0.1 | 0.163 | 1 |
| ARPC2 | 0.000268769 | 0.145765186 | 0.1 | 0.184 | 1 |
| TPT1 | 0.000269278 | 0.172968023 | 0.2 | 0.216 | 1 |
| GLS | 0.000277471 | 0.377352561 | 0.3 | 0.417 | 1 |
| HAGH | 0.000282962 | 0.196107712 | 0.2 | 0.242 | 1 |
| ACO2 | 0.000287876 | 0.144770599 | 0.2 | 0.247 | 1 |
| ACADVL | 0.000294964 | 0.209770887 | 0.1 | 0.19 | 1 |
| LHFPL3 | 0.000299075 | 0.428113745 | 0.2 | 0.119 | 1 |
| PTPN4 | 0.000318353 | 0.29525255 | 0.3 | 0.378 | 1 |
| MAN2C1 | 0.000332303 | 0.145612302 | 0.1 | 0.154 | 1 |
| SPPL2B | 0.000333255 | 0.231322836 | 0.2 | 0.214 | 1 |
| VPS35 | 0.000343036 | 0.164380433 | 0.1 | 0.175 | 1 |
| SERPINI1 | 0.000346636 | 0.353531886 | 0.3 | 0.418 | 1 |
| FAM169A | 0.000349454 | 0.150198538 | 0.2 | 0.227 | 1 |
| ATP6V1C1 | 0.000351158 | 0.233132684 | 0.2 | 0.213 | 1 |
| RNPS1 | 0.000351929 | 0.170159688 | 0.2 | 0.249 | 1 |
| STRAP | 0.000352046 | 0.158799616 | 0.1 | 0.142 | 1 |
| CTSB | 0.000372156 | 0.591495499 | 0.1 | 0.087 | 1 |
| GPRASP1 | 0.000377167 | 0.545921747 | 0.1 | 0.076 | 1 |
| HSPA4 | 0.000381351 | 0.161872497 | 0.1 | 0.193 | 1 |
| RPL15 | 0.000391854 | 0.404625922 | 0.2 | 0.284 | 1 |
| YWHAZ | 0.000391972 | 0.443182503 | 0.3 | 0.417 | 1 |
| CCDC186 | 0.00039965 | 0.158925859 | 0.1 | 0.174 | 1 |
| C16orf72 | 0.000422215 | 0.219232717 | 0.2 | 0.218 | 1 |
| EIF3A | 0.000437055 | 0.205084244 | 0.2 | 0.238 | 1 |
| ULK1 | 0.000444587 | 0.146794134 | 0.1 | 0.148 | 1 |
| ATP6V1E1 | 0.00045315 | 0.245298259 | 0.1 | 0.186 | 1 |
| KIF21A | 0.000453827 | 0.343061163 | 0.3 | 0.403 | 1 |
| INPP4A | 0.000463496 | 0.55169137 | 0.2 | 0.132 | 1 |
| CPEB3 | 0.000464572 | 0.260822129 | 0.4 | 0.452 | 1 |
| GTF3A | 0.000490731 | 0.144872159 | 0.1 | 0.115 | 1 |
| HSPD1 | 0.000492789 | 0.270200348 | 0.2 | 0.23 | 1 |
| SPCS1 | 0.00049303 | 0.147433982 | 0.1 | 0.154 | 1 |
| DENND5B | 0.000493588 | 0.255544809 | 0.4 | 0.436 | 1 |
| ATP6V1B2 | 0.000511049 | 0.780566369 | 0.2 | 0.127 | 1 |
| LGI4 | 0.000515506 | 0.495379178 | 0.1 | 0.075 | 1 |
| COLEC12 | 0.000529848 | 0.417388751 | 0.1 | 0.097 | 1 |
| AHI1 | 0.000554501 | 0.245059186 | 0.5 | 0.624 | 1 |
| GON4L | 0.000628014 | 0.187393752 | 0.2 | 0.246 | 1 |
| CDC37 | 0.000642734 | 0.204843417 | 0.1 | 0.17 | 1 |
| ARPP21 | 0.000655624 | 0.620467218 | 0.3 | 0.317 | 1 |
| UHMK1 | 0.000665231 | 0.197045206 | 0.1 | 0.169 | 1 |
| CLEC16A | 0.000665715 | 0.23032419 | 0.3 | 0.396 | 1 |
| EXTL3 | 0.000690407 | 0.154212727 | 0.1 | 0.161 | 1 |
| KDM1A | 0.00070217 | 0.165734065 | 0.1 | 0.168 | 1 |
| PHF21B | 0.000710863 | 0.405968003 | 0.1 | 0.078 | 1 |
| WDR60 | 0.00071933 | 0.285216303 | 0.3 | 0.316 | 1 |
| PKD1 | 0.000738726 | 0.437155595 | 0.3 | 0.316 | 1 |
| RFX3 | 0.000750884 | 0.256309532 | 0.4 | 0.53 | 1 |
| PPP3CB | 0.000755437 | 0.27626694 | 0.2 | 0.284 | 1 |
| MAP1LC3A | 0.000760018 | 0.22770663 | 0.1 | 0.16 | 1 |
| TSPAN3 | 0.00078842 | 0.2389383 | 0.1 | 0.19 | 1 |
| KIF1A | 0.000789162 | 0.361101389 | 0.3 | 0.354 | 1 |
| RABL6 | 0.000809218 | 0.175095443 | 0.2 | 0.253 | 1 |
| CLCN7 | 0.000817449 | 0.167563069 | 0.1 | 0.189 | 1 |
| CAMSAP2 | 0.000820343 | 0.265542277 | 0.4 | 0.497 | 1 |
| SARAF | 0.000820566 | 0.572566502 | 0.3 | 0.386 | 1 |

| | | | | | |
|---|---|---|---|---|---|
| SCP2 | 0.000821877 | 0.247139836 | 0.1 | 0.177 | 1 |
| MAPK1 | 0.000835647 | 0.508040927 | 0.3 | 0.373 | 1 |
| TAB2 | 0.000848952 | 0.275160873 | 0.3 | 0.311 | 1 |
| BTBD2 | 0.000905315 | 0.160045779 | 0.1 | 0.14 | 1 |
| UBA5 | 0.000914903 | 0.161257957 | 0.1 | 0.143 | 1 |
| APBA2 | 0.000918823 | 0.24355077 | 0.3 | 0.35 | 1 |
| SGCD | 0.000953919 | 0.166198519 | 0.3 | 0.43 | 1 |
| MT-ND2 | 0.00099218 | 0.437807144 | 0.6 | 0.7 | 1 |
| EIF4G1 | 0.001005536 | 0.172260304 | 0.1 | 0.13 | 1 |
| PRPF8 | 0.001038765 | 0.216511064 | 0.2 | 0.226 | 1 |
| PELP1 | 0.001055704 | 0.139410731 | 0.1 | 0.128 | 1 |
| SLC26A3 | 0.001059838 | 0.570306132 | 0.5 | 0.59 | 1 |
| C11orf49 | 0.001073013 | 0.149782265 | 0.3 | 0.336 | 1 |
| DCAF6 | 0.001078436 | 0.272877782 | 0.2 | 0.289 | 1 |
| MDH1 | 0.001083537 | 0.674811732 | 0.2 | 0.156 | 1 |
| DYNC2LI1 | 0.00109239 | 0.137701231 | 0.1 | 0.186 | 1 |
| RPS4X | 0.001099809 | 0.154595487 | 0.1 | 0.136 | 1 |
| RAD23A | 0.001112747 | 0.183599086 | 0.1 | 0.163 | 1 |
| PARK7 | 0.001160099 | 0.163082996 | 0.1 | 0.128 | 1 |
| MTFMT | 0.001171279 | 0.137596532 | 0.1 | 0.104 | 1 |
| PEBP1 | 0.001204334 | 0.564308948 | 0.4 | 0.532 | 1 |
| MAEA | 0.001217969 | 0.165597046 | 0.1 | 0.126 | 1 |
| TMSB10 | 0.001272657 | 0.680462751 | 0.1 | 0.104 | 1 |
| PSD3 | 0.001291554 | 0.220746818 | 0.6 | 0.736 | 1 |
| INO80D | 0.001340679 | 0.44752951 | 0.5 | 0.62 | 1 |
| PCDH10 | 0.001347175 | 0.235390658 | 0.2 | 0.284 | 1 |
| NUAK1 | 0.001381357 | 0.36351913 | 0.2 | 0.306 | 1 |
| DHX30 | 0.001398289 | 0.142002365 | 0.2 | 0.211 | 1 |
| ELOC | 0.001402545 | 0.187286581 | 0.1 | 0.102 | 1 |
| SCAI | 0.001424769 | 0.156087782 | 0.2 | 0.228 | 1 |
| RPL23A | 0.001443997 | 0.201755562 | 0.1 | 0.138 | 1 |
| ATMIN | 0.001466457 | 0.186218636 | 0.1 | 0.115 | 1 |
| SLC35B4 | 0.001475631 | 0.182552264 | 0.1 | 0.156 | 1 |
| ACTB | 0.001499961 | 0.464138781 | 0.6 | 0.728 | 1 |
| CREG2 | 0.00150628 | 0.766276254 | 0.2 | 0.145 | 1 |
| TRIM9 | 0.001581651 | 0.161420158 | 0.5 | 0.581 | 1 |
| FAM168B | 0.001584356 | 0.280080832 | 0.2 | 0.236 | 1 |
| ETV6 | 0.001585289 | 0.355190791 | 0.2 | 0.217 | 1 |
| VDAC3 | 0.00161476 | 0.28783648 | 0.2 | 0.194 | 1 |
| EHMT2 | 0.001635067 | 0.158738918 | 0.1 | 0.172 | 1 |
| TRIR | 0.001637927 | 0.205835412 | 0.1 | 0.138 | 1 |
| KIF3A | 0.00164108 | 0.382937477 | 0.3 | 0.359 | 1 |
| SKP1 | 0.001730741 | 0.472639532 | 0.2 | 0.278 | 1 |
| C18orf25 | 0.001783833 | 0.160435377 | 0.1 | 0.151 | 1 |
| ZFP36L2 | 0.00181139 | 0.216484393 | 0.1 | 0.164 | 1 |
| NSMAF | 0.001877399 | 0.170243677 | 0.2 | 0.222 | 1 |
| C1orf61 | 0.001886215 | 0.523272046 | 0.1 | 0.12 | 1 |
| FAM126B | 0.001887396 | 0.278680842 | 0.2 | 0.279 | 1 |
| H2AFV | 0.001890782 | 0.168353485 | 0.2 | 0.204 | 1 |
| PSMG2 | 0.001917985 | 0.144597142 | 0.2 | 0.191 | 1 |
| YWHAB | 0.002027829 | 0.684785765 | 0.2 | 0.302 | 1 |
| ERO1A | 0.002031738 | 0.167737823 | 0.1 | 0.131 | 1 |
| SOD1 | 0.002114039 | 0.263304009 | 0.2 | 0.202 | 1 |
| SRI | 0.002189975 | 0.184698375 | 0.1 | 0.131 | 1 |
| ACTR1A | 0.002203768 | 0.206019438 | 0.1 | 0.158 | 1 |
| PTPRE | 0.002214859 | 0.168566781 | 0.2 | 0.231 | 1 |
| STRBP | 0.002245462 | 0.470796039 | 0.2 | 0.169 | 1 |
| AF117829.1 | 0.002331445 | 0.188585444 | 0.2 | 0.208 | 1 |
| AL078639.1 | 0.002340852 | 0.427320515 | 0.1 | 0.098 | 1 |
| AZIN1 | 0.002373033 | 0.267913871 | 0.1 | 0.162 | 1 |
| STMN1 | 0.002381319 | 0.640947609 | 0.3 | 0.4 | 1 |
| SCPEP1 | 0.002431015 | 0.175463175 | 0.1 | 0.137 | 1 |
| ASRGL1 | 0.002448321 | 0.157220998 | 0.1 | 0.174 | 1 |
| NME7 | 0.002493827 | 0.148142813 | 0.2 | 0.219 | 1 |
| BRD2 | 0.002668012 | 0.173250567 | 0.2 | 0.207 | 1 |
| PCDH9-AS1 | 0.002668987 | 0.16623848 | 0.1 | 0.112 | 1 |
| MYCBP2 | 0.002716682 | 0.344172722 | 0.5 | 0.648 | 1 |
| G3BP2 | 0.002723573 | 0.335147234 | 0.2 | 0.235 | 1 |
| ERC1 | 0.002728624 | 0.266010284 | 0.4 | 0.458 | 1 |
| SCN1B | 0.002746131 | 0.551502184 | 0.1 | 0.088 | 1 |
| DUT | 0.002757895 | 0.193052258 | 0.2 | 0.191 | 1 |
| DHRSX | 0.002809734 | 0.240199531 | 0.2 | 0.212 | 1 |
| PANK2 | 0.002860568 | 0.148302241 | 0.1 | 0.135 | 1 |
| GOLGA8A | 0.002907 | 0.230506565 | 0.1 | 0.132 | 1 |
| WDR17 | 0.002978584 | 0.425204894 | 0.2 | 0.127 | 1 |
| TMEM246 | 0.003034288 | 0.287666818 | 0.1 | 0.11 | 1 |
| SERINC1 | 0.003036695 | 0.786053475 | 0.3 | 0.436 | 1 |
| HPCAL1 | 0.003054143 | 0.527772474 | 0.1 | 0.091 | 1 |
| ERH | 0.003055787 | 0.23076037 | 0.1 | 0.145 | 1 |
| CCT2 | 0.003094138 | 0.185620933 | 0.1 | 0.109 | 1 |
| PSMD3 | 0.003117241 | 0.183451567 | 0.1 | 0.131 | 1 |
| SLC25A37 | 0.003165122 | 0.225063762 | 0.2 | 0.205 | 1 |
| SIN3B | 0.003296043 | 0.277026327 | 0.1 | 0.184 | 1 |
| RHBDD2 | 0.003444296 | 0.479493695 | 0.1 | 0.093 | 1 |
| MAGED1 | 0.003478659 | 0.145930687 | 0.2 | 0.217 | 1 |
| DHFR | 0.003486364 | 0.409766996 | 0.2 | 0.204 | 1 |
| KDM7A | 0.003546874 | 0.190030976 | 0.1 | 0.133 | 1 |
| AGAP3 | 0.003606558 | 0.283649115 | 0.2 | 0.283 | 1 |
| ZFPM2 | 0.003615116 | 0.302407325 | 0.2 | 0.206 | 1 |
| PDE4DIP | 0.003621102 | 0.599970978 | 0.5 | 0.553 | 1 |
| SMARCA2 | 0.003657669 | 0.293977752 | 0.3 | 0.415 | 1 |
| LRRC4C | 0.003664658 | 0.594642481 | 0.5 | 0.549 | 1 |
| NISCH | 0.003705739 | 0.254716916 | 0.1 | 0.188 | 1 |
| PLEKHB2 | 0.003759476 | 0.289452496 | 0.1 | 0.175 | 1 |
| ACSS1 | 0.003801552 | 0.225975006 | 0.2 | 0.235 | 1 |
| EXTL2 | 0.003843881 | 0.251622297 | 0.1 | 0.112 | 1 |
| RIC3 | 0.003857189 | 0.163305132 | 0.1 | 0.142 | 1 |
| ARNT2 | 0.003955709 | 0.288538555 | 0.4 | 0.448 | 1 |
| BMPR2 | 0.003963784 | 0.578657751 | 0.3 | 0.248 | 1 |
| SFMBT2 | 0.003970754 | 0.291237307 | 0.3 | 0.329 | 1 |
| BAALC | 0.003972536 | 0.622898279 | 0.2 | 0.199 | 1 |
| PLD3 | 0.003995917 | 0.432455557 | 0.3 | 0.39 | 1 |
| TARBP1 | 0.004006083 | 0.348105523 | 0.2 | 0.269 | 1 |
| GNA12 | 0.004020717 | 0.278126703 | 0.2 | 0.208 | 1 |
| ELOB | 0.004044718 | 0.144932049 | 0.1 | 0.116 | 1 |
| PJA2 | 0.004051221 | 0.516030597 | 0.3 | 0.419 | 1 |
| FASN | 0.004084174 | 0.196388913 | 0.1 | 0.128 | 1 |
| MAP2K2 | 0.004120405 | 0.183335915 | 0.1 | 0.185 | 1 |
| PGP | 0.004196995 | 0.229606482 | 0.1 | 0.172 | 1 |
| TOMM20 | 0.004256681 | 0.458291972 | 0.2 | 0.225 | 1 |
| ZYG11B | 0.004259809 | 0.362102314 | 0.2 | 0.262 | 1 |
| RPS24 | 0.004305586 | 0.175744751 | 0.1 | 0.171 | 1 |
| CCDC6 | 0.004409628 | 0.145140066 | 0.1 | 0.173 | 1 |
| TLN2 | 0.004434972 | 0.221207686 | 0.3 | 0.302 | 1 |
| GPX4 | 0.004451359 | 0.301852925 | 0.1 | 0.151 | 1 |
| SSB | 0.004533644 | 0.162282173 | 0.1 | 0.136 | 1 |
| DPY19L1 | 0.004608304 | 0.142455547 | 0.1 | 0.176 | 1 |
| BCL7A | 0.004625845 | 0.182108666 | 0.1 | 0.136 | 1 |
| HACD3 | 0.004640336 | 0.191209145 | 0.2 | 0.242 | 1 |
| ARHGAP32 | 0.004730134 | 0.639800438 | 0.4 | 0.408 | 1 |
| CHPT1 | 0.004844051 | 0.520024202 | 0.2 | 0.17 | 1 |
| TNIK | 0.004956647 | 0.554105273 | 0.5 | 0.51 | 1 |
| NECAP1 | 0.004962664 | 0.209593581 | 0.1 | 0.109 | 1 |
| SLC44A2 | 0.00499029 | 0.164631927 | 0.1 | 0.144 | 1 |
| GPT2 | 0.005210755 | 0.192254766 | 0.1 | 0.158 | 1 |
| AC007388.1 | 0.00523643 | 0.459811211 | 0.1 | 0.105 | 1 |
| IDH3A | 0.005467467 | 0.168510973 | 0.1 | 0.118 | 1 |
| ACTR3B | 0.005607152 | 0.147241925 | 0.1 | 0.149 | 1 |
| PREPL | 0.005710949 | 0.683166006 | 0.4 | 0.484 | 1 |
| RNF167 | 0.005723998 | 0.151615596 | 0.1 | 0.11 | 1 |
| TCAF1 | 0.005781859 | 0.3635589 | 0.2 | 0.27 | 1 |
| RAB31 | 0.006060564 | 0.397619009 | 0.1 | 0.081 | 1 |
| RLIM | 0.006086882 | 0.175093839 | 0.1 | 0.1 | 1 |
| RASSF8 | 0.006295418 | 0.147562383 | 0.2 | 0.22 | 1 |
| SMIM13 | 0.006323252 | 0.220140804 | 0.1 | 0.123 | 1 |
| COX6B1 | 0.00643327 | 0.159650685 | 0.1 | 0.106 | 1 |
| FAR2 | 0.006482262 | 0.229329722 | 0.1 | 0.083 | 1 |
| SEMA4B | 0.006510786 | 0.215166613 | 0.1 | 0.153 | 1 |

| | | | | | |
|---|---|---|---|---|---|
| SPIDR | 0.006696048 | 0.210927905 | 0.4 | 0.453 | 1 |
| B4GAT1 | 0.006727012 | 0.595501059 | 0.1 | 0.082 | 1 |
| FKBP4 | 0.00685659 | 0.265630496 | 0.1 | 0.129 | 1 |
| STUB1 | 0.007148553 | 0.142850054 | 0.1 | 0.111 | 1 |
| MADD | 0.007162942 | 0.240907695 | 0.1 | 0.162 | 1 |
| DYNC2H1 | 0.007185464 | 0.264459165 | 0.2 | 0.281 | 1 |
| SH3BGRL | 0.007291523 | 0.141215187 | 0.2 | 0.187 | 1 |
| TUBB | 0.007492687 | 0.649454491 | 0.1 | 0.094 | 1 |
| RGS11 | 0.007709691 | 0.217219947 | 0.1 | 0.105 | 1 |
| UPF1 | 0.007744698 | 0.193173001 | 0.1 | 0.12 | 1 |
| PRKCI | 0.007842507 | 0.162482251 | 0.2 | 0.184 | 1 |
| ADGRL3 | 0.007906007 | 0.317087463 | 0.7 | 0.768 | 1 |
| SOX9 | 0.008270319 | 0.419656019 | 0.1 | 0.083 | 1 |
| ATP6AP2 | 0.008386591 | 0.364274182 | 0.2 | 0.188 | 1 |
| GRINA | 0.008476301 | 0.479564751 | 0.1 | 0.09 | 1 |
| CIZ1 | 0.008503326 | 0.183721727 | 0.1 | 0.135 | 1 |
| EIF1B | 0.008575459 | 0.223318813 | 0.1 | 0.133 | 1 |
| FADS3 | 0.008666949 | 0.204714215 | 0.1 | 0.142 | 1 |
| SATB2 | 0.008748773 | 0.507500781 | 0.2 | 0.129 | 1 |
| TECPR1 | 0.008843677 | 0.14559866 | 0.1 | 0.107 | 1 |
| FKBP8 | 0.008865406 | 0.395956282 | 0.2 | 0.283 | 1 |
| NUDT21 | 0.009055334 | 0.254684144 | 0.1 | 0.173 | 1 |
| ITGB4 | 0.009094615 | 0.449161157 | 0.1 | 0.096 | 1 |
| NAV2 | 0.009178848 | 0.392695677 | 0.4 | 0.405 | 1 |
| GPD2 | 0.009267536 | 0.139722404 | 0.2 | 0.2 | 1 |
| NNT | 0.009483641 | 0.193542334 | 0.2 | 0.247 | 1 |
| ARMC2 | 0.009534084 | 0.363471755 | 0.1 | 0.089 | 1 |
| HSPA1B | 0.009803631 | 0.213781968 | 0.1 | 0.116 | 1 |
| SSU72 | 0.009813314 | 0.236508639 | 0.1 | 0.122 | 1 |
| PPM1L | 0.010047022 | 0.423876927 | 0.2 | 0.201 | 1 |
| CCNDBP1 | 0.010157969 | 0.182407006 | 0.1 | 0.126 | 1 |
| FAM20C | 0.010655662 | 0.337336174 | 0.1 | 0.155 | 1 |
| STK25 | 0.010844399 | 0.164882595 | 0.1 | 0.179 | 1 |
| CREM | 0.011515185 | 0.142163092 | 0.1 | 0.167 | 1 |
| NBEA | 0.011780279 | 0.618330211 | 0.5 | 0.539 | 1 |
| REV3L | 0.012153702 | 0.362396675 | 0.4 | 0.479 | 1 |
| ATL2 | 0.012289735 | 0.178502285 | 0.1 | 0.134 | 1 |
| TTC33 | 0.012357374 | 0.181124815 | 0.1 | 0.175 | 1 |
| SLC25A4 | 0.012879285 | 0.220260348 | 0.1 | 0.1 | 1 |
| COPG1 | 0.012947682 | 0.165256571 | 0.1 | 0.105 | 1 |
| UBC | 0.013143652 | 0.663050511 | 0.3 | 0.358 | 1 |
| SRP14 | 0.013290652 | 0.393805594 | 0.2 | 0.211 | 1 |
| HPRT1 | 0.013353043 | 0.235249996 | 0.2 | 0.182 | 1 |
| OCRL | 0.013415832 | 0.1540114 | 0.1 | 0.112 | 1 |
| NQO2 | 0.013474104 | 0.174401965 | 0.1 | 0.118 | 1 |
| WDR6 | 0.013606226 | 0.189240131 | 0.1 | 0.14 | 1 |
| CCPG1 | 0.013740118 | 0.315004542 | 0.3 | 0.304 | 1 |
| GPR137B | 0.01384521 | 0.151132501 | 0.1 | 0.113 | 1 |
| PER1 | 0.013982811 | 0.149208759 | 0.1 | 0.179 | 1 |
| CACNA1E | 0.014228897 | 0.396535432 | 0.1 | 0.104 | 1 |
| ATP6AP1 | 0.014969927 | 0.53614851 | 0.1 | 0.184 | 1 |
| MT3 | 0.015998993 | 0.405781945 | 0.2 | 0.271 | 1 |
| SLC25A3 | 0.016212527 | 0.379796396 | 0.1 | 0.163 | 1 |
| SLC38A6 | 0.016222866 | 0.160420574 | 0.1 | 0.153 | 1 |
| PCNX2 | 0.016256948 | 0.589329396 | 0.2 | 0.193 | 1 |
| CHCHD2 | 0.016366852 | 0.271233893 | 0.1 | 0.163 | 1 |
| DCTN5 | 0.01652273 | 0.250806271 | 0.1 | 0.15 | 1 |
| SUB1 | 0.016937067 | 0.248109598 | 0.2 | 0.186 | 1 |
| REPS1 | 0.017051521 | 0.289683581 | 0.3 | 0.317 | 1 |
| AC096711.2 | 0.017596303 | 0.190691668 | 0.2 | 0.213 | 1 |
| RPS20 | 0.017668894 | 0.159864075 | 0.1 | 0.102 | 1 |
| ARF1 | 0.017676379 | 0.432819323 | 0.2 | 0.267 | 1 |
| TCP1 | 0.017778423 | 0.163608065 | 0.1 | 0.125 | 1 |
| GAN | 0.017936637 | 0.21566404 | 0.2 | 0.186 | 1 |
| STAU2 | 0.018029543 | 0.316553475 | 0.3 | 0.297 | 1 |
| NOP56 | 0.018224341 | 0.140964737 | 0.1 | 0.14 | 1 |
| UBAC2 | 0.018921558 | 0.25391752 | 0.2 | 0.285 | 1 |
| PSAP | 0.018933872 | 0.498003239 | 0.5 | 0.594 | 1 |
| RAC1 | 0.01895998 | 0.421042089 | 0.3 | 0.318 | 1 |
| ATP6V1A | 0.019990062 | 0.316772863 | 0.2 | 0.239 | 1 |
| ZNF483 | 0.020042721 | 0.475981262 | 0.3 | 0.398 | 1 |
| P4HB | 0.020065551 | 0.264888642 | 0.1 | 0.165 | 1 |
| KIF3C | 0.020223341 | 0.566877843 | 0.1 | 0.119 | 1 |
| TTN-AS1 | 0.020460446 | 0.183790983 | 0.2 | 0.242 | 1 |
| SCMH1 | 0.020648987 | 0.173091745 | 0.2 | 0.271 | 1 |
| SRSF8 | 0.020681328 | 0.209274838 | 0.1 | 0.106 | 1 |
| C16orf45 | 0.02100022 | 0.266464123 | 0.2 | 0.297 | 1 |
| PRPSAP1 | 0.021045925 | 0.142254548 | 0.1 | 0.156 | 1 |
| YWHAG | 0.021442045 | 0.991959816 | 0.3 | 0.256 | 1 |
| FEM1B | 0.021902761 | 0.218799776 | 0.1 | 0.172 | 1 |
| ATXN7L3B | 0.02259175 | 0.248126546 | 0.1 | 0.152 | 1 |
| WASF3 | 0.022988235 | 0.24486589 | 0.1 | 0.18 | 1 |
| MAP1A | 0.023038847 | 0.453928461 | 0.3 | 0.336 | 1 |
| CALM1 | 0.023480924 | 1.215847247 | 0.5 | 0.607 | 1 |
| PPP2R2C | 0.023921346 | 0.359199689 | 0.2 | 0.24 | 1 |
| TEF | 0.023937097 | 0.144477011 | 0.1 | 0.147 | 1 |
| CNN3 | 0.025351904 | 0.277498081 | 0.1 | 0.165 | 1 |
| IRAK1BP1 | 0.025651413 | 0.146399657 | 0.1 | 0.155 | 1 |
| ERGIC3 | 0.025790488 | 0.235181186 | 0.1 | 0.123 | 1 |
| RGS12 | 0.026575035 | 0.202021906 | 0.2 | 0.201 | 1 |
| MT2A | 0.026797753 | 0.205845013 | 0.1 | 0.158 | 1 |
| CIC | 0.027927891 | 0.138374546 | 0.1 | 0.117 | 1 |
| DPY19L3 | 0.028437563 | 0.142410694 | 0.1 | 0.167 | 1 |
| AC027117.2 | 0.029034051 | 0.264056174 | 0.1 | 0.106 | 1 |
| ZNF106 | 0.029373846 | 0.255694252 | 0.2 | 0.202 | 1 |
| RETREG2 | 0.029449841 | 0.329619506 | 0.1 | 0.123 | 1 |
| GPAA1 | 0.030484872 | 0.146703904 | 0.1 | 0.112 | 1 |
| CBX5 | 0.030490298 | 0.287287167 | 0.2 | 0.218 | 1 |
| PEA15 | 0.030621665 | 0.414109476 | 0.2 | 0.212 | 1 |
| SCG5 | 0.030818369 | 0.482835541 | 0.2 | 0.213 | 1 |
| DEF8 | 0.032035487 | 0.147837953 | 0.1 | 0.116 | 1 |
| CDKL5 | 0.032317718 | 0.278578079 | 0.3 | 0.316 | 1 |
| RAP2A | 0.032364343 | 0.318707098 | 0.1 | 0.168 | 1 |
| LPIN2 | 0.03252484 | 0.151597285 | 0.1 | 0.141 | 1 |
| LMBRD2 | 0.033105556 | 0.284130862 | 0.2 | 0.192 | 1 |
| DEAF1 | 0.033388585 | 0.182610593 | 0.1 | 0.144 | 1 |
| PRNP | 0.033969363 | 0.807601172 | 0.4 | 0.518 | 1 |
| PARD3B | 0.034387173 | 0.497813529 | 0.3 | 0.26 | 1 |
| MRFAP1 | 0.034752438 | 0.289615585 | 0.1 | 0.135 | 1 |
| ARHGEF9 | 0.034876222 | 0.293282512 | 0.2 | 0.187 | 1 |
| NSF | 0.035746821 | 0.739353929 | 0.3 | 0.326 | 1 |
| SUFU | 0.035752508 | 0.13920577 | 0.1 | 0.142 | 1 |
| ABAT | 0.036655552 | 0.207403818 | 0.1 | 0.162 | 1 |
| SYNGR1 | 0.037259174 | 0.488498504 | 0.1 | 0.113 | 1 |
| ARFGAP1 | 0.037514861 | 0.201369903 | 0.1 | 0.167 | 1 |
| PITHD1 | 0.037765628 | 0.204608961 | 0.1 | 0.125 | 1 |
| HIVEP1 | 0.037887268 | 0.204887001 | 0.2 | 0.197 | 1 |
| SIRT5 | 0.038077458 | 0.159431431 | 0.1 | 0.129 | 1 |
| TTTY14 | 0.038124161 | 0.421202074 | 0.2 | 0.174 | 1 |
| NAPA | 0.038976614 | 0.373010143 | 0.1 | 0.179 | 1 |
| SNRK | 0.038991258 | 0.168890384 | 0.1 | 0.116 | 1 |
| ACLY | 0.039506727 | 0.185719609 | 0.1 | 0.105 | 1 |
| TSHZ1 | 0.04055594 | 0.33555962 | 0.1 | 0.092 | 1 |
| ARPP19 | 0.041221206 | 0.45366754 | 0.2 | 0.281 | 1 |
| BCAP29 | 0.042087957 | 0.277898719 | 0.2 | 0.185 | 1 |
| ZFYVE28 | 0.042365756 | 0.555242566 | 0.2 | 0.164 | 1 |
| UBA6-AS1 | 0.042450207 | 0.141984255 | 0.3 | 0.324 | 1 |
| SCOC | 0.042867733 | 0.258919566 | 0.1 | 0.15 | 1 |
| LHFPL6 | 0.043756345 | -0.243717781 | 0.2 | 0.25 | 1 |
| INPP5F | 0.043917101 | 0.594966942 | 0.2 | 0.189 | 1 |
| PNPLA6 | 0.044071753 | 0.22888222 | 0.1 | 0.162 | 1 |
| DNAJA1 | 0.044129489 | 0.29583636 | 0.2 | 0.196 | 1 |
| FAM19A5 | 0.044161457 | 0.215991684 | 0.3 | 0.317 | 1 |
| PGAM1 | 0.044455409 | 0.687875455 | 0.1 | 0.126 | 1 |
| FBXO34 | 0.044804779 | 0.222181538 | 0.1 | 0.164 | 1 |
| AC098829.1 | 0.044864468 | 0.28442417 | 0.1 | 0.144 | 1 |
| MAPK3 | 0.04593138 | 0.162000717 | 0.1 | 0.115 | 1 |
| KIF9-AS1 | 0.046411689 | 0.25144481 | 0.2 | 0.198 | 1 |

| | | | | | |
|---|---|---|---|---|---|
| PRKCB | 0.04700985 | 0.641752229 | 0.3 | 0.311 | 1 |
| GTF3C1 | 0.049003288 | 0.28495576 | 0.2 | 0.192 | 1 |
| DGKE | 0.049010506 | 0.289768404 | 0.1 | 0.111 | 1 |
| RPS6KA3 | 0.049241018 | 0.175248832 | 0.1 | 0.158 | 1 |
| SUGP2 | 0.049422786 | 0.321379083 | 0.3 | 0.304 | 1 |
| FAM217B | 0.050283455 | 0.377011078 | 0.2 | 0.193 | 1 |
| KATNAL2 | 0.050921806 | 0.159583769 | 0.1 | 0.153 | 1 |
| RAPGEF1 | 0.051964696 | 0.145294012 | 0.2 | 0.192 | 1 |
| SLITRK5 | 0.052260258 | 0.368750825 | 0.1 | 0.085 | 1 |
| CATSPER2 | 0.052857091 | 0.220530361 | 0.2 | 0.194 | 1 |
| RELN | 0.054010499 | 0.371277484 | 0.1 | 0.122 | 1 |
| WSB2 | 0.054645644 | 0.267956879 | 0.1 | 0.116 | 1 |
| CYFIP1 | 0.055529314 | 0.193472477 | 0.1 | 0.102 | 1 |
| PFDN5 | 0.055584938 | 0.200618115 | 0.1 | 0.125 | 1 |
| NADK2 | 0.055954123 | 0.281589485 | 0.1 | 0.14 | 1 |
| CRELD1 | 0.056916107 | 0.191271466 | 0.1 | 0.139 | 1 |
| INTU | 0.059403537 | 0.534586744 | 0.2 | 0.154 | 1 |
| H1FX | 0.059765755 | 0.222873314 | 0.1 | 0.14 | 1 |
| MLF2 | 0.061580656 | 0.393872306 | 0.1 | 0.142 | 1 |
| LETM1 | 0.06247367 | 0.167780001 | 0.1 | 0.107 | 1 |
| TMEM170B | 0.062788387 | 0.146309269 | 0.1 | 0.1 | 1 |
| C14orf132 | 0.062958095 | 0.241156878 | 0.1 | 0.172 | 1 |
| CHCHD6 | 0.063644212 | 0.209243952 | 0.1 | 0.141 | 1 |
| SYT14 | 0.063706144 | 0.196092127 | 0.2 | 0.245 | 1 |
| AES | 0.063992491 | 0.442432344 | 0.2 | 0.242 | 1 |
| TOLLIP | 0.064060403 | 0.598432124 | 0.1 | 0.112 | 1 |
| CYB5B | 0.065054342 | 0.265038518 | 0.2 | 0.177 | 1 |
| CCDC3 | 0.065848467 | 0.212905649 | 0.1 | 0.099 | 1 |
| ATP2B4 | 0.06597015 | 0.293434314 | 0.3 | 0.303 | 1 |
| PPM1K | 0.068111853 | 0.430835499 | 0.3 | 0.351 | 1 |
| WIPF2 | 0.069560898 | 0.239719184 | 0.2 | 0.26 | 1 |
| OSBPL11 | 0.071433466 | 0.141460846 | 0.1 | 0.117 | 1 |
| KLC1 | 0.07200631 | 0.6131461 | 0.3 | 0.371 | 1 |
| DLGAP4 | 0.072743799 | 0.305420394 | 0.1 | 0.108 | 1 |
| EVL | 0.073508837 | 0.382495432 | 0.2 | 0.284 | 1 |
| CBX6 | 0.075202905 | 0.355697537 | 0.2 | 0.2 | 1 |
| FAM199X | 0.075329578 | 0.234034739 | 0.1 | 0.115 | 1 |
| SIRPA | 0.075508198 | 0.609863778 | 0.2 | 0.235 | 1 |
| YPEL5 | 0.075647049 | 0.187442445 | 0.1 | 0.115 | 1 |
| LONP1 | 0.07654636 | 0.150387704 | 0.1 | 0.122 | 1 |
| GDI1 | 0.076549457 | 0.544329298 | 0.3 | 0.308 | 1 |
| HSPA8 | 0.077076977 | 0.424019696 | 0.2 | 0.179 | 1 |
| MSI2 | 0.077721689 | 0.468760633 | 0.4 | 0.505 | 1 |
| UBA52 | 0.079426049 | 0.234878895 | 0.1 | 0.122 | 1 |
| AP1M1 | 0.080754781 | 0.189349535 | 0.1 | 0.11 | 1 |
| TMA7 | 0.080808232 | 0.21111741 | 0.1 | 0.11 | 1 |
| PACRG | 0.081333765 | 0.369329569 | 0.1 | 0.125 | 1 |
| CAMTA1 | 0.081582311 | 0.345068097 | 0.6 | 0.677 | 1 |
| TSPAN14 | 0.084032203 | 0.170088366 | 0.1 | 0.109 | 1 |
| ENSA | 0.085103438 | 0.491122014 | 0.2 | 0.208 | 1 |
| PTPN9 | 0.087162621 | 0.435158428 | 0.1 | 0.123 | 1 |
| RTL8C | 0.088527442 | 0.36724496 | 0.1 | 0.151 | 1 |
| GRID2 | 0.088684838 | 0.213677021 | 0.4 | 0.38 | 1 |
| NORAD | 0.088976319 | 0.807597734 | 0.3 | 0.249 | 1 |
| TLK1 | 0.089535325 | 0.392068164 | 0.2 | 0.227 | 1 |
| USP9Y | 0.090498743 | 0.350361669 | 0.1 | 0.166 | 1 |
| SRGAP3 | 0.091297346 | 0.326454785 | 0.3 | 0.346 | 1 |
| MCRIP1 | 0.091773055 | 0.261302617 | 0.1 | 0.118 | 1 |
| GLCCI1 | 0.092724929 | 0.384534856 | 0.2 | 0.228 | 1 |
| LMTK3 | 0.09398349 | 0.317799729 | 0.1 | 0.1 | 1 |
| ATP1A1 | 0.094788102 | 0.713692591 | 0.3 | 0.309 | 1 |
| BNIP3 | 0.096624645 | 0.34061139 | 0.1 | 0.156 | 1 |
| SERGEF | 0.097074908 | 0.263036095 | 0.2 | 0.204 | 1 |
| CRY2 | 0.100523925 | 0.175108771 | 0.1 | 0.138 | 1 |
| NAPB | 0.102803448 | 0.685216601 | 0.2 | 0.159 | 1 |
| HSPH1 | 0.103918837 | 0.625709862 | 0.2 | 0.185 | 1 |
| JAK1 | 0.104068139 | 0.244822361 | 0.2 | 0.213 | 1 |
| SLC25A12 | 0.105529368 | 0.464611424 | 0.2 | 0.202 | 1 |
| UBE2V2 | 0.105991694 | 0.248895911 | 0.1 | 0.137 | 1 |
| EIF2AK2 | 0.106103168 | 0.190831596 | 0.1 | 0.165 | 1 |
| PPIA | 0.106695974 | 0.567701462 | 0.2 | 0.262 | 1 |
| FP671120.1 | 0.106783426 | 0.771696159 | 0.4 | 0.392 | 1 |
| ATP6V1H | 0.107776392 | 0.35804783 | 0.2 | 0.241 | 1 |
| CLIP3 | 0.108247877 | 0.463777755 | 0.1 | 0.113 | 1 |
| LPIN1 | 0.109091893 | 0.314675505 | 0.2 | 0.24 | 1 |
| RAD9A | 0.111247974 | 0.329986835 | 0.1 | 0.109 | 1 |
| RRAGA | 0.112287504 | 0.184130765 | 0.1 | 0.104 | 1 |
| LDHB | 0.11270276 | 0.357087773 | 0.1 | 0.168 | 1 |
| RNF24 | 0.114289972 | 0.215218684 | 0.1 | 0.152 | 1 |
| ATP6V0D1 | 0.115937953 | 0.280469107 | 0.1 | 0.156 | 1 |
| GABARAPL1 | 0.116175204 | 0.274509313 | 0.1 | 0.159 | 1 |
| PPIP5K1 | 0.121755838 | 0.185458387 | 0.1 | 0.101 | 1 |
| COX7A2L | 0.121797961 | 0.298437059 | 0.1 | 0.12 | 1 |
| CALN1 | 0.12311197 | 0.319666203 | 0.2 | 0.177 | 1 |
| MAP2K1 | 0.123161162 | 0.192033252 | 0.1 | 0.151 | 1 |
| COX6C | 0.125675437 | 0.196000668 | 0.1 | 0.1 | 1 |
| ATP13A2 | 0.125980327 | 0.507065121 | 0.1 | 0.124 | 1 |
| FLOT1 | 0.126393725 | 0.332097492 | 0.1 | 0.143 | 1 |
| BRSK2 | 0.126800191 | 0.19952982 | 0.1 | 0.145 | 1 |
| WDR82 | 0.128823705 | 0.292340988 | 0.1 | 0.169 | 1 |
| ITSN1 | 0.130379639 | 0.584552433 | 0.3 | 0.323 | 1 |
| P4HTM | 0.130680636 | 0.366207156 | 0.1 | 0.135 | 1 |
| AP2M1 | 0.13182215 | 0.457326281 | 0.2 | 0.199 | 1 |
| DDX24 | 0.132158834 | 0.516723097 | 0.2 | 0.274 | 1 |
| SCAF1 | 0.133891375 | 0.159192851 | 0.1 | 0.108 | 1 |
| OAT | 0.134725131 | 0.348239666 | 0.2 | 0.187 | 1 |
| OPTN | 0.135852134 | 0.287312582 | 0.1 | 0.136 | 1 |
| FBXW5 | 0.13868908 | 0.157535073 | 0.1 | 0.118 | 1 |
| ERBB4 | 0.139637368 | 0.181939798 | 0.7 | 0.837 | 1 |
| PNKD | 0.13989695 | 0.32376404 | 0.1 | 0.132 | 1 |
| ANKRD46 | 0.144029885 | 0.156858642 | 0.1 | 0.121 | 1 |
| PRKCA | 0.147053462 | 0.457578233 | 0.5 | 0.508 | 1 |
| DCXR | 0.149768586 | 0.215893826 | 0.1 | 0.105 | 1 |
| DANT2 | 0.151318886 | 0.465706696 | 0.4 | 0.382 | 1 |
| ABHD2 | 0.152197286 | 0.308991153 | 0.3 | 0.318 | 1 |
| PQLC1 | 0.153024589 | 0.293792421 | 0.1 | 0.15 | 1 |
| MICALL2 | 0.15452755 | 0.375289301 | 0.1 | 0.104 | 1 |
| SLC38A1 | 0.155891944 | 0.314423894 | 0.2 | 0.244 | 1 |
| AC092683.1 | 0.156871468 | 0.423779556 | 0.2 | 0.2 | 1 |
| HIF1A | 0.159856636 | 0.314141718 | 0.2 | 0.183 | 1 |
| CDR2 | 0.163360757 | 0.273286592 | 0.1 | 0.11 | 1 |
| GOLGB1 | 0.164086968 | 0.381001391 | 0.4 | 0.414 | 1 |
| MALT1 | 0.164095568 | 0.213360805 | 0.1 | 0.107 | 1 |
| DGKI | 0.164506696 | 0.355419183 | 0.3 | 0.319 | 1 |
| TMX4 | 0.16501125 | 0.46961126 | 0.2 | 0.174 | 1 |
| CPT1C | 0.167502667 | 0.207539077 | 0.1 | 0.101 | 1 |
| GOLIM4 | 0.168663235 | 0.335135365 | 0.1 | 0.16 | 1 |
| GAPDH | 0.171935105 | 0.821174255 | 0.4 | 0.459 | 1 |
| GPR75-ASB3 | 0.172971535 | 0.143867231 | 0.2 | 0.227 | 1 |
| BCL6 | 0.173703343 | 0.469351006 | 0.3 | 0.304 | 1 |
| SMAD1 | 0.173741337 | 0.402318438 | 0.1 | 0.108 | 1 |
| FAM3C | 0.178289136 | 0.412480324 | 0.1 | 0.12 | 1 |
| P4HA1 | 0.179480813 | 0.205992448 | 0.2 | 0.196 | 1 |
| TPM1 | 0.17981701 | 0.145498235 | 0.1 | 0.101 | 1 |
| LINC00461 | 0.180169101 | 0.344633352 | 0.3 | 0.34 | 1 |
| PPP1R13B | 0.183288422 | 0.358966543 | 0.1 | 0.139 | 1 |
| PACS1 | 0.184229982 | 0.458910942 | 0.2 | 0.215 | 1 |
| BMP2K | 0.184356762 | 0.374454763 | 0.2 | 0.259 | 1 |
| RAPH1 | 0.184512471 | 0.240529836 | 0.1 | 0.122 | 1 |
| EML5 | 0.185601852 | 0.198356589 | 0.2 | 0.209 | 1 |
| BCR | 0.186496055 | 0.363340974 | 0.1 | 0.107 | 1 |
| ADGRL1 | 0.188183202 | 0.315043553 | 0.2 | 0.181 | 1 |
| NEDD4L | 0.189559508 | 0.409495227 | 0.2 | 0.198 | 1 |
| CTBP1 | 0.192019438 | 0.273232793 | 0.2 | 0.171 | 1 |
| COL19A1 | 0.192829158 | 0.221984035 | 0.1 | 0.094 | 1 |
| AP1B1 | 0.193442394 | 0.273260281 | 0.1 | 0.104 | 1 |
| CCDC30 | 0.193693423 | 0.151852987 | 0.2 | 0.167 | 1 |
| INTS1 | 0.195209378 | 0.180816247 | 0.1 | 0.126 | 1 |

| | | | | | |
|---|---|---|---|---|---|
| CAPRIN2 | 0.195453296 | 0.197953731 | 0.1 | 0.145 | 1 |
| ICA1L | 0.196296725 | 0.281519465 | 0.2 | 0.174 | 1 |
| CCDC136 | 0.19768995 | 0.235420504 | 0.1 | 0.101 | 1 |
| ATP5IF1 | 0.201447908 | 0.246218649 | 0.1 | 0.114 | 1 |
| NTM | 0.206007119 | 0.213536174 | 0.7 | 0.809 | 1 |
| DMXL2 | 0.206898923 | 0.418035606 | 0.2 | 0.209 | 1 |
| OTUB1 | 0.206927116 | 0.225131299 | 0.1 | 0.111 | 1 |
| AC104806.2 | 0.207227977 | 0.160351077 | 0.1 | 0.119 | 1 |
| MAN1C1 | 0.208550223 | 0.492132436 | 0.1 | 0.114 | 1 |
| ARHGEF3 | 0.211210913 | 0.345397231 | 0.2 | 0.174 | 1 |
| SHC2 | 0.213175154 | 0.194373193 | 0.1 | 0.128 | 1 |
| MAP1LC3B | 0.216476138 | 0.355405208 | 0.1 | 0.16 | 1 |
| PKIG | 0.221758678 | 0.154762736 | 0.1 | 0.132 | 1 |
| GOT1 | 0.224283993 | 0.384550796 | 0.1 | 0.145 | 1 |
| 3-Mar | 0.226123486 | 0.175686798 | 0.1 | 0.114 | 1 |
| GRIPAP1 | 0.229992487 | 0.218689069 | 0.1 | 0.141 | 1 |
| RAP1GAP | 0.231053631 | 0.522054361 | 0.1 | 0.119 | 1 |
| HIVEP3 | 0.231671991 | 0.15027785 | 0.2 | 0.246 | 1 |
| SMIM8 | 0.236204646 | 0.146421985 | 0.1 | 0.1 | 1 |
| DYNLL1 | 0.236863383 | 0.697143258 | 0.3 | 0.314 | 1 |
| PYGB | 0.237092928 | 0.314659394 | 0.1 | 0.139 | 1 |
| MTRNR2L12 | 0.243377638 | 0.305873303 | 0.1 | 0.13 | 1 |
| TRAK1 | 0.245367225 | 0.457080175 | 0.2 | 0.172 | 1 |
| DGKZ | 0.250188159 | 0.386796104 | 0.2 | 0.144 | 1 |
| PRKCE | 0.254844871 | 0.411465866 | 0.4 | 0.412 | 1 |
| FGD4 | 0.259519093 | 0.231770692 | 0.2 | 0.245 | 1 |
| PIN1 | 0.260797854 | 0.303221802 | 0.1 | 0.107 | 1 |
| HECTD2 | 0.261831939 | 0.344494469 | 0.1 | 0.104 | 1 |
| RNF11 | 0.262143643 | 0.481350955 | 0.2 | 0.179 | 1 |
| NUCB1 | 0.265554144 | 0.288132716 | 0.1 | 0.109 | 1 |
| RUNX1T1 | 0.265985578 | 0.14078823 | 0.2 | 0.173 | 1 |
| CCDC57 | 0.266535372 | 0.239534992 | 0.1 | 0.102 | 1 |
| ZNF770 | 0.267697834 | 0.220789293 | 0.1 | 0.105 | 1 |
| TRAF3 | 0.26849214 | 0.15231683 | 0.1 | 0.145 | 1 |
| RNF19A | 0.269305437 | 0.476995745 | 0.2 | 0.23 | 1 |
| PCYOX1 | 0.272593993 | 0.310576966 | 0.1 | 0.11 | 1 |
| KCTD1 | 0.274938481 | 0.176622351 | 0.2 | 0.204 | 1 |
| CNIH1 | 0.280726118 | 0.180875061 | 0.1 | 0.113 | 1 |
| ABI1 | 0.282434888 | 0.495728304 | 0.2 | 0.21 | 1 |
| FBXW7 | 0.284496328 | 0.44021838 | 0.3 | 0.316 | 1 |
| MRPL41 | 0.286943801 | 0.389556472 | 0.1 | 0.131 | 1 |
| FOCAD | 0.289136987 | 0.555934163 | 0.2 | 0.209 | 1 |
| KHDRBS3 | 0.291358709 | 0.49684045 | 0.3 | 0.312 | 1 |
| BEX3 | 0.299004539 | 0.639171887 | 0.2 | 0.19 | 1 |
| EFR3B | 0.305724162 | 0.331484662 | 0.2 | 0.198 | 1 |
| ATP5MPL | 0.312894244 | 0.249993907 | 0.1 | 0.11 | 1 |
| LMTK2 | 0.316434551 | 0.300454958 | 0.2 | 0.188 | 1 |
| ATP5F1B | 0.322281202 | 0.545750048 | 0.1 | 0.149 | 1 |
| PPP5C | 0.325959111 | 0.260850799 | 0.1 | 0.121 | 1 |
| FGFR1OP | 0.326900952 | 0.266348135 | 0.1 | 0.118 | 1 |
| LINGO1 | 0.327630573 | 0.795998293 | 0.6 | 0.781 | 1 |
| ATN1 | 0.327824471 | 0.377868695 | 0.2 | 0.171 | 1 |
| SLC22A17 | 0.32813024 | 0.620922027 | 0.3 | 0.292 | 1 |
| SLC39A10 | 0.330767876 | 0.607448651 | 0.2 | 0.235 | 1 |
| NDUFA4 | 0.331465699 | 0.512060744 | 0.1 | 0.11 | 1 |
| IRS2 | 0.334506383 | 0.354292921 | 0.2 | 0.255 | 1 |
| SAE1 | 0.336621444 | 0.194754528 | 0.1 | 0.13 | 1 |
| CDH13 | 0.336951013 | 0.37471251 | 0.3 | 0.285 | 1 |
| APC2 | 0.33809878 | 0.336919077 | 0.2 | 0.17 | 1 |
| MARK1 | 0.341650609 | 0.298394269 | 0.2 | 0.162 | 1 |
| EPB41L5 | 0.34614411 | 0.140662473 | 0.2 | 0.167 | 1 |
| AL357153.2 | 0.347581046 | 0.209953435 | 0.1 | 0.118 | 1 |
| TPI1 | 0.348910484 | 0.484204815 | 0.1 | 0.111 | 1 |
| ATP5F1A | 0.351866144 | 0.302938921 | 0.1 | 0.135 | 1 |
| MTPN | 0.351903981 | 0.478442833 | 0.1 | 0.113 | 1 |
| SMARCA1 | 0.354136542 | 0.275353436 | 0.1 | 0.112 | 1 |
| SCRN1 | 0.35699578 | 0.496804945 | 0.1 | 0.154 | 1 |
| RTF1 | 0.362116747 | 0.297071089 | 0.2 | 0.178 | 1 |
| GFOD1 | 0.362734269 | 0.44150088 | 0.2 | 0.181 | 1 |
| NR3C2 | 0.363507283 | 0.380018558 | 0.3 | 0.309 | 1 |
| CYSTM1 | 0.364714943 | 0.256231026 | 0.1 | 0.111 | 1 |
| GHITM | 0.369402256 | 0.47433247 | 0.1 | 0.097 | 1 |
| NRXN2 | 0.373590696 | 0.38895229 | 0.1 | 0.124 | 1 |
| BSG | 0.373794521 | 0.407541637 | 0.2 | 0.213 | 1 |
| CCDC171 | 0.37521587 | 0.300725184 | 0.2 | 0.17 | 1 |
| PIP4K2B | 0.37655302 | 0.369308975 | 0.2 | 0.154 | 1 |
| MAPK9 | 0.376786686 | 0.244722868 | 0.2 | 0.186 | 1 |
| STXBP5 | 0.377068455 | 0.568611628 | 0.4 | 0.476 | 1 |
| FBXL17 | 0.38052402 | 0.39847072 | 0.4 | 0.446 | 1 |
| TMEM63B | 0.381952057 | 0.229209071 | 0.1 | 0.1 | 1 |
| ARMC9 | 0.388194976 | 0.294657454 | 0.1 | 0.094 | 1 |
| NDUFA5 | 0.390279693 | 0.333346375 | 0.1 | 0.101 | 1 |
| TENM3 | 0.411409121 | 0.22193765 | 0.2 | 0.169 | 1 |
| GPR155 | 0.41316556 | 0.556766218 | 0.2 | 0.174 | 1 |
| PITPNA | 0.419679774 | 0.340620767 | 0.1 | 0.099 | 1 |
| ISPD | 0.419717972 | 0.202167661 | 0.1 | 0.147 | 1 |
| PURG | 0.427933426 | 0.274024699 | 0.1 | 0.149 | 1 |
| MXD4 | 0.429534691 | 0.188942507 | 0.1 | 0.116 | 1 |
| ZFYVE9 | 0.429895513 | 0.368805165 | 0.2 | 0.222 | 1 |
| ATP6V1G1 | 0.435090942 | 0.241293824 | 0.1 | 0.109 | 1 |
| LINC00623 | 0.43550778 | 0.348668513 | 0.1 | 0.144 | 1 |
| KLHL23 | 0.439137941 | 0.20160937 | 0.1 | 0.105 | 1 |
| CRB1 | 0.45945538 | 0.346086639 | 0.2 | 0.249 | 1 |
| PPP2R1A | 0.460071623 | 0.546663898 | 0.2 | 0.215 | 1 |
| EIF4A2 | 0.460111926 | 0.900889785 | 0.3 | 0.342 | 1 |
| AC004076.1 | 0.460512899 | 0.170511845 | 0.1 | 0.1 | 1 |
| RASGEF1B | 0.47226619 | 0.617276085 | 0.6 | 0.755 | 1 |
| SKI | 0.478422878 | 0.215326527 | 0.2 | 0.178 | 1 |
| ABLIM2 | 0.482052675 | 0.332814197 | 0.1 | 0.105 | 1 |
| TMED8 | 0.485001949 | 0.310806976 | 0.1 | 0.102 | 1 |
| PDXK | 0.485026805 | 0.343779626 | 0.1 | 0.118 | 1 |
| ZMAT3 | 0.489579465 | 0.254612102 | 0.1 | 0.108 | 1 |
| HS6ST1 | 0.497070645 | 0.274678452 | 0.1 | 0.102 | 1 |
| C2orf72 | 0.498458836 | 0.289119841 | 0.1 | 0.13 | 1 |
| R3HDM1 | 0.504402627 | 0.488165255 | 0.5 | 0.496 | 1 |
| PREX2 | 0.507892522 | 0.431817663 | 0.4 | 0.396 | 1 |
| NDRG3 | 0.515561385 | 0.437549408 | 0.2 | 0.167 | 1 |
| CDH2 | 0.518843529 | 0.248528332 | 0.3 | 0.315 | 1 |
| PISD | 0.523560307 | 0.191196818 | 0.1 | 0.105 | 1 |
| FUT9 | 0.524691942 | 0.313972294 | 0.4 | 0.485 | 1 |
| ST7 | 0.530288347 | 0.295221322 | 0.2 | 0.224 | 1 |
| LRRC8A | 0.53167168 | 0.26875904 | 0.1 | 0.103 | 1 |
| KCNH1 | 0.535743089 | 0.229171589 | 0.1 | 0.104 | 1 |
| CRKL | 0.538741391 | 0.254408756 | 0.1 | 0.134 | 1 |
| SLC3A2 | 0.541852878 | 0.300396341 | 0.1 | 0.108 | 1 |
| PDE10A | 0.542355875 | 0.338604446 | 0.2 | 0.151 | 1 |
| HS2ST1 | 0.542615889 | 0.193126997 | 0.1 | 0.114 | 1 |
| IQCJ-SCHIP1 | 0.54826523 | 0.444595243 | 0.5 | 0.536 | 1 |
| RTN3 | 0.552728592 | 0.730051404 | 0.6 | 0.653 | 1 |
| DPP7 | 0.555439941 | 0.367203782 | 0.2 | 0.166 | 1 |
| PRKACA | 0.556533004 | 0.258225155 | 0.1 | 0.111 | 1 |
| CMIP | 0.558404518 | 0.521094048 | 0.2 | 0.236 | 1 |
| KIFAP3 | 0.55888681 | 0.487826991 | 0.2 | 0.248 | 1 |
| ENO1 | 0.560100759 | 0.427068256 | 0.2 | 0.182 | 1 |
| ZBTB18 | 0.562135449 | 0.45315192 | 0.1 | 0.106 | 1 |
| SPATS2 | 0.573060087 | 0.192865144 | 0.1 | 0.136 | 1 |
| ME1 | 0.581722324 | 0.242190023 | 0.1 | 0.131 | 1 |
| DIAPH2 | 0.591254905 | 0.226016682 | 0.1 | 0.113 | 1 |
| EHBP1 | 0.591393115 | 0.376015377 | 0.4 | 0.392 | 1 |
| GGA2 | 0.593758536 | 0.190246642 | 0.1 | 0.123 | 1 |
| PNPLA7 | 0.594707587 | 0.226433004 | 0.2 | 0.159 | 1 |
| ULK4 | 0.595765337 | 0.249043658 | 0.1 | 0.137 | 1 |
| CEP126 | 0.596392619 | 0.276015367 | 0.1 | 0.152 | 1 |
| HINT1 | 0.601797849 | 0.514651289 | 0.2 | 0.176 | 1 |
| NRG2 | 0.603065223 | 0.202385504 | 0.2 | 0.164 | 1 |
| LRP12 | 0.603632992 | 0.247794108 | 0.1 | 0.101 | 1 |
| AP2A2 | 0.603703866 | 0.447902725 | 0.2 | 0.207 | 1 |
| PRELID2 | 0.607357144 | 0.175235702 | 0.1 | 0.105 | 1 |

| | | | | | | | | | | | |
|---|---|---|---|---|---|---|---|---|---|---|---|
| APMAP | 0.609012371 | 0.317255828 | 0.1 | 0.103 | 1 | | | | | | |
| SPRED2 | 0.61104832 | 0.191353793 | 0.1 | 0.13 | 1 | | | | | | |
| LMO7 | 0.618338505 | 0.221462654 | 0.2 | 0.16 | 1 | | | | | | |
| AC087500.1 | 0.619945358 | 0.17418193 | 0.1 | 0.111 | 1 | | | | | | |
| BDH1 | 0.631431012 | 0.219413651 | 0.1 | 0.106 | 1 | | | | | | |
| ARHGEF12 | 0.637001747 | 0.428611123 | 0.3 | 0.347 | 1 | | | | | | |
| FP700111.1 | 0.646594265 | 0.428320707 | 0.1 | 0.146 | 1 | | | | | | |
| AL589740.1 | 0.648637026 | 0.345819827 | 0.4 | 0.392 | 1 | | | | | | |
| ATP5F1D | 0.651323153 | 0.299292871 | 0.1 | 0.122 | 1 | | | | | | |
| TNRC18 | 0.653667664 | 0.187059512 | 0.2 | 0.184 | 1 | | | | | | |
| FRY | 0.661142866 | 0.192164812 | 0.3 | 0.251 | 1 | | | | | | |
| DTX3 | 0.663572886 | 0.272643869 | 0.1 | 0.115 | 1 | | | | | | |
| TRIM37 | 0.667085529 | 0.342434381 | 0.2 | 0.169 | 1 | | | | | | |
| MAP3K13 | 0.673425198 | 0.330667146 | 0.2 | 0.196 | 1 | | | | | | |
| NEK11 | 0.677738988 | 0.336516817 | 0.1 | 0.136 | 1 | | | | | | |
| FAM85B | 0.678751411 | 0.202974274 | 0.2 | 0.216 | 1 | | | | | | |
| BLCAP | 0.682545047 | 0.360165634 | 0.1 | 0.122 | 1 | | | | | | |
| KAZN | 0.684073826 | 0.328363776 | 0.6 | 0.705 | 1 | | | | | | |
| SYNM | 0.693476755 | 0.432023452 | 0.1 | 0.108 | 1 | | | | | | |
| SOCS7 | 0.71165033 | 0.188758488 | 0.1 | 0.144 | 1 | | | | | | |
| ACOT7 | 0.727624678 | 0.30409357 | 0.1 | 0.128 | 1 | | | | | | |
| MCF2L2 | 0.751072563 | 0.250967893 | 0.2 | 0.195 | 1 | | | | | | |
| EFR3A | 0.751080952 | 0.485414912 | 0.2 | 0.262 | 1 | | | | | | |
| NMD3 | 0.751584723 | 0.250073967 | 0.1 | 0.104 | 1 | | | | | | |
| NPTN | 0.752237672 | 0.628274096 | 0.2 | 0.243 | 1 | | | | | | |
| GALNT11 | 0.752366586 | 0.378680572 | 0.1 | 0.119 | 1 | | | | | | |
| ISCU | 0.756014936 | 0.422327829 | 0.1 | 0.103 | 1 | | | | | | |
| SLF1 | 0.772009439 | 0.296572162 | 0.1 | 0.104 | 1 | | | | | | |
| MT-ND4L | 0.780747418 | 0.283470511 | 0.2 | 0.228 | 1 | | | | | | |
| WASF1 | 0.78081015 | 0.483614821 | 0.2 | 0.206 | 1 | | | | | | |
| NFE2L1 | 0.782297123 | 0.487899114 | 0.1 | 0.112 | 1 | | | | | | |
| TMSB4X | 0.800140217 | 0.671378127 | 0.3 | 0.278 | 1 | | | | | | |
| USP11 | 0.800521175 | 0.619467448 | 0.2 | 0.216 | 1 | | | | | | |
| CRIM1 | 0.816723658 | 0.301699118 | 0.2 | 0.178 | 1 | | | | | | |
| APLP2 | 0.828195718 | 0.660907474 | 0.2 | 0.212 | 1 | | | | | | |
| MAPRE3 | 0.829073153 | 0.439581871 | 0.2 | 0.171 | 1 | | | | | | |
| MIR99AHG | 0.835513095 | 0.200676608 | 0.7 | 0.8 | 1 | | | | | | |
| WDR19 | 0.835795458 | 0.175615445 | 0.1 | 0.129 | 1 | | | | | | |
| GNPTAB | 0.837490241 | 0.242474466 | 0.1 | 0.128 | 1 | | | | | | |
| CDIP1 | 0.844739882 | 0.463006717 | 0.1 | 0.122 | 1 | | | | | | |
| TSC22D1 | 0.848313879 | 0.596803101 | 0.4 | 0.417 | 1 | | | | | | |
| MT-ND5 | 0.850636437 | 0.370459152 | 0.4 | 0.493 | 1 | | | | | | |
| GLB1 | 0.852050641 | 0.222667205 | 0.1 | 0.124 | 1 | | | | | | |
| ROCK2 | 0.856186909 | 0.435045423 | 0.3 | 0.283 | 1 | | | | | | |
| MIB2 | 0.862911761 | 0.147733938 | 0.1 | 0.1 | 1 | | | | | | |
| SEZ6L2 | 0.877774532 | 0.442066603 | 0.1 | 0.135 | 1 | | | | | | |
| GPR37L1 | 0.888798389 | 0.325965188 | 0.1 | 0.145 | 1 | | | | | | |
| FAIM2 | 0.891690786 | 0.830952925 | 0.3 | 0.35 | 1 | | | | | | |
| KIAA0825 | 0.896308579 | 0.14151711 | 0.2 | 0.167 | 1 | | | | | | |
| ANKRD26 | 0.896878879 | 0.353228374 | 0.2 | 0.264 | 1 | | | | | | |
| SSX2IP | 0.90284648 | 0.259702658 | 0.1 | 0.114 | 1 | | | | | | |
| BEND6 | 0.910551093 | 0.192522685 | 0.1 | 0.114 | 1 | | | | | | |
| HSP90AB1 | 0.912216595 | 0.850250044 | 0.4 | 0.473 | 1 | | | | | | |
| RBFOX2 | 0.913168538 | 0.385699318 | 0.4 | 0.415 | 1 | | | | | | |
| PGK1 | 0.92061781 | 0.4277281 | 0.1 | 0.122 | 1 | | | | | | |
| WDR47 | 0.921372831 | 0.324957545 | 0.1 | 0.148 | 1 | | | | | | |
| PATJ | 0.924783756 | 0.218688708 | 0.2 | 0.167 | 1 | | | | | | |
| MTMR1 | 0.942763027 | 0.269639782 | 0.1 | 0.145 | 1 | | | | | | |
| FAM171B | 0.944975738 | 0.447293745 | 0.2 | 0.253 | 1 | | | | | | |
| FP236383.1 | 0.951408749 | 0.51832509 | 0.3 | 0.347 | 1 | | | | | | |
| SNU13 | 0.95857961 | 0.385293139 | 0.1 | 0.103 | 1 | | | | | | |
| SLC9A6 | 0.960579296 | 0.516025216 | 0.1 | 0.138 | 1 | | | | | | |
| BEX2 | 0.964546726 | 0.42474182 | 0.1 | 0.116 | 1 | | | | | | |
| JUN | 0.965989184 | 0.52948819 | 0.1 | 0.12 | 1 | | | | | | |
| B4GALNT4 | 0.969572426 | 0.409155576 | 0.1 | 0.153 | 1 | | | | | | |
| CRTC1 | 0.970456373 | 0.272224686 | 0.2 | 0.166 | 1 | | | | | | |
| ZNF346 | 0.972398519 | 0.267350435 | 0.2 | 0.166 | 1 | | | | | | |
| TPST1 | 0.973876166 | 0.78919134 | 0.2 | 0.25 | 1 | | | | | | |
| ADCY9 | 0.978329175 | 0.22589351 | 0.2 | 0.152 | 1 | | | | | | |
| TNRC6A | 0.986638108 | 0.406192493 | 0.6 | 0.745 | 1 | | | | | | |
| DPP6 | 0.988120172 | 0.152693254 | 0.5 | 0.506 | 1 | | | | | | |
| SLC27A6 | 0.989087063 | 0.385771904 | 0.1 | 0.149 | 1 | | | | | | |
| LRIG1 | 0.993207272 | 0.506391757 | 0.3 | 0.356 | 1 | | | | | | |
| BAG4 | 0.996394474 | 0.26568382 | 0.1 | 0.102 | 1 | | | | | | |
| SPAG16 | 0.999631456 | 0.425249457 | 0.2 | 0.209 | 1 | | | | | | |
| Figure7F_correlation_analysis_of_hOL3_mOL3 | | | | | | | | | | | |
| | | hOL3 DEGs versus others | | | | | mOL3 DEGs versus others | | | | |
| | gene | p_val.x | log2 | pct.1.x | pct.2.x | p_val_adj.x | p_val.y | avg_log2FC.y | pct.1.y | pct.2.y | p_val_adj.y |
| 1 | Nrxn1 | 5.57E-300 | 2.9 | 0.706 | 0.212 | 0 | 1.53E-117 | 0.9386433 | 0.397 | 0.286 | 4.11E-113 |
| 2 | Tenm2 | 5.57E-300 | 2 | 0.393 | 0.068 | 0 | 2.53E-88 | 0.7878718 | 0.491 | 0.383 | 6.82E-84 |
| 3 | Celf2 | 5.57E-300 | 1.9 | 0.468 | 0.098 | 0 | 1.49E-156 | 1.1296813 | 0.401 | 0.197 | 4.00E-152 |
| 4 | Nrg3 | 5.57E-300 | 3.3 | 0.693 | 0.155 | 0 | 2.27E-184 | 1.3627809 | 0.367 | 0.152 | 6.10E-180 |
| 5 | Syt1 | 5.57E-300 | 3 | 0.585 | 0.177 | 0 | 1.82E-125 | 1.0108495 | 0.298 | 0.181 | 4.91E-121 |
| 6 | Dlgap1 | 5.57E-300 | 2.3 | 0.501 | 0.122 | 0 | 6.19E-114 | 1.0053564 | 0.273 | 0.127 | 1.67E-109 |
| 7 | Opcml | 1.89E-259 | 2.3 | 0.441 | 0.121 | 5.45E-255 | 1.84E-62 | 0.4650881 | 0.469 | 0.44 | 4.97E-58 |
| 8 | Rbfox1 | 1.64E-223 | 2.2 | 0.597 | 0.258 | 4.73E-219 | 1.97E-202 | 1.4164724 | 0.352 | 0.124 | 5.30E-198 |
| 9 | Cadps | 3.60E-217 | 1.6 | 0.276 | 0.055 | 1.04E-212 | 1.65E-29 | 0.3646701 | 0.253 | 0.237 | 4.44E-25 |
| 10 | Ccser1 | 2.79E-211 | 1.7 | 0.348 | 0.089 | 8.08E-207 | 1.31E-26 | 0.2051003 | 0.35 | 0.366 | 3.52E-22 |
| 11 | Kcnip4 | 7.78E-184 | 2.3 | 0.628 | 0.326 | 2.25E-179 | 3.50E-256 | 1.6381486 | 0.485 | 0.189 | 9.43E-252 |
| 12 | Ctnna2 | 2.95E-170 | 1.4 | 0.622 | 0.289 | 8.52E-166 | 5.38E-24 | -0.3020753 | 0.492 | 0.604 | 1.45E-19 |
| 13 | Pclo | 4.30E-167 | 1.5 | 0.304 | 0.081 | 1.24E-162 | 6.27E-49 | 0.2343999 | 0.23 | 0.254 | 1.69E-44 |
| 14 | Pip4k2a | 1.03E-166 | -0.8 | 0.669 | 0.967 | 2.99E-162 | 1.06E-57 | -0.0709371 | 0.506 | 0.629 | 2.85E-53 |
| 15 | Cdh10 | 1.37E-163 | 1 | 0.157 | 0.023 | 3.95E-159 | 2.91E-35 | -0.0173067 | 0.405 | 0.493 | 7.82E-31 |
| 16 | Slc8a1 | 2.62E-156 | 1.8 | 0.278 | 0.073 | 7.58E-152 | 4.33E-25 | -0.1156015 | 0.675 | 0.77 | 1.17E-20 |
| 17 | Csmd1 | 4.81E-151 | 2.2 | 0.655 | 0.427 | 1.39E-146 | 6.45E-188 | 1.3842016 | 0.37 | 0.142 | 1.74E-183 |
| 18 | Elmo1 | 4.09E-142 | -0.7 | 0.664 | 0.958 | 1.18E-137 | 2.75E-53 | -0.3009512 | 0.673 | 0.818 | 7.40E-49 |
| 19 | Grik2 | 1.36E-141 | 1.6 | 0.338 | 0.11 | 3.93E-137 | 1.13E-33 | 0.0782481 | 0.64 | 0.702 | 3.03E-29 |
| 20 | Camk1d | 9.29E-139 | 1.2 | 0.237 | 0.058 | 2.68E-134 | 1.19E-73 | 0.2956243 | 0.286 | 0.308 | 3.20E-69 |
| 21 | Plp1 | 1.82E-138 | -0.9 | 0.662 | 0.926 | 5.25E-134 | 2.41E-38 | -0.1824402 | 0.965 | 0.988 | 6.50E-34 |
| 22 | Rnf220 | 5.33E-134 | -0.8 | 0.638 | 0.964 | 1.54E-129 | 1.39E-49 | -0.2448315 | 0.687 | 0.824 | 3.74E-45 |
| 23 | Qk | 5.59E-134 | -0.6 | 0.773 | 0.987 | 1.62E-129 | 2.85E-12 | -0.0672984 | 0.956 | 0.979 | 7.66E-08 |
| 24 | Mbp | 4.60E-132 | -0.7 | 0.676 | 0.963 | 1.33E-127 | 1.03E-60 | -0.2266098 | 0.967 | 0.991 | 2.76E-56 |
| 25 | Map7 | 1.91E-129 | -0.7 | 0.645 | 0.943 | 5.52E-125 | 6.99E-29 | -0.1596245 | 0.842 | 0.918 | 1.88E-24 |
| 26 | St18 | 9.36E-129 | -0.8 | 0.647 | 0.952 | 2.71E-124 | 2.09E-26 | -0.1608128 | 0.926 | 0.97 | 5.63E-22 |
| 27 | Il1rapl1 | 1.03E-128 | -0.7 | 0.764 | 0.967 | 2.97E-124 | 1.58E-26 | 0.1317922 | 0.591 | 0.637 | 4.25E-22 |
| 28 | Dock5 | 2.95E-125 | -0.7 | 0.609 | 0.941 | 8.52E-121 | 1.89E-62 | -0.1985786 | 0.232 | 0.352 | 5.09E-58 |
| 29 | Dock10 | 3.46E-124 | -0.7 | 0.621 | 0.948 | 1.00E-119 | 3.41E-18 | -0.0145419 | 0.897 | 0.947 | 9.19E-14 |
| 30 | Slc44a1 | 1.54E-120 | -0.7 | 0.672 | 0.962 | 4.44E-116 | 6.57E-72 | -0.2126273 | 0.453 | 0.62 | 1.77E-67 |
| 31 | Frmd5 | 3.12E-120 | -0.7 | 0.673 | 0.954 | 9.01E-116 | 7.97E-11 | -0.087797 | 0.923 | 0.956 | 2.14E-06 |
| 32 | Ctnna3 | 3.48E-120 | -0.7 | 0.687 | 0.957 | 1.01E-115 | 2.64E-46 | -0.3029403 | 0.653 | 0.792 | 7.10E-42 |
| 33 | Ugt8a | 3.57E-119 | -0.8 | 0.499 | 0.838 | 1.03E-114 | 8.17E-60 | -0.1345834 | 0.477 | 0.616 | 2.20E-55 |
| 34 | Pde4b | 2.90E-116 | -0.6 | 0.74 | 0.966 | 8.39E-112 | 1.48E-08 | -0.0662269 | 0.996 | 0.998 | 0.000399651 |
| 35 | Phlpp1 | 7.81E-116 | -0.7 | 0.655 | 0.958 | 2.26E-111 | 3.60E-12 | 0.0040292 | 0.907 | 0.944 | 9.69E-08 |
| 36 | Plcl1 | 9.12E-116 | -0.8 | 0.643 | 0.945 | 2.63E-111 | 2.56E-30 | -0.2050348 | 0.95 | 0.98 | 6.88E-26 |
| 37 | Frmd4b | 4.43E-115 | -0.7 | 0.605 | 0.935 | 1.28E-110 | 9.43E-50 | -0.0611923 | 0.545 | 0.661 | 2.54E-45 |
| 38 | D7Ertd443e | 1.77E-114 | -0.7 | 0.589 | 0.923 | 5.13E-110 | 7.05E-63 | -0.5714155 | 0.245 | 0.418 | 1.90E-58 |
| 39 | Mog | 3.31E-114 | -0.7 | 0.504 | 0.846 | 9.56E-110 | 2.84E-48 | -0.0850814 | 0.658 | 0.782 | 7.65E-44 |
| 40 | Unc5c | 9.65E-114 | -0.8 | 0.651 | 0.938 | 2.79E-109 | 4.12E-22 | -0.0546625 | 0.728 | 0.807 | 1.11E-17 |
| 41 | Edil3 | 2.36E-113 | -0.7 | 0.667 | 0.955 | 6.82E-109 | 2.32E-15 | -0.0979972 | 0.952 | 0.978 | 6.25E-11 |
| 42 | Slain1 | 1.27E-112 | -0.8 | 0.599 | 0.914 | 3.68E-108 | 1.74E-55 | -0.1009684 | 0.427 | 0.55 | 4.68E-51 |
| 43 | Dnm3 | 1.52E-111 | -0.6 | 0.694 | 0.961 | 4.39E-107 | 2.33E-45 | -0.1376641 | 0.739 | 0.857 | 6.27E-41 |
| 44 | Plekhh1 | 2.11E-111 | -0.7 | 0.558 | 0.898 | 6.10E-107 | 1.71E-74 | -0.1042169 | 0.439 | 0.58 | 4.61E-70 |
| 45 | Bcas1 | 7.85E-111 | -0.7 | 0.521 | 0.865 | 2.27E-106 | 3.13E-69 | -0.2513162 | 0.455 | 0.625 | 8.43E-65 |
| 46 | Erbin | 3.68E-110 | -0.7 | 0.671 | 0.959 | 1.06E-105 | 1.34E-43 | 0.063689 | 0.653 | 0.731 | 3.60E-39 |
| 47 | Pde8a | 1.49E-109 | -0.7 | 0.599 | 0.931 | 4.31E-105 | 1.03E-54 | -0.0389966 | 0.524 | 0.64 | 2.78E-50 |
| 48 | Lrrtm4 | 3.24E-109 | 1.4 | 0.549 | 0.287 | 9.37E-105 | 7.60E-148 | 1.2133241 | 0.274 | 0.095 | 2.05E-143 |
| 49 | Shtn1 | 4.38E-108 | -0.7 | 0.64 | 0.936 | 1.27E-103 | 7.96E-59 | -0.0479687 | 0.472 | 0.589 | 2.14E-54 |
| 50 | Sik3 | 3.53E-106 | -0.6 | 0.72 | 0.979 | 1.02E-101 | 1.00E-39 | -0.1165769 | 0.641 | 0.759 | 2.71E-35 |
| 51 | Picalm | 3.78E-105 | -0.7 | 0.567 | 0.905 | 1.09E-100 | 4.90E-64 | -0.1651552 | 0.33 | 0.462 | 1.32E-59 |
| 52 | Dlg1 | 4.79E-104 | -0.7 | 0.635 | 0.945 | 1.38E-99 | 7.89E-54 | 0.0640857 | 0.538 | 0.619 | 2.12E-49 |
| 53 | Spock3 | 9.65E-104 | -0.7 | 0.637 | 0.93 | 2.79E-99 | 2.38E-39 | -0.2653518 | 0.413 | 0.546 | 6.40E-35 |
| 54 | Pde1c | 1.07E-103 | -0.7 | 0.623 | 0.932 | 3.09E-99 | 9.22E-34 | 0.0240339 | 0.256 | 0.312 | 2.48E-29 |
| 55 | Slc24a2 | 1.24E-103 | -0.6 | 0.713 | 0.953 | 3.57E-99 | 4.66E-24 | -0.1413356 | 0.979 | 0.989 | 1.25E-19 |
| 56 | Enpp2 | 4.87E-103 | -0.6 | 0.57 | 0.903 | 1.41E-98 | 2.34E-27 | -0.0708267 | 0.758 | 0.846 | 6.31E-23 |
| 57 | Zfp536 | 1.34E-102 | -0.7 | 0.599 | 0.923 | 3.87E-98 | 3.82E-30 | -0.2077166 | 0.765 | 0.86 | 1.03E-25 |
| 58 | Xkr4 | 1.99E-102 | 0.9 | 0.152 | 0.033 | 5.75E-98 | 3.35E-31 | 0.2338968 | 0.391 | 0.402 | 9.01E-27 |
| 59 | Ptprk | 3.48E-102 | -0.7 | 0.656 | 0.949 | 1.01E-97 | 4.32E-35 | -0.1879204 | 0.653 | 0.773 | 1.16E-30 |

| | | | | | | | | | | | |
|---|---|---|---|---|---|---|---|---|---|---|---|
| 60 | Fryl | 6.13E-102 | -0.6 | 0.639 | 0.953 | 1.77E-97 | 3.38E-53 | -0.0174996 | 0.436 | 0.537 | 9.11E-49 |
| 61 | Clmn | 8.66E-101 | -0.6 | 0.598 | 0.925 | 2.50E-96 | 2.88E-45 | 0.0401942 | 0.498 | 0.577 | 7.75E-41 |
| 62 | Pcdh9 | 1.37E-99 | -0.5 | 0.931 | 0.976 | 3.95E-95 | 1.74E-33 | -0.170995 | 0.997 | 0.999 | 4.68E-29 |
| 63 | Fa2h | 2.18E-99 | -0.6 | 0.475 | 0.804 | 6.31E-95 | 1.05E-61 | -0.1030979 | 0.477 | 0.613 | 2.82E-57 |
| 64 | Sgk1 | 8.08E-99 | -0.7 | 0.589 | 0.911 | 2.33E-94 | 6.77E-64 | -0.2897447 | 0.257 | 0.403 | 1.82E-59 |
| 65 | Dtna | 5.08E-97 | 2 | 0.559 | 0.343 | 1.47E-92 | 1.32E-58 | 0.14615 | 0.341 | 0.395 | 3.56E-54 |
| 66 | Mag | 7.02E-97 | -0.7 | 0.423 | 0.729 | 2.03E-92 | 3.98E-66 | -0.173423 | 0.553 | 0.711 | 1.07E-61 |
| 67 | Creb5 | 1.05E-96 | -0.7 | 0.572 | 0.902 | 3.02E-92 | 3.58E-56 | -0.0127032 | 0.314 | 0.404 | 9.63E-52 |
| 68 | Dock1 | 1.82E-96 | -0.6 | 0.564 | 0.892 | 5.26E-92 | 7.20E-60 | 0.0061035 | 0.464 | 0.566 | 1.94E-55 |
| 69 | Tcf12 | 1.96E-96 | -0.6 | 0.661 | 0.961 | 5.66E-92 | 7.60E-49 | -0.1005121 | 0.615 | 0.742 | 2.05E-44 |
| 70 | Qdpr | 1.46E-95 | -0.7 | 0.527 | 0.845 | 4.22E-91 | 4.10E-55 | -0.0139393 | 0.219 | 0.293 | 1.10E-50 |
| 71 | Clasp2 | 2.30E-95 | -0.6 | 0.685 | 0.955 | 6.64E-91 | 4.32E-25 | 0.0307389 | 0.736 | 0.8 | 1.16E-20 |
| 72 | Agap1 | 4.43E-95 | -0.6 | 0.671 | 0.969 | 1.28E-90 | 3.71E-54 | -0.1300071 | 0.584 | 0.724 | 9.98E-50 |
| 73 | Dnajc6 | 9.20E-95 | -0.6 | 0.627 | 0.909 | 2.66E-90 | 3.23E-42 | 0.1097139 | 0.251 | 0.295 | 8.69E-38 |
| 74 | Tmtc2 | 3.39E-94 | -0.7 | 0.671 | 0.946 | 9.81E-90 | 3.52E-57 | -0.1475286 | 0.325 | 0.454 | 9.47E-53 |
| 75 | Slco3a1 | 5.07E-94 | -0.6 | 0.574 | 0.901 | 1.47E-89 | 5.60E-58 | -0.5773188 | 0.141 | 0.282 | 1.51E-53 |
| 76 | Synj2 | 5.70E-94 | -0.7 | 0.51 | 0.817 | 1.65E-89 | 1.73E-46 | -0.0607497 | 0.27 | 0.35 | 4.65E-42 |
| 77 | Ttll7 | 2.42E-93 | -0.6 | 0.671 | 0.951 | 6.98E-89 | 4.55E-61 | -0.0298941 | 0.556 | 0.679 | 1.23E-56 |
| 78 | Abca2 | 2.94E-93 | -0.6 | 0.582 | 0.894 | 8.50E-89 | 2.23E-70 | -0.1842323 | 0.199 | 0.316 | 6.01E-66 |
| 79 | Adipor2 | 2.70E-92 | -0.7 | 0.489 | 0.813 | 7.80E-88 | 7.56E-55 | -0.2864704 | 0.535 | 0.695 | 2.04E-50 |
| 80 | Dip2b | 7.95E-90 | -0.6 | 0.624 | 0.941 | 2.30E-85 | 1.41E-46 | -0.0990459 | 0.498 | 0.615 | 3.80E-42 |
| 81 | Gab1 | 1.89E-89 | -0.6 | 0.605 | 0.918 | 5.47E-85 | 5.73E-56 | -0.1617144 | 0.455 | 0.595 | 1.54E-51 |
| 82 | Gpm6b | 7.50E-89 | -0.6 | 0.687 | 0.93 | 2.17E-84 | 1.23E-43 | -0.2204632 | 0.686 | 0.816 | 3.32E-39 |
| 83 | Mobp | 3.28E-88 | -0.7 | 0.556 | 0.867 | 9.48E-84 | 7.03E-63 | -0.2924214 | 0.642 | 0.803 | 1.89E-58 |
| 84 | Spock1 | 4.42E-88 | -0.6 | 0.694 | 0.929 | 1.28E-83 | 1.05E-24 | -0.2618217 | 0.39 | 0.499 | 2.84E-20 |
| 85 | Jam3 | 4.73E-85 | -0.6 | 0.486 | 0.797 | 1.37E-80 | 4.15E-82 | -0.0113133 | 0.282 | 0.388 | 1.12E-77 |
| 86 | Zeb2 | 4.28E-84 | -0.6 | 0.651 | 0.949 | 1.24E-79 | 5.98E-15 | -0.0245753 | 0.839 | 0.893 | 1.61E-10 |
| 87 | Dock9 | 6.85E-84 | -0.6 | 0.554 | 0.873 | 1.98E-79 | 5.98E-59 | -0.1968971 | 0.397 | 0.541 | 1.61E-54 |
| 88 | Apbb2 | 7.48E-84 | -0.6 | 0.6 | 0.918 | 2.16E-79 | 7.04E-47 | -0.1264008 | 0.348 | 0.463 | 1.89E-42 |
| 89 | Cntn2 | 7.97E-84 | -0.6 | 0.534 | 0.842 | 2.30E-79 | 1.19E-71 | 0.0318049 | 0.359 | 0.452 | 3.21E-67 |
| 90 | Cdk19 | 8.92E-84 | -0.6 | 0.566 | 0.879 | 2.58E-79 | 1.20E-31 | -0.088156 | 0.671 | 0.772 | 3.22E-27 |
| 91 | Usp54 | 1.05E-83 | -0.6 | 0.558 | 0.877 | 3.04E-79 | 3.26E-67 | -0.164656 | 0.432 | 0.579 | 8.79E-63 |
| 92 | Phldb1 | 2.40E-83 | -0.6 | 0.479 | 0.803 | 6.93E-79 | 1.38E-60 | 0.0880161 | 0.526 | 0.606 | 3.71E-56 |
| 93 | Hip1 | 2.22E-82 | -0.6 | 0.533 | 0.856 | 6.40E-78 | 6.06E-54 | -0.112619 | 0.232 | 0.329 | 1.63E-49 |
| 94 | Hipk2 | 6.09E-82 | -0.6 | 0.556 | 0.879 | 1.76E-77 | 1.04E-58 | 0.0051366 | 0.478 | 0.584 | 2.80E-54 |
| 95 | Lpgat1 | 8.75E-82 | -0.6 | 0.526 | 0.827 | 2.53E-77 | 1.30E-58 | 0.008436 | 0.403 | 0.5 | 3.49E-54 |
| 96 | Dip2c | 4.50E-81 | -0.5 | 0.643 | 0.935 | 1.30E-76 | 1.96E-46 | 0.1224394 | 0.263 | 0.314 | 5.27E-42 |
| 97 | Nfasc | 7.57E-81 | -0.5 | 0.578 | 0.892 | 2.19E-76 | 5.91E-35 | -0.0535606 | 0.576 | 0.672 | 1.59E-30 |
| 98 | Pex5l | 1.94E-80 | -0.6 | 0.663 | 0.934 | 5.60E-76 | 1.20E-48 | -0.3319779 | 0.774 | 0.887 | 3.23E-44 |
| 99 | Map4k4 | 2.21E-80 | -0.5 | 0.65 | 0.937 | 6.40E-76 | 2.38E-62 | -0.0966885 | 0.352 | 0.469 | 6.41E-58 |
| 100 | Sh3d19 | 6.72E-80 | -0.6 | 0.516 | 0.824 | 1.94E-75 | 1.94E-55 | -0.0624868 | 0.433 | 0.552 | 5.23E-51 |
| 101 | Ttyh2 | 1.15E-79 | -0.5 | 0.441 | 0.742 | 3.33E-75 | 1.11E-66 | -0.0071454 | 0.354 | 0.456 | 2.99E-62 |
| 102 | Hapln2 | 1.29E-79 | -0.7 | 0.382 | 0.678 | 3.71E-75 | 7.79E-100 | -0.329748 | 0.246 | 0.427 | 2.10E-95 |
| 103 | Kif13b | 1.49E-79 | -0.6 | 0.457 | 0.756 | 4.31E-75 | 1.74E-49 | -0.1480739 | 0.532 | 0.666 | 4.69E-45 |
| 104 | Arhgap21 | 7.32E-79 | -0.5 | 0.673 | 0.938 | 2.11E-74 | 8.69E-63 | -0.1792271 | 0.307 | 0.438 | 2.34E-58 |
| 105 | Myo1d | 1.46E-78 | -0.6 | 0.51 | 0.825 | 4.23E-74 | 6.52E-68 | 0.0137545 | 0.406 | 0.509 | 1.75E-63 |
| 106 | Ank3 | 1.55E-78 | -0.5 | 0.721 | 0.954 | 4.49E-74 | 1.61E-20 | 0.0195728 | 0.817 | 0.875 | 4.35E-16 |
| 107 | Ano4 | 3.38E-78 | -0.6 | 0.574 | 0.863 | 9.76E-74 | 3.56E-21 | 0.118353 | 0.821 | 0.851 | 9.59E-17 |
| 108 | Daam2 | 4.01E-78 | -0.5 | 0.526 | 0.847 | 1.16E-73 | 3.45E-70 | -0.3271233 | 0.463 | 0.646 | 9.30E-66 |
| 109 | 2010111I01Rik | 1.16E-76 | -0.5 | 0.563 | 0.884 | 3.34E-72 | 7.63E-17 | 0.1074859 | 0.295 | 0.31 | 2.05E-12 |
| 110 | Lsamp | 1.43E-76 | 0.9 | 0.861 | 0.841 | 4.13E-72 | 8.06E-104 | 0.7707859 | 0.632 | 0.514 | 2.17E-99 |
| 111 | Palm2 | 8.50E-74 | -0.7 | 0.532 | 0.812 | 2.46E-69 | 3.96E-05 | 0.0883368 | 0.505 | 0.496 | 1 |
| 112 | Dscaml1 | 9.51E-74 | -0.6 | 0.531 | 0.833 | 2.75E-69 | 1.20E-43 | -0.1761428 | 0.596 | 0.731 | 3.23E-39 |
| 113 | Fmnl2 | 9.60E-74 | -0.5 | 0.713 | 0.958 | 2.77E-69 | 7.46E-41 | -0.1680331 | 0.467 | 0.593 | 2.01E-36 |
| 114 | Tulp4 | 1.09E-73 | -0.5 | 0.54 | 0.848 | 3.14E-69 | 1.55E-42 | -0.064918 | 0.535 | 0.641 | 4.16E-38 |
| 115 | Slc12a2 | 1.86E-73 | -0.5 | 0.447 | 0.752 | 5.36E-69 | 5.48E-48 | -0.0865047 | 0.499 | 0.616 | 1.47E-43 |
| 116 | Atg4c | 8.00E-73 | -0.5 | 0.457 | 0.758 | 2.31E-68 | 1.82E-57 | -0.1166266 | 0.181 | 0.273 | 4.91E-53 |
| 117 | Pkp4 | 8.29E-73 | -0.6 | 0.541 | 0.825 | 2.40E-68 | 5.28E-54 | -0.0071086 | 0.479 | 0.584 | 1.42E-49 |
| 118 | Tjp1 | 9.45E-73 | -0.5 | 0.647 | 0.94 | 2.73E-68 | 1.39E-50 | 0.0867287 | 0.32 | 0.383 | 3.73E-46 |
| 119 | Kcnq3 | 1.14E-72 | 1.2 | 0.241 | 0.09 | 3.31E-68 | 3.56E-19 | -0.0076785 | 0.633 | 0.7 | 9.58E-15 |
| 120 | Map4k5 | 1.69E-72 | -0.5 | 0.565 | 0.874 | 4.88E-68 | 1.22E-52 | 0.0424062 | 0.349 | 0.426 | 3.29E-48 |
| 121 | Amph | 1.35E-71 | 0.8 | 0.107 | 0.023 | 3.89E-67 | 2.26E-52 | 0.1214457 | 0.291 | 0.338 | 6.09E-48 |
| 122 | Cdh19 | 1.57E-71 | -0.6 | 0.42 | 0.696 | 4.55E-67 | 1.65E-45 | -0.3081319 | 0.39 | 0.535 | 4.45E-41 |
| 123 | Fgfr2 | 7.14E-70 | -0.5 | 0.528 | 0.813 | 2.06E-65 | 2.70E-35 | -0.0767324 | 0.825 | 0.912 | 7.27E-31 |
| 124 | Trim2 | 8.89E-70 | -0.5 | 0.669 | 0.929 | 2.57E-65 | 7.02E-44 | 0.0049134 | 0.539 | 0.632 | 1.89E-39 |
| 125 | Clic4 | 9.40E-69 | -0.5 | 0.423 | 0.717 | 2.72E-64 | 2.36E-55 | -0.0745339 | 0.257 | 0.35 | 6.37E-51 |
| 126 | Kdm6a | 1.41E-68 | -0.5 | 0.461 | 0.759 | 4.08E-64 | 7.78E-58 | -0.0318446 | 0.276 | 0.369 | 2.09E-53 |
| 127 | Anln | 1.60E-68 | -0.6 | 0.438 | 0.727 | 4.63E-64 | 4.89E-112 | -0.7209217 | 0.237 | 0.471 | 1.32E-107 |
| 128 | Akap6 | 4.25E-68 | -0.5 | 0.667 | 0.924 | 1.23E-63 | 5.35E-49 | 0.0343556 | 0.38 | 0.46 | 1.44E-44 |
| 129 | Slc22a23 | 1.49E-67 | -0.5 | 0.47 | 0.764 | 4.30E-63 | 1.30E-35 | 0.0139591 | 0.232 | 0.286 | 3.49E-31 |
| 130 | Atp8a1 | 2.53E-67 | -0.5 | 0.63 | 0.905 | 7.31E-63 | 5.97E-35 | -0.078848 | 0.665 | 0.768 | 1.61E-30 |
| 131 | Aatk | 5.36E-67 | -0.5 | 0.485 | 0.787 | 1.55E-62 | 2.54E-66 | 0.1105356 | 0.467 | 0.545 | 6.83E-62 |
| 132 | Alcam | 1.09E-66 | -0.5 | 0.603 | 0.889 | 3.14E-62 | 3.08E-35 | -0.114206 | 0.183 | 0.259 | 8.30E-31 |
| 133 | Sema6a | 1.71E-66 | -0.5 | 0.479 | 0.764 | 4.95E-62 | 1.65E-51 | 0.1556478 | 0.327 | 0.378 | 4.45E-47 |
| 134 | Tmem63a | 2.87E-66 | -0.5 | 0.362 | 0.647 | 8.28E-62 | 9.98E-55 | -0.0365316 | 0.215 | 0.292 | 2.69E-50 |
| 135 | Igsf11 | 3.50E-66 | -0.5 | 0.463 | 0.747 | 1.01E-61 | 8.31E-62 | -0.15587 | 0.175 | 0.273 | 2.24E-57 |
| 136 | Aplp1 | 6.54E-66 | -0.5 | 0.557 | 0.822 | 1.89E-61 | 3.05E-61 | -0.0796262 | 0.381 | 0.501 | 8.21E-57 |
| 137 | Dlg2 | 1.02E-65 | -0.3 | 0.847 | 0.984 | 2.96E-61 | 3.72E-28 | 0.0930667 | 0.81 | 0.87 | 1.00E-23 |
| 138 | Fut8 | 1.76E-65 | -0.5 | 0.606 | 0.891 | 5.09E-61 | 1.19E-48 | -0.0600516 | 0.199 | 0.277 | 3.20E-44 |
| 139 | Megf10 | 2.52E-65 | -0.5 | 0.467 | 0.745 | 7.29E-61 | 2.59E-67 | -0.1496424 | 0.205 | 0.316 | 6.96E-63 |
| 140 | Elf2 | 4.92E-65 | -0.5 | 0.489 | 0.786 | 1.42E-60 | 2.61E-52 | -0.1055079 | 0.169 | 0.25 | 7.03E-48 |
| 141 | Arhgap23 | 1.81E-64 | -0.5 | 0.396 | 0.668 | 5.24E-60 | 1.97E-45 | -0.0808918 | 0.657 | 0.775 | 5.29E-41 |
| 142 | Mal | 1.41E-63 | -0.6 | 0.37 | 0.642 | 4.06E-59 | 1.46E-61 | -0.1688908 | 0.363 | 0.501 | 3.93E-57 |
| 143 | Magi2 | 1.63E-63 | -0.4 | 0.824 | 0.983 | 4.70E-59 | 0.036742626 | -0.0095749 | 0.992 | 0.995 | 1 |
| 144 | Arap2 | 3.27E-63 | -0.5 | 0.649 | 0.933 | 9.44E-59 | 3.41E-45 | 0.0200739 | 0.325 | 0.396 | 9.18E-41 |
| 145 | Ncam2 | 5.30E-63 | -0.5 | 0.787 | 0.953 | 1.53E-58 | 1.39E-37 | -0.1687714 | 0.63 | 0.75 | 3.73E-33 |
| 146 | Ppp2r2a | 6.90E-63 | -0.5 | 0.456 | 0.755 | 1.99E-58 | 6.38E-74 | -0.0940762 | 0.259 | 0.373 | 1.72E-69 |
| 147 | Cobl | 1.21E-62 | -0.5 | 0.577 | 0.869 | 3.48E-58 | 6.85E-46 | -0.0003585 | 0.217 | 0.283 | 1.84E-41 |
| 148 | Kif13a | 1.22E-62 | -0.5 | 0.515 | 0.819 | 3.54E-58 | 9.34E-49 | 0.0598413 | 0.495 | 0.577 | 2.51E-44 |
| 149 | Rnf13 | 2.23E-62 | -0.5 | 0.47 | 0.751 | 6.45E-58 | 9.75E-67 | -0.0411203 | 0.386 | 0.498 | 2.63E-62 |
| 150 | Zbtb20 | 2.37E-62 | -0.4 | 0.749 | 0.981 | 6.84E-58 | 1.72E-41 | 0.0578109 | 0.747 | 0.831 | 4.62E-37 |
| 151 | Slc25a13 | 2.71E-62 | -0.6 | 0.4 | 0.664 | 7.83E-58 | 4.94E-57 | -0.1864302 | 0.252 | 0.371 | 1.33E-52 |
| 152 | Macrod2 | 3.03E-62 | 1.1 | 0.362 | 0.183 | 8.76E-58 | 2.72E-20 | 0.1738508 | 0.37 | 0.377 | 7.31E-16 |
| 153 | Stk39 | 3.89E-62 | -0.5 | 0.428 | 0.696 | 1.13E-57 | 6.10E-52 | -0.0479789 | 0.219 | 0.298 | 1.64E-47 |
| 154 | Limch1 | 1.47E-61 | -0.5 | 0.651 | 0.921 | 4.24E-57 | 3.88E-40 | 0.0119157 | 0.595 | 0.684 | 1.04E-35 |
| 155 | Ptprd | 1.51E-61 | -0.3 | 0.803 | 0.96 | 4.36E-57 | 0.02131165 | 0.0217936 | 0.983 | 0.99 | 1 |
| 156 | Tmeff2 | 3.03E-61 | -0.5 | 0.584 | 0.858 | 8.75E-57 | 2.57E-07 | -0.0425151 | 0.963 | 0.981 | 0.00692046 |
| 157 | Bin1 | 9.44E-61 | -0.4 | 0.51 | 0.814 | 2.73E-56 | 1.34E-48 | -0.0316206 | 0.609 | 0.72 | 3.62E-44 |
| 158 | Kat2b | 1.41E-60 | -0.4 | 0.443 | 0.722 | 4.08E-56 | 1.05E-59 | -2.57E-05 | 0.268 | 0.352 | 2.84E-55 |
| 159 | Ptk2 | 2.17E-60 | -0.5 | 0.675 | 0.947 | 6.27E-56 | 3.02E-64 | -0.0820365 | 0.473 | 0.606 | 8.13E-60 |
| 160 | Grm3 | 2.55E-60 | -0.5 | 0.594 | 0.871 | 7.36E-56 | 6.53E-14 | -0.0418252 | 0.788 | 0.848 | 1.76E-09 |
| 161 | Gnai1 | 2.88E-60 | -0.5 | 0.455 | 0.74 | 8.33E-56 | 1.28E-52 | -0.0050212 | 0.281 | 0.361 | 3.43E-48 |
| 162 | Agpat4 | 4.23E-60 | -0.5 | 0.442 | 0.711 | 1.22E-55 | 6.53E-64 | -0.1865425 | 0.225 | 0.341 | 1.76E-59 |
| 163 | Pik3c2b | 5.31E-60 | -0.5 | 0.331 | 0.597 | 1.53E-55 | 4.65E-64 | -0.0084176 | 0.274 | 0.363 | 1.25E-59 |
| 164 | Osbpl1a | 5.86E-60 | -0.5 | 0.564 | 0.837 | 1.69E-55 | 7.41E-63 | -0.0496575 | 0.311 | 0.413 | 1.99E-58 |
| 165 | Nav3 | 1.38E-59 | 1.4 | 0.572 | 0.426 | 3.99E-55 | 3.58E-31 | 0.4196039 | 0.257 | 0.223 | 9.63E-27 |
| 166 | Tnrc6c | 6.79E-59 | -0.4 | 0.491 | 0.795 | 1.96E-54 | 1.76E-62 | 0.0922954 | 0.335 | 0.404 | 4.72E-58 |
| 167 | Wipf1 | 4.28E-58 | -0.5 | 0.311 | 0.561 | 1.24E-53 | 2.10E-49 | -0.1124785 | 0.255 | 0.349 | 5.64E-45 |
| 168 | Ccp110 | 8.90E-58 | -0.4 | 0.468 | 0.751 | 2.57E-53 | 8.80E-52 | -0.0479436 | 0.35 | 0.446 | 2.37E-47 |
| 169 | Myo6 | 1.02E-57 | -0.4 | 0.555 | 0.853 | 2.94E-53 | 1.49E-59 | -0.0544741 | 0.486 | 0.607 | 4.01E-55 |
| 170 | Ankib1 | 1.05E-57 | -0.4 | 0.559 | 0.847 | 3.03E-53 | 1.16E-63 | 0.0773432 | 0.352 | 0.429 | 3.13E-59 |
| 171 | Pam | 1.13E-57 | 0.7 | 0.166 | 0.055 | 3.26E-53 | 1.97E-38 | 0.0703692 | 0.207 | 0.255 | 5.31E-34 |
| 172 | Oxr1 | 1.21E-57 | 1.3 | 0.388 | 0.224 | 3.51E-53 | 8.40E-30 | -0.0805802 | 0.524 | 0.617 | 2.26E-25 |
| 173 | Sort1 | 3.46E-57 | -0.4 | 0.603 | 0.875 | 1.00E-52 | 3.98E-50 | -0.0316386 | 0.255 | 0.337 | 1.07E-45 |
| 174 | Itch | 3.76E-57 | -0.4 | 0.487 | 0.774 | 1.09E-52 | 4.80E-68 | -0.0929091 | 0.413 | 0.543 | 1.29E-63 |
| 175 | Nkain2 | 9.52E-57 | -0.6 | 0.692 | 0.896 | 2.75E-52 | 1.09E-60 | -0.2088735 | 0.998 | 0.999 | 2.93E-56 |
| 176 | Zfp638 | 2.05E-56 | -0.4 | 0.656 | 0.938 | 5.92E-52 | 3.13E-45 | -0.003644 | 0.475 | 0.566 | 8.42E-41 |
| 177 | Ermn | 2.57E-56 | -0.5 | 0.327 | 0.585 | 7.42E-52 | 3.91E-70 | -0.2153406 | 0.256 | 0.392 | 1.05E-65 |
| 178 | Cttnbp2 | 3.90E-56 | -0.4 | 0.441 | 0.721 | 1.13E-51 | 1.23E-63 | 0.1699917 | 0.458 | 0.521 | 3.30E-59 |
| 179 | Ncam1 | 1.46E-55 | -0.4 | 0.73 | 0.94 | 4.23E-51 | 6.66E-37 | -0.1543664 | 0.769 | 0.874 | 1.79E-32 |
| 180 | St6galnac3 | 3.41E-55 | -0.6 | 0.47 | 0.73 | 9.84E-51 | 1.87E-32 | -0.1962129 | 0.816 | 0.903 | 5.05E-28 |
| 181 | Arid4b | 4.69E-55 | -0.4 | 0.501 | 0.778 | 1.36E-50 | 5.01E-69 | -0.0276706 | 0.331 | 0.435 | 1.35E-64 |
| 182 | Zdhhc20 | 1.21E-54 | -0.6 | 0.42 | 0.67 | 3.50E-50 | 1.13E-54 | -0.271361 | 0.455 | 0.612 | 3.03E-50 |
| 183 | Chd7 | 1.31E-54 | -0.4 | 0.409 | 0.675 | 3.78E-50 | 2.34E-52 | 0.082183 | 0.282 | 0.341 | 6.30E-48 |
| 184 | Zbtb16 | 1.76E-54 | -0.5 | 0.499 | 0.775 | 5.10E-50 | 6.41E-70 | -0.268392 | 0.269 | 0.42 | 1.73E-65 |
| 185 | Opalin | 2.36E-54 | -0.5 | 0.368 | 0.618 | 6.81E-50 | 1.52E-42 | 0.1734436 | 0.297 | 0.325 | 4.08E-38 |
| 186 | Faf1 | 3.72E-54 | -0.4 | 0.55 | 0.831 | 1.08E-49 | 1.81E-57 | 0.000283 | 0.213 | 0.286 | 4.87E-53 |
| 187 | Prr5l | 4.66E-54 | -0.5 | 0.36 | 0.609 | 1.35E-49 | 4.17E-56 | -0.3011935 | 0.811 | 0.916 | 1.12E-51 |
| 188 | Secisbp2l | 4.68E-54 | -0.4 | 0.512 | 0.783 | 1.35E-49 | 1.79E-63 | -0.1821828 | 0.185 | 0.289 | 4.81E-59 |
| 189 | Epn2 | 6.45E-54 | -0.4 | 0.525 | 0.803 | 1.86E-49 | 8.62E-62 | -0.1058763 | 0.337 | 0.453 | 2.32E-57 |
| 190 | Mbnl2 | 6.94E-54 | -0.4 | 0.612 | 0.899 | 2.01E-49 | 1.10E-38 | -0.0898594 | 0.683 | 0.794 | 2.97E-34 |
| 191 | Ypel2 | 9.97E-54 | -0.5 | 0.367 | 0.622 | 2.88E-49 | 2.35E-51 | -0.0792899 | 0.541 | 0.662 | 6.34E-47 |
| 192 | Kif6 | 1.63E-53 | -0.5 | 0.421 | 0.679 | 4.72E-49 | 1.12E-36 | -0.0785613 | 0.182 | 0.251 | 3.03E-32 |
| 193 | Auts2 | 2.11E-53 | -0.4 | 0.688 | 0.942 | 6.11E-49 | 6.68E-27 | 0.1623899 | 0.683 | 0.707 | 1.80E-22 |
| 194 | Ppp1r16b | 2.58E-53 | -0.4 | 0.446 | 0.707 | 7.45E-49 | 2.09E-39 | -0.0501191 | 0.628 | 0.735 | 5.64E-35 |
| 195 | Wdr20 | 9.96E-53 | -0.4 | 0.384 | 0.625 | 2.88E-48 | 2.14E-51 | -0.0811896 | 0.194 | 0.273 | 5.75E-47 |
| 196 | Pacs2 | 1.72E-52 | -0.4 | 0.395 | 0.654 | 4.96E-48 | 1.13E-62 | -0.0533503 | 0.238 | 0.326 | 3.05E-58 |
| 197 | Kirrel3 | 8.77E-52 | -0.5 | 0.561 | 0.824 | 2.53E-47 | 1.95E-22 | 0.3644775 | 0.594 | 0.51 | 5.25E-18 |

| | | | | | | | | | | | |
|---|---|---|---|---|---|---|---|---|---|---|---|
| 198 | Apod | 1.84E-51 | -0.6 | 0.307 | 0.543 | 5.31E-47 | 1.86E-66 | -0.5037021 | 0.351 | 0.537 | 5.00E-62 |
| 199 | Ssh2 | 2.56E-51 | -0.4 | 0.579 | 0.861 | 7.39E-47 | 3.95E-47 | -0.1074311 | 0.537 | 0.663 | 1.06E-42 |
| 200 | Exoc6b | 4.01E-51 | -0.4 | 0.673 | 0.938 | 1.16E-46 | 3.34E-36 | -0.1094245 | 0.628 | 0.739 | 8.99E-32 |
| 201 | Car2 | 8.03E-51 | -0.5 | 0.259 | 0.5 | 2.32E-46 | 3.88E-42 | -0.0266784 | 0.588 | 0.693 | 1.04E-37 |
| 202 | Nsmce2 | 8.63E-51 | -0.4 | 0.474 | 0.74 | 2.49E-46 | 5.80E-57 | -0.0132596 | 0.176 | 0.25 | 1.56E-52 |
| 203 | Pard3 | 4.05E-50 | 0.8 | 0.23 | 0.099 | 1.17E-45 | 6.86E-70 | -0.1987452 | 0.35 | 0.501 | 1.85E-65 |
| 204 | Ccdc88a | 5.65E-50 | -0.4 | 0.671 | 0.939 | 1.63E-45 | 8.19E-62 | 0.0552955 | 0.407 | 0.493 | 2.20E-57 |
| 205 | Lama2 | 6.41E-50 | -1 | 0.335 | 0.56 | 1.85E-45 | 1.28E-28 | -0.1221547 | 0.277 | 0.358 | 3.45E-24 |
| 206 | Aspa | 1.85E-49 | -0.4 | 0.394 | 0.65 | 5.36E-45 | 7.96E-49 | -0.0291961 | 0.65 | 0.766 | 2.14E-44 |
| 207 | Nlgn1 | 3.25E-49 | -0.5 | 0.705 | 0.906 | 9.40E-45 | 2.70E-41 | 0.3044787 | 0.568 | 0.581 | 7.26E-37 |
| 208 | Otud7a | 4.20E-49 | -0.4 | 0.623 | 0.893 | 1.21E-44 | 5.78E-42 | -0.2495687 | 0.44 | 0.581 | 1.56E-37 |
| 209 | Eps15 | 4.92E-49 | -0.3 | 0.498 | 0.762 | 1.42E-44 | 1.87E-62 | 0.0261904 | 0.257 | 0.335 | 5.03E-58 |
| 210 | Kansl1 | 6.19E-49 | -0.3 | 0.506 | 0.775 | 1.79E-44 | 1.11E-57 | 0.0223473 | 0.342 | 0.429 | 2.99E-53 |
| 211 | Atp11a | 1.48E-48 | -0.3 | 0.387 | 0.642 | 4.26E-44 | 6.68E-61 | -0.0626928 | 0.181 | 0.264 | 1.80E-56 |
| 212 | Cyth1 | 1.51E-48 | -0.4 | 0.371 | 0.623 | 4.37E-44 | 3.49E-72 | -0.0994491 | 0.298 | 0.42 | 9.39E-68 |
| 213 | Dennd5a | 2.84E-48 | -0.3 | 0.467 | 0.721 | 8.22E-44 | 1.88E-64 | -0.1052866 | 0.234 | 0.338 | 5.06E-60 |
| 214 | Fnbp1 | 3.16E-48 | -0.4 | 0.587 | 0.876 | 9.15E-44 | 1.43E-18 | -0.0459778 | 0.819 | 0.883 | 3.84E-14 |
| 215 | Hecw2 | 7.47E-48 | -0.4 | 0.526 | 0.796 | 2.16E-43 | 3.18E-45 | -0.2091863 | 0.708 | 0.836 | 8.56E-41 |
| 216 | Ralgapa1 | 1.08E-47 | -0.3 | 0.51 | 0.777 | 3.12E-43 | 3.02E-61 | 0.090587 | 0.3 | 0.368 | 8.13E-57 |
| 217 | Plekhg3 | 1.10E-47 | -0.4 | 0.319 | 0.544 | 3.17E-43 | 8.54E-64 | -0.0123449 | 0.222 | 0.301 | 2.30E-59 |
| 218 | Sorcs1 | 7.85E-47 | 1 | 0.129 | 0.042 | 2.27E-42 | 1.37E-47 | -0.0964395 | 0.536 | 0.659 | 3.69E-43 |
| 219 | Ppp2r3a | 3.00E-46 | -0.4 | 0.362 | 0.593 | 8.66E-42 | 3.50E-47 | -0.0996595 | 0.53 | 0.652 | 9.43E-43 |
| 220 | Spag9 | 4.37E-46 | -0.3 | 0.508 | 0.771 | 1.26E-41 | 1.04E-69 | -0.0243414 | 0.362 | 0.47 | 2.80E-65 |
| 221 | Rere | 5.39E-46 | -0.3 | 0.624 | 0.895 | 1.56E-41 | 5.59E-49 | -0.142765 | 0.504 | 0.637 | 1.50E-44 |
| 222 | Nrd1 | 5.40E-46 | -0.3 | 0.354 | 0.589 | 1.56E-41 | 3.10E-64 | -0.0626595 | 0.217 | 0.308 | 8.35E-60 |
| 223 | Sox10 | 6.13E-46 | -0.5 | 0.267 | 0.489 | 1.77E-41 | 5.71E-58 | -0.0002333 | 0.277 | 0.362 | 1.54E-53 |
| 224 | Retreg1 | 6.73E-46 | -0.4 | 0.465 | 0.723 | 1.95E-41 | 3.90E-63 | -0.040659 | 0.277 | 0.376 | 1.05E-58 |
| 225 | Wnk1 | 1.66E-45 | -0.3 | 0.516 | 0.781 | 4.81E-41 | 9.45E-58 | 0.0221701 | 0.524 | 0.624 | 2.54E-53 |
| 226 | Ppp1r21 | 1.68E-45 | -0.3 | 0.446 | 0.697 | 4.84E-41 | 3.13E-60 | 0.0059434 | 0.215 | 0.288 | 8.42E-56 |
| 227 | Grm7 | 2.49E-45 | 1 | 0.26 | 0.129 | 7.18E-41 | 1.15E-07 | 0.0051722 | 0.457 | 0.477 | 0.00308767 |
| 228 | Tacc1 | 2.87E-45 | -0.5 | 0.494 | 0.757 | 8.28E-41 | 5.54E-58 | -0.2403284 | 0.19 | 0.306 | 1.49E-53 |
| 229 | Zfp280d | 7.63E-45 | -0.3 | 0.4 | 0.637 | 2.20E-40 | 2.45E-71 | -0.0544467 | 0.234 | 0.335 | 6.59E-67 |
| 230 | Stxbp3 | 1.07E-44 | -0.3 | 0.334 | 0.562 | 3.10E-40 | 1.66E-60 | -0.1505994 | 0.226 | 0.334 | 4.47E-56 |
| 231 | Nipbl | 1.48E-44 | -0.3 | 0.498 | 0.762 | 4.28E-40 | 4.86E-74 | 0.0344934 | 0.288 | 0.378 | 1.31E-69 |
| 232 | Pcdh15 | 7.66E-44 | 0.9 | 0.128 | 0.044 | 2.21E-39 | 1.89E-29 | 0.2436337 | 0.303 | 0.311 | 5.08E-25 |
| 233 | Rasgrp3 | 1.65E-43 | -0.5 | 0.273 | 0.482 | 4.75E-39 | 3.79E-49 | -0.2256922 | 0.161 | 0.258 | 1.02E-44 |
| 234 | Bcas3 | 1.98E-43 | -0.4 | 0.534 | 0.788 | 5.73E-39 | 4.49E-67 | -0.1437303 | 0.327 | 0.463 | 1.21E-62 |
| 235 | Neo1 | 4.50E-43 | -0.3 | 0.554 | 0.822 | 1.30E-38 | 1.22E-65 | 0.0315571 | 0.349 | 0.439 | 3.27E-61 |
| 236 | Csrp1 | 4.53E-43 | -0.4 | 0.367 | 0.6 | 1.31E-38 | 1.21E-62 | -0.1277487 | 0.308 | 0.427 | 3.25E-58 |
| 237 | Srcin1 | 4.54E-43 | -0.4 | 0.474 | 0.721 | 1.31E-38 | 5.99E-62 | -0.0603082 | 0.321 | 0.425 | 1.61E-57 |
| 238 | Zfp704 | 7.36E-43 | -0.4 | 0.489 | 0.743 | 2.13E-38 | 2.61E-41 | -0.0169313 | 0.214 | 0.278 | 7.03E-37 |
| 239 | Prox1 | 1.12E-42 | -0.4 | 0.217 | 0.421 | 3.22E-38 | 2.80E-50 | -0.0908718 | 0.177 | 0.257 | 7.54E-46 |
| 240 | Fam13b | 1.25E-42 | -0.3 | 0.448 | 0.694 | 3.60E-38 | 2.04E-51 | -0.0315013 | 0.198 | 0.269 | 5.49E-47 |
| 241 | Kmt2e | 2.36E-42 | -0.2 | 0.494 | 0.752 | 6.83E-38 | 1.30E-57 | -0.0087649 | 0.278 | 0.361 | 3.49E-53 |
| 242 | Tbc1d5 | 2.95E-42 | -0.3 | 0.58 | 0.844 | 8.52E-38 | 7.63E-30 | -0.1505717 | 0.702 | 0.807 | 2.05E-25 |
| 243 | Rap1a | 7.09E-42 | -0.3 | 0.377 | 0.619 | 2.05E-37 | 3.13E-54 | -0.0497844 | 0.27 | 0.358 | 8.43E-50 |
| 244 | Cpm | 1.11E-41 | -0.5 | 0.315 | 0.533 | 3.20E-37 | 1.71E-53 | -0.0928981 | 0.187 | 0.271 | 4.60E-49 |
| 245 | Enox1 | 1.62E-41 | -0.4 | 0.609 | 0.861 | 4.68E-37 | 3.53E-41 | -0.1061929 | 0.575 | 0.694 | 9.51E-37 |
| 246 | Gnao1 | 1.79E-41 | -0.4 | 0.596 | 0.858 | 5.16E-37 | 1.01E-54 | -0.2244945 | 0.56 | 0.713 | 2.71E-50 |
| 247 | Tanc1 | 1.86E-41 | -0.4 | 0.295 | 0.514 | 5.37E-37 | 7.28E-49 | -0.0122896 | 0.204 | 0.271 | 1.96E-44 |
| 248 | Gatm | 2.34E-41 | -0.3 | 0.293 | 0.512 | 6.78E-37 | 1.17E-61 | 0.036254 | 0.518 | 0.62 | 3.15E-57 |
| 249 | Cpeb2 | 3.87E-41 | -0.3 | 0.427 | 0.676 | 1.12E-36 | 1.19E-72 | -0.0068742 | 0.244 | 0.335 | 3.20E-68 |
| 250 | Snx29 | 4.81E-41 | -0.3 | 0.395 | 0.623 | 1.39E-36 | 2.46E-58 | -0.0909593 | 0.238 | 0.336 | 6.61E-54 |
| 251 | Lrch3 | 2.70E-40 | -0.3 | 0.411 | 0.657 | 7.80E-36 | 7.40E-63 | -0.1597467 | 0.275 | 0.394 | 1.99E-58 |
| 252 | Otud7b | 4.62E-40 | -0.3 | 0.422 | 0.66 | 1.34E-35 | 1.59E-73 | -0.1588955 | 0.421 | 0.572 | 4.28E-69 |
| 253 | Trip12 | 5.83E-40 | -0.2 | 0.392 | 0.618 | 1.68E-35 | 4.31E-54 | 0.0428177 | 0.226 | 0.289 | 1.16E-49 |
| 254 | Ago3 | 5.87E-40 | -0.2 | 0.373 | 0.607 | 1.70E-35 | 6.34E-53 | -0.0646527 | 0.222 | 0.304 | 1.71E-48 |
| 255 | Rap1gds1 | 9.67E-40 | -0.3 | 0.45 | 0.695 | 2.79E-35 | 8.03E-42 | -0.14801 | 0.178 | 0.26 | 2.16E-37 |
| 256 | Ankrd28 | 1.72E-39 | -0.3 | 0.498 | 0.756 | 4.96E-35 | 2.71E-64 | -0.1252172 | 0.449 | 0.587 | 7.29E-60 |
| 257 | Acyp2 | 2.60E-39 | -0.3 | 0.488 | 0.732 | 7.51E-35 | 4.69E-26 | 0.0014024 | 0.255 | 0.308 | 1.26E-21 |
| 258 | Ubr3 | 3.18E-39 | -0.2 | 0.501 | 0.756 | 9.20E-35 | 1.93E-61 | 0.0523273 | 0.245 | 0.313 | 5.19E-57 |
| 259 | Hepacam | 3.52E-39 | -0.3 | 0.391 | 0.627 | 1.02E-34 | 1.13E-55 | -0.1160855 | 0.301 | 0.407 | 3.05E-51 |
| 260 | Ywhaq | 4.13E-39 | -0.3 | 0.366 | 0.583 | 1.19E-34 | 1.25E-63 | -0.0832635 | 0.313 | 0.424 | 3.37E-59 |
| 261 | Abhd17b | 1.46E-38 | -0.3 | 0.335 | 0.565 | 4.21E-34 | 1.46E-54 | -0.0430084 | 0.47 | 0.582 | 3.93E-50 |
| 262 | Ralgps1 | 1.61E-38 | -0.3 | 0.313 | 0.526 | 4.66E-34 | 1.49E-50 | -0.0826196 | 0.327 | 0.427 | 4.02E-46 |
| 263 | Sltm | 1.57E-37 | -0.2 | 0.335 | 0.541 | 4.55E-33 | 1.19E-63 | 0.0143242 | 0.296 | 0.38 | 3.21E-59 |
| 264 | Ikzf2 | 2.63E-37 | -0.3 | 0.343 | 0.557 | 7.60E-33 | 4.00E-57 | -0.1352411 | 0.196 | 0.299 | 1.08E-52 |
| 265 | Vmp1 | 2.65E-37 | -0.3 | 0.332 | 0.549 | 7.66E-33 | 2.36E-56 | -0.1874729 | 0.642 | 0.789 | 6.35E-52 |
| 266 | Ube2e2 | 3.38E-37 | -0.4 | 0.608 | 0.833 | 9.77E-33 | 8.38E-24 | 0.0970897 | 0.677 | 0.719 | 2.26E-19 |
| 267 | Rtn4 | 5.80E-37 | -0.3 | 0.703 | 0.93 | 1.68E-32 | 1.15E-38 | -0.0355315 | 0.717 | 0.816 | 3.09E-34 |
| 268 | Arglu1 | 1.17E-36 | -0.1 | 0.519 | 0.781 | 3.38E-32 | 1.92E-55 | 0.0370539 | 0.331 | 0.41 | 5.17E-51 |
| 269 | Ptn | 1.36E-36 | -0.3 | 0.291 | 0.501 | 3.92E-32 | 3.60E-58 | -0.203623 | 0.394 | 0.538 | 9.69E-54 |
| 270 | Kidins220 | 2.12E-36 | -0.2 | 0.486 | 0.74 | 6.14E-32 | 3.78E-55 | 0.00612 | 0.19 | 0.255 | 1.02E-50 |
| 271 | Vps13b | 2.64E-36 | -0.2 | 0.432 | 0.66 | 7.62E-32 | 6.03E-60 | 0.0958801 | 0.286 | 0.346 | 1.62E-55 |
| 272 | Ptma | 2.72E-36 | -0.2 | 0.474 | 0.716 | 7.86E-32 | 2.80E-55 | -0.0103108 | 0.197 | 0.268 | 7.52E-51 |
| 273 | Gatad2b | 5.11E-36 | -0.2 | 0.37 | 0.591 | 1.48E-31 | 5.85E-52 | -0.0813657 | 0.189 | 0.271 | 1.58E-47 |
| 274 | Snx30 | 8.56E-36 | -0.3 | 0.343 | 0.545 | 2.48E-31 | 9.70E-55 | -0.1025206 | 0.237 | 0.332 | 2.61E-50 |
| 275 | Tra2a | 9.11E-36 | -0.2 | 0.456 | 0.703 | 2.63E-31 | 7.93E-73 | -0.0092362 | 0.323 | 0.428 | 2.13E-68 |
| 276 | Phactr3 | 1.01E-35 | -0.3 | 0.563 | 0.804 | 2.91E-31 | 2.08E-49 | 0.0176935 | 0.242 | 0.309 | 5.59E-45 |
| 277 | Tmcc3 | 1.14E-35 | -0.4 | 0.225 | 0.411 | 3.28E-31 | 4.64E-65 | -0.2736271 | 0.354 | 0.511 | 1.25E-60 |
| 278 | Ninj2 | 8.11E-35 | -0.4 | 0.275 | 0.467 | 2.34E-30 | 2.08E-40 | 0.0273036 | 0.38 | 0.457 | 5.59E-36 |
| 279 | Rps6ka5 | 8.83E-35 | -0.3 | 0.368 | 0.567 | 2.55E-30 | 1.90E-55 | -0.0800472 | 0.221 | 0.31 | 5.12E-51 |
| 280 | Zfp532 | 6.03E-34 | -0.2 | 0.409 | 0.624 | 1.74E-29 | 4.20E-68 | -0.1432558 | 0.235 | 0.351 | 1.13E-63 |
| 281 | Dnm2 | 6.50E-34 | -0.3 | 0.297 | 0.496 | 1.88E-29 | 2.33E-65 | -0.0816464 | 0.189 | 0.282 | 6.28E-61 |
| 282 | Krit1 | 7.08E-34 | -0.3 | 0.262 | 0.457 | 2.05E-29 | 6.47E-61 | 0.0057323 | 0.214 | 0.289 | 1.74E-56 |
| 283 | Phip | 7.20E-34 | -0.2 | 0.523 | 0.77 | 2.08E-29 | 3.75E-55 | 0.1851417 | 0.247 | 0.282 | 1.01E-50 |
| 284 | Cdc42bpa | 8.64E-34 | -0.3 | 0.683 | 0.93 | 2.50E-29 | 3.01E-25 | -0.1731163 | 0.755 | 0.846 | 8.10E-21 |
| 285 | App | 1.01E-33 | -0.2 | 0.66 | 0.913 | 2.91E-29 | 3.57E-51 | -0.0217443 | 0.664 | 0.776 | 9.61E-47 |
| 286 | Kmt2c | 1.12E-33 | -0.2 | 0.578 | 0.843 | 3.24E-29 | 7.54E-70 | -0.0114218 | 0.307 | 0.408 | 2.03E-65 |
| 287 | Dmxl1 | 1.36E-33 | -0.2 | 0.376 | 0.586 | 3.94E-29 | 2.97E-65 | 0.1516643 | 0.295 | 0.348 | 7.99E-61 |
| 288 | Sec14l5 | 1.48E-33 | -0.4 | 0.269 | 0.459 | 4.27E-29 | 2.49E-68 | -0.2203959 | 0.496 | 0.662 | 6.70E-64 |
| 289 | Fto | 1.69E-33 | -0.2 | 0.511 | 0.745 | 4.90E-29 | 2.56E-63 | -0.0468237 | 0.286 | 0.384 | 6.90E-59 |
| 290 | Slc38a2 | 1.99E-33 | -0.2 | 0.325 | 0.524 | 5.74E-29 | 9.36E-50 | -0.0409386 | 0.366 | 0.465 | 2.52E-45 |
| 291 | Kif1b | 2.11E-33 | -0.3 | 0.624 | 0.894 | 6.10E-29 | 1.26E-43 | -0.0726716 | 0.645 | 0.759 | 3.38E-39 |
| 292 | Sik2 | 2.47E-33 | -0.3 | 0.398 | 0.609 | 7.15E-29 | 6.77E-42 | -0.0664349 | 0.187 | 0.255 | 1.82E-37 |
| 293 | Tspan5 | 2.64E-33 | -0.3 | 0.54 | 0.764 | 7.62E-29 | 8.37E-51 | -0.1438505 | 0.324 | 0.441 | 2.25E-46 |
| 294 | Arhgef2 | 2.76E-33 | -0.2 | 0.306 | 0.506 | 7.97E-29 | 1.40E-50 | 0.0205994 | 0.195 | 0.254 | 3.76E-46 |
| 295 | Mtmr2 | 3.05E-33 | -0.3 | 0.314 | 0.507 | 8.82E-29 | 3.61E-53 | -0.0740815 | 0.239 | 0.326 | 9.71E-49 |
| 296 | Jmjd1c | 3.34E-33 | -0.2 | 0.58 | 0.83 | 9.65E-29 | 1.66E-55 | 0.0430379 | 0.266 | 0.338 | 4.46E-51 |
| 297 | Fth1 | 5.70E-33 | -0.3 | 0.591 | 0.81 | 1.65E-28 | 3.52E-50 | -0.0165005 | 0.419 | 0.522 | 9.49E-46 |
| 298 | Tom1l2 | 1.09E-32 | -0.2 | 0.463 | 0.693 | 3.14E-28 | 2.84E-52 | -0.0834183 | 0.28 | 0.378 | 7.65E-48 |
| 299 | Slc9a9 | 1.13E-32 | -0.3 | 0.496 | 0.723 | 3.26E-28 | 7.77E-46 | 0.0132959 | 0.237 | 0.301 | 2.09E-41 |
| 300 | Fbxo7 | 1.57E-32 | -0.3 | 0.257 | 0.45 | 4.53E-28 | 4.27E-53 | -0.0310127 | 0.191 | 0.262 | 1.15E-48 |
| 301 | Lrrc8b | 2.55E-32 | -0.2 | 0.303 | 0.505 | 7.36E-28 | 1.69E-52 | -0.0249802 | 0.211 | 0.284 | 4.56E-48 |
| 302 | Mat2a | 5.72E-32 | -0.2 | 0.379 | 0.586 | 1.65E-27 | 6.92E-75 | -0.1813845 | 0.313 | 0.457 | 1.86E-70 |
| 303 | Atf7ip | 6.68E-32 | -0.2 | 0.31 | 0.505 | 1.93E-27 | 1.02E-60 | -0.0943957 | 0.228 | 0.325 | 2.75E-56 |
| 304 | Add3 | 6.99E-32 | -0.2 | 0.521 | 0.747 | 2.02E-27 | 3.67E-56 | -0.2898469 | 0.241 | 0.372 | 9.88E-52 |
| 305 | Ank | 9.51E-32 | -0.2 | 0.361 | 0.572 | 2.75E-27 | 5.40E-57 | -0.0641012 | 0.249 | 0.344 | 1.45E-52 |
| 306 | Nfix | 9.97E-32 | -0.1 | 0.454 | 0.683 | 2.88E-27 | 8.01E-56 | -0.0422696 | 0.194 | 0.271 | 2.16E-51 |
| 307 | Bcl2l1 | 1.04E-31 | -0.2 | 0.294 | 0.486 | 3.02E-27 | 2.48E-65 | -0.1615715 | 0.339 | 0.473 | 6.67E-61 |
| 308 | Rbm5 | 1.37E-31 | -0.2 | 0.319 | 0.515 | 3.96E-27 | 4.64E-64 | 0.0230067 | 0.263 | 0.343 | 1.25E-59 |
| 309 | Stag1 | 1.45E-31 | -0.3 | 0.446 | 0.667 | 4.19E-27 | 2.93E-54 | -0.0516272 | 0.29 | 0.382 | 7.89E-50 |
| 310 | Dnajb2 | 1.53E-31 | -0.2 | 0.316 | 0.518 | 4.41E-27 | 9.01E-54 | 0.1344963 | 0.232 | 0.274 | 2.43E-49 |
| 311 | Cdc42se2 | 2.41E-31 | -0.2 | 0.345 | 0.542 | 6.97E-27 | 8.13E-52 | -0.0099733 | 0.229 | 0.3 | 2.19E-47 |
| 312 | Abl1 | 4.13E-31 | -0.3 | 0.287 | 0.479 | 1.19E-26 | 4.75E-56 | -0.087308 | 0.172 | 0.253 | 1.28E-51 |
| 313 | Dip2a | 4.53E-31 | -0.2 | 0.32 | 0.511 | 1.31E-26 | 7.85E-67 | -0.0985434 | 0.399 | 0.527 | 2.11E-62 |
| 314 | Itsn2 | 8.09E-31 | -0.2 | 0.379 | 0.585 | 2.34E-26 | 1.08E-46 | 0.0003961 | 0.449 | 0.541 | 2.91E-42 |
| 315 | Klf13 | 1.06E-30 | -0.2 | 0.259 | 0.442 | 3.08E-26 | 1.93E-74 | -0.1030116 | 0.288 | 0.411 | 5.19E-70 |
| 316 | Dixdc1 | 1.47E-30 | -0.2 | 0.367 | 0.566 | 4.24E-26 | 2.85E-66 | -0.0595883 | 0.282 | 0.388 | 7.66E-62 |
| 317 | Ubr2 | 2.21E-30 | -0.1 | 0.337 | 0.532 | 6.40E-26 | 2.34E-61 | 0.092395 | 0.255 | 0.318 | 6.29E-57 |
| 318 | Actn4 | 2.65E-30 | -0.2 | 0.343 | 0.543 | 7.66E-26 | 6.30E-61 | -0.227845 | 0.151 | 0.256 | 1.70E-56 |
| 319 | Mical3 | 2.74E-30 | -0.2 | 0.498 | 0.733 | 7.93E-26 | 9.18E-49 | -0.0376927 | 0.185 | 0.253 | 2.47E-44 |
| 320 | Fxr1 | 5.20E-30 | -0.2 | 0.335 | 0.529 | 1.50E-25 | 4.59E-54 | -0.0071166 | 0.211 | 0.279 | 1.24E-49 |
| 321 | Inf2 | 7.01E-30 | -0.2 | 0.263 | 0.453 | 2.03E-25 | 8.55E-55 | -0.0498494 | 0.182 | 0.258 | 2.30E-50 |
| 322 | Elovl7 | 9.95E-30 | -0.4 | 0.161 | 0.318 | 2.88E-25 | 3.97E-54 | -0.2583056 | 0.186 | 0.298 | 1.07E-49 |
| 323 | Ankrd12 | 1.21E-29 | -0.2 | 0.557 | 0.799 | 3.50E-25 | 3.08E-58 | 0.1041953 | 0.314 | 0.377 | 8.29E-54 |
| 324 | Wdfy3 | 1.30E-29 | -0.2 | 0.552 | 0.802 | 3.76E-25 | 3.60E-71 | 0.0442475 | 0.26 | 0.339 | 9.69E-67 |
| 325 | Mdm4 | 1.38E-29 | -0.2 | 0.349 | 0.56 | 3.98E-25 | 6.41E-57 | -0.0067945 | 0.197 | 0.268 | 1.73E-52 |
| 326 | Rock1 | 2.57E-29 | -0.2 | 0.322 | 0.508 | 7.44E-25 | 1.48E-67 | 0.0608789 | 0.226 | 0.294 | 3.98E-63 |
| 327 | Srpk2 | 3.44E-29 | -0.2 | 0.591 | 0.842 | 9.94E-25 | 2.63E-55 | -0.0114214 | 0.19 | 0.261 | 7.08E-51 |
| 328 | Fus | 4.08E-29 | -0.2 | 0.328 | 0.522 | 1.18E-24 | 3.41E-64 | -0.1297064 | 0.255 | 0.366 | 9.19E-60 |
| 329 | Rcan2 | 5.94E-29 | -0.2 | 0.398 | 0.594 | 1.72E-24 | 2.84E-81 | -0.3984821 | 0.288 | 0.471 | 7.64E-77 |
| 330 | Lgi3 | 6.25E-29 | -0.2 | 0.231 | 0.401 | 1.81E-24 | 8.58E-62 | 0.070743 | 0.234 | 0.298 | 2.31E-57 |
| 331 | Dis3l2 | 6.35E-29 | -0.2 | 0.371 | 0.562 | 1.83E-24 | 1.62E-45 | -0.1095222 | 0.256 | 0.35 | 4.35E-41 |
| 332 | Jakmip3 | 7.01E-29 | -0.3 | 0.221 | 0.385 | 2.03E-24 | 4.00E-57 | -0.148622 | 0.212 | 0.313 | 1.08E-52 |
| 333 | Litaf | 9.66E-29 | -0.3 | 0.158 | 0.31 | 2.79E-24 | 4.14E-70 | -0.1953274 | 0.276 | 0.41 | 1.11E-65 |
| 334 | Insr | 1.42E-28 | -0.2 | 0.416 | 0.63 | 4.10E-24 | 1.71E-53 | -0.0661959 | 0.186 | 0.264 | 4.60E-49 |
| 335 | Pan3 | 1.86E-28 | -0.2 | 0.39 | 0.594 | 5.36E-24 | 7.47E-51 | 0.0121216 | 0.21 | 0.275 | 2.01E-46 |

| | | | | | | | | | | | |
|---|---|---|---|---|---|---|---|---|---|---|---|
| 336 | Slc20a2 | 2.28E-28 | -0.2 | 0.358 | 0.561 | 6.59E-24 | 5.18E-52 | -0.1141975 | 0.226 | 0.318 | 1.40E-47 |
| 337 | Plxdc2 | 2.67E-28 | -0.3 | 0.616 | 0.832 | 7.71E-24 | 2.27E-40 | -0.0763909 | 0.696 | 0.803 | 6.11E-36 |
| 338 | Cadm2 | 2.67E-28 | -0.3 | 0.887 | 0.961 | 7.73E-24 | 3.81E-25 | 0.151705 | 0.714 | 0.751 | 1.03E-20 |
| 339 | Cdh20 | 3.53E-28 | -0.3 | 0.643 | 0.873 | 1.02E-23 | 1.05E-53 | -0.3217167 | 0.465 | 0.627 | 2.82E-49 |
| 340 | E130308A19Rik | 6.01E-28 | -0.2 | 0.377 | 0.57 | 1.74E-23 | 4.52E-51 | -0.0450111 | 0.258 | 0.341 | 1.22E-46 |
| 341 | Tmem178b | 8.75E-28 | 0.8 | 0.242 | 0.136 | 2.53E-23 | 1.62E-29 | -0.0861317 | 0.658 | 0.756 | 4.37E-25 |
| 342 | Exoc4 | 1.04E-27 | -0.2 | 0.502 | 0.716 | 3.00E-23 | 1.78E-65 | 0.0197964 | 0.38 | 0.479 | 4.80E-61 |
| 343 | Fign | 1.46E-27 | -0.3 | 0.38 | 0.563 | 4.22E-23 | 2.41E-43 | -0.015052 | 0.259 | 0.332 | 6.49E-39 |
| 344 | Mbnl1 | 1.92E-27 | -0.2 | 0.53 | 0.744 | 5.54E-23 | 6.44E-55 | -0.0484303 | 0.28 | 0.367 | 1.73E-50 |
| 345 | Cdkal1 | 2.68E-27 | -0.2 | 0.405 | 0.602 | 7.75E-23 | 1.80E-57 | -0.1555497 | 0.166 | 0.26 | 4.86E-53 |
| 346 | Robo1 | 3.06E-27 | -0.2 | 0.559 | 0.798 | 8.83E-23 | 2.63E-15 | -0.3386715 | 0.194 | 0.266 | 7.08E-11 |
| 347 | Baz2b | 4.56E-27 | -0.1 | 0.61 | 0.867 | 1.32E-22 | 3.50E-57 | -0.1940205 | 0.509 | 0.66 | 9.42E-53 |
| 348 | Fbxl5 | 8.48E-27 | -0.1 | 0.324 | 0.511 | 2.45E-22 | 3.17E-53 | 0.082093 | 0.329 | 0.387 | 8.53E-49 |
| 349 | Arid1b | 1.30E-26 | -0.2 | 0.602 | 0.837 | 3.75E-22 | 3.21E-60 | -0.0772159 | 0.257 | 0.358 | 8.63E-56 |
| 350 | Naaladl2 | 2.49E-26 | -0.3 | 0.451 | 0.641 | 7.21E-22 | 5.08E-39 | -0.0743638 | 0.438 | 0.546 | 1.37E-34 |
| 351 | Usp31 | 4.44E-26 | -0.2 | 0.286 | 0.457 | 1.28E-21 | 3.93E-60 | -0.3700107 | 0.158 | 0.286 | 1.06E-55 |
| 352 | Fam172a | 5.42E-26 | -0.2 | 0.495 | 0.718 | 1.57E-21 | 6.84E-57 | 0.012032 | 0.318 | 0.403 | 1.84E-52 |
| 353 | Myo18a | 7.92E-26 | -0.2 | 0.245 | 0.404 | 2.29E-21 | 2.84E-46 | 0.0209346 | 0.28 | 0.344 | 7.64E-42 |
| 354 | Ambra1 | 1.55E-25 | -0.1 | 0.39 | 0.584 | 4.48E-21 | 7.88E-60 | -0.0145543 | 0.257 | 0.343 | 2.12E-55 |
| 355 | Ephb1 | 4.04E-25 | 0.5 | 0.123 | 0.054 | 1.17E-20 | 1.44E-65 | -0.3167241 | 0.265 | 0.416 | 3.87E-61 |
| 356 | Wwox | 4.55E-25 | -0.3 | 0.653 | 0.86 | 1.31E-20 | 4.43E-30 | -0.1656445 | 0.26 | 0.348 | 1.19E-25 |
| 357 | Crebbp | 1.67E-24 | -0.2 | 0.345 | 0.528 | 4.82E-20 | 1.46E-66 | 0.1099107 | 0.27 | 0.333 | 3.94E-62 |
| 358 | Ppfibp2 | 1.67E-24 | -0.3 | 0.222 | 0.374 | 4.84E-20 | 1.09E-59 | -0.0863592 | 0.245 | 0.343 | 2.92E-55 |
| 359 | Lrp1b | 2.14E-24 | 0.7 | 0.782 | 0.773 | 6.18E-20 | 1.07E-08 | -0.0572349 | 0.961 | 0.979 | 0.000289259 |
| 360 | Lcorl | 2.52E-24 | -0.2 | 0.32 | 0.487 | 7.29E-20 | 5.21E-46 | 0.0907885 | 0.242 | 0.29 | 1.40E-41 |
| 361 | Cep97 | 2.64E-24 | -0.2 | 0.273 | 0.435 | 7.63E-20 | 1.13E-48 | 0.0581392 | 0.213 | 0.265 | 3.05E-44 |
| 362 | Ddx17 | 2.79E-24 | -0.2 | 0.664 | 0.922 | 8.08E-20 | 1.72E-71 | 0.0845891 | 0.378 | 0.46 | 4.62E-67 |
| 363 | Zcchc7 | 3.17E-24 | -0.2 | 0.397 | 0.577 | 9.18E-20 | 6.87E-65 | -0.1393419 | 0.402 | 0.542 | 1.85E-60 |
| 364 | Add1 | 3.99E-23 | -0.2 | 0.521 | 0.741 | 1.15E-18 | 1.86E-51 | -0.1152215 | 0.291 | 0.393 | 5.02E-47 |
| 365 | Stxbp6 | 5.77E-23 | -0.2 | 0.376 | 0.545 | 1.67E-18 | 2.30E-49 | -0.0669167 | 0.408 | 0.519 | 6.18E-45 |
| 366 | Atad2b | 5.93E-23 | -0.2 | 0.279 | 0.438 | 1.71E-18 | 2.73E-52 | -0.1049772 | 0.241 | 0.335 | 7.34E-48 |
| 367 | Prkcq | 1.02E-22 | -0.3 | 0.199 | 0.342 | 2.94E-18 | 5.44E-46 | -0.0519638 | 0.192 | 0.263 | 1.46E-41 |
| 368 | Vti1a | 1.05E-22 | -0.2 | 0.386 | 0.562 | 3.05E-18 | 4.43E-51 | -0.0059633 | 0.247 | 0.325 | 1.19E-46 |
| 369 | Stox2 | 1.86E-22 | -0.2 | 0.355 | 0.53 | 5.38E-18 | 1.08E-39 | 0.0423613 | 0.237 | 0.294 | 2.92E-35 |
| 370 | Eml6 | 3.90E-22 | 0.6 | 0.141 | 0.069 | 1.13E-17 | 1.88E-35 | 0.1020221 | 0.249 | 0.283 | 5.07E-31 |
| 371 | Slc25a27 | 9.19E-22 | -0.2 | 0.267 | 0.418 | 2.66E-17 | 9.93E-55 | -0.261771 | 0.218 | 0.338 | 2.67E-50 |
| 372 | Zdhhc14 | 2.36E-21 | -0.2 | 0.309 | 0.467 | 6.81E-17 | 8.28E-43 | -0.071398 | 0.459 | 0.561 | 2.23E-38 |
| 373 | Rsrc1 | 2.66E-21 | -0.1 | 0.296 | 0.456 | 7.70E-17 | 1.15E-53 | 0.03975 | 0.229 | 0.294 | 3.09E-49 |
| 374 | Ttll5 | 8.74E-21 | -0.2 | 0.246 | 0.391 | 2.52E-16 | 1.44E-58 | -0.077717 | 0.293 | 0.395 | 3.89E-54 |
| 375 | Kcnab1 | 1.31E-20 | -0.5 | 0.262 | 0.398 | 3.78E-16 | 2.91E-44 | -0.1457059 | 0.411 | 0.534 | 7.82E-40 |
| 376 | Tmcc1 | 1.63E-20 | -0.2 | 0.402 | 0.569 | 4.72E-16 | 6.23E-52 | -0.1294187 | 0.579 | 0.715 | 1.68E-47 |
| 377 | Rffl | 4.79E-20 | -0.2 | 0.132 | 0.249 | 1.39E-15 | 2.50E-72 | -0.1692788 | 0.275 | 0.407 | 6.72E-68 |
| 378 | Dmd | 1.32E-19 | -0.2 | 0.562 | 0.756 | 3.82E-15 | 2.20E-35 | 0.2064272 | 0.263 | 0.286 | 5.94E-31 |
| 379 | Daam1 | 4.58E-19 | 0.8 | 0.229 | 0.145 | 1.32E-14 | 3.71E-53 | 0.0718304 | 0.473 | 0.548 | 9.98E-49 |
| 380 | Mdga2 | 4.74E-19 | 0.6 | 0.339 | 0.233 | 1.37E-14 | 5.01E-25 | 0.0736486 | 0.722 | 0.782 | 1.35E-20 |
| 381 | Adgrb3 | 6.90E-19 | 0.5 | 0.799 | 0.848 | 1.99E-14 | 1.34E-35 | 0.4230345 | 0.331 | 0.305 | 3.59E-31 |
| 382 | St3gal4 | 1.90E-18 | -0.2 | 0.175 | 0.294 | 5.50E-14 | 7.24E-32 | -0.1834833 | 0.198 | 0.281 | 1.95E-27 |
| 383 | Negr1 | 4.25E-18 | 0.8 | 0.54 | 0.462 | 1.23E-13 | 3.59E-36 | 0.1822356 | 0.227 | 0.261 | 9.67E-32 |
| 384 | Heg1 | 1.08E-17 | -0.2 | 0.077 | 0.172 | 3.13E-13 | 4.99E-47 | -0.2290411 | 0.16 | 0.256 | 1.34E-42 |
| 385 | Camkmt | 1.89E-17 | -0.3 | 0.331 | 0.47 | 5.46E-13 | 2.17E-36 | -0.0492775 | 0.201 | 0.268 | 5.85E-32 |
| 386 | Dock3 | 4.95E-17 | -0.2 | 0.63 | 0.819 | 1.43E-12 | 2.90E-58 | 0.0087703 | 0.332 | 0.421 | 7.80E-54 |
| 387 | Ppp3ca | 5.66E-16 | 0.9 | 0.526 | 0.485 | 1.64E-11 | 1.67E-62 | 0.2574951 | 0.438 | 0.475 | 4.50E-58 |
| 388 | Dnajc1 | 6.02E-16 | -0.2 | 0.299 | 0.429 | 1.74E-11 | 3.88E-64 | 0.0202336 | 0.222 | 0.299 | 1.04E-59 |
| 389 | Dock4 | 3.12E-15 | -0.2 | 0.701 | 0.917 | 9.02E-11 | 3.73E-45 | 0.1577595 | 0.328 | 0.37 | 1.00E-40 |
| 390 | Phactr1 | 8.15E-15 | 0.8 | 0.439 | 0.365 | 2.36E-10 | 9.76E-47 | 0.0781071 | 0.545 | 0.628 | 2.63E-42 |
| 391 | Trim59 | 8.67E-15 | -0.2 | 0.09 | 0.177 | 2.51E-10 | 3.28E-51 | -0.0188524 | 0.202 | 0.275 | 8.83E-47 |
| 392 | Syt11 | 2.01E-13 | 0.2 | 0.461 | 0.62 | 5.81E-09 | 6.32E-61 | -0.0054171 | 0.315 | 0.406 | 1.70E-56 |
| 393 | Apoe | 3.34E-13 | 0.8 | 0.141 | 0.086 | 9.65E-09 | 4.26E-68 | 0.144747 | 0.448 | 0.539 | 1.15E-63 |
| 394 | Itm2b | 3.38E-13 | 0.3 | 0.386 | 0.546 | 9.77E-09 | 4.46E-74 | -0.0378801 | 0.222 | 0.322 | 1.20E-69 |
| 395 | Son | 1.45E-12 | 0.1 | 0.474 | 0.645 | 4.19E-08 | 8.16E-65 | 0.0074378 | 0.403 | 0.506 | 2.20E-60 |
| 396 | Nrxn3 | 3.24E-12 | -0.2 | 0.778 | 0.888 | 9.38E-08 | 4.20E-149 | 1.2176007 | 0.31 | 0.129 | 1.13E-144 |
| 397 | Mast4 | 1.21E-11 | 0.7 | 0.304 | 0.237 | 3.51E-07 | 3.16E-10 | -0.0789196 | 0.919 | 0.952 | 8.50E-06 |
| 398 | Immp1l | 3.41E-11 | -0.1 | 0.154 | 0.238 | 9.85E-07 | 3.72E-52 | -0.0491158 | 0.23 | 0.312 | 1.00E-47 |
| 399 | Atp2b1 | 1.35E-10 | 1.1 | 0.357 | 0.31 | 3.91E-06 | 2.89E-78 | 0.2972586 | 0.261 | 0.294 | 7.78E-74 |
| 400 | Samd4 | 2.25E-10 | 0.7 | 0.216 | 0.157 | 6.50E-06 | 8.22E-41 | 0.0774302 | 0.24 | 0.289 | 2.21E-36 |
| 401 | Herc1 | 3.43E-10 | 0.2 | 0.5 | 0.676 | 9.91E-06 | 6.63E-62 | 0.1331456 | 0.225 | 0.275 | 1.78E-57 |
| 402 | Prrc2c | 1.56E-09 | 0.2 | 0.502 | 0.678 | 4.50E-05 | 5.51E-81 | -0.0627594 | 0.246 | 0.359 | 1.48E-76 |
| 403 | Csmd3 | 2.28E-09 | 0.8 | 0.389 | 0.334 | 6.60E-05 | 5.40E-25 | -0.1161065 | 0.774 | 0.86 | 1.45E-20 |
| 404 | Slc6a1 | 2.49E-09 | 0.2 | 0.39 | 0.517 | 7.19E-05 | 4.66E-59 | -0.0891115 | 0.189 | 0.277 | 1.26E-54 |
| 405 | Ppp1r12a | 3.58E-09 | 0.1 | 0.361 | 0.485 | 0.000103325 | 1.48E-66 | 0.0674207 | 0.277 | 0.35 | 4.00E-62 |
| 406 | Xrcc6 | 7.73E-09 | 0.2 | 0.13 | 0.203 | 0.000223489 | 3.98E-51 | -0.0067452 | 0.322 | 0.409 | 1.07E-46 |
| 407 | Nkain1 | 9.35E-09 | -0.2 | 0.11 | 0.174 | 0.000270182 | 1.64E-67 | -0.0833248 | 0.267 | 0.373 | 4.43E-63 |
| 408 | Tmod2 | 1.35E-08 | 0.2 | 0.299 | 0.409 | 0.000391502 | 2.95E-47 | -0.16705 | 0.529 | 0.666 | 7.94E-43 |
| 409 | Srek1 | 1.86E-08 | 0.2 | 0.264 | 0.357 | 0.000538945 | 8.71E-69 | 0.0470544 | 0.207 | 0.28 | 2.34E-64 |
| 410 | Atp6v0b | 3.71E-08 | 0.2 | 0.166 | 0.245 | 0.00107168 | 5.42E-65 | 0.0921348 | 0.27 | 0.341 | 1.46E-60 |
| 411 | Dcc | 5.74E-08 | 0.3 | 0.283 | 0.222 | 0.001658474 | 4.72E-25 | 0.1532658 | 0.37 | 0.4 | 1.27E-20 |
| 412 | Epb41l3 | 5.77E-08 | 0.2 | 0.244 | 0.34 | 0.001666744 | 5.24E-66 | 0.0679267 | 0.339 | 0.422 | 1.41E-61 |
| 413 | Atrx | 2.33E-07 | 0.2 | 0.53 | 0.707 | 0.006740378 | 3.57E-68 | 0.0355634 | 0.295 | 0.384 | 9.62E-64 |
| 414 | Dst | 2.82E-07 | 0.3 | 0.821 | 0.978 | 0.008139287 | 6.93E-59 | 0.0091118 | 0.589 | 0.7 | 1.87E-54 |
| 415 | Rasal2 | 1.27E-06 | 0.5 | 0.264 | 0.214 | 0.036733369 | 9.17E-43 | -0.0177002 | 0.206 | 0.275 | 2.47E-38 |
| 416 | Nckap1 | 1.95E-06 | 0.2 | 0.326 | 0.427 | 0.056490119 | 1.51E-54 | -0.0016965 | 0.265 | 0.342 | 4.07E-50 |
| 417 | Dpysl2 | 7.92E-06 | 0.2 | 0.617 | 0.786 | 0.228766601 | 2.32E-72 | -0.07877 | 0.336 | 0.456 | 6.24E-68 |
| 418 | Macf1 | 8.88E-06 | 0.8 | 0.747 | 0.892 | 0.256576179 | 2.46E-56 | 0.0384003 | 0.377 | 0.464 | 6.63E-52 |
| 419 | Myo9a | 1.03E-05 | 0.2 | 0.411 | 0.516 | 0.297020978 | 6.49E-59 | 0.1352156 | 0.319 | 0.375 | 1.75E-54 |
| 420 | Prickle2 | 1.03E-05 | 0.5 | 0.151 | 0.113 | 0.297511815 | 3.84E-24 | -0.1148606 | 0.712 | 0.805 | 1.03E-19 |
| 421 | Apc | 1.92E-05 | 0.3 | 0.362 | 0.454 | 0.555473394 | 1.03E-58 | 0.0725347 | 0.38 | 0.453 | 2.77E-54 |
| 422 | Akap8l | 1.92E-05 | 0.2 | 0.217 | 0.289 | 0.556032318 | 2.31E-54 | -0.0626281 | 0.225 | 0.312 | 6.21E-50 |
| 423 | Scamp5 | 2.16E-05 | 0.6 | 0.184 | 0.146 | 0.623350733 | 3.26E-62 | -0.0807366 | 0.184 | 0.272 | 8.78E-58 |
| 424 | Bicd1 | 2.61E-05 | 0.2 | 0.414 | 0.528 | 0.754958418 | 4.11E-53 | -0.0749154 | 0.27 | 0.365 | 1.11E-48 |
| 425 | Adam22 | 3.72E-05 | 0.2 | 0.292 | 0.376 | 1 | 6.21E-51 | 0.0175324 | 0.221 | 0.289 | 1.67E-46 |
| 426 | Pak7 | 5.14E-05 | 0.4 | 0.121 | 0.089 | 1 | 1.14E-32 | -0.0239021 | 0.642 | 0.734 | 3.08E-28 |
| 427 | Thra | 5.33E-05 | 0.2 | 0.231 | 0.297 | 1 | 2.81E-68 | 0.1781811 | 0.284 | 0.332 | 7.55E-64 |
| 428 | Gria2 | 6.20E-05 | 0.7 | 0.562 | 0.598 | 1 | 2.18E-115 | 0.7917753 | 0.353 | 0.276 | 5.87E-111 |
| 429 | Nfat5 | 7.92E-05 | 0.3 | 0.437 | 0.556 | 1 | 5.69E-46 | 0.0839112 | 0.235 | 0.288 | 1.53E-41 |
| 430 | Plekha5 | 0.000183396 | 0.6 | 0.28 | 0.244 | 1 | 2.27E-62 | 0.182853 | 0.272 | 0.317 | 6.11E-58 |
| 431 | Gria4 | 0.000188891 | 0.7 | 0.365 | 0.338 | 1 | 2.71E-31 | 0.2254313 | 0.32 | 0.334 | 7.30E-27 |
| 432 | Kif21a | 0.000453827 | 0.3 | 0.32 | 0.403 | 1 | 6.91E-45 | 0.0412293 | 0.558 | 0.64 | 1.86E-40 |
| 433 | Cpeb3 | 0.000464572 | 0.3 | 0.365 | 0.452 | 1 | 1.53E-56 | -0.0306414 | 0.247 | 0.331 | 4.13E-52 |
| 434 | Dennd5b | 0.000493588 | 0.3 | 0.352 | 0.436 | 1 | 3.98E-58 | -0.1490826 | 0.236 | 0.344 | 1.07E-53 |
| 435 | Psd3 | 0.001291554 | 0.2 | 0.592 | 0.736 | 1 | 1.56E-43 | 0.0975906 | 0.264 | 0.312 | 4.21E-39 |
| 436 | Strbp | 0.002245462 | 0.5 | 0.199 | 0.169 | 1 | 1.41E-35 | -0.0376747 | 0.387 | 0.474 | 3.79E-31 |
| 437 | Mycbp2 | 0.002716682 | 0.3 | 0.524 | 0.648 | 1 | 3.56E-54 | 0.0558312 | 0.579 | 0.67 | 9.60E-50 |
| 438 | Pde4dip | 0.003621102 | 0.6 | 0.504 | 0.553 | 1 | 1.88E-55 | 0.1014016 | 0.198 | 0.25 | 5.05E-51 |
| 439 | Lrrc4c | 0.003664658 | 0.6 | 0.521 | 0.549 | 1 | 2.98E-25 | -0.1566244 | 0.644 | 0.745 | 8.02E-21 |
| 440 | Arnt2 | 0.003955709 | 0.3 | 0.373 | 0.448 | 1 | 7.33E-58 | -0.025446 | 0.255 | 0.337 | 1.97E-53 |
| 441 | Gna12 | 0.004020717 | 0.3 | 0.166 | 0.208 | 1 | 1.36E-62 | -0.1375866 | 0.252 | 0.364 | 3.65E-58 |
| 442 | Dpy19l1 | 0.004608304 | 0.1 | 0.141 | 0.176 | 1 | 3.54E-66 | -0.0372403 | 0.202 | 0.286 | 9.52E-62 |
| 443 | Rab31 | 0.006060564 | 0.4 | 0.101 | 0.081 | 1 | 2.74E-76 | -0.3169854 | 0.486 | 0.675 | 7.38E-72 |
| 444 | Nbea | 0.011780279 | 0.6 | 0.501 | 0.539 | 1 | 4.57E-42 | 0.0049072 | 0.623 | 0.721 | 1.23E-37 |
| 445 | Rev3l | 0.012153702 | 0.4 | 0.395 | 0.479 | 1 | 1.51E-41 | 0.0365558 | 0.223 | 0.276 | 4.07E-37 |
| 446 | Inpp5f | 0.043917101 | 0.6 | 0.201 | 0.189 | 1 | 4.26E-65 | -0.1532861 | 0.222 | 0.332 | 1.15E-60 |
| 447 | Pacrg | 0.081333765 | 0.4 | 0.138 | 0.125 | 1 | 1.28E-67 | -0.4013012 | 0.502 | 0.687 | 3.45E-63 |
| 448 | Grid2 | 0.088684838 | 0.2 | 0.395 | 0.38 | 1 | 2.44E-45 | -0.0409379 | 0.565 | 0.681 | 6.56E-41 |
| 449 | Erbb4 | 0.139637368 | 0.2 | 0.746 | 0.837 | 1 | 8.02E-13 | -0.0157006 | 0.761 | 0.807 | 2.16E-08 |
| 450 | Prkca | 0.147053462 | 0.5 | 0.477 | 0.508 | 1 | 3.28E-55 | -0.1111533 | 0.329 | 0.448 | 8.83E-51 |
| 451 | Dgki | 0.164506696 | 0.4 | 0.282 | 0.319 | 1 | 5.13E-40 | -0.2001188 | 0.518 | 0.65 | 1.38E-35 |
| 452 | Bmp2k | 0.184356762 | 0.4 | 0.229 | 0.259 | 1 | 1.19E-62 | -0.0419164 | 0.181 | 0.261 | 3.21E-58 |
| 453 | Fbxl17 | 0.38052402 | 0.4 | 0.416 | 0.446 | 1 | 5.52E-48 | 0.0428832 | 0.257 | 0.32 | 1.49E-43 |
| 454 | Ablim2 | 0.482052675 | 0.3 | 0.108 | 0.105 | 1 | 6.20E-61 | 0.3586125 | 0.323 | 0.315 | 1.67E-56 |
| 455 | R3hdm1 | 0.504402627 | 0.5 | 0.45 | 0.496 | 1 | 1.15E-58 | 0.1433469 | 0.376 | 0.432 | 3.09E-54 |
| 456 | Cdh2 | 0.518843529 | 0.2 | 0.29 | 0.315 | 1 | 8.07E-62 | -0.0544175 | 0.344 | 0.452 | 2.17E-57 |
| 457 | Rtn3 | 0.552728592 | 0.7 | 0.555 | 0.653 | 1 | 1.06E-57 | 0.0233924 | 0.269 | 0.347 | 2.84E-53 |
| 458 | Ehbp1 | 0.591393115 | 0.4 | 0.375 | 0.392 | 1 | 9.34E-51 | 0.0997952 | 0.229 | 0.281 | 2.51E-46 |
| 459 | Arhgef12 | 0.637001747 | 0.4 | 0.325 | 0.347 | 1 | 4.92E-54 | -0.0109288 | 0.218 | 0.293 | 1.32E-49 |
| 460 | Kazn | 0.684073826 | 0.3 | 0.624 | 0.705 | 1 | 2.03E-59 | -0.0453059 | 0.216 | 0.298 | 5.46E-55 |
| 461 | Aplp2 | 0.828195718 | 0.7 | 0.196 | 0.212 | 1 | 2.99E-77 | -0.0945446 | 0.191 | 0.294 | 8.04E-73 |
| 462 | Sez6l2 | 0.877774532 | 0.4 | 0.132 | 0.135 | 1 | 1.86E-68 | -0.2455685 | 0.271 | 0.414 | 5.02E-64 |
