## Supplementary table 8 for "DAP12 deficiency alters microglia-oligodendrocyte communication and enhances resilience against tau toxicity"

### snRNAseq

| n sample | brain bank | brain bank ID | Dx | ClinicalDx | TREM2 | Braak stage | APOE | Age | Age of onset | Sex | Froz | Mid Frontal Ctx | PMI |
|---|---|---|---|---|---|---|---|---|---|---|---|---|---|
| 43 | Upenn | 108953 | AD | Dementia with Lewy Bodies | WT | 6 | E3/E3 | 83 | 77 | F | Yes | Yes | 4 |
| 44 | Upenn | 105007 | AD | Alzheimer's Disease Probable | WT | 6 | E3/E3 | 87 | 78 | F | Yes | Yes | 16 |
| 45 | Upenn | 102475 | AD | Probable Alzheimer's Disease | WT | 5 | E4/E4 | 72 | 66 | F | Yes | Yes | 5 |
| 46 | Upenn | 107928 | AD | Dementia of undetermined etiology | WT | 5 | E3E3 | 82 | 74 | M | Yes | Yes | 39 |
| 47 | Upenn | 122417 | AD | Probable Alzheimer's Disease | WT | 6 | E3/E4 | 64 | 60 | M | Yes | Yes | 6 |
| 48 | Upenn | 111593 | AD | Alzheimer's Disease Probable | WT | 6 | E3/E4 | 71 | 62 | M | Yes | Yes | 4.5 |
| 49 | Upenn | 107833 | AD | Alzheimer's Disease Probable | WT | 6 | E3/E4 | 78 | 64 | M | Yes | Yes | 6.5 |
| 50 | Upenn | 100436 | AD | Probable Alzheimer's Disease | WT | 6 | E4/E4 | 78 | 70 | M | Yes | Yes | 6 |
| 59 | Upenn | 119534 | Normal | | | | E2/E3 | 72 | | F | Yes | Yes | 6 |
| 60 | Upenn | 112090 | Normal | | | | E2/E3 | 83 | | F | Yes | Yes | 3 |
| 61 | Upenn | 125061 | Normal | | | | NA | 75 | | F | Yes | Yes | 12 |
| 62 | Upenn | 118709 | Normal | | | | E2/E3 | 68 | | M | Yes | Yes | 14 |
| 63 | Upenn | 120927 | Normal | | | | E2/E3 | 72 | | M | Yes | Yes | 17 |
| 64 | Upenn | 100786 | Normal | | | | E2/E4 | 61 | | M | Yes | Yes | 6 |
| 65 | Upenn | 113464 | Normal | | | | E3/E3 | 75 | | M | Yes | Yes | 17 |
| 66 | Upenn | 119767 | Normal | | | | E3/E3 | 83 | | M | Yes | Yes | 6 |

### Immunohistochemistry

lount saini

| Barcode | Sub Num | StorageType | Brain Region | UoM | Size | Age | Sex | Race | PMI min. | DX | B&B Alz | ApoE |
|---|---|---|---|---|---|---|---|---|---|---|---|---|
| 261539 | 11539 | Fixed Slide | BM-9 | ea | 1 | 97 | Female | White | 140 | Contro | 2 | 3-Feb |
| 261440 | 755 | Fixed Slide | BM-9 | ea | 1 | 102 | Female | White | 423 | Contro | 5 | 3-Mar |
| 261879 | 50580 | Fixed Slide | BM-9 | ea | 1 | 80 | Female | White | 285 | Contro | 2 | 3-Feb |
| 261958 | 108267 | Fixed Slide | BM-9 | ea | 1 | 54 | Male | White | 1185 | Contro | 9 | 3-Mar |
| 261422 | 10795 | Fixed Slide | BM-9 | ea | 1 | 79 | Female | Hispanic | 460 | Contro | 3 | 3-Feb |
| 261296 | 81898 | Fixed Slide | BM-9 | ea | 1 | 94 | Female | White | 260 | AD | 6 | 4-Mar |
| 262039 | 1119 | Fixed Slide | BM-9 | ea | 1 | 91 | Female | White | 160 | AD | 6 | 3-Mar |
| 261787 | 82158 | Fixed Slide | BM-9 | ea | 1 | 86 | Female | White | 185 | AD | 6 | 4-Mar |
| 261843 | 47108 | Fixed Slide | BM-9 | ea | 1 | 90 | Female | White | 380 | AD | 6 | 4-Mar |
| 261483 | 53593 | Fixed Slide | BM-9 | ea | 1 | 70 | Female | White | 1605 | AD | 0 | 4-Mar |

Uppen

| INDDID | Sex | Age | ge Ons | PMI | Braak06 | NPDx1 | APOE | INDDID | ClinicalDx1 |
|---|---|---|---|---|---|---|---|---|---|
| 107928 | Male | 82 | 74 | 39 | 5 | AD | E3/E3 | 100786 | Dementia of undetermined etiology |
| 122417 | Male | 64 | 60 | 6 | 6 | AD | E3/E4 | 118709 | Probable Alzheimer's Disease |
| 111593 | Male | 71 | 62 | 4.5 | 6 | AD | E3/E4 | 120927 | Alzheimer's Disease Probable |
| 107833 | Male | 78 | 64 | 6.5 | 6 | AD | E3/E4 | 113464 | Alzheimer's Disease Probable |
| 100436 | Male | 78 | 70 | 6 | 6 | AD | E4/E4 | 119767 | Probable Alzheimer's Disease |
| 118709 | Male | 68 | NA | 14 | 0 | Normal | E2/E3 | 100957 | |
| 120927 | Male | 72 | NA | 17 | 1 | Normal | E2/E3 | 105703 | |
| 100786 | Male | 61 | NA | 6 | 0 | Normal | E2/E4 | 107970 | |
| 113464 | Male | 75 | NA | 17 | 1 | Normal | E3/E3 | 107342 | |
| 119767 | Male | 83 | NA | 6 | 2 | Normal | E3/E3 | 112403 | |
